## Supplementary material for "Biosynthesis and Bioactivity of Anti-Inflammatory Triterpenoids in *Calendula officinalis* (pot marigold)": Combined Supplementary Data

**Supplementary Materials**

Supplementary Figure 1: Metabolite analysis of *Calendula officinalis* by GC-MS.
Supplementary Figure 2: Metabolite analysis of *Calendula officinalis* by LC-MS.
Supplementary Figure 3: GC-MS standards and mass spectra for triterpene monols.
Supplementary Figure 4: GC-MS standards and mass spectra for triterpene diols and acids.
Supplementary Figure 5: GC-MS standards and mass spectra for faradiol fatty acid esters.
Supplementary Figure 6: Semi-preparative uHPLC chromatograms of the methanol extract of *Calendula officinalis* ray florets.
Supplementary Figure 7: GC-MS analysis of compounds purified from *C. officinalis* ray floret extracts.
Supplementary Figure 8: GC-MS analysis of *Calendula officinalis* ray floret fractions.
Supplementary Figure 9: Structure of 3-*O*-palmitoyl faradiol (faradiol palmitate).
Supplementary Figure 10: $^{1}$H NMR spectra of faradiol palmitate.
Supplementary Figure 11: $^{13}$C NMR spectra of faradiol palmitate.
Supplementary Figure 12: $^{1}$H-$^{1}$H COSY NMR (600 MHz, CDCl3, 298 K) of faradiol palmitate.
Supplementary Figure 13: $^{1}$H-$^{13}$C-HSQC-edited NMR (600 MHz, CDCl3, 298 K) of faradiol palmitate.
Supplementary Figure 14: $^{1}$H-$^{13}$C HMBC NMR (600 MHz, CDCl3, 298 K) of faradiol palmitate.
Supplementary Figure 15: Effects of *C. officinalis* extracts and triterpenoids on the viability of human monocytic (THP-1) cells.
Supplementary Figure 16. Concentration-dependent responses to floral extracts and faradiol.
Supplementary Figure 17: Phylogenetic analysis of plant oxidosqualene cyclases.
Supplementary Figure 18: Chromosomal location of *Calendula officinalis* oxidosqualene cyclases.
Supplementary Figure 19: Differential expression of *Calendula officinalis* oxidosqualene cyclases candidate genes.
Supplementary Figure 20: GC-MS analysis of *N. benthamiana* leaf extracts infiltrated with constructs expressing oxidosqualene cyclases.
Supplementary Figure 21: Bayesian phylogenetic analysis of oxidosqualene cyclases .
Supplementary Figure 22: GC-MS profile of *Calendula officinalis* tissues .
Supplementary Figure 23: GM-MS profile of *Taraxacum kok-saghyz* tissues.
Supplementary Figure 24: GC-MS analysis (total ion chromatograms) of *N. benthamiana* leaf extracts infiltrated expressing wild type and mutant taraxasterol synthases .
Supplementary Figure 25: Phylogenetic analysis of cytochrome p450s.
Supplementary Figure 26: Differential expression of *Calendula officinalis* cytochrome p450 candidate genes.
Supplementary Figure 27. Genomic location and synteny of *Calendula officinalis* (pot marigold) cytochrome P450 genes encoding CoCYP716A392 and CoCYP716A393.
Supplementary Figure 28:GC-MS analysis of *Nicotiana benthamiana* expressing cytochrome P450s (CYPs).
Supplementary Figure 29: GC-MS profiling of floral extracts from Asteraceae species.
Supplementary Figure 30: GC-MS analysis of *Nicotiana benthamiana* expressing cytochrome P450 mutants.
Supplementary Figure 31. Maximum-likelihood tree of plant acyltransferases.
Supplementary Figure 32: ACT Diff gene expression.
Supplementary Figure 33: Characterisation of pot marigold triterpene acyl transferases.
Supplementary Figure 34: Characterisation of hydroxylated triterpene acyl transferases.
Supplementary Figure 35: Genomic location and synteny of *Calendula officinalis* (pot marigold) acyltransferases .
Supplementary Table 1: Triterpenes detected in *Calendula officinalis.*
Supplementary Table 2: NMR data for faradiol palmitate .
Supplementary Table 3: Comparison of experimental and literature assignment of NMR data for faradiol palmitate.
Supplementary Table 4: Tables of statistics for Figure 1.
Supplementary Table 5: Table of genome assembly statistics.

Supplementary Table 6: Table of candidate *oxidosqualene cyclase* genes (*OSCs*) identified in the *Calendula officinalis* genome.
Supplementary Table 7: Table of *Calendula officinalis* oxidosqualene cycle transcripts.
Supplementary Table 8: Table of coding and protein sequences of *Calendula officinalis* oxidosqualene cyclases.
Supplementary Table 9: Table of all oxidosqualene cyclases with accession numbers.
Supplementary Table 10: Table of mixed amyrin synthases and taraxasterol synthases.
Supplementary Table 11: Statistical tests used in Figure 3.
Supplementary Table 12: Statistical tests used in Figure 4.
Supplementary Table 13: Statistical tests used in Figure 5.
Supplementary Table 14: Statistical tests used in Figure 6.
Supplementary Table 15: List of plasmids used in this study.
Supplementary Table 16: List of primers used in this study.
Extended Methods.

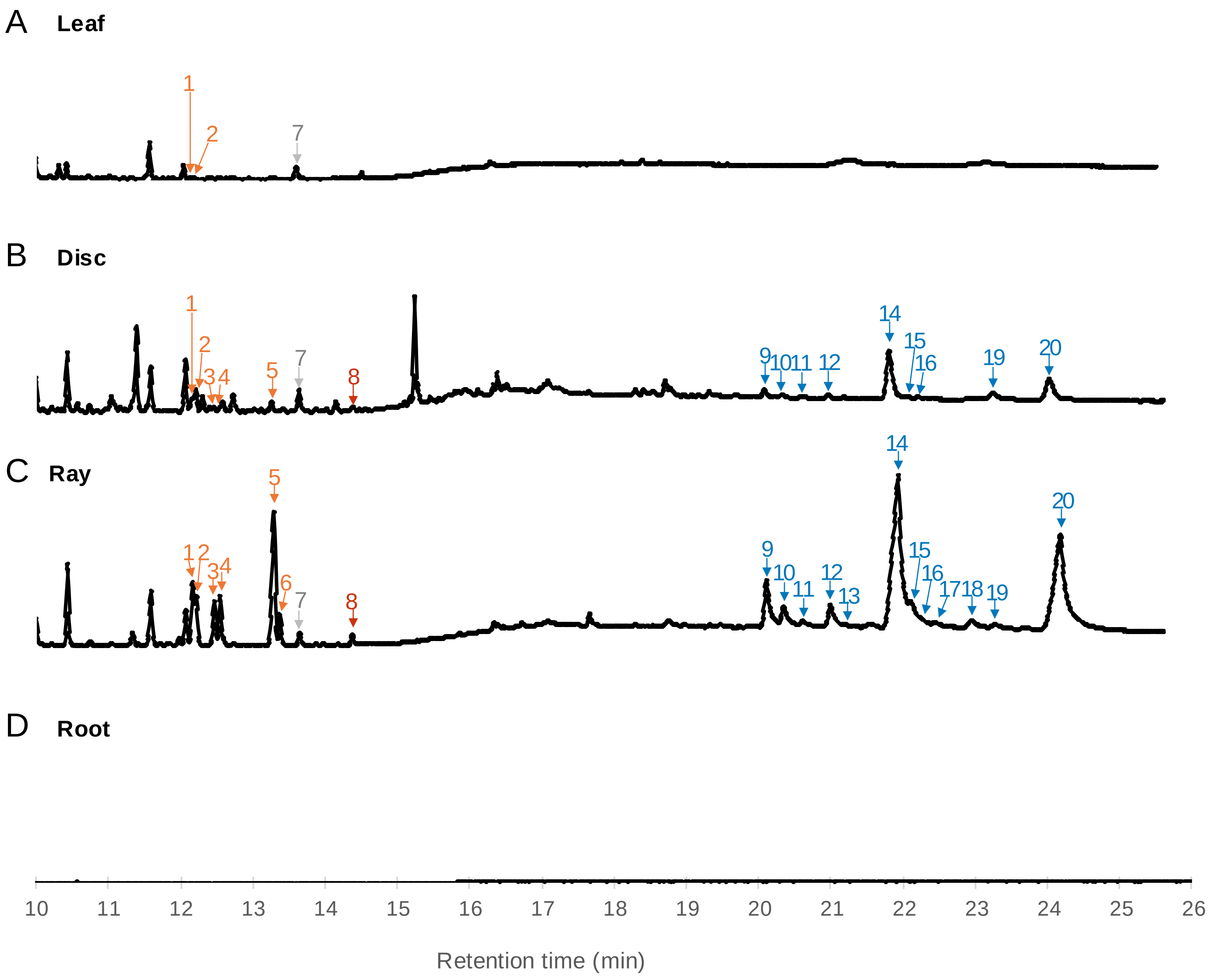

**Triterpene monols**

1. β-amyrin
2. isofucosterol
3. α-amyrin
4. lupeol
5. ψ-taraxasterol
6. taraxasterol

7. friedelin (internal standard)

**Triterpene diols**

8. faradiol

**Triterpene fatty acid esters**

9. faradiol/arnidiol laurate
10. β-amyrin myristate
11. manaladiol myristate
12. calenduladiol myristate
13. ψ-taraxasterol/taraxasterol myristate
14. faradiol/arnidiol myristate
15. β-amyrin palmitate
16. lupeol palmitate
17. manaladiol palmitate
18. calenduladiol palmitate
19. ψ-taraxasterol/taraxasterol palmitate
20. faradiol/arnidiol palmitate

**Supplementary Figure 1. Metabolite analysis of *Calendula officinalis* by GC-MS. A.** Total Ion Chromatogram (TIC) of derivatised leaf extract. **B.** TIC of derivatised disc floret extract. **C.** TIC of derivatised ray floret extract. **D.** TIC of derivatised root extract. Labelled peaks = triterpene monols (orange), triterpene diols (red), triterpene fatty acid esters (blue) and internal standard (grey).

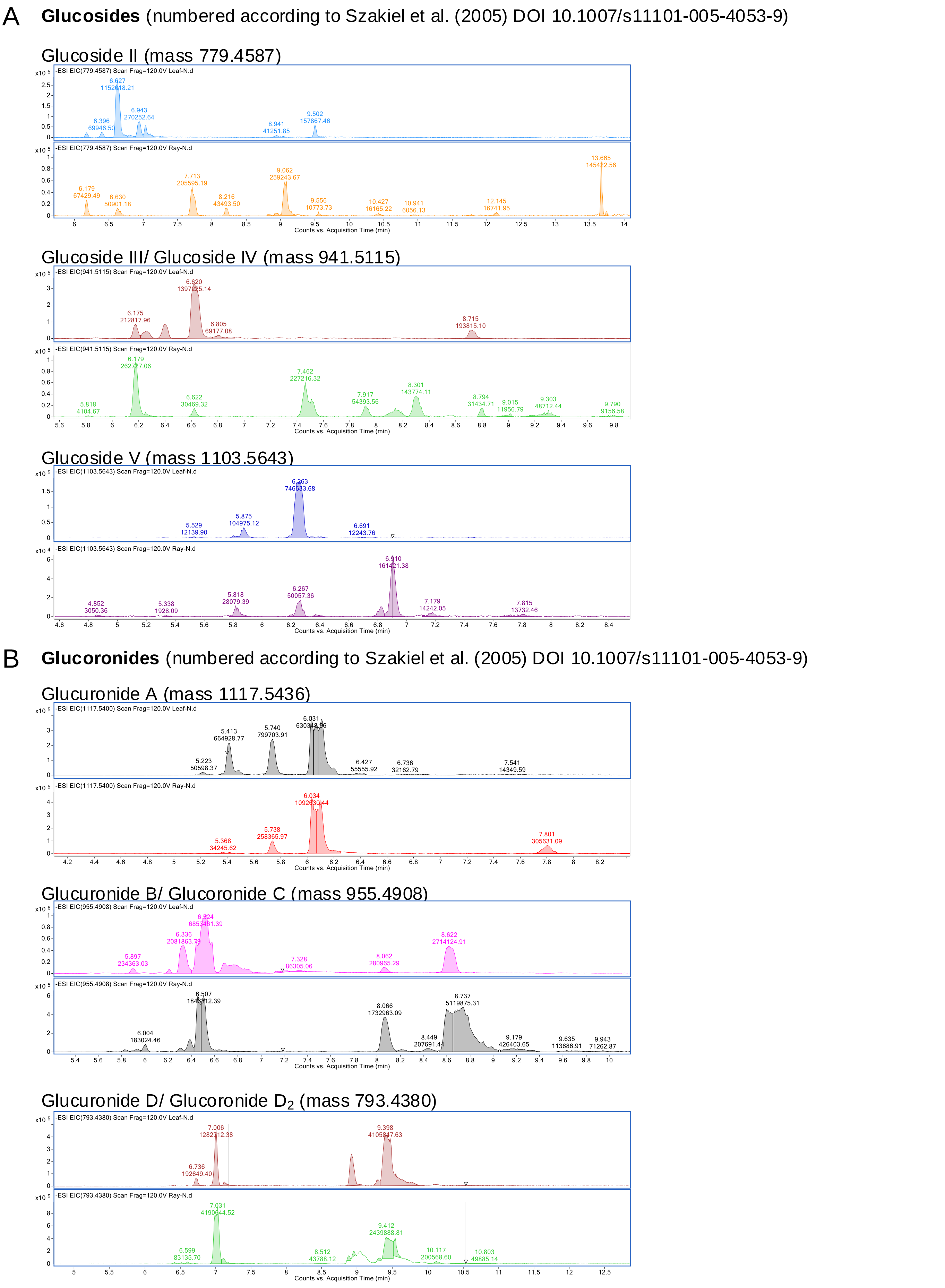

**Supplementary Figure 2. Metabolite analysis of Calendula officinalis by LC-MS**
**A.** Extracted ion chromatograms (EIC) of *Calendula officinalis* leaf and ray extracts corresponding to masses of oleanolic acid glucosides. **B.** Extracted ion chromatograms (EIC) of *Calendula officinalis* leaf and ray extracts corresponding to masses of oleanolic acid glucuronides. Compounds are numbered according to Szakiel et al. (2005) DOI 10.1007/s11101-005-4053-9.

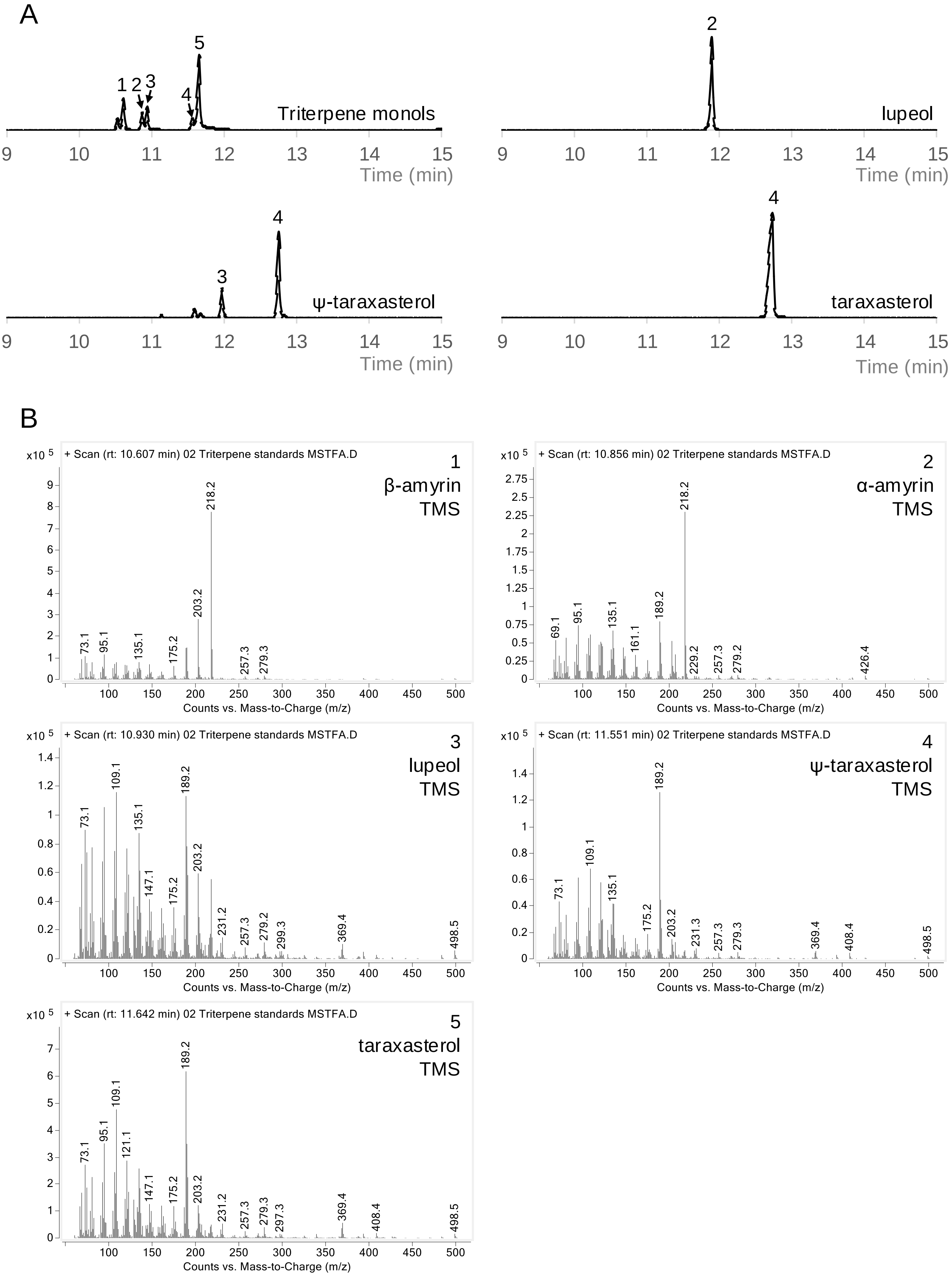

**Supplementary Figure 3. GC-MS standards and mass spectra for triterpene monols. A.** GC-MS chromatograms of trimethylsilyl (TMS) derivatised triterpene monols. Triterpene monols is a mixture of 1. β-amyrin, 2. α-amyrin, 3. lupeol, 4. ψ-taraxasterol, 5. taraxasterol. β-amyrin, α-amyrin and lupeol were kindly provided by Anne Osbourn, John Innes Centre, Norwich, UK; taraxasterol were purchased from Merck, Darmstadt, Germany (PHL84272-10MG). ψ-taraxasterol was purified from *Calendula officinalis* (Supplementary Figures 6-8) **B.** Mass spectra for each of the TMS derivatised triterpene monols.

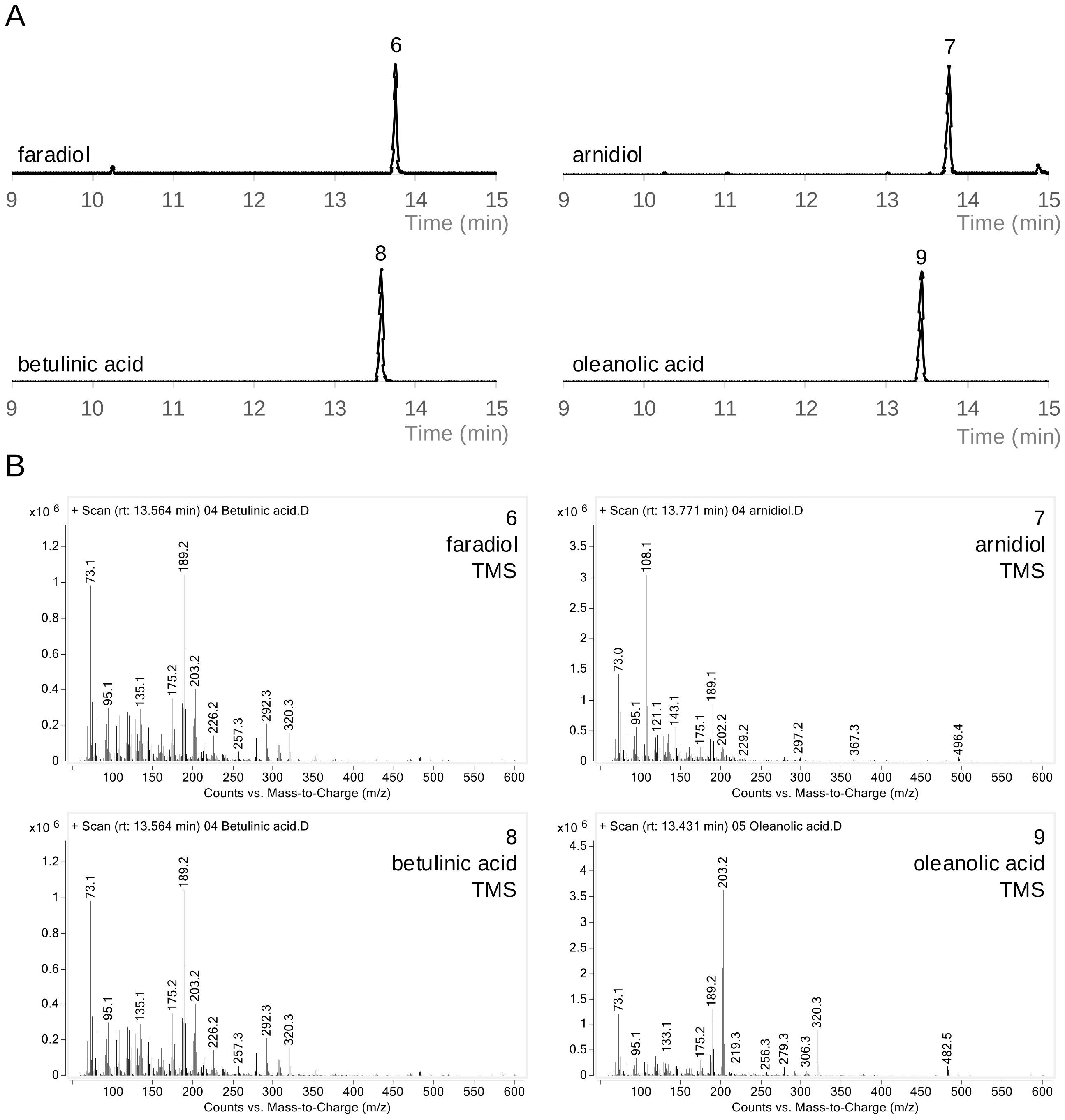

**Supplementary Figure 4. GC-MS standards and mass spectra for triterpene diols and acids. A.** GC-MS chromatograms of TMS derivatised triterpene diol and acids. Faradiol, betulinic acid and oleanolic acid were purchased from Merck, Darmstadt, Germany (PHL82536-10MG, 91466-10MG and 42515-10MG respectively). Arnidiol was purchased from MedChem Express, New Jersey, USA (HY-N4165-1mg) **B.** Mass spectra for each of the TMS derivatised triterpene diols and acids.

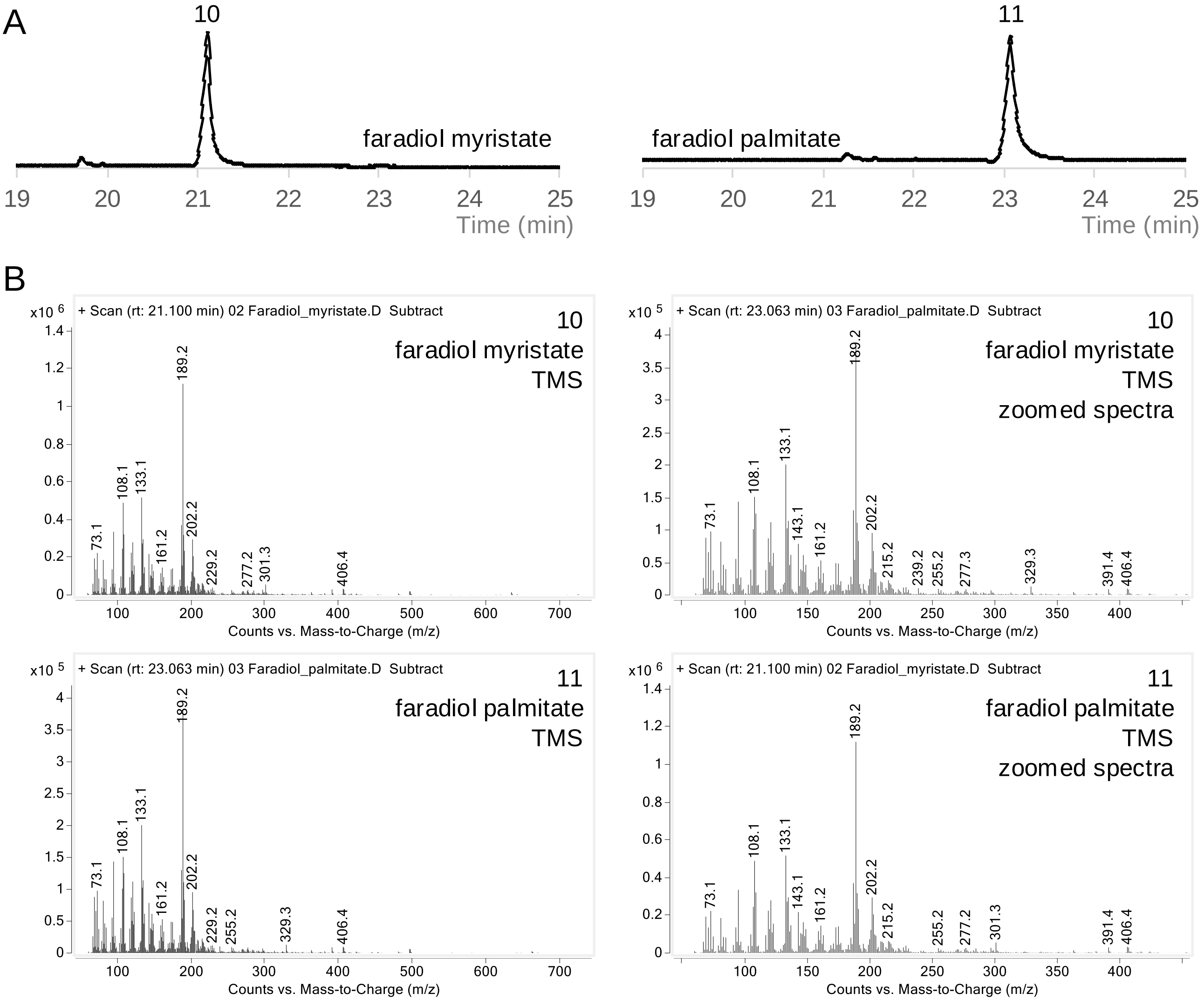

**Supplementary Figure 5. GC-MS standards and mass spectra for faradiol fatty acid esters. A.** GC-MS chromatograms of TMS derivatised faradiol fatty acid esters. Faradiol palmitate and faradiol myristate were purified from *Calendula officinalis* flowers and verified by NMR (Supplementary Figures 6-14; Supplementary Tables 2-3) **B.** Mass spectra for each of the TMS derivatised faradiol fatty acid esters.

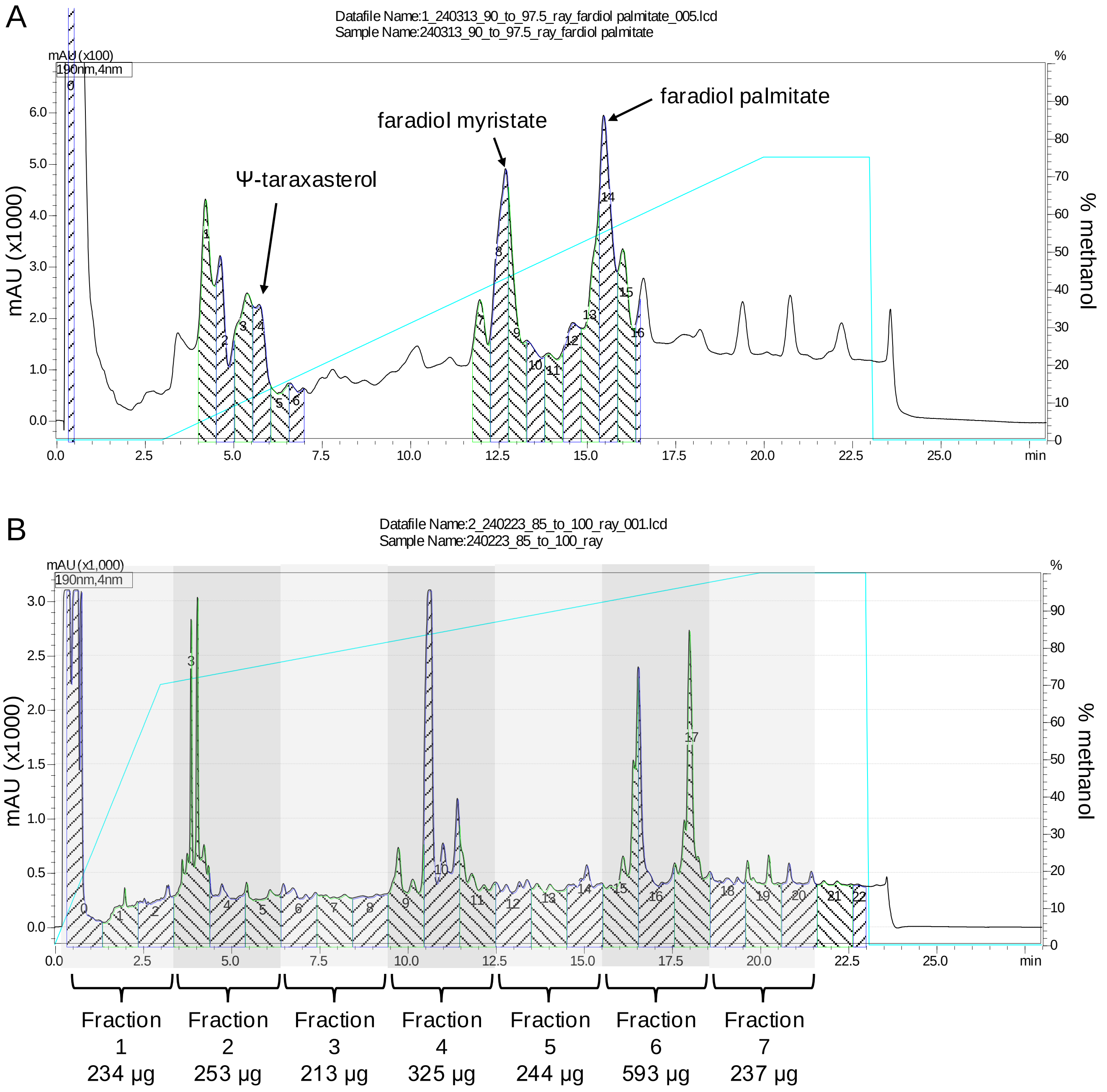

**Supplementary Figure 6. Semi-preparative uHPLC chromatograms of methanol extracts of *Calendula officinalis* ray florets**. The methanol gradient used for separation is shown as a cyan line. **A - Fractionation for purification.** Selected regions were collected as fractions (as indicated in green and blue shading). Multiple runs were performed and fractions corresponding to compounds of interest were pooled, verified by GC-MS analysis and dried down for NMR confirmation. In this run, the following fractions were taken for the compounds of interest: Ψ-taraxasterol (fraction 4); faradiol myristate (fractions 8 and 9); and faradiol palmitate (fraction 14). **B - Fractionation for bioassays.** Selected regions were collected as fractions (as indicated in green and blue shading). A total of 8 runs were performed and fractions were pooled into seven groups of three fractions and dried down for use in bioassays. The yield of each pooled fraction is also shown.

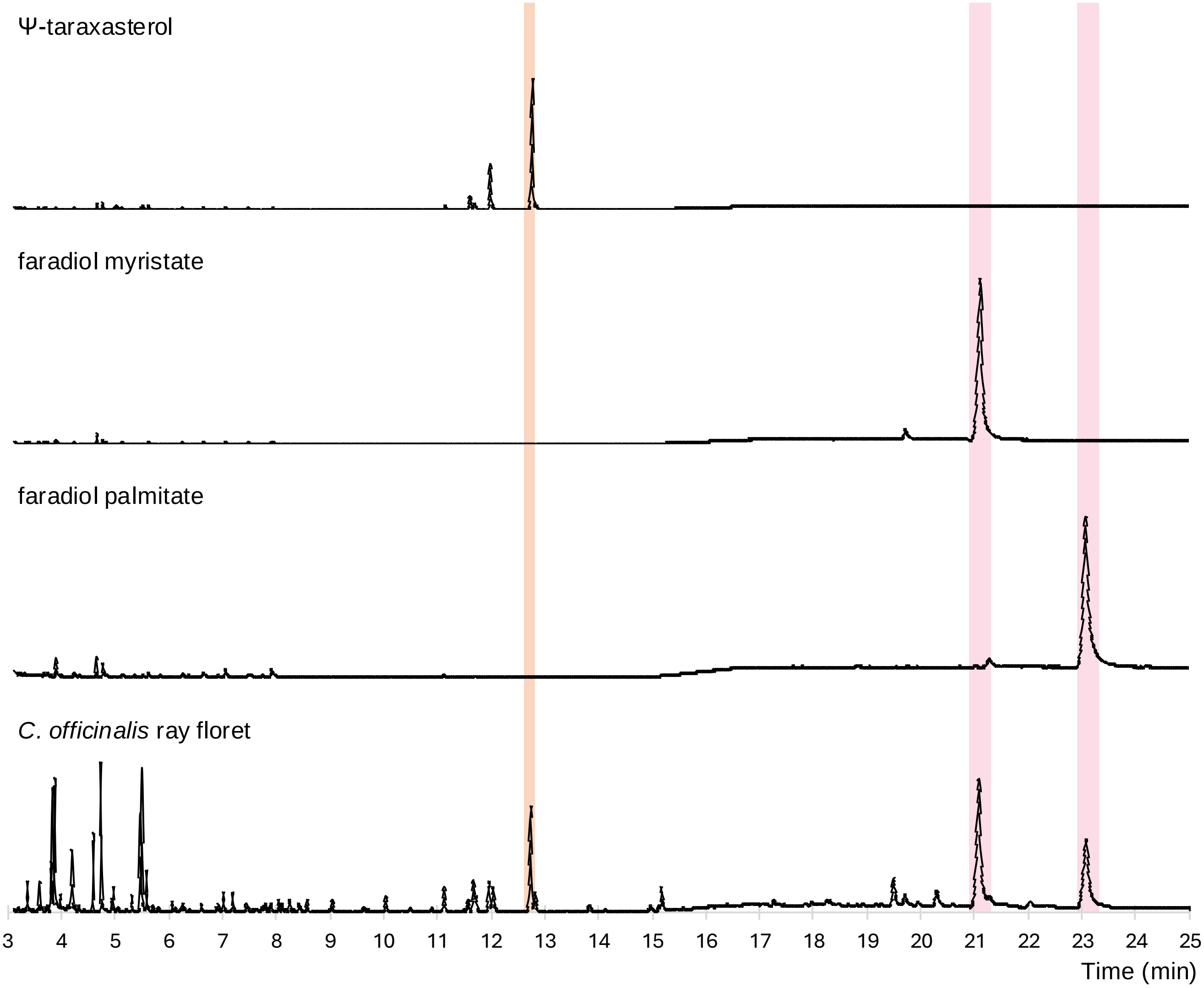

**Supplementary Figure 7. GC-MS analysis of compounds purified from *Calendula officinalis* ray floret extracts.** GC-MS chromatograms of TMS derivatised pooled fractions from purification of triterpene compounds from *C. officinalis* ray floret extract. The ψ-taraxasterol fraction contains a mixture of ψ-taraxasterol (62%) and α-amyrin (19%). The purity of the faradiol myristate and faradiol palmitate fractions is 96% and 98% respectively. Triterpene monols were identified by comparing retention times and mass fragmentation patterns with those of authentic standards (Supplementary Figures 3-5). Faradiol palmitate was confirmed by NMR analysis (Supplementary Figures 9-14; Supplementary Tables 2-3).

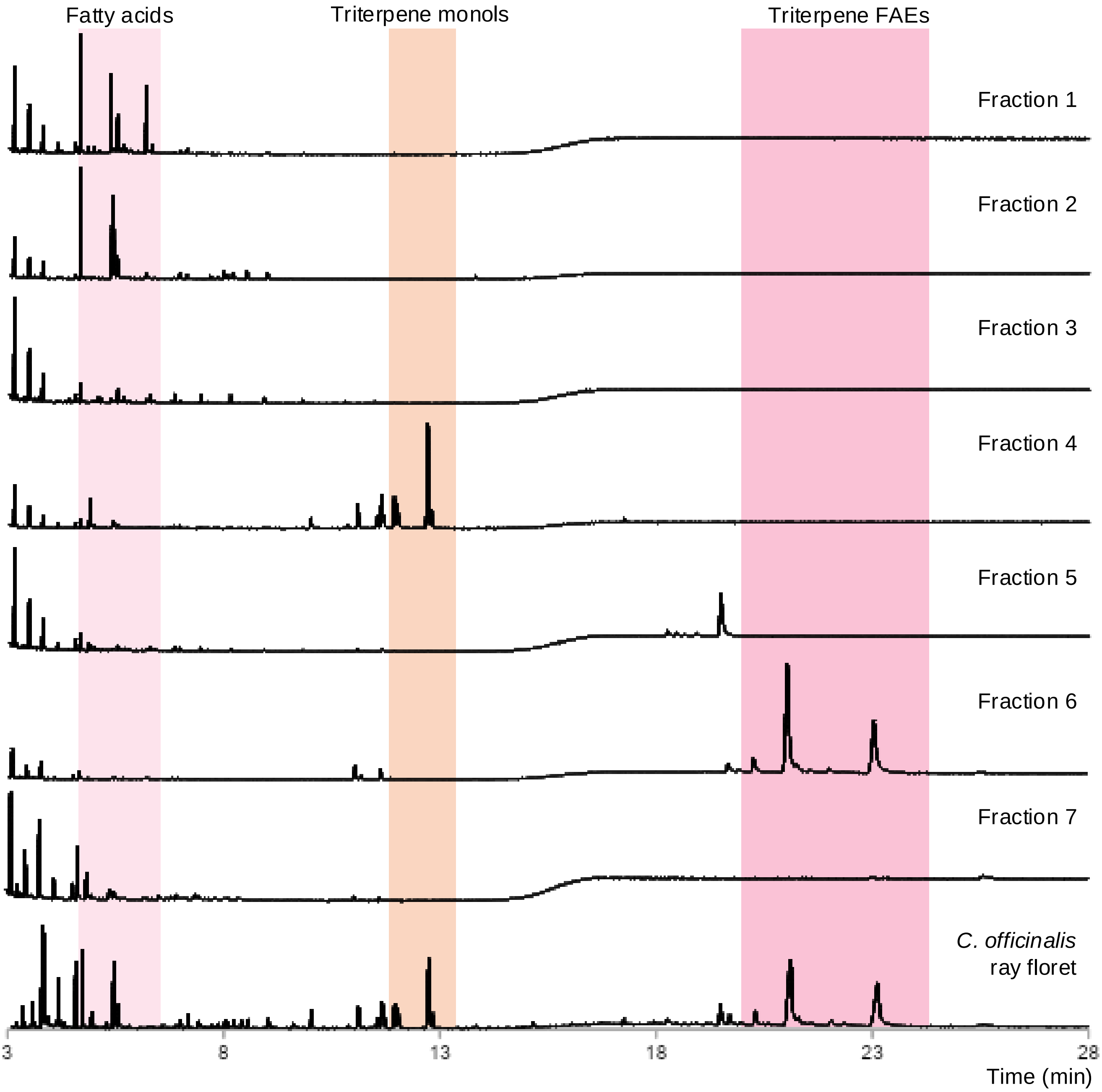

**Supplementary Figure 8. GC-MS analysis of *Calendula officinalis* ray floret fractions.** GC-MS chromatograms of TMS derivatised pooled fractions from fractionation of the *C. officinalis* ray floret extract. Fatty acids were found in fractions 1 and 2; triterpene monols in fraction 4 and triterpene fatty acid esters in fractions 5 and 6. Triterpene monols and diols were identified by comparing retention times and mass fragmentation patterns with those of authentic standards (Supplementary Figures 3-5).

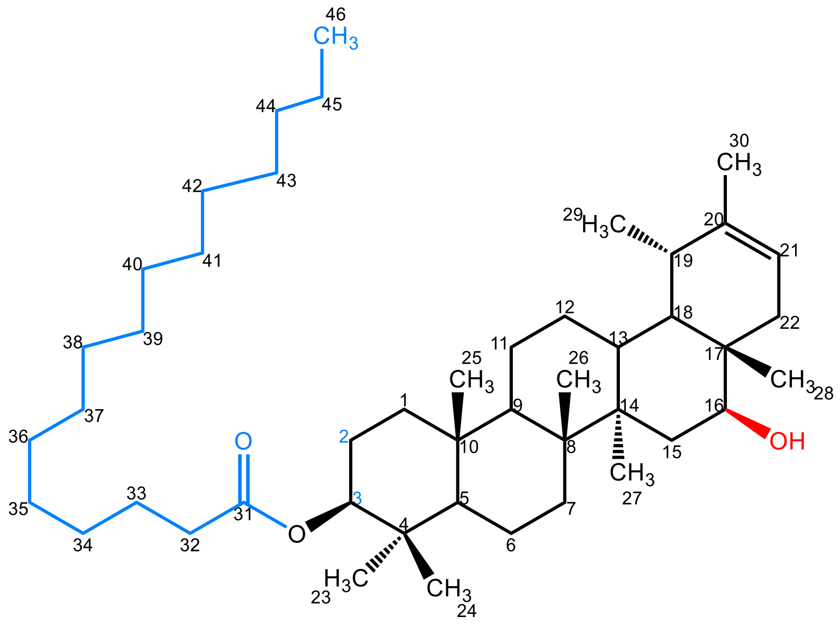

**Supplementary Figure 9. Structure of 3-*O*-palmitoyl faradiol (faradiol palmitate)**

The structure of purified 3-O-palmitoyl faradiol was determined by 1D and 2D NMR -1H, 13 C, COSY, HMQC, and HMBC. The 1H and 13C NMR spectra of 3-O-palmitoyl faradiol were in a good agreement with spectra published by Ezzat et al (2017) Nat Prod Res. 31(6):676-680. Using HMBC spectra it was possible to identify the positions of C31, C32, C45 and C46 of palmitoyl residue that were not previously assigned. Furthermore, cross peaks of 4.48 ppm (in proton spectra dimension) with 173.7 (C31), 37.8 (C-4), 28.0 (C23) and 16.6 (C24) in HMBC spectra indicate that the signal at 4.48 ppm belongs to the resonance of H3 thus establishing the position 3 as a position of esterification of faradiol. In 1H, the NMR spectra resonance of methine proton H3 (4.48 ppm) appears 1 ppm downfield compared to the resonance of H16 (3.44 ppm) of hydromethine group. This characteristic downfield shift due to esterification confirms the position of the palmitoyl attachment to O-3.

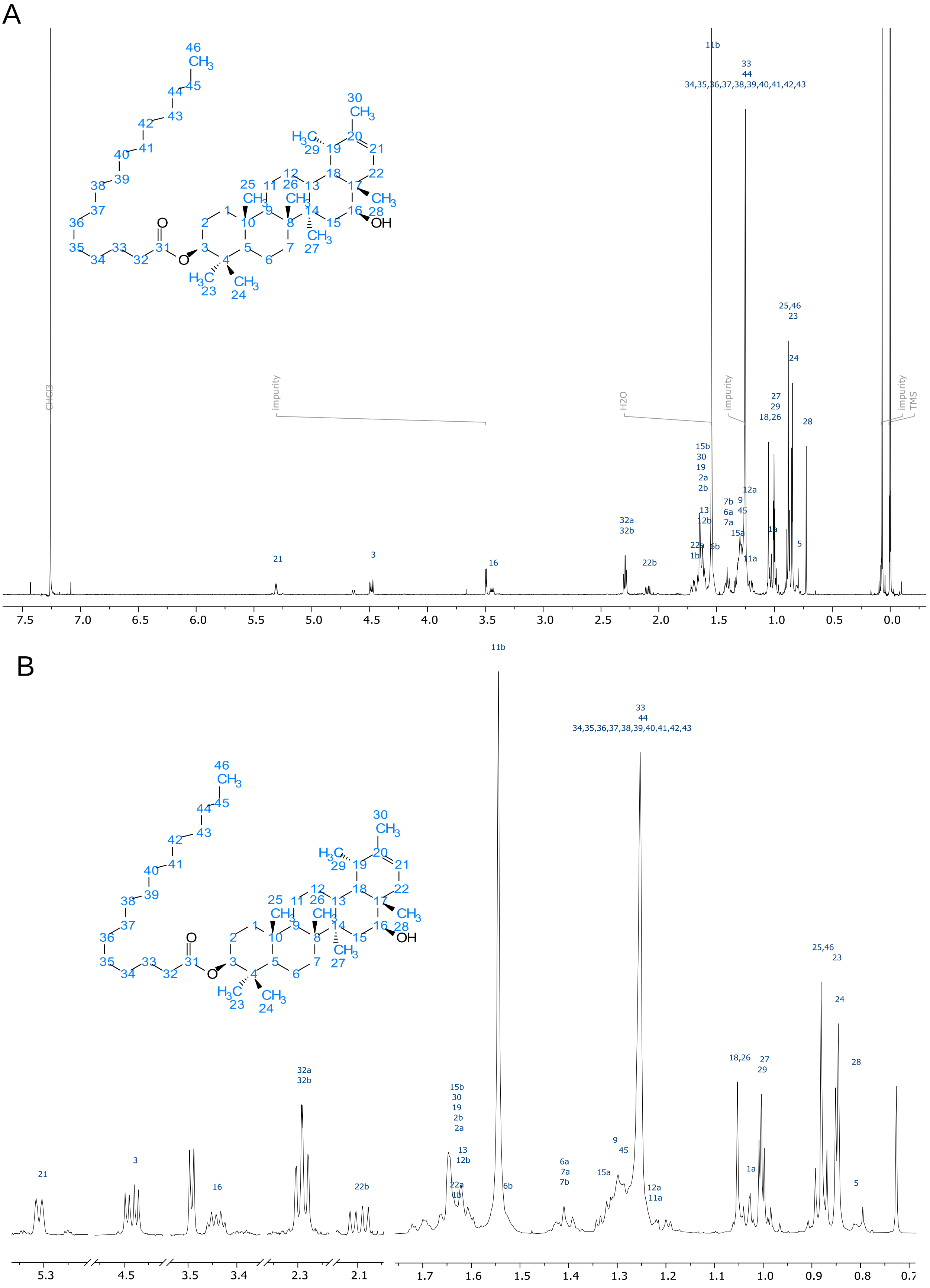

**Supplementary Figure 10. $^{1}$H NMR spectra of faradiol palmitate. A.** Overview of $^{1}$H NMR (600 MHz, CDCl3, 298 K) of 3-O-palmitoyl faradiol. **B.** Expansion of 3-O-palmitoyl faradiol resonances in $^{1}$H NMR (600 MHz, CDCl3, 298 K).

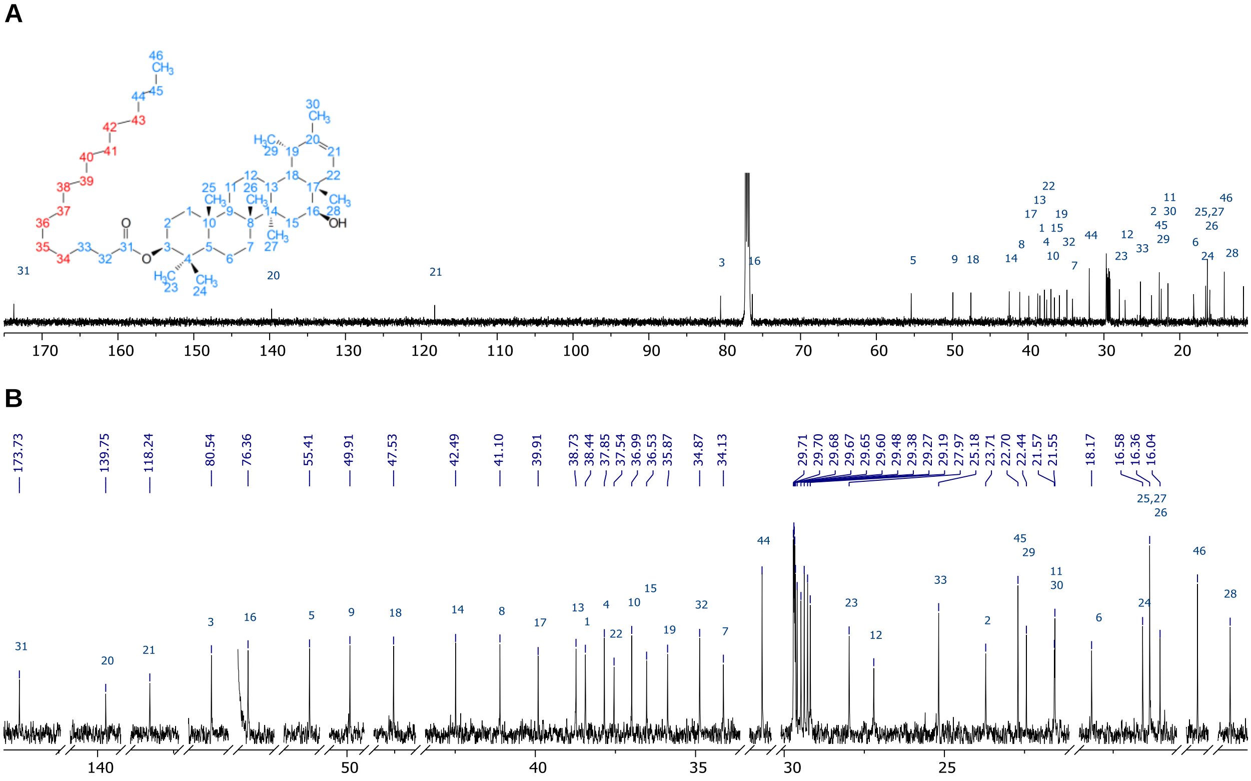

**Supplementary Figure 11. $^{13}$C NMR spectra of faradiol palmitate. A.** Overview; **B.** Expansion.

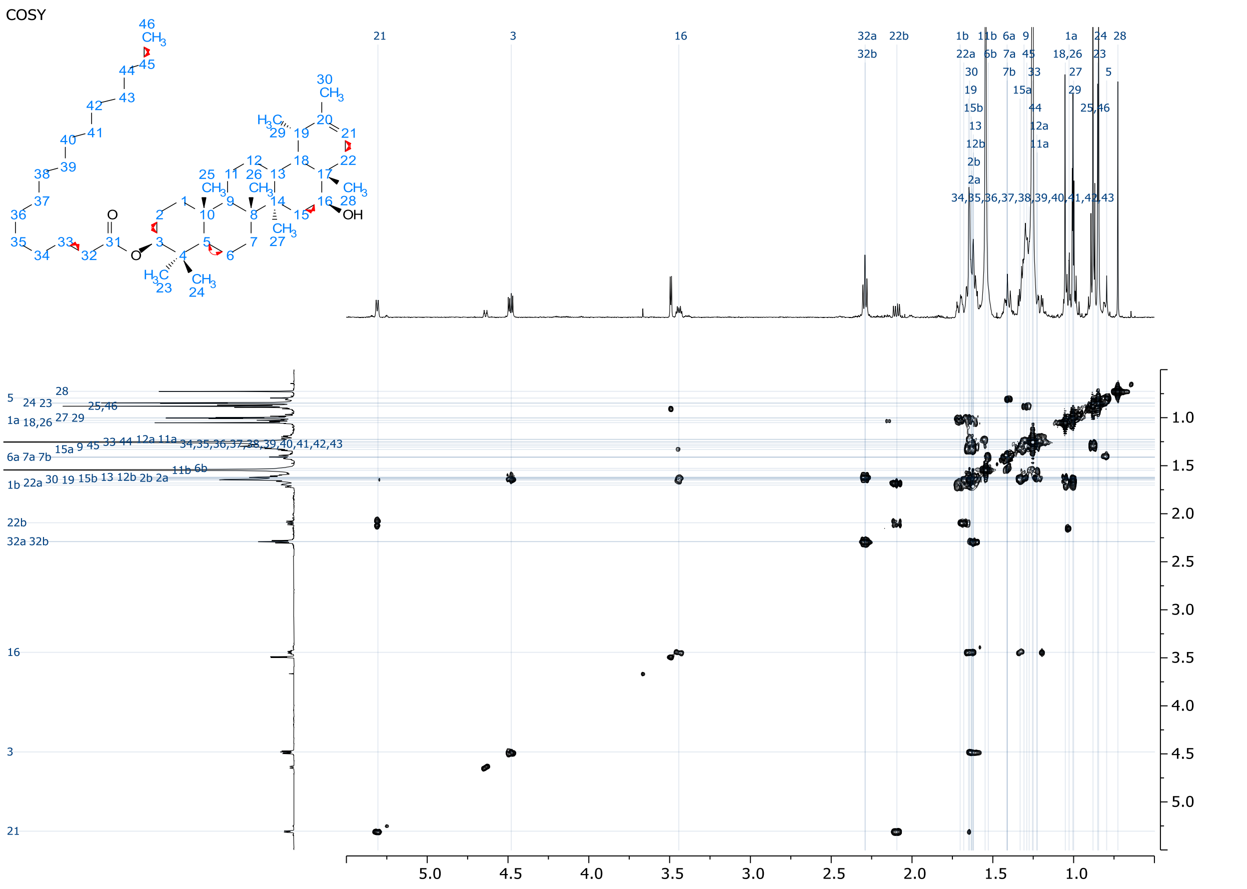

**Supplementary Figure 12. $^{1}$H-$^{1}$H COSY NMR (600 MHz, CDCl3, 298 K) of faradiol palmitate.**
Only limited number of correlations can be found due to signals overlap.

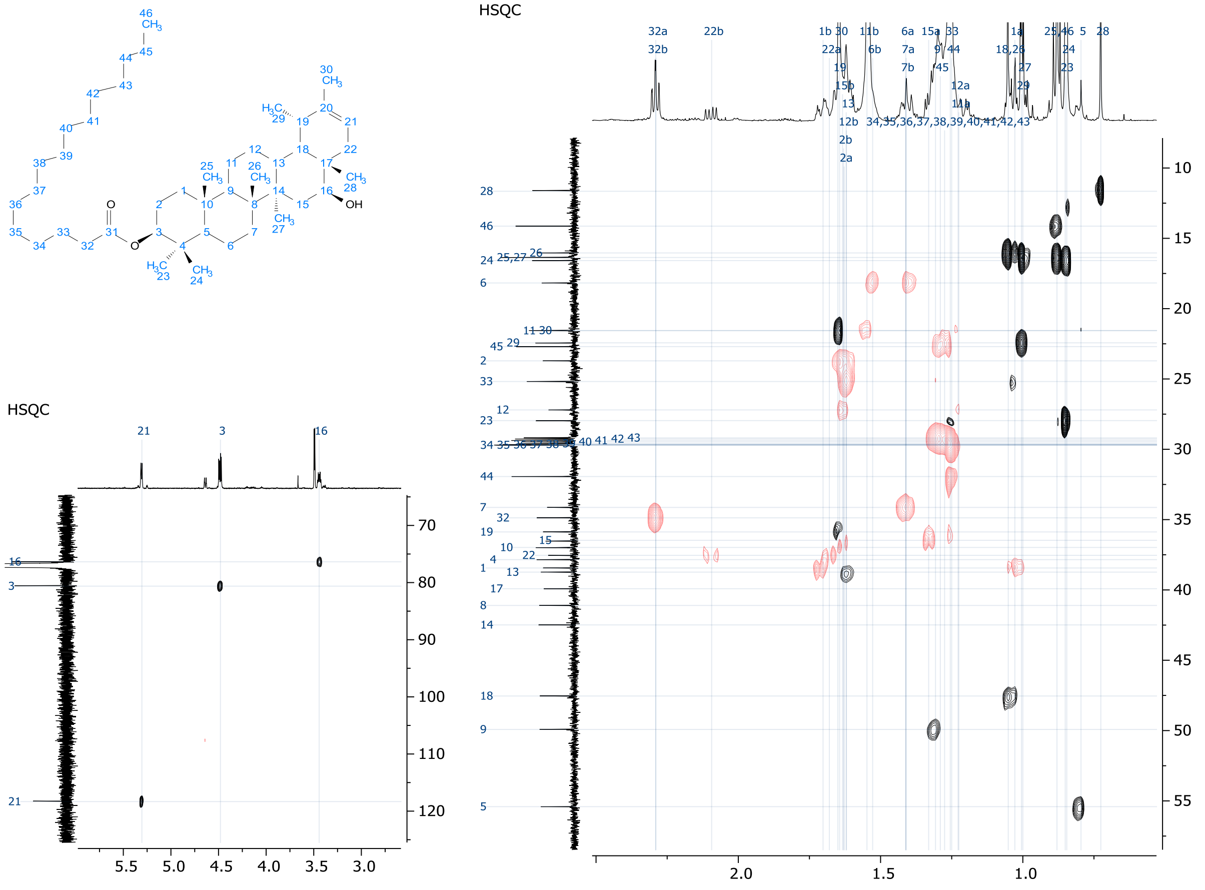

**Supplementary Figure 13. $^{1}H$-$^{13}C$-HSQC-edited NMR (600 MHz, CDCl3, 298 K) of faradiol palmitate.**

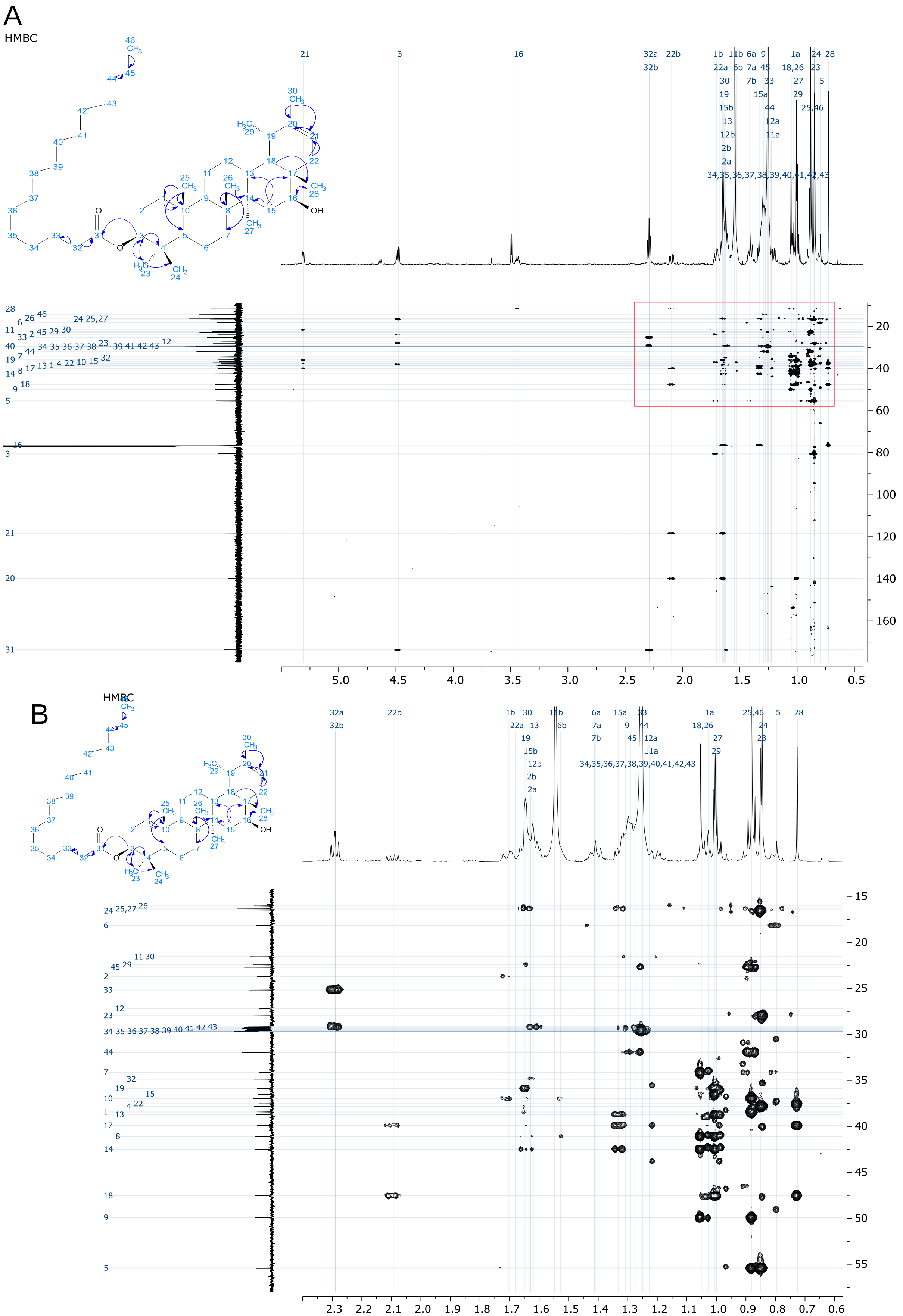

**Supplementary Figure 14.** $^{1}H$-$^{13}C$ **HMBC NMR (600 MHz, CDCl3, 298 K) of faradiol palmitate. A.** Overview **B.** Expansion of area highlighted in A. Curly arrows indicate observed HMBC interactions.

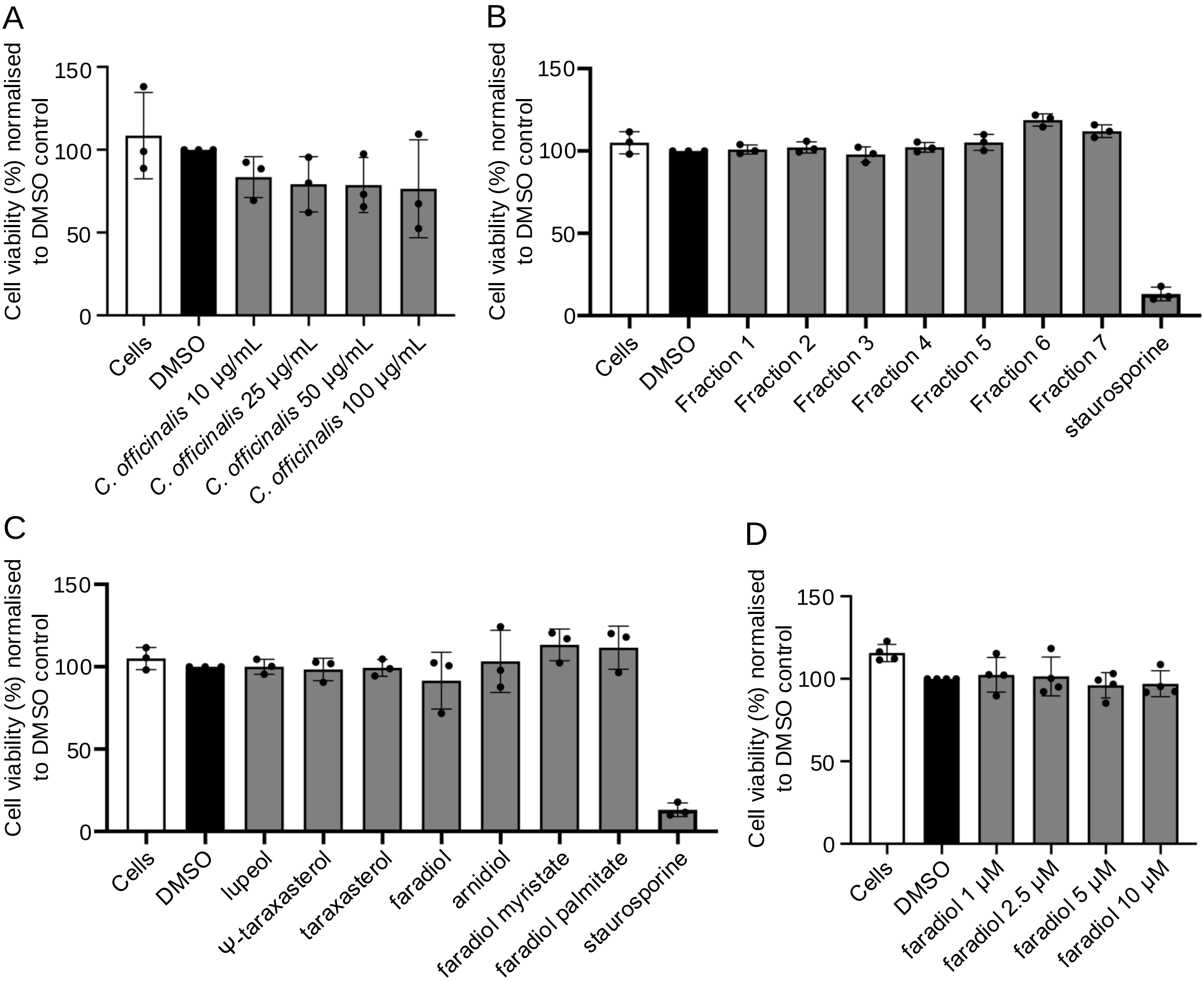

**Supplementary Figure 15. Effects of Calendula officinalis (pot marigold) extracts and triterpenoids on the viability of human monocytic (THP-1) cells. A** Cell viability following application of pot marigold floral extracts at different concentrations. **B** Cell viability following application of extract fractions. C Cell viability following application of triterpenoid compounds (20 µM). D Cell viability following application of faradiol at different concentrations.

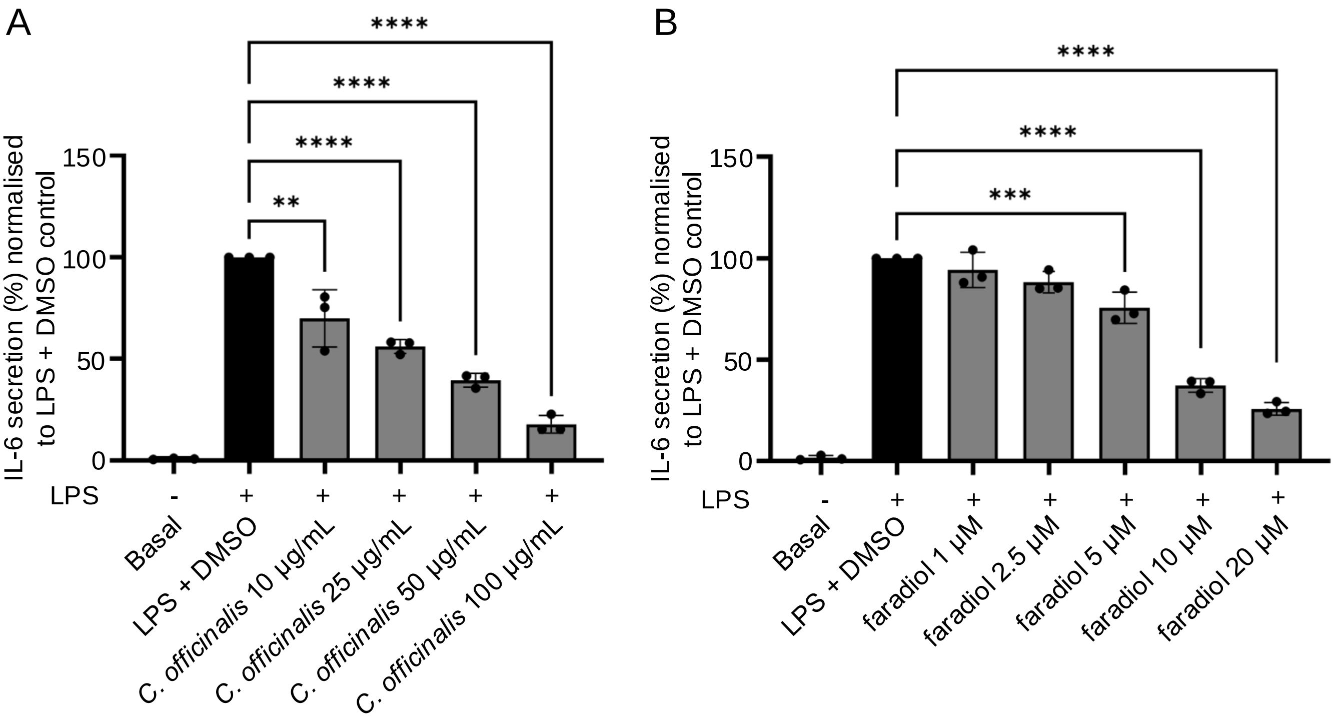

**Supplementary Figure 16. Concentration-dependent responses to floral extracts and faradiol.** Concentration-dependent anti-inflammatory effect of: **A.** Calendula officinalis (pot marigold) floral extracts and **B.** Faradiol , on the release of IL-6 in (LPS)-activated human monocytic (THP-1) cells.

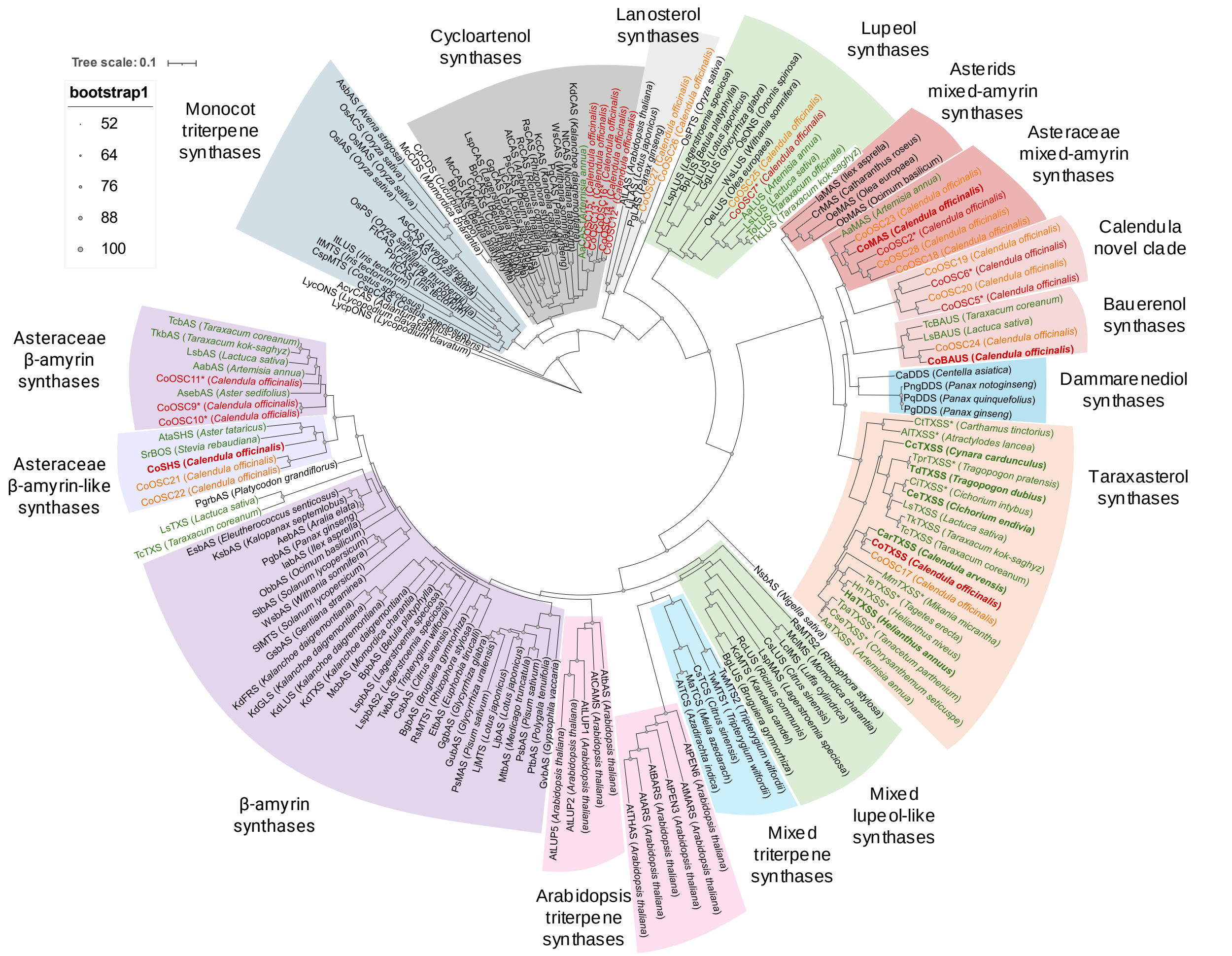

**Supplementary Figure 17. Maximum-likelihood tree of plant oxidosqualene cyclases (OSCs)**. Functionally characterised non-Asteraceae and Asteraceae plant OSCs are shown in black and green respectively. *Calendula officinalis* plant OSCs that are expressed are shown in red with likely pseudogenes shown in orange. Genes that were functionally characterised in this paper are shown in bold and any uncharacterised genes are indicated with an asterisk (*). The maximum likelihood tree was constructed in IQ-Tree using the JTT matrix-based model and visualised using Interactive Tree of Life (iTOL) v6. Filled grey circles indicate bootstrap supports for each node. Scale bar represents the number of substitutions per site. Clades are highlighted and labelled according to product specificity.

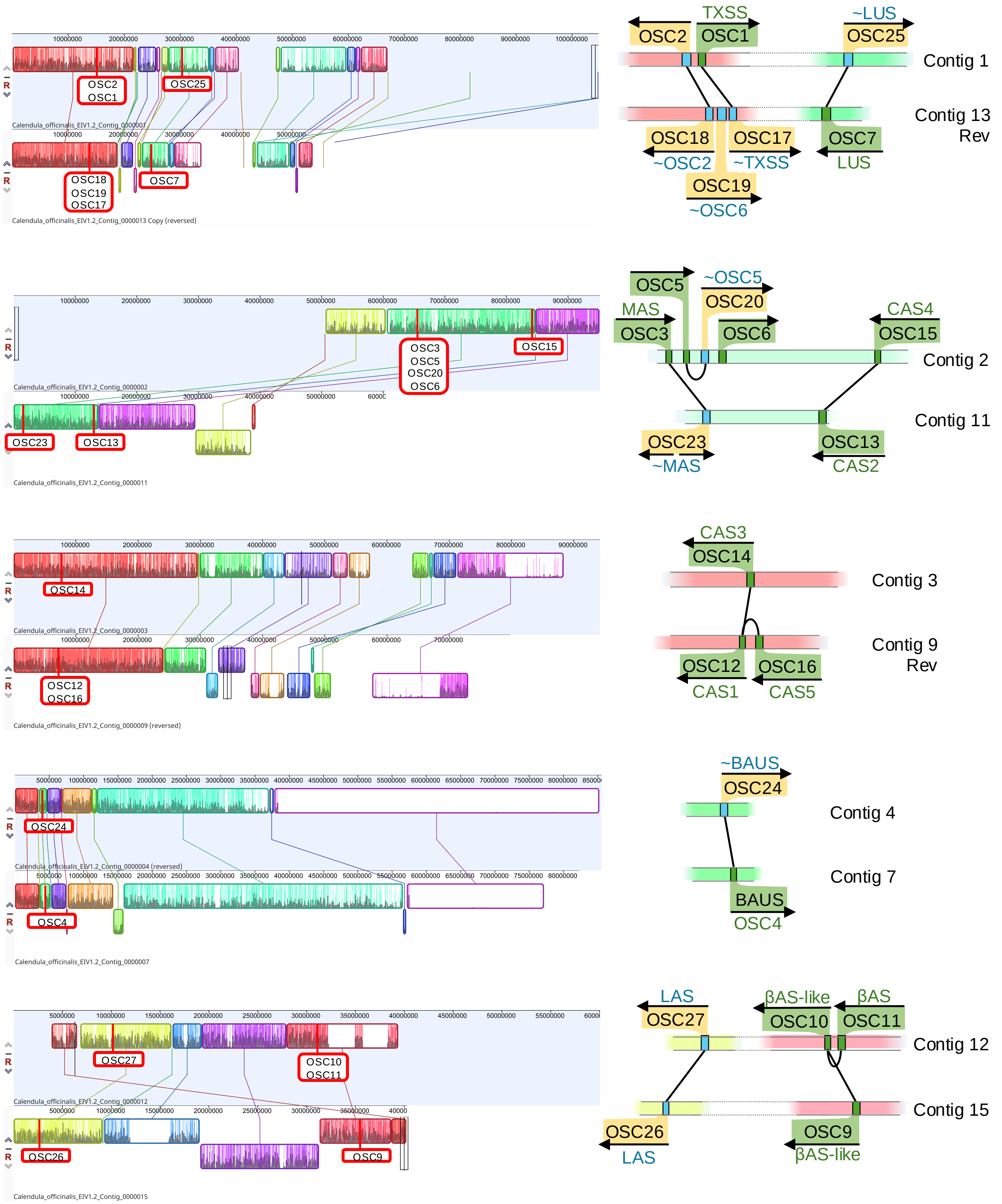

**Supplementary Figure 18. Genomic locations of *Calendula officinalis* (pot marigold) genes encoding oxidosqualene cyclases (OSCs).** A graphical representation of the relative position, orientation and similarity of *CoOSC* genes. Contigs are paired to show synteny between genes found on homeologous contigs. Genes that were expressed in the pot marigold transcriptome are shown in green. Those for which expression was not detected are shown in yellow and those not expressed and with mutations indicating pseudogenisation are prefixed "~". Functional annotations based on phylogenetic relationships to previously characterised OSCs: TXSS = taraxasterol synthase; βAS = β-amyrin synthase; LAS = Lanosterol synthase; BAUS = baurenol synthase; CAS = cycloartenol synthase; MAS = mixed amyrin synthase. LUS = lupeol synthase.

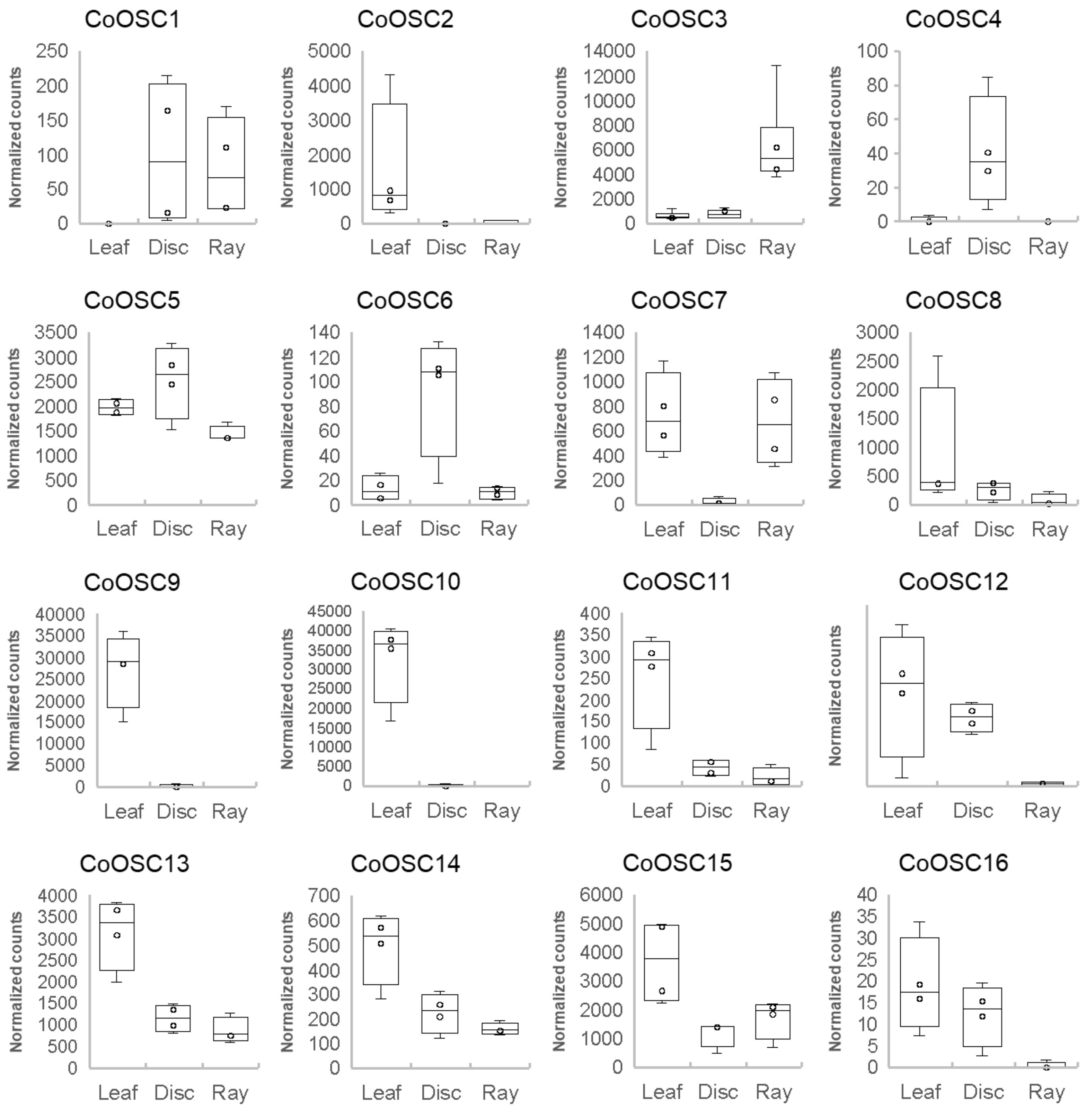

**Supplementary Figure 19. Differential expression of *Calendula officinalis OSC* genes.** Gene expression of 16 *OSC* genes expressed in leaf, disc and ray floret tissues. Gene expression for each tissue is displayed in a box plot as normalised counts following transcript quantification using Salmon v1.2.0 and differential gene expression analysis using DESeq2. The centre line of the boxplot denotes the median, the box denotes the 25th to 75th percentiles of the data and the whiskers show the minimum and maximum of the data.

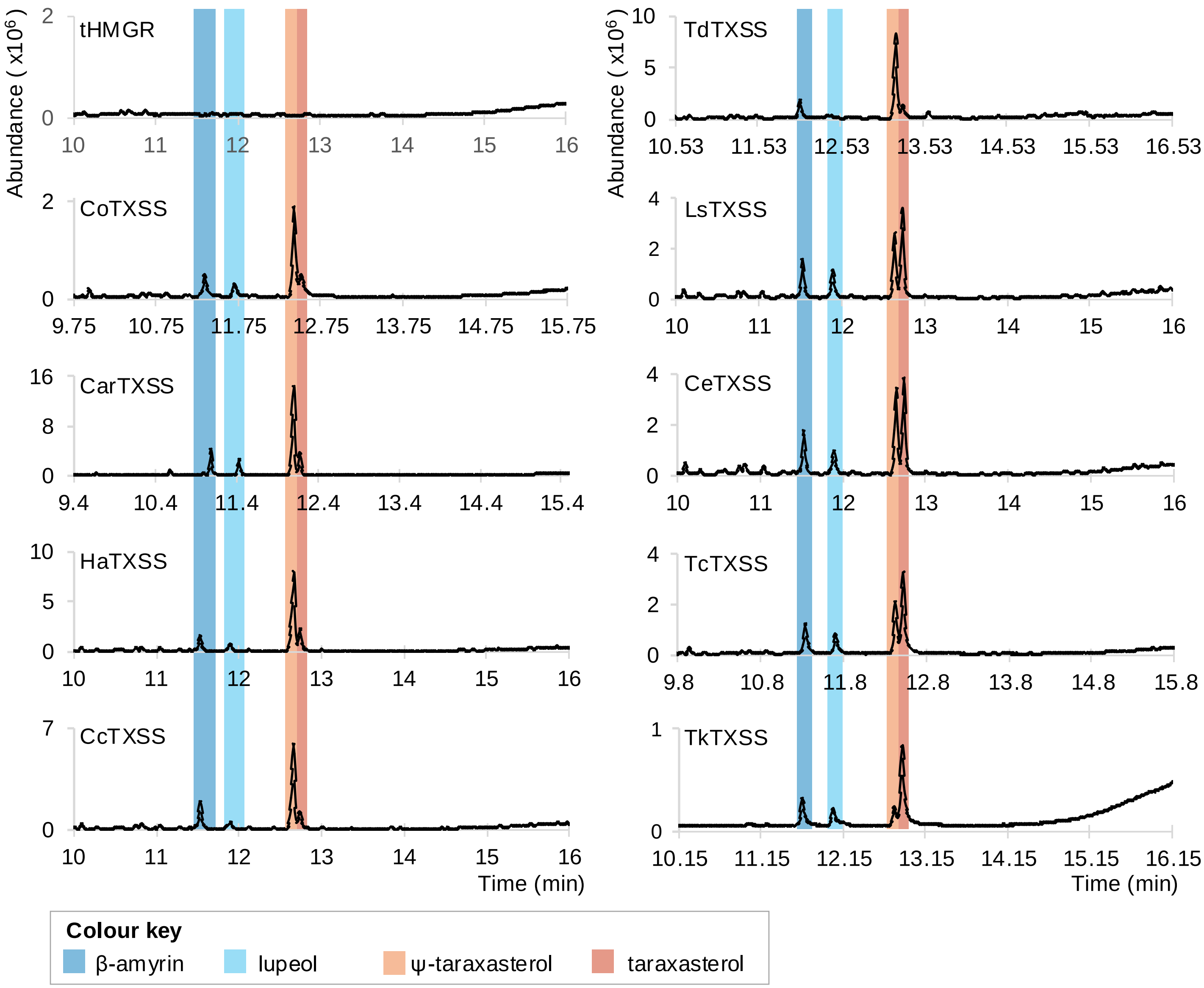

**Supplementary Figure 20. GC-MS analysis of *Nicotiana benthamiana* expressing taraxasterol synthases (TXSS).** Representative total ion chromatograms of ethyl acetate extracts of *Nicotiana benthamiana* leaves transiently expressing *TXSS* genes from Asteraceae species. *CoTXSS* and *CarTXSS* were mined from data generated for this manuscript; *HaTXSS, CcTXSS, TdTXSS* and *CeTXSS* were mined from publicly available datasets; *LsTXSS*, *TcTXSS* and *TkTXSS* were previously characterised. Key metabolites are highlighted and detailed in the colour key. *Co = Calendula officinalis; Car = Calendula arvensis; Ha = Helianthus annuus; Cc = Cynara cardunculus; Td = Tragopogon dubius; Ls = Lactuca sativa; Ce = Cichorium endivia; Tc = Taraxacum coreanum; TkTXSS = Taraxacum kok-saghyz*.

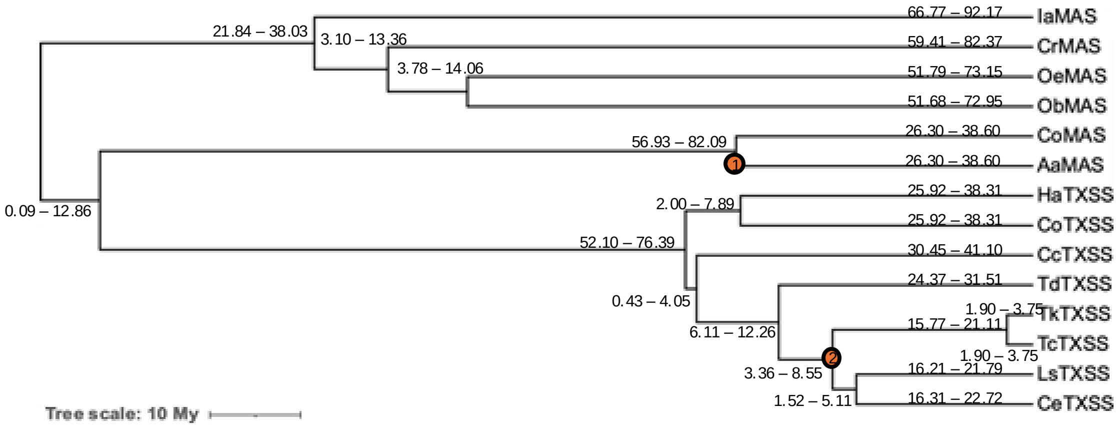

**Supplementary Figure 21. Bayesian phylogenetic tree of mixed amyrin synthases (MAS) and taraxasterol synthases (TXSS).** Bayesian phylogenetic analysis was performed using BEAST2 with a GTR model and an *Artemisia* fossil (orange circle; 1.31 mya) and a *Cichorium intybus* type fossil (orange circle; 2.22 mya) as calibrators. Numbers show branch length range. Ia=*Ilex asprella ;* Cr=*Catharanthus roseus;* Oe=*Olea europaea;* Ob=*Ocimum basilicum;* Co=*Calendula officinalis;* Aa=*Artemisia annua;* Ha=*Helianthus annuus;* Cc=*Cynara cardunculus;* Td=*Tragopogon dubius;* Tk=*Taraxacum kok-saghyz;* Tc=*Taraxacum coreanum;* Ls=*Lactuca sativa;* Ce=*Cichorium endivia*.

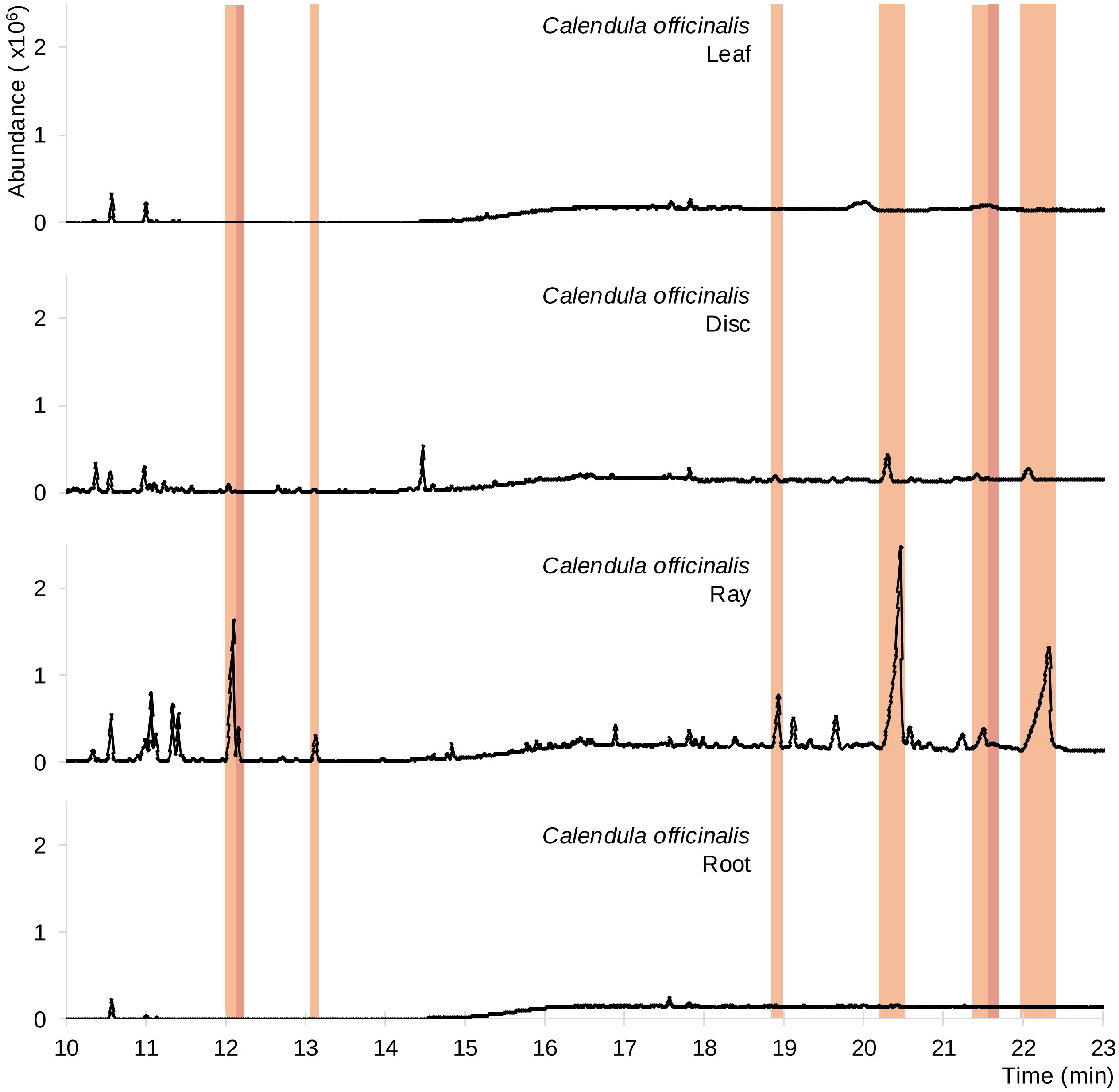

**Supplementary Figure 22. GC-MS profile of *Calendula officinalis* (pot marigold) tissues.** Representative total ion chromatograms of ethyl acetate extracts of pot marigold tissues to identify ψ-taraxasterol and taraxasterol metabolites and their derivatives. ψ-taraxasterol and its derivatives are highlighted in orange and taraxasterol and its derivatives are highlighted in red.

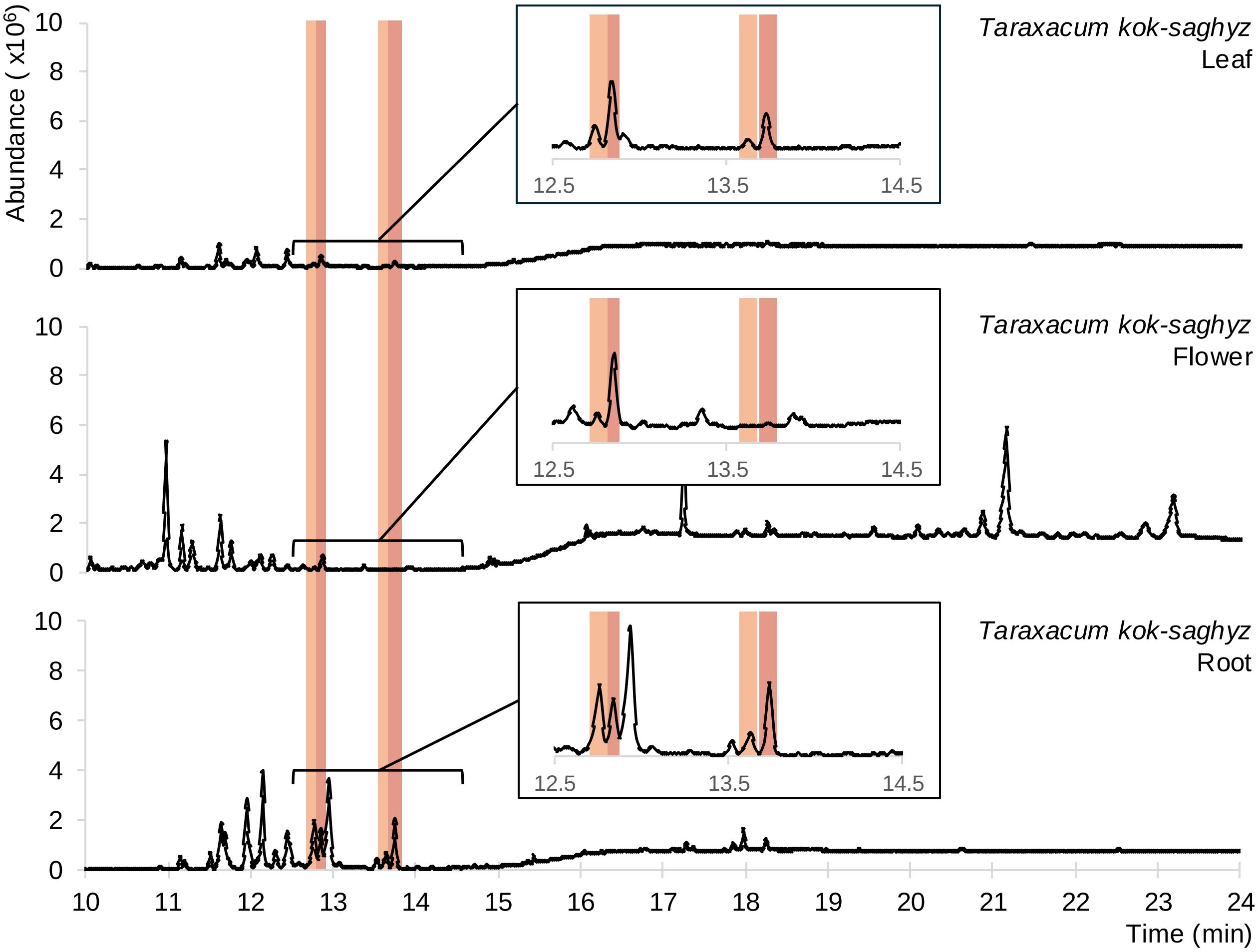

**Supplementary Figure 23. GC-MS profile of *Taraxacum kok-saghyz* (Russian dandelion) tissues.** Representative total ion chromatograms of ethyl acetate extracts of Russian dandelion tissues to identify ψ-taraxasterol and taraxasterol metabolites and their derivatives. ψ-taraxasterol and its derivatives are highlighted in orange and taraxasterol and its derivatives are highlighted in red.

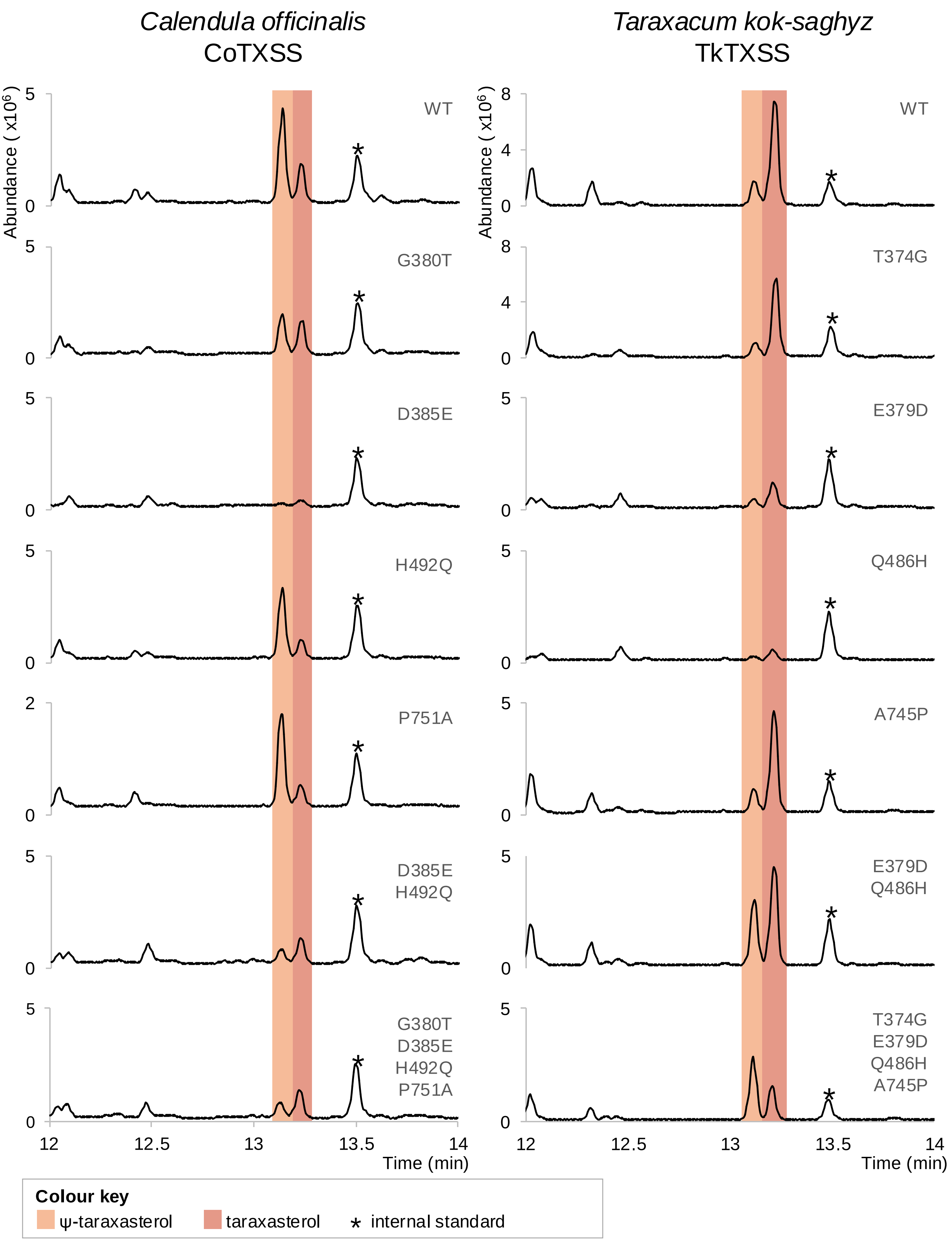

**Supplementary Figure 24. GC-MS analysis of *Nicotiana benthamiana* expressing taraxasterol synthase (TXSS) mutants.** Total ion chromatograms of extracts of *N. benthamiana* leaves transiently expressing wild type and mutated CoTXSS and TkTXSS. Key metabolites are highlighted and detailed in the colour key.

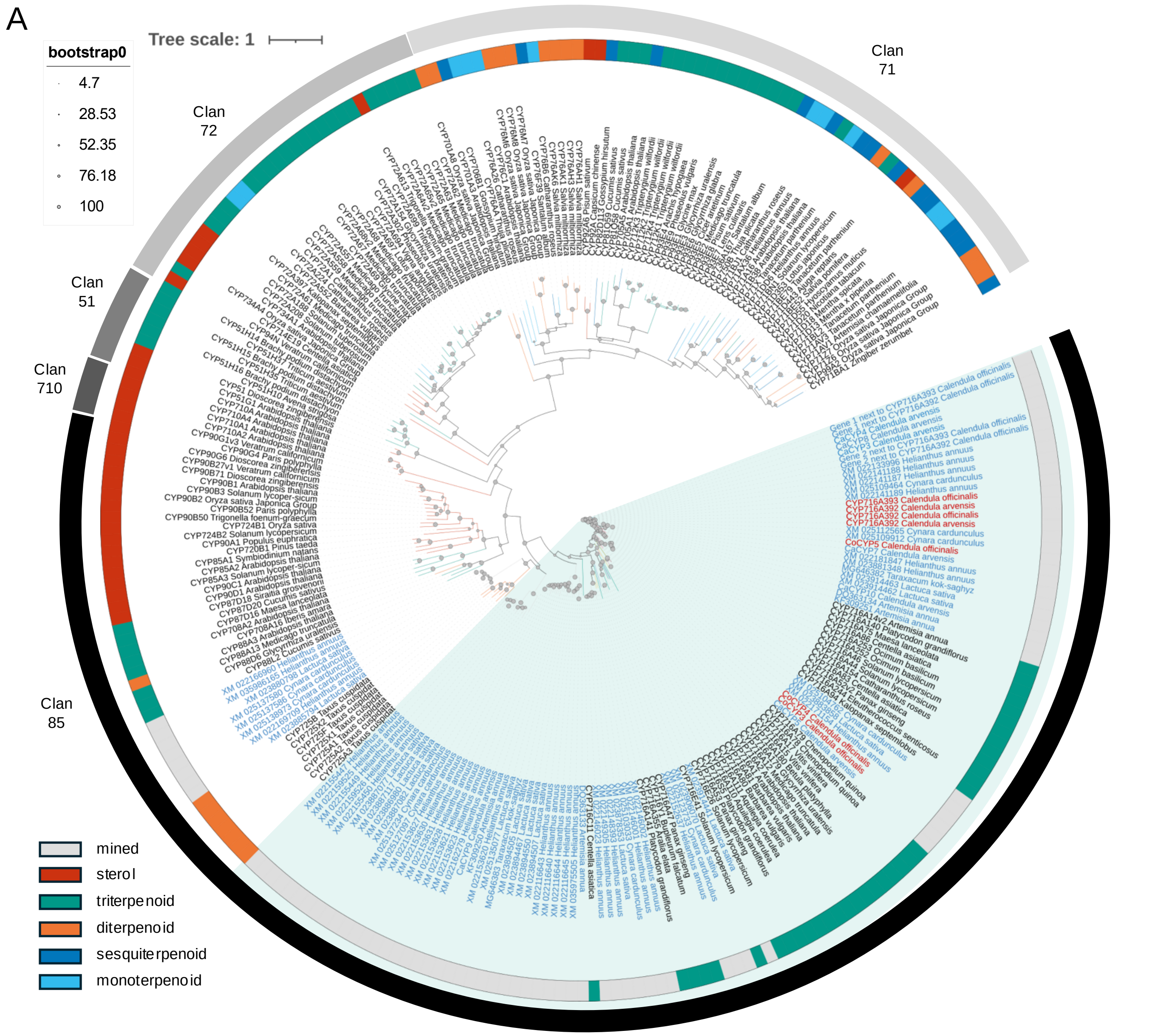

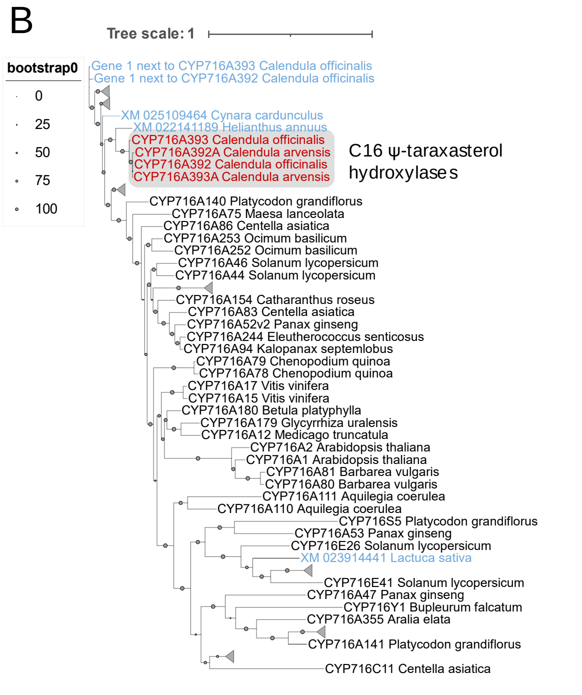

**Supplementary Figure 25. Maximum-likelihood tree of plant cytochrome p450s (CYPs).**
**A.** Maximum likelihood tree was constructed in IQ-Tree using the LG+F+I+G4 matrix-based model and visualised using Interactive Tree of Life (iTOL) v6. Substrate specificity is shown in the inner ring; CYPs clans are shown in the outer ring, CYP716 family triterpenoid modifying clade is highlighted in light teal**.** Filled grey circles indicate bootstrap supports for each node. The scale bar represents the number of substitutions per site. Functionally characterised taxa are shown in black. CYPs sequences mined from the publicly available genomes of Asteraceae, *Calendula officinalis* genome and transcriptome datasets, and *Calendula arvensis* transcriptome, are shown in blue. *C. officinalis* and *C. arvensis* C16 ψ-taraxasterol hydroxylases are shown in red. **B.** Extracted CYP716 family showing characterised p450s (black text), uncharacterised genes (blue text) and C16 ψ-taraxasterol hydroxylases (red text, grey shading)

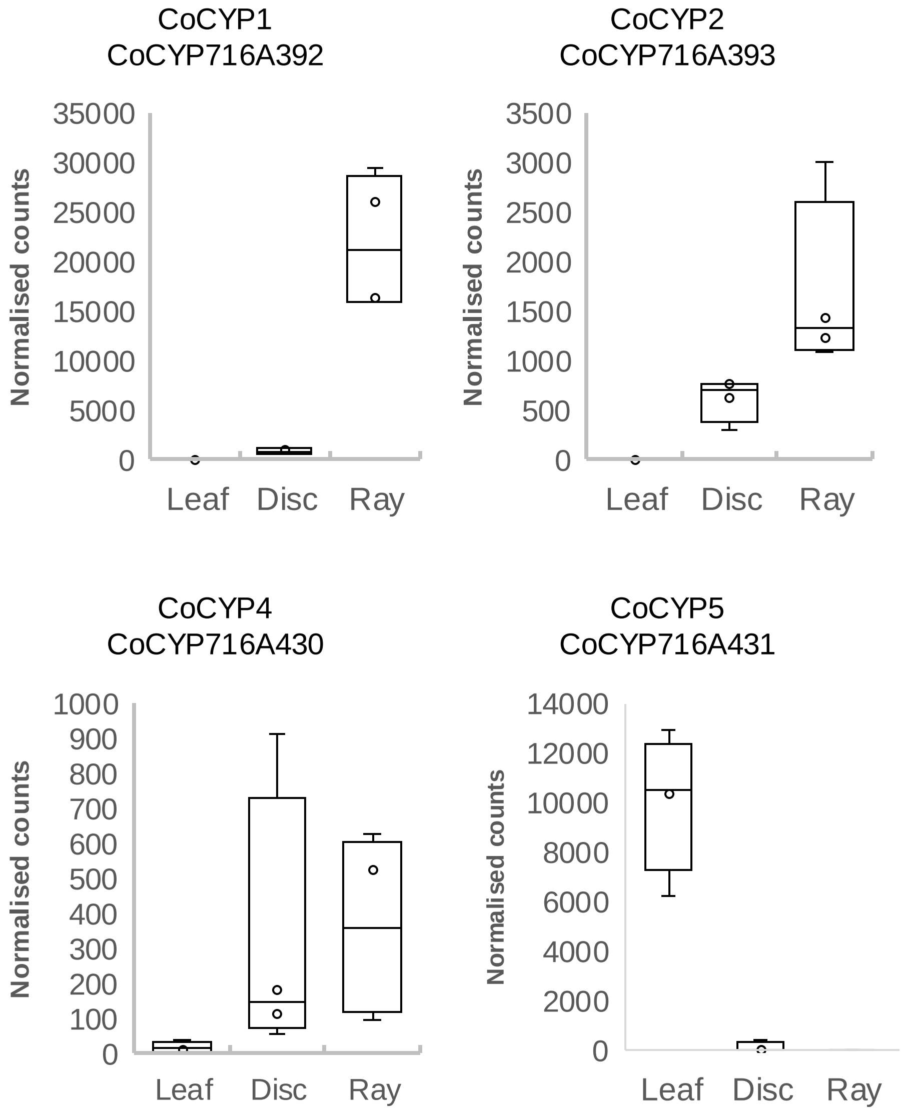

**Supplementary Figure 26. Differential expression of *Calendula officinalis* (pot marigold) CYP genes.** Gene expression of five cytochrome P450 (CYP) genes expressed in leaf, disc and ray floret tissues. Gene expression for each tissue is displayed in a box plot as normalised counts following transcript quantification using Salmon v1.2.0 and differential gene expression analysis using DESeq2. The centre line of the boxplot denotes the median, the box denotes the 25th to 75th percentiles of the data and the whiskers show the minimum and maximum of the data.

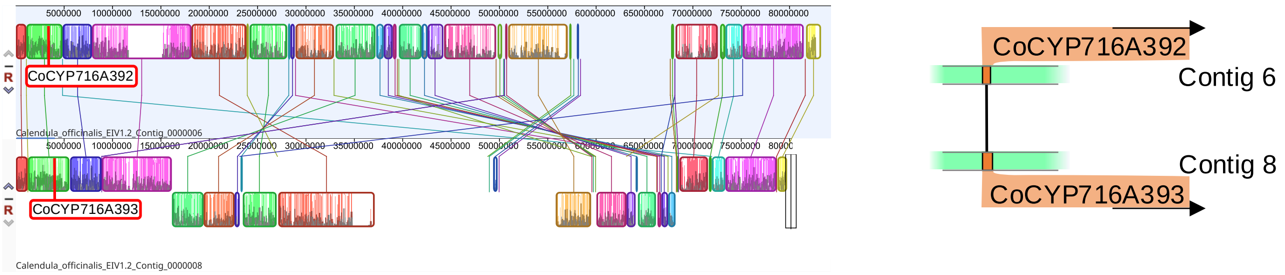

**Supplementary Figure 27. Genomic location and synteny of *Calendula officinalis* (pot marigold) cytochrome P450 genes encoding CoCYP716A392 and CoCYP716A393.** Chromosome synteny and graphical representation of the contig positions of cytochrome P450 and acyltransferase genes involved in the triterpene tatty acid ester (TFAE) pathway. Contigs are paired to show synteny between genes found on homeologous contigs. A graphical representation of the relative position, orientation and similarity of genes within each contig pair are shown to the right of the contig synteny plot.

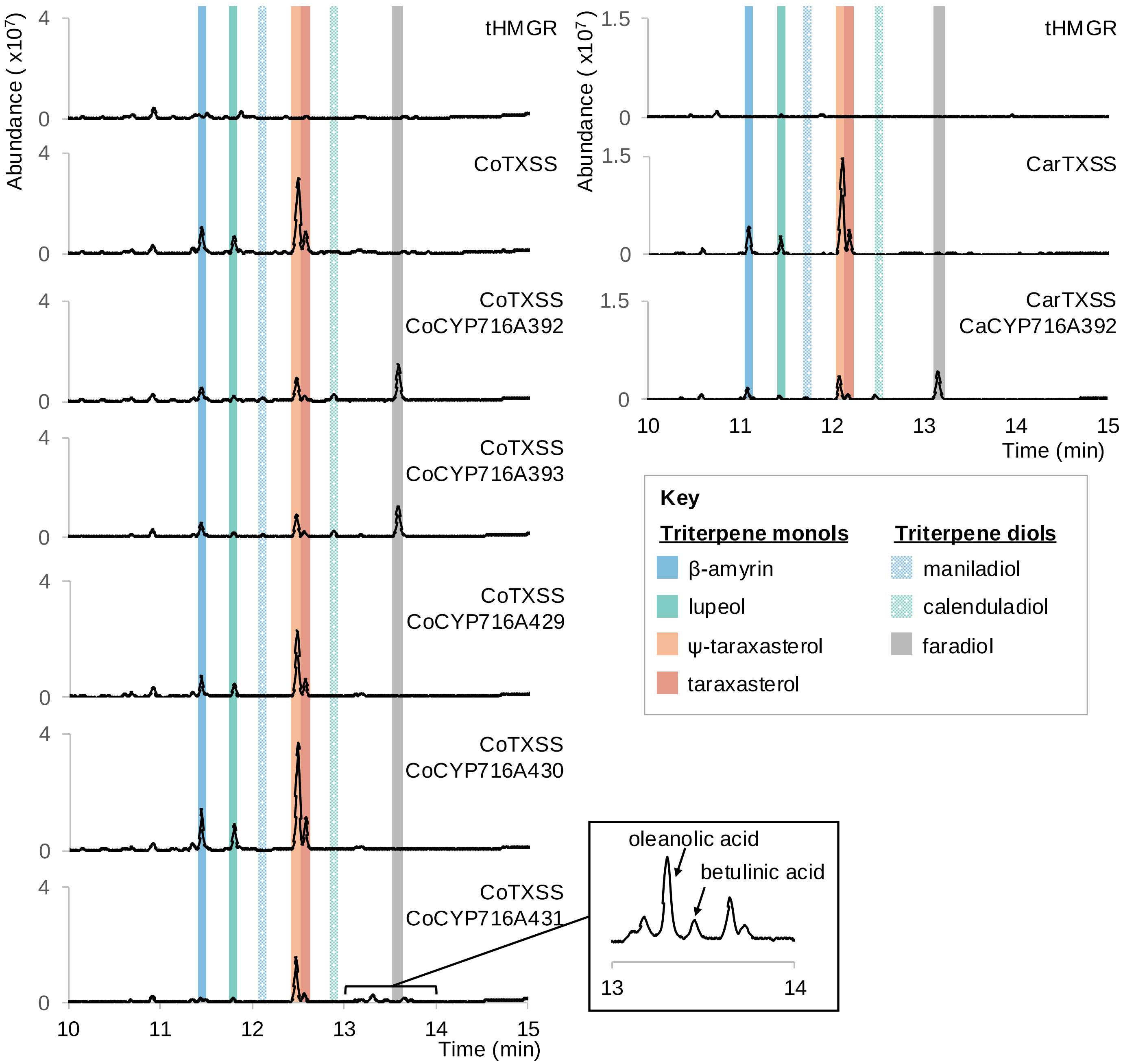

**Supplementary Figure 28. GC-MS analysis of *Nicotiana benthamiana* expressing cytochrome P450s (CYPs).** Representative total ion chromatograms of ethyl acetate extracts of *N. benthamiana* leaves transiently expressing *TARAXASTEROL SYNTHASE* (*TXSS)* genes and *CYP* candidate genes mined from *Calendula officinalis* and *Calendula arvensis* transcriptomes.

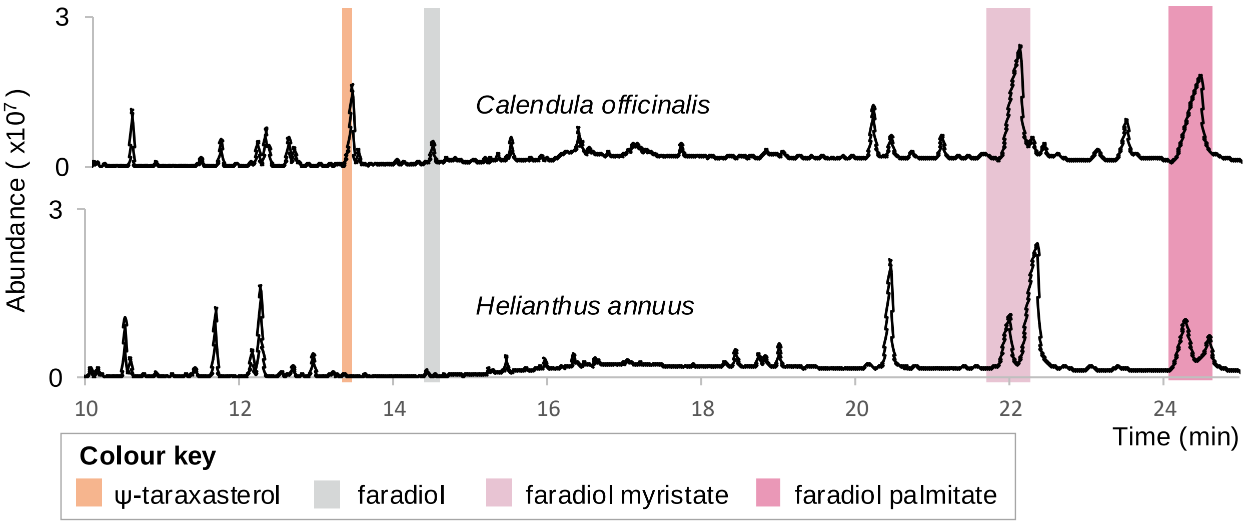

**Supplementary Figure 29. GC-MS profiling of floral extracts from Asteraceae species.** Total ion chromatograms of ethyl acetate extracts of floral tissues from five Asteraceae species identified to encode a protein with high similarity to CoCYP716A392/CoCYP716A393. Faradiol is highlighted in grey; its fatty acid derivatives, faradiol myristate and faradiol palmitate, are highlighted in light pink and pink respectively; its precursor ψ-taraxasterol is highlighted in orange.

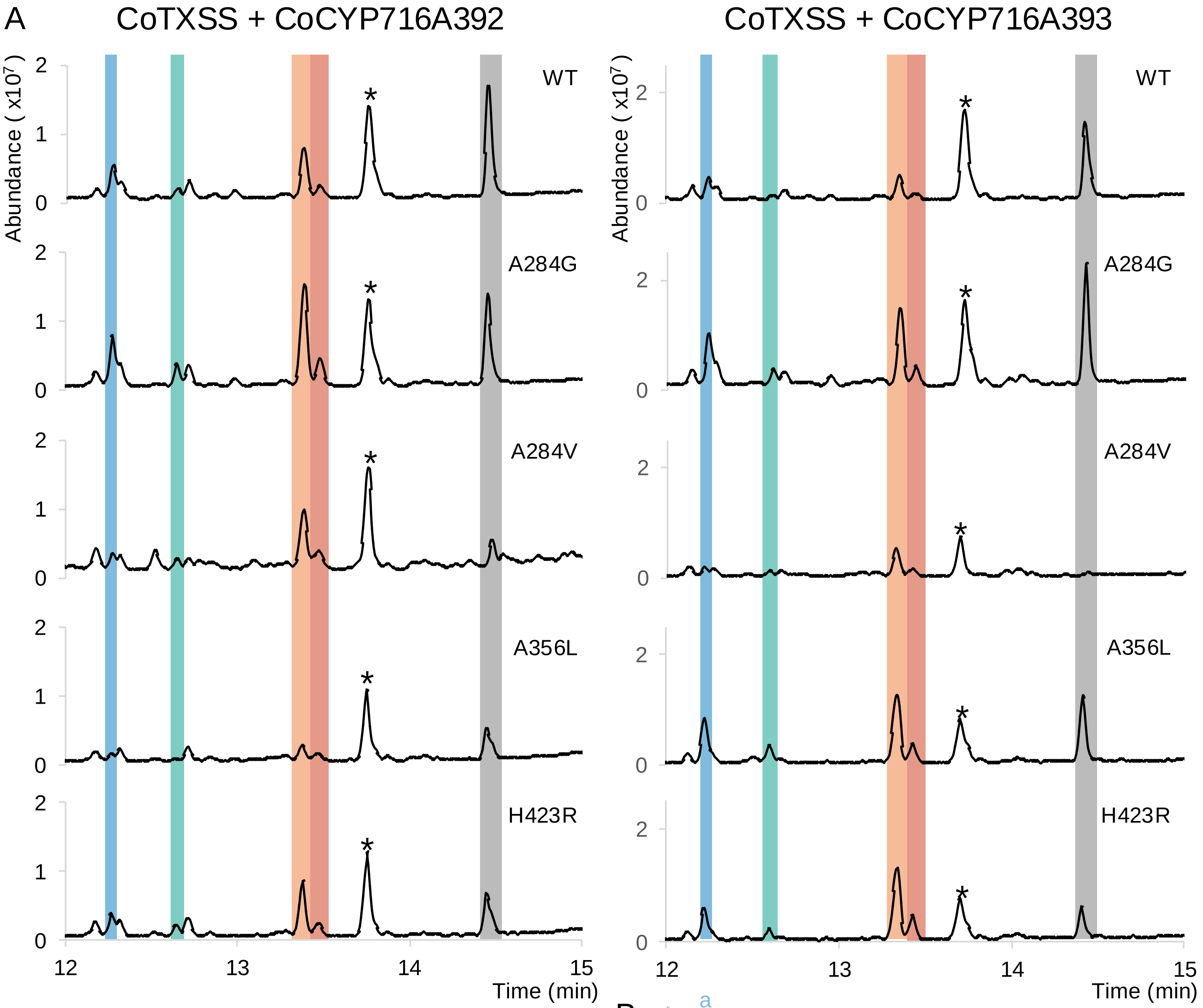

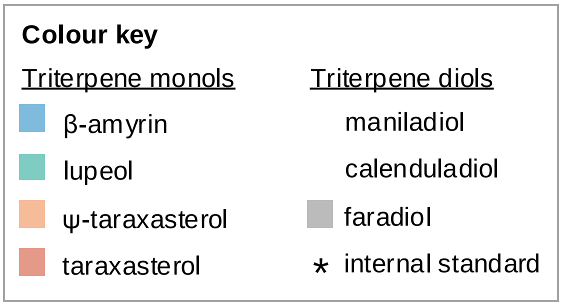

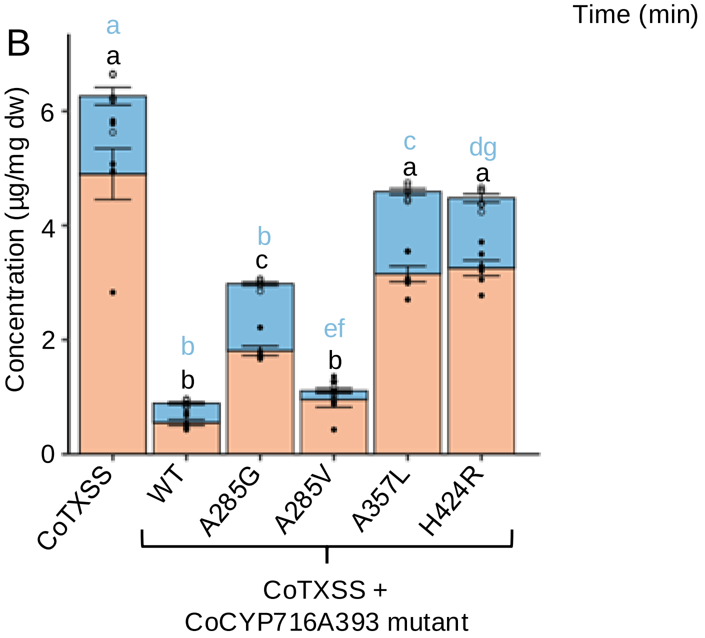

**Supplementary Figure 30. GC-MS analysis of *Nicotiana benthamiana* expressing cytochrome P450 mutants. A.** Total ion chromatograms of extracts of *N. benthamiana* leaves transiently co-expressing *CoTXSS with wild type and mutated Calendula officinalis* P450s (CoCYP716A392 and CoCYP716A393).

**B.** Quantification of triterpenes depleted by wild type and mutated CoCYP716A392; n=6; error bars indicate standard error. Significant differences in total Ψ-taraxasterol/β-amyrin content compared to wild type CoCYP716A393 (black lowercase letters), and significant differences in taraxasterol/β-amyrin ratio compared to wild type CoCYP716A393 (blue lowercase letters) were analysed using a Kruskal-Wallis test followed by post-hoc Wilcoxon rank sum test with a Benjamini-Hochberg correction (Supplementary Table 11). Samples that do not share the same lower-case letter are significantly different from each other (p<0.05).

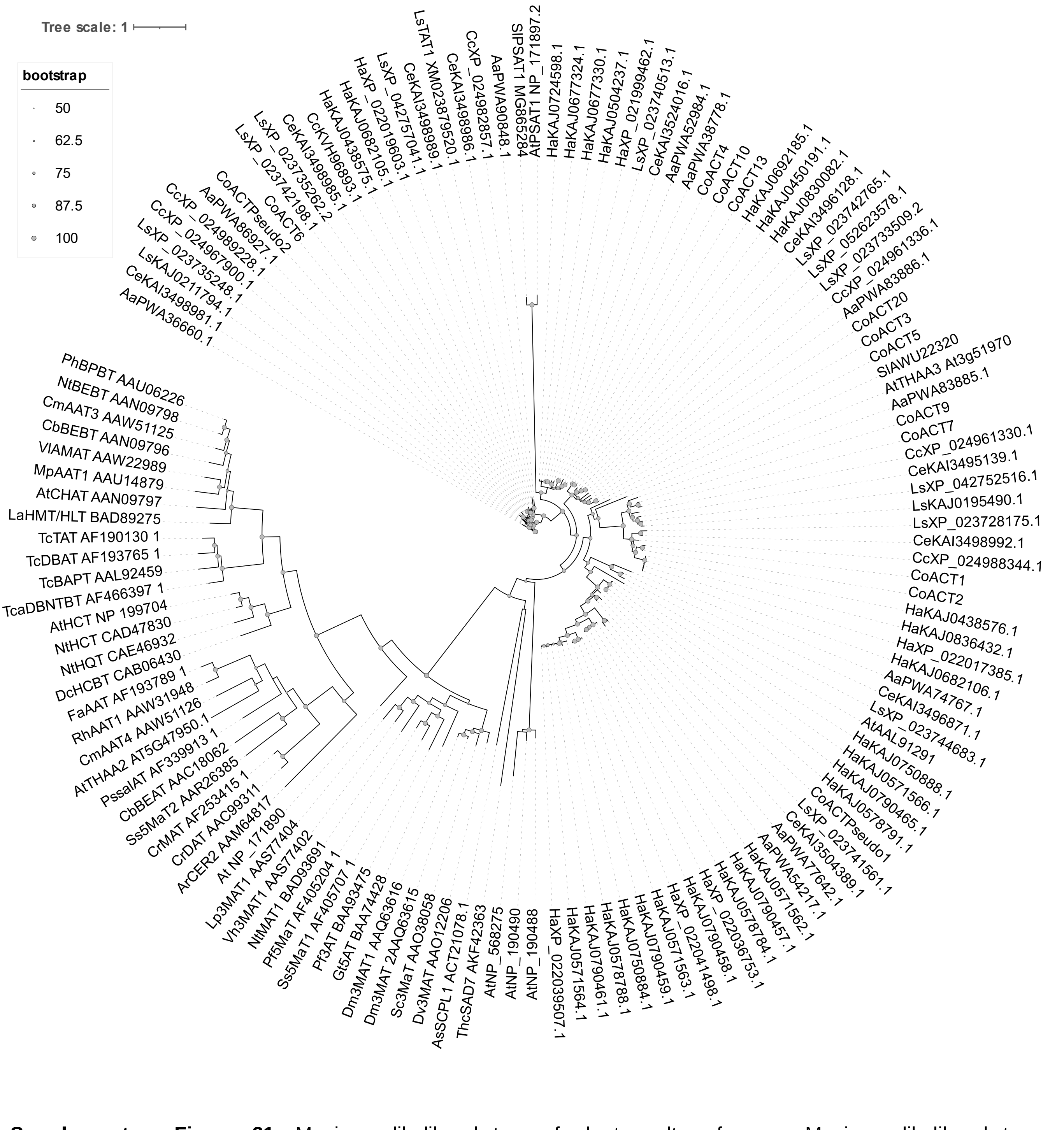

**Supplementary Figure 31.** Maximum-likelihood tree of plant acyltransferases. Maximum-likelihood tree of characterised plant ACTs constructed in IQTree using the JTT matrix-based model. Taxa annotated with an asterisk are likely pseudogenes. Filled grey circles indicate bootstrap supports for each node. The scale bar represents the number of substitutions per site. Accession numbers are indicated by prefixe indicating the species: Aa, *Artemisia annua*; As, *Avena strigosa*; At, *Arabidopsis thaliana*; Cb, *Clarkia breweri;* Cc, *Cynara cardunculus* var. scolymus; Ce, *Cichorium endivia*; Cm, *Cucumis melo*; Co, *Calendula officinalis*; Cr, *Catharanthus roseus*; Dc, *Dianthus caryophyllus*; Dm, *Dendranthema x morifolium*; Dv, *Dahlia variabilis;* Fa, *Fragaria x ananassa*; Gt, *Gentiana triflora*; Ha, *Helianthus annuus*; La, *Lupinus albus*; Lp, *Lamium purpureum*; Ls, *Lactuca sativa*; Mp, *Malus domestica*; Nt, *Nicotiana tabacum*; Pf, *Perilla frutescens*; Ph, *Petunia x hybrida*; Ps, *Papaver somniferum*; Rh, Rosa hybrid cultivar; Sc, *Senecio cruentus*; Sl, *Solanum lycopersicum*; Ss, *Salvia splendens*; Tc, *Taxus cuspidata*; Tca, *Taxus canadensis*; Thc, *Theobroma cacao*; Vh, *Verbena hybrida*; Vl, *Vitis labrusca*.

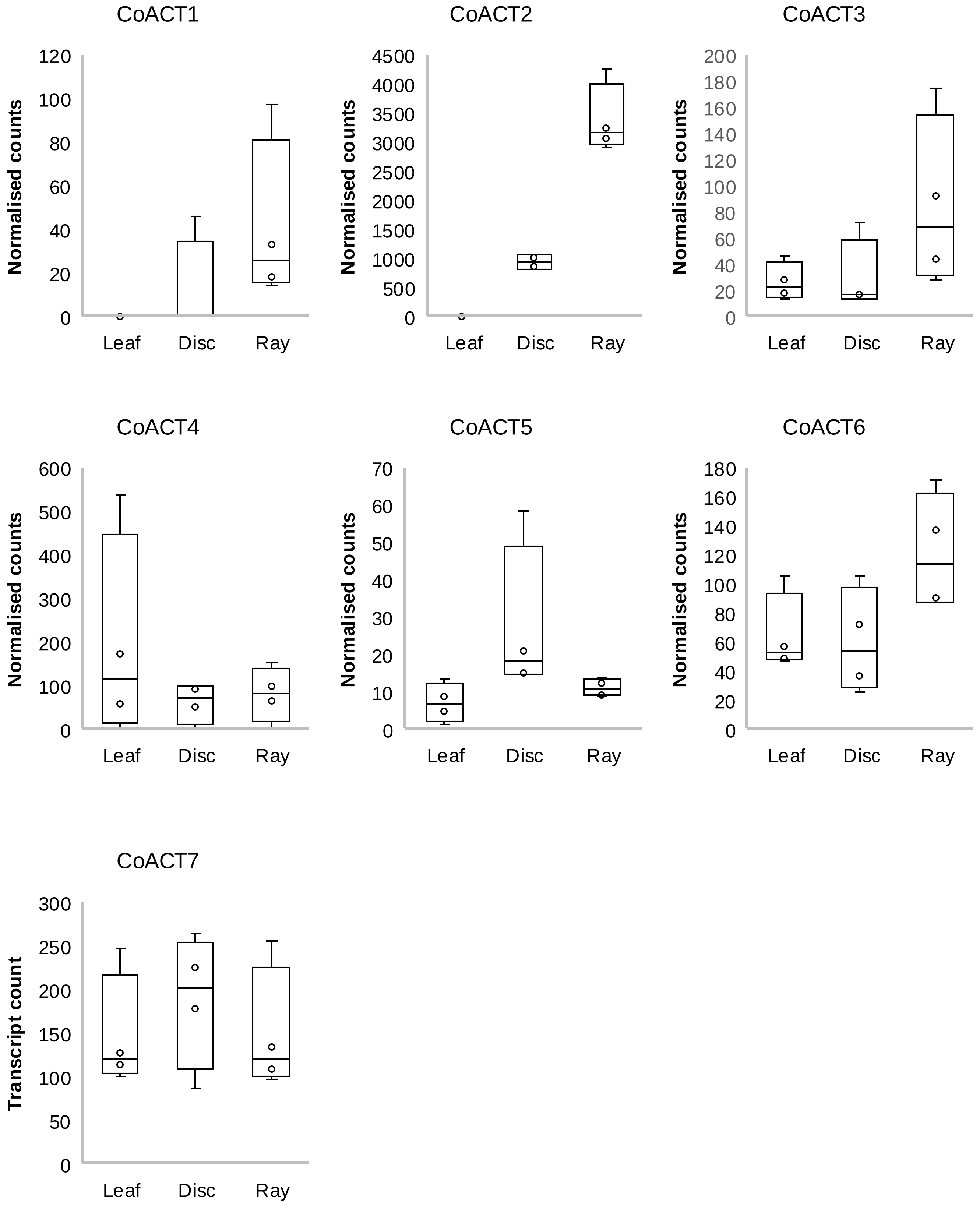

**Supplementary Figure 32. Differential expression of *Calendula officinalis ACYLTRANSFERASE (ACT)* genes.** Gene expression of seven *ACT* genes expressed in leaf, disc and ray floret tissues. Gene expression for each tissue is displayed in a box plot as normalised counts following transcript quantification using Salmon v1.2.0 and differential gene expression analysis using DESeq2. The centre line of the boxplot denotes the median, the box denotes the 25th to 75th percentiles of the data and the whiskers show the minimum and maximum of the data.

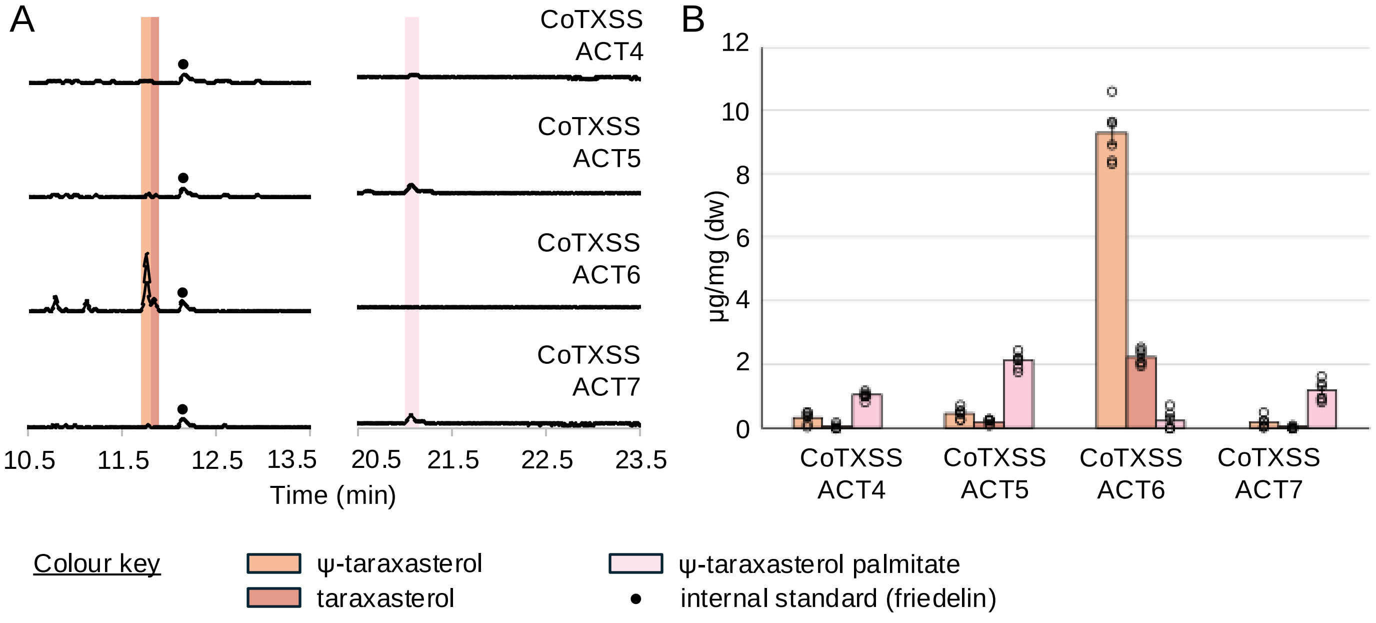

**Supplementary Figure 33. Characterisation of pot marigold triterpene acyl transferases.**
**A.** Total ion chromatograms of extracts of *N. benthamiana* leaves transiently co-expressing CoTXSS with CoACT4-7. **B.** Peak area analysis showing products of pathway reconstruction in *N. benthamiana* through agroinfiltration of indicated genes, quantified compared to friedelin internal standard, n=6, error bars represent standard error.

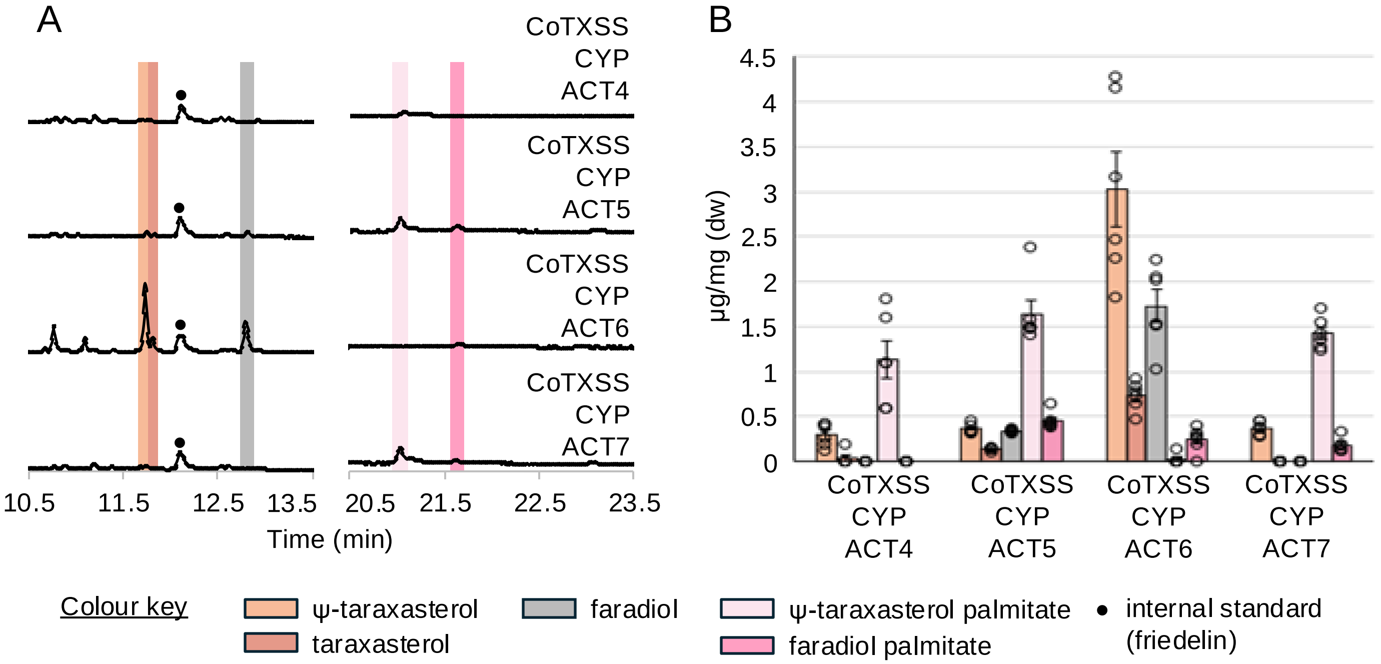

**Supplementary Figure 34. Characterisation of hydroxylated triterpene acyl transferases.**
**A.** Total ion chromatograms of extracts of *N. benthamiana* leaves transiently co-expressing CoTXSS and CoCYP716A392 with CoACT4-7. **B.** Peak area analysis showing products of pathway reconstruction in *N. benthamiana* through agroinfiltration of indicated genes, quantified compared to friedelin internal standard, n=6, error bars represent standard error.

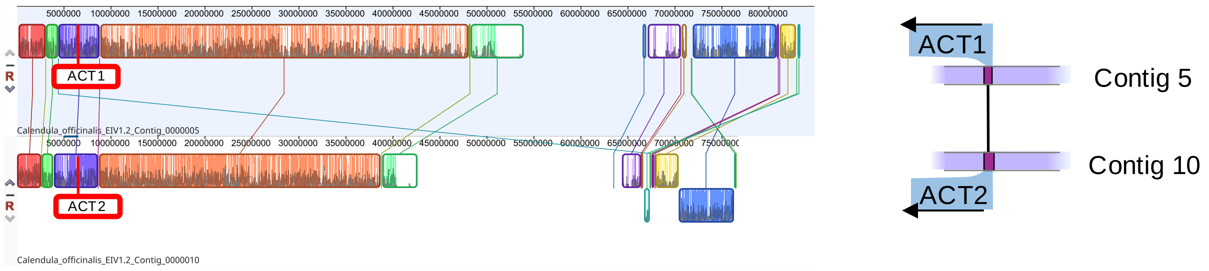

**Supplementary Figure 35. Genomic location and synteny of *Calendula officinalis* (pot marigold) *ACYLTRANSFERASES* (*ACTs*).** A graphical representation of the relative genomic position, orientation and similarity of *CoACT1* and *CoACT2*, which add fatty acid groups to faradiol C3.

**Supplementary Table 1. Triterpenes detected in *Calendula officinalis.***

Content and distribution of triterpenes detected in extracts of leaf, disc floret, ray floret and root tissue. Metabolite concentrations (µg/mg dry weight) were determined by GC-MS peak area analysis using friedelin as an internal standard. Data are means ± SD of 4 independent plants. n.d. = not detected.

| Metabolite | Leaf | Disc floret | Ray floret | Root |
|---|---|---|---|---|
| µg/mg dry weight ± S.D. | | | | |
| **Triterpene scaffold** | | | | |
| β-amyrin | 0.02 ± 0.001 | 0.246 ± 0.069 | 1.965 ± 0.443 | n.d. |
| isofucosterol | 0.052 ± 0.015 | 0.33 ± 0.02 | 1.068 ± 0.303 | n.d. |
| α-amyrin | 0.018 ± 0.003 | 0.082 ± 0.029 | 1.126 ± 0.213 | n.d. |
| lupeol | 0.02 ± 0.002 | 0.056 ± 0.021 | 1.46 ± 0.513 | n.d. |
| ψ-taraxasterol | n.d. | 0.209 ± 0.065 | 4.645 ± 1.104 | n.d. |
| taraxasterol | n.d. | n.d. | 0.848 ± 0.244 | n.d. |
| **Hydroxylated triterpenes** | | | | |
| faradiol | n.d. | 0.07 ± 0.006 | 0.294 ± 0.071 | n.d. |
| **Triterpene fatty acid esters** | | | | |
| faradiol/arnidiol laurate | n.d. | 0.255 ± 0.231 | 2.486 ± 0.186 | n.d. |
| β-amyrin myristate | n.d. | 0.062 ± 0.06 | 1.311 ± 0.161 | n.d. |
| manaladiol myristate | n.d. | 0.046 ± 0.049 | 0.401 ± 0.098 | n.d. |
| calenduladiol myristate | n.d. | 0.062 ± 0.072 | 1.282 ± 0.197 | n.d. |
| ψ-taraxasterol/taraxasterol palmitate | n.d. | n.d. | 0.298 ± 0.143 | n.d. |
| faradiol/arnidiol myristate | n.d. | 2.257 ± 2.201 | 13.178 ± 1.171 | n.d. |
| β-amyrin palmitate | n.d. | 0.059 ± 0.064 | 1.769 ± 0.307 | n.d. |
| lupeol palmitate | n.d. | 0.094 ± 0.076 | 0.735 ± 0.225 | n.d. |
| manaladiol palmitate | n.d. | n.d. | 0.505 ± 0.132 | n.d. |
| calenduladiol palmitate | n.d. | n.d. | 0.782 ± 0.168 | n.d. |
| ψ-taraxasterol/taraxasterol palmitate | n.d. | 0.595 ± 0.565 | 1.125 ± 0.983 | n.d. |
| faradiol/arnidiol palmitate | n.d. | 1.082 ± 1.163 | 12.463 ± 0.569 | n.d. |

**Supplementary Table 2. NMR data for faradiol palmitate.** Notes: [a]Assignments of C34 – C43 can be interchanged. [b]Only six resonances of palmitoyl residue were listed in Ezzat et al (2017) Nat Prod Res. 31(6):676-680, with assignment of C33 – C36 being incorrect. [b]Carbon resonances of the palmitoyl residue were not assigned Zitterl-Eglseer et al (1997) J Ethnopharmacol. 57(2):139-44. [d]Group of six unassigned signals at 29.6 – 29.2 ppm reported in Zitterl-Eglseer et al (1997) J Ethnopharmacol. 57(2):139-44 corresponds to C34-C43 with some resonances apparently overlapping; all assignments in that group can be interchanged.

| No | $\delta_H$ (Multiplicity, *J*) | $\delta_C$ | HSQC | HMBC | COSY |
|---:|---:|---:|---:|---:|---:|
| 1a | 1.03 (m) | 38.4 | - | - | 1[b] |
| 1b | 1.7 (m) | 38.4 | - | 10 | 1[a] |
| 2 | 1.63 (m) | 23.7 | 2 | - | 3 |
| 3 | 4.48 (dd, 11.3,5.2 Hz) | 80.6 | 3 | 2b, 23, 24, 31 | 2[b], 2[a] |
| 4 | – | 37.8 | - | - | - |
| 5 | 0.80 (d, 8.4 Hz) | 55.4 | 5 | 6[b] | 6a |
| 6a | 1.41 (m) | 18.2 | 6 | - | 5 |
| 6b | 1.53 (m) | 18.2 | 6 | - | - |
| 7 | 1.41 (m) | 34.1 | 7 | - | - |
| 8 | – | 41.1 | - | 26, 27 | - |
| 9 | 1.31 (m) | 49.9 | 9 | - | - |
| 10 | – | 37.0 | - | 1[b], 25 | - |
| 11a | 1.23 (m) | 21.6 | 11 | - | 11[b] |
| 11b | 1.55 (m) | 21.6 | 11 | - | 11[a] |
| 12a | 1.23 (m) | 27.2 | 12 | - | - |
| 12b | 1.62 (m) | 27.2 | 12 | - | - |
| 13 | 1.62 (m) | 38.7 | 13 | - | - |
| 14 | – | 42.5 | - | 15[a], 26, 27 | - |
| 15a | 1.33 (m) | 36.5 | 15 | 13,14, 17 | 16 |
| 15b | 1.63 (m) | 36.5 | 15 | - | 16 |
| 16 | 3.44 (dt, 10.8,5.0 Hz) | 76.4 | 16 | 28 | 15[b], 15[a] |
| 17 | – | 39.9 | - | 15a, 28 | - |
| 18 | 1.05 (s) | 47.5 | 18 | 28 | - |
| 19 | 1.65 (m) | 35.9 | 19 | - | - |
| 20 | – | 139.8 | - | 22b, 22a, 29 | - |
| 21 | 5.31 (d, 7.2 Hz) | 118.3 | 21 | - | 22b, 22[a] |

| | | | | | |
|---|---|---|---|---|---|
| 22a | 1.68 (m) | 37.5 | 22 | 20,21, 28 | 21,22[b] |
| 22b | 2.09 (m) | 37.5 | 22 | 20,21, 28 | 21,22[a] |
| 23 | 0.85 (s) | 28.0 | 23 | 3, 5 | - |
| 24 | 0.85 (s) | 16.6 | 24 | 3, 5 | - |
| 25 | 0.88 (s) | 16.4 | 25 | 1[b],5, 10 | - |
| 26 | 1.05 (s) | 16.0 | 26 | 7[b], 8, 14 | - |
| 27 | 1.00 (m) | 16.4 | 27 | 8, 13, 14, 15[b], 19 | - |
| 28 | 0.73 (s) | 11.7 | 28 | 16, 17, 22[b] | - |
| 29 | 1.00 (m) | 22.4 | 29 | 19,20 | - |
| 30 | 1.64 (s) | 21.6 | 30 | - | - |
| 31 | – | 173.7 | - | 3, 32b, 32a | - |
| 32 | 2.29 (td, 7.3,1.2 Hz) | 34.9 | 32 | 31, 33 | 33 |
| 33 | 1.25 (m) | 25.2 | 33 | - | 32b, 32[a] |
| 34 | 1.34-1.21 (m) | 29.71[a] | - | - | - |
| 35 | 1.34-1.21 (m) | 29.70[a] | - | - | - |
| 36 | 1.34-1.21 (m) | 29.68[a] | - | - | - |
| 37 | 1.34-1.21 (m) | 29.67[a] | - | - | - |
| 38 | 1.34-1.21 (m) | 29.65[a] | - | - | - |
| 39 | 1.34-1.21 (m) | 29.6[a] | - | - | - |
| 40 | 1.34-1.21 (m) | 29.5[a] | - | - | - |
| 41 | 1.34-1.21 (m) | 29.4[a] | - | - | - |
| 42 | 1.34-1.21 (m) | 29.3[a] | - | - | - |
| 43 | 1.34-1.21 (m) | 29.2[a] | - | - | - |
| 44 | 1.25 (m) | 31.9[a] | 44 | 45 | - |
| 45 | 1.29 (m) | 22.7[a] | 45 | - | 46 |
| 46 | 0.88 (t, 6.9 Hz) | 14.1 | 46 | 45 | 45 |

**Supplementary Table 3. Comparison of experimental and literature assignment of NMR data for faradiol palmitate.** Notes: [a]Assignments of C34 – C43 can be interchanged. [b]Only six resonances of palmitoyl residue were listed in Ezzat et al (2017) Nat Prod Res. 31(6):676-680, with assignment of C33 – C36 being incorrect. [b]Carbon resonances of the palmitoyl residue were not assigned Zitterl-Eglseer et al (1997) J Ethnopharmacol. 57(2):139-44. [d]Group of six unassigned signals at 29.6 – 29.2 ppm reported in Zitterl-Eglseer et al (1997) J Ethnopharmacol. 57(2):139-44 corresponds to C34-C43 with some resonances apparently overlapping; all assignments in that group can be interchanged.

| | δ $_H$ (Multiplicity, *J*) | | δ $_C$ | | |
|---|---|---|---|---|---|
| | Experimental | Lit.[1] | Experimental | Lit.[1] | Lit.[2] |
| 1a | 1.03 (m) | 1.02 | 38.4 | 38.4 | 38.8 |
| 1b | 1.7 (m) | 1.71 | – | – | – |
| 2a | 1.63 (m) | 1.62 | 23.7 | 23.7 | 27.2 |
| 3 | 4.48 (dd, 11.3,5.2 Hz) | 4.44 (dd, 5.5,11) | 80.6 | 80.7 | 80.6 |
| 4 | – | – | 37.8 | 37.9 | 38.5 |
| 5 | 0.80 (d, 8.4 Hz) | 0.8 | 55.4 | 55.4 | 55.5 |
| 6a | 1.41 (m) | 1.4 | 18.2 | 18.2 | 18.2 |
| 6b | 1.53 (m) | 1.52 | – | – | – |
| 7 | 1.41 (m) | 1.4 | 34.1 | 34 | 34.2 |
| 8 | – | – | 41.1 | 41.1 | 41.1 |
| 9 | 1.31 (m) | 1.3 | 49.9 | 49.9 | 50 |
| 10 | – | – | 37 | 37.3 | 37.8 |
| 11a | 1.23 (m) | 1.23 | 21.6 | 21.5 | 22.6 |
| 11b | 1.55 (m) | 1.54 | – | – | – |
| 12a | 1.23 (m) | 1.22 | 27.2 | 27.1 | 28 |
| 12b | 1.62 (m) | 1.62 | – | – | – |
| 13 | 1.62 (m) | 1.6 | 38.7 | 38.7 | 37.6 |
| 14 | – | – | 42.5 | 42.3 | 42.5 |
| 15a | 1.33 (m) | 1.32 | 36.5 | 36.5 | 37 |
| 15b | 1.63 (m) | 1.64 | – | – | – |
| 16 | 3.44 (dt, 10.8,5.0 Hz) | 3.42 (dd, 5,12) | 76.4 | 76.9 | 76.2 |
| 17 | – | – | 39.9 | 39.9 | 39.9 |
| 18 | 1.05 (s) | 1.04 | 47.5 | 47.6 | 47.7 |
| 19 | 1.65 (m) | 1.65 | 35.9 | 35.7 | 35.9 |
| 20 | – | – | 139.8 | 139.8 | 139.6 |
| 21 | 5.31 (d, 7.2 Hz) | 5.29 (d, 7) | 118.3 | 118.3 | 118.4 |

| | | | | | |
|---|---|---|---|---|---|
| 22a | 1.68 (m) | 1.68 | 37.5 | 37.5 | 36.5 |
| 22b | 2.09 (m) | 2.1 | – | – | – |
| 23 | 0.85 (s) | 0.84 (s) | 28.0 | 28.1 | 29.2 |
| 24 | 0.85 (s) | 0.84 (s) | 16.6 | 16.5 | 14 |
| 25 | 0.88 (s) | 0.87 (s) | 16.4 | 16.4 | 16.3 |
| 26 | 1.05 (s) | 1.05 (s) | 16.0 | 15.9 | 16 |
| 27 | 1.00 (m) | 0.99 (s) | 16.4 | 16.3 | 16.5 |
| 28 | 0.73 (s) | 0.72 (s) | 11.7 | 11.3 | 11.6 |
| 29 | 1.00 (m) | 1.00 (d,6.3) | 22.4 | 22.4 | 22.4 |
| 30 | 1.64 (s) | 1.64 | 21.6 | 21.5 | 21.6 |
| 31 | – | – | 173.7 | 171 | 173.6c |
| 32 | 2.29 (td, 7.3,1.2 Hz) | 2.29 (t, 7 Hz) | 34.9 | 34.8 | 34.8c |
| 33 | 1.25 (m) | n.d. | 25.2 | 29.5b | 25.1c |
| 34 | 1.34-1.21 (m) | n.d. | 29.71[a] | 30.17[b] | n.d.e |
| 35 | 1.34-1.21 (m) | n.d. | 29.70[a] | 31.2[b] | n.d.e |
| 36 | 1.34-1.21 (m) | n.d. | 29.68a | 14.17[b] | n.d.e |
| 37 | 1.34-1.21 (m) | n.d. | 29.67[a] | n.d. | n.d.e |
| 38 | 1.34-1.21 (m) | n.d. | 29.65a | n.d. | 29.6[c,d] |
| 39 | 1.34-1.21 (m) | n.d. | 29.6[a] | n.d. | 29.5[c,d] |
| 40 | 1.34-1.21 (m) | n.d. | 29.5a | n.d. | 29.4[c,d] |
| 41 | 1.34-1.21 (m) | n.d. | 29.4[a] | n.d. | 29.3[c,d] |
| 42 | 1.34-1.21 (m) | n.d. | 29.3a | n.d. | 29.2[c,d] |
| 43 | 1.34-1.21 (m) | n.d. | 29.2[a] | n.d. | 29.1[c,d] |
| 44 | 1.25 (m) | n.d. | 31.9a | n.d. | 31.8c |
| 45 | 1.29 (m) | n.d. | 22.7[a] | n.d. | 23.7c |
| 46 | 0.88 (t, 6.9 Hz) | n.d. | 14.1 | n.d. | 16.3c |

**Supplementary Table 4.** Statistical tests used in Figure 1. *=P<0.05, ** P<0.01,*** P<0.0001

| Experiment | Sample | T-value | P value |
|---|---|---|---|
| ELISA on THP cells for IL6 in response to *C. officinalis* extract (One-way ANOVA (N=4) with a Post-hoc Dunnetts test. | LPS+DMSO-BAY | -10.091 | <0.0001 (***) |
| | LPS+DMSO- C. officinalis extract | -6.995 | <0.0002 (***) |
| | LPS+DMSO- Basal | -11.096 | <0.0001 (***) |
| ELISA on THP cells for TNF in response to *C. officinalis* extract (One-way ANOVA (N=4) with a Post-hoc Dunnetts test. | LPS+DMSO-BAY | -12.318 | <0.001 (***) |
| | LPS+DMSO- C.officinalis extract | -6.303 | <0.001 (***) |
| | LPS+DMSO- Basal | -13.599 | <0.001 (***) |
| ELISA on THP cells for IL6 with fractions of *C. officinalis* extract (One-way ANOVA (N=4) with a Post-hoc Dunnetts test. | LPS+DMSO-Basal | -8.939 | <0.001 (***) |
| | LPS+DMSO-BAY | -8.918 | <0.001 (***) |
| | LPS+DMSO- Fraction 1 | -3.306 | 0.02336 (*) |
| | LPS+DMSO- Fraction 2 | -1.602 | 0.51842 |
| | LPS+DMSO- Fraction 3 | -2.470 | 0.12804 |
| | LPS+DMSO- Fraction 4 | -2.347 | 0.16083 |
| | LPS+DMSO- Fraction 5 | -3.038 | 0.04141 (*) |
| | LPS+DMSO- Fraction 6 | -4.344 | 0.00241 (**) |
| | LPS+DMSO- Fraction 7 | -4.108 | 0.00395 (**) |
| ELISA on THP cells for IL6 with 20 µM of pure compound (One-way ANOVA (N=4) | LPS+DMSO- Basal | -12.639 | <0.0001 (***) |
| | LPS+DMSO- BAY | -12.519 | <0.0001 (***) |

| | | | |
|---|---|---|---|
| with a Post-hoc Dunnetts test. | LPS+DMSO- Psitaraxasterol | -4.283 | 0.001377 (**) |
| | LPS+DMSO- taraxasterol | 10.829 | <0.0001 (***) |
| | LPS+DMSO- Faradiol | -7.526 | <0.0001 (***) |
| | LPS+DMSO- Arnidiol | -7.752 | <0.0001 (***) |
| | LPS+DMSO- Faradiol myristate | -6.072 | <0.0001 (***) |
| | LPS+DMSO- Faradiol palmitate | -5.970 | <0.0001 (***) |
| | LPS+DMSO- Faradiol myristate/faradiol palmitate | -4.991 | 0.000167 (***) |

**Supplementary Table 5.** Table of genome assembly statistics

| **Statistics** | **Final Calendula officinalis genome**<br>Calendula_officinalis_EIV1.2 |
|---|---|
| Genome size | 1,301,655,687 |
| GC content | 36.5% |
| Contigs | 2,811 |
| Largest contig | 104,728,146 |
| N50 | 80.2 Mb |
| Complete BUSCOs | 2256 (96.9%) |
| Single Copy BUSCOs | 586 (25.2%) |
| Duplicated BUSCOs | 1670 (71.8%) |
| Fragmented BUSCOs | 10 (0.4%) |
| Missing BUSCOs | 60 (2.6%) |

**Supplementary Table 6. Table of candidate *oxidosqualene cyclase* genes (*OSCs*) identified in the *Calendula officinalis* genome.** Genes for which transcripts were present in the transcriptome datasets are shaded in grey. Corresponding gene and transcript IDs from the functional annotation of the genome and transcriptome respectively are shown. Manually curated CDS sequences for all *OSC* genes are provided in Supplementary Table 8. **CoOSC28* and *CoOSC2* are from haplotypes that were collapsed to generate the final genome assembly. *CoOSC28* is the sequence found in the genome and *CoOSC2* represents the sequence observed in the transcriptome.

| Gene name | Length (bp) | Gene ID from functional annotation (CALOF41496_EIv1.0) | Transcript ID from transcriptome (EIV1.1.mikado.loci.cds) | Predicted gene function | Predicted to be functional / non-functional | Sequence features | Putative homeologue |
|---|---|---|---|---|---|---|---|
| *CoOSC1/ CoTXSS* | 2331 | 0016890.1 | 0435770.1 | Taraxasterol synthase | Functional | - | **CoOSC17** |
| *CoOSC2* | 2280 | - | 0275220.1* | Mixed amyrin synthase | Functional | Sequence in transcriptome, from alt contig in genome | **CoOSC18** |
| *CoOSC3/ CoMAS* | 2274 | 0132450.1 | 0109510.1 | Mixed amyrin synthase | Functional | - | **CoOSC23** |
| *CoOSC4* | 2256 | 0450140.1 | 0055930.1 | Bauerenol synthase | Functional | - | **CoOSC24** |
| *CoOSC5* | 2265 | 0132490.1 | 0279660.1 | Novel triterpene synthase | Functional | - | CoOSC20 |
| *CoOSC6* | 2277 | 0132510.1 | 0762650.1 | Novel triterpene synthase | Functional | - | - |
| *CoOSC7* | 2070 | 0830400.1 | 0553310.1 | Lupeol synthase | Non-functional | Missing exon 1 | **CoOSC25** |
| *CoOSC8* | 2280 | 0214270.1 | 0142220.1 | Shionone synthase | Functional | - | CoOSC21/CoOSC22 |
| *CoOSC9* | 2289 | 0921050.1 | 0358550.1 | β-amyrin synthase-like | Functional | - | **CoOSC10** |
| *CoOSC10* | 2292 | 0776440.1 | 0553470.1 | β-amyrin synthase-like | Functional | - | **CoOSC9** |
| *CoOSC11* | 2280 | 0776460.1 | 0264370.1 | β-amyrin synthase | Functional | - | - |
| *CoOSC12* | 2265 | 0625860.1 | 0025710.1 | Cycloartenol synthase | Functional | - | **CoOSC14/**CoOSC16 |
| *CoOSC13* | 2286 | 0710970.1 | 0135220.1 | Cycloartenol synthase | Functional | - | **CoOSC15** |
| *CoOSC14* | 2268 | 0171410.1 | 0433710.1 | Cycloartenol synthase | Functional | - | **CoOSC12** |
| *CoOSC15* | 2277 | 0147700.1 | 0546580.1 | Cycloartenol synthase | Functional | - | **CoOSC13** |
| *CoOSC16* | 2280 | 0625850.1 | 0553310.1 | Cycloartenol synthase | Functional | - | CoOSC12 |
| *CoOSC17* | 1998 | 0840750.1 | - | Taraxasterol synthase | Non-functional | Multiple small and large deletions | **CoOSC1** |
| *CoOSC18* | 2247 | 0841920.1 | - | Mixed amyrin synthase | Non-functional | Small deletion, 2 in-frame STOP codons | **CoOSC2/CoOSC28*** |

| | | | | | | | |
|---|---|---|---|---|---|---|---|
| *CoOSC19* | 2280 | 0841840.1 | - | Novel triterpene synthase | Unknown | DCTSE motif, small substitutions | - |
| *CoOSC20* | 2265 | 0132500.1 | - | Novel triterpene synthase | Unknown | Small substitutions | CoOSC5 |
| *CoOSC21* | 2280 | 0214250.1 | - | Shionone synthase | Non-functional | DCTAD motif, in-frame STOP codons | CoOSC8/CoOSC22 |
| *CoOSC22* | 2277 | 0214200.1 | - | Shionone synthase | Unknown | DCTAY motif, small substitutions | CoOSC8/CoOSC21 |
| *CoOSC23* | 756 + 1692 | 0697250.1 + 0697210.1 | - | Mixed amyrin synthase | Non-functional | Gene split and inverted in genome, in-frame STOP codon | **CoOSC3** |
| *CoOSC24* | 2010 | 0318520.1 | - | Bauerenol synthase | Non-functional | Multiple short deletions | **CoOSC4** |
| *CoOSC25* | 1998 | 0027700.1, 0027710.1 | - | Lupeol synthase | Non-functional | Missing exon 1, multiple short deletions | **CoOSC7** |
| *CoOSC26* | 2280 | 0893670.1 | - | Lanosterol synthase | Non-functional | Few deletions, substantial substitutions | **CoOSC27** |
| *CoOSC27* | 2268 | 0752710.1 | - | Lanosterol synthase | Unknown | - | **CoOSC26** |
| *CoOSC28* | 2274 | 0015710.1* | - | Mixed amyrin synthase | Functional | - | **CoOSC18** |

**Supplementary Table 7. Table of *Calendula officinalis oxidosqualene cycle* (*OSC*) transcripts.** *OSC* genes identified in the *Calendula officinalis* transcriptome (Calendula_officinalis_EIV1.1.mikado.loci.cds.fa) and the *Calendula arvensis* transcriptome (CombineCarv.combined.cdhit98.fa.transdecoder.cds.cds). Genes that were functionally characterised have their functions shown in bold, predicted functions are in normal text.

| **Species** | **Gene name** | **Transcript number** | **Characterised/**predicted function |
|---|---|---|---|
| *Calendula officinalis* | CoOSC1 | 0435770.1 | **Taraxasterol synthase** |
| *Calendula officinalis* | CoOSC2 | 0275220.1 | Mixed-amyrin synthase |
| *Calendula officinalis* | CoOSC3 | 0109510.1 | **Mixed-amyrin synthase** |
| *Calendula officinalis* | CoOSC4 | 0055930.1 | **Bauerenol synthase** |
| *Calendula officinalis* | CoOSC5 | 0279660.1 | Novel/multifunctional triterpene synthase |
| *Calendula officinalis* | CoOSC6 | 0762650.1 | Novel/multifunctional triterpene synthase |
| *Calendula officinalis* | CoOSC7 | 0553310.1 | Lupeol synthase |
| *Calendula officinalis* | CoOSC8 | 0142220.1 | **Shionone synthase** |
| *Calendula officinalis* | CoOSC9 | 0358550.1 | β-amyrin-like synthase |
| *Calendula officinalis* | CoOSC10 | 0553470.1 | β-amyrin-like synthase |
| *Calendula officinalis* | CoOSC11 | 0264370.1 | β-amyrin synthase |
| *Calendula officinalis* | CoOSC12 | 0025710.1 | Cycloartenol synthase |
| *Calendula officinalis* | CoOSC13 | 0135220.1 | Cycloartenol synthase |
| *Calendula officinalis* | CoOSC14 | 0433710.1 | Cycloartenol synthase |
| *Calendula officinalis* | CoOSC15 | 0546580.1 | Cycloartenol synthase |
| *Calendula officinalis* | CoOSC16 | 0553310.1 | Cycloartenol synthase |
| *Calendula arvensis* | CarOSC1 | DN42097_c0_g2_i3 | **Taraxasterol synthase** |

**Supplementary Table 8. Table of coding and protein sequences of *Calendula officinalis oxidosqualene cyclases*.** DNA sequences that were identified from the transcriptome have the suffix _CDS, and those that were identified from the *C. officinalis* genome have the suffix _CDS(gDNA). Protein sequences are a direct translation of the CDS sequences and were subsequently used in the phylogenetic analysis.

| | |
|---|---|
| >CoOSC1_CDS | ATGTGGAAATTAAAAATTGGTGAAAAAAATGGAAAGCTGGAAATTGGTGACGGAAATGGCGATGAATATTTGT<br>ATAGTACCAACAATTTCGTAGGGAGACAAACATGGGAGTTTGATCCAAGTGCAGGCACACAGGAAGAACGTGA<br>CCAAATCGAAAGGATTCGAGAGCAATTCTTGAACAATAAGAAGAAACTCGATATCCATTGTTGTGGTGACTTG<br>CTCATGCGAACCCAGCTTATCAAGGAAAGCCAGATTGATCTTACTAGCGAACTTCTAGTGAGACTCAAAGATG<br>AAGAAGATGTTAACTATGAAGCTGTGACAATGGCGGTAAAAAAAGCGGTCCTCTTGAACCGTGCAATTCAAGC<br>TTGGGACGGTCATTGGCCAGCTGAAAATGCCGGTCCCCTTTTTTTTACCCCTCCTCTGATAATTGCCTTGTAC<br>ATTAGTGGTACAGTGGATACAATTTTAACAAAAGAACACCAAAAGGAGATGATTCGCTACATGTACATCCATC<br>AGAATGAAGATGGAGGATGGGGATTCTATATATCGGGGAAAAGCACGATGATTGGAAGTGCATTGAACTATGT<br>GGCTCTAAGACTTCTTGGAGAAGCTTCACCAACTGATGATGATGACATTGGTGCTCTTGCTAGAGGCCATAAA<br>TGGATATTTGATCACGGCGGTGCTACATCCATCCCTTCATGGGGCAAGGTTTATCTCGCGGTGCTTGGGGTAT<br>ATGAATGGGAAGGCTGCAATCCTCTGCCTCCTGAATTTTGGCTATTTCCTTCCTTTTTGCCTTATCATCCAGC<br>AAAAATGTGGTGTTACTGTAGAACGACATACATGCCGATGTCACACTTGTATGGTAAAAGCTACCATGGGCCT<br>ATTACTGATCTTGTTTTGTCTTTAAGAAAAGAGATTCACACCCTTCCTTACCACCAGATTAACTGGAACAAAC<br>AACGACATAATTGTTGTAAGTTGGATCTATACTACCCTCATTCTTCCATACAAGATATGTTATGGGATAGTCT<br>GCACTATTTTTGTGAACCACTTGTCAAAAAATGGCCCTTGAATAAGCTAAGAGAGAAGGGTCTCAAAAGAGTG<br>GTTGACCTAATGCGTTATGGATCCGAAGAAGGACGTTACATAGGCATGGGTTGTGTTGACAAGGCTTTACAAA<br>TGATGTGTTTTTATGCCGAGGATCCAAATGGGATTGACTTCAAACACCACCTGGCTAGAGTCCCTGATTACTT<br>GTGGCTGGCAGAAGATGGTATGAAGATGCAAAGTTTCGGTAGTCAGTTATGGGATTGCACTCTTGTAACTCAA<br>GCAATTATGGCCAGTAATATGGTTGATGAGTATGGAGATTCACTCAAAAAAGCCCACTTTTACTTAAAACAGT<br>CACAGATTAAAGAGAACCCTAAAGGCGATTTCACAAAAATGTGTCGTTTATTCACGAAAGGGGCATGGACTTT<br>CTCTGATCAAGACCATGGATGGGTTGTATCAGATTGCACTGCAGAAGCTTTGATTTGTTTATTGGCATTGTCA<br>CAAATGCCACAAGATATTGCGGGCGAAAAGGCTGAAGTTGATAGATTATATGATGCTGTGAATGTCCTCCTTT<br>ACCTACAAAGTCCTGAAAGTGGTGGTTTCGCTATTTGGGAGCCACCAGTTCCAAAACCATATTTACAGATGTT<br>GAACCCTTCAGAACTTTTTGCAGACATAGTGGTTGAGAAAGAGCATGTTGAATGTACTGGATCCATAATCCAA<br>GCATTGAATTCATTCAAAAATCTGCACCCACGACACCGTGAGAAAGAAATAGAAGCTGCTATCGAGAAAGGCA<br>TACGTTTTTTGGAAAACAAACAACAAAATGATGGTTCATGGTACGGTTATTGGGGTATTTGTTTTCTCTATGG<br>CACATTTTTTGTGCTGCAAGGTTTAGTATCATGCGGGAAAACATATGAAAATAGTGAAACCGTTCGAAAAGCT<br>GTCAATTTTTTGCTCTCAACACAGAATTCAGAAGGTGGTTGGGGAGAGAGTTTTGAGTCATGCCCTCAAGAGA<br>AATTCATACCTTTGGAAGGAAATAGAACAAATTTGGTGCAAACTTCATGGGCCATGCTTGGTCTTCTCTATGG<br>TGGACAGGTTGAAAGGGATGTAACACCGTTACACAAGGCAGCAAAATTGCTCATTAATGGACAATTGGATAAT<br>GGAGATTTTCCTCAACAGGAAATAACGGGAGTGTACATGAAAAACTGCATGTTACATTATCCAGAATATAGGA<br>ACACTTTTCCGTTATGGGCGTTAGGAGAATACCGCAACCGTGTTTGGTTGCCAAAGCAACAAGTCTAA |
| >CoOSC2_CDS | ATGTGGAGCTTGAAGATAGCAGAAGGGAATGACCATCCTTACTTGTTTAGCACCAACAATTTTGTTGGTAGGC<br>AGATTTGGGAGTTTGATCCCGATGCTGGTACTTCCGAAGAAAAACAACAAGTCAAAAACGTTCGTCAACATTT<br>TAGAAGCAATCTAAGGGGAGGTGTTCATCCATGCAGTGATTTGCTTATGCGTATGCAGTTGATAAAAGAGAAT<br>GGAATAGATTTATTGAGCATACGACCAGCAAGATTGGGAGAGAATGAGGAAGTGAATTATGAAGCGATAACGA<br>CAGCAGTGAAGAAAGCAGTCCGATTGAACCGTGCGATTCAAGCAAAGGATGGTCATTGGCCTGCTGAAAATGC<br>TGGCTCTATGATTTCCACTCCTGCCCTCCTTATTGCTATGTATATAAGTGGAACCATCAACACGCATTTAACC<br>AAAGAACACAAGACCGAAATGATACGTTATATCTACAACCATCAGAACGAAGATGGAGGGTGGGGATTTTATA<br>TCGGGGGGCAGAGCATCATGATCGGGTCTGCTTTGAGCTATGTAGCCCTAAGGTTACTAGGAGAGGGACCCGA<br>TGATGGAAATGGTGCGGTTGGCCGAGCCAGGAAGTGGATACTTGACCATGGTGGTGCAACCGCCATTCCCTTT<br>TTGGGAAAAATTTATCTCTCGGTTCTTGGAGTATATGAATGGGAAGGATGCAACCCACTGCCACCAGAAATTT<br>GGCTTTCCCCTCAAGCTTTACCATATCATCCATCAAAAATGTTGAGCCATTTCCGGACAACCTATATGCCTAT<br>GTCATACCTGTACGGGAGAAAAATTCATGGTCCAATCACTGATCTTGTTCAACAACTCCGAAATGAAATTCAC<br>GTGATCCCATACAACAATATAAATTGGAATAAACAGCGACATAACTGTTGCAAGGAAGATCTCTACTACCCTC<br>ATTCAACGGTACAAGATCTATTGTGGGATGGTCTTCATTACTTAAGTGAGCCAATTCTTAAGTTTTGGCCTTT<br>TACAAAGTTAAGAGAAAGAGGTCTCAAAAGAGCAGTTGAATTAATGCGATATGGTGCTCAAGAAAGCAGATAT<br>ATCACCACTGGATATATCTCAAAGAGCTTGCAAATAATGTGTTGGTGGGCGGAGAACCCAAATGGGGATGAAT<br>TCAAACGTCATCTTGCTAGAGTGCCTGATTATTTGTGGTTAGCAGAAGATGGAATGAAGATGCAAAGTTTTGG<br>AAGCCAATTATGGGATTGTGCACTTGCAACACAAGCAATAATAGCTAGTGATATGGTCGATGAATATGGCGAT<br>TCACTTAAGAAAGCCAATTTCTACCTCAAAGAATCTCAAATCAAGAATAATCCGAGTGGTGATTTTAGTAAAA<br>TGTGCCGGCAATTTACTAAAGGGTCGTGGACTTTGTCTGACCAAGATCAAGGTTGGGTTGTATCAGACTGCAC<br>AGCTGAAGCGATGAAGTGCCTTTTGTTACTATCCCAAATGCCAGACGAAATTACGGGAGAAAAAGCAGAAACT<br>GAGCGATTATATGAAGCTGTTAATGTCCTTCTTTACTTACAGAGTCCTATAAGTGGAGGTTTTGCTGTTTGGG |

| | |
|---|---|
| | AGCCACCTGTCCCACAACCATATTTACAGATGTTGAATCCTTCAGAACTTTTTGCAGATATTATTGTTGAGAA<br>AGAGCATGTTGAGTGCACTGCATCAGTTATTCAAGCCCTTTTAGCCTTCAACCGGTTGCACCCACAGCACAGA<br>GAGAAAGAAATAGAAATTTCTGTGGCAAAAGCGGTTTCTTTTTTGGAGGAAAAACAACAGCATGATGGTTCAT<br>GGTATGGTTATTGGGGAGTATGCTTTTTATATGGCACATTCTTTGCTATAGGAGGGTTAGAAGCTGCTGGAAA<br>AACATACAAAAACAGTGAAACAATTCGTAAAGCGGTTAATTTTTTTCTTTTGACACAAAATGAAGAAGGTGGT<br>TGGGGAGAAAGCATCAAATCCTGCCCTAGTGAAGTATACTCACCGTTGGATGGAAATCGAACAAATCTAGTTC<br>AAACATCATGGGCTATGCTTGGTCTTATGTTAGGTGGACAGATTGAGAGAGATCCAACACCCTTGCATAAAGC<br>CGCAAAGATATTAATTAATGCACAGATGGATAATGGAGATTTTCCTCAACAGGAAATTACTGGAGTCTACATG<br>AAAAATTGCATGCTACATTATGCGGAATACAGGAACATTTTCCCGCTTTGGGCACTTGGGGAATATCGCAAAC<br>GGGTTTGGGTCAACTAA |
| >CoOSC<br>3_CDS | ATGTGGAAGTTGAAGATAGCAGAAGGGAATGATCCTTACTTATTTAGTACCAACAATTTCGTCGGTAGGCAAA<br>TTTGGGAATTTGATCCCGACGCTGGTACTCCCGAAGAAAAACAAGAAGTTGAAAATGCTCGTCAACATTATAG<br>AAACAATCGAAAGGAAGATGTTCATCCATGCAGTGATTTGCTTATGCGGATGCAGTTGATCAAAGAGAATGGC<br>ATAGATCTACTGAGTATTCCACCAGCAAGGTTAGGAGATAATGAGCAAGTAAATTATGAAGCAGTGACGAGAT<br>CAGTGACGAAAGCAGTTCGATTGAACCGTGCGATTCAAGCAAAGGATGGTCACTGGCCTGCAGAAAACGCTGG<br>CCCTATGTTTTTCACTCCTCCCCTCCTTATTGCTATGTACATCAGTGGAGCCATCGCCACACATTTAACCAAA<br>GAACACAAGACCGAAATGATACGTTATATCTACAACCATCAAAACGATGATGGAGGGTGGGGATTTTATATCG<br>AGGGACACAGTACCATGATCGGGTCTGCTTTGAGCTATGTAGCCCTAAGGTTACTAGGAGAGGGACCTCATGA<br>TGGAAATGGTGCGGTTGACCGAGCCAGAAAGTGGATAATTGACCATGGTGGTGCAACCTCAATTCCCTCATGG<br>GGCAAAACTTATCTATCGGTTCTTGGAGTATATGAATGGGAAGGATGCAATCCACTTCCACCAGAATTTTGGA<br>TTTTCCCTGAAACTTTACCATATCATCCGGCGAAAATGTGGTGCTATTGCCGGACAACATACATGCCTATGTC<br>ATACTTATATGGGACAAAATTTCATGGTCCGATCACTGATCTTGTTCTACAACTTAGACAAGAAATTCATGCG<br>ATCCCGTACAATGAGATAAATTGGAATAAACAACGACATAACTGTTGTAAGGAAGATTTGTATTACCCTCATT<br>CAACGTTGCAAGATCTATTGTGGGATGGACTTCATTACTTAAGTGAACCGTTTCTTAAATATTGGCCTTTTAA<br>AAAGCTAAGAGAAAGAGGTCTCAAAAGAGCAGTTGAATTAATGCGATATGGTGCTCAACAAAGCAGATATATG<br>ACTATTGGATGTGTTGAAAAGAGCTTGCAAATGATGTGTTGGTGGGCAGAGAACCCAAATGGGGATGAATTCA<br>AGCACCATCTTGCTAGAGTGCCTGATTATTTGTGGTTAGCAGAAGATGGAATGAAGATGCAAAGTTTTGGGAG<br>CCAATTATGGGATTGCACACTTGCAACACAAGCAATAATAGCTACTGATATGGTTGAAGAATATGGTGATTCA<br>CTTAAAAAAGCTCATTTTTATATCAAAGAATCTCAAGTTAAACAAAATCCTTCGGGCGATTTTAGTCAAATGT<br>GCAGACAGTTTACTATAGGGTCGTGGACTTTTTCTGACCAAGATCATGGTTGGGTTGTCTCAGATTGCACAGC<br>TGAAGCATTAAAGTGTCTTTTGTTACTATCCCAAATGCCAGAGGAAATTGTGGGAGAAAAAGCTGATAATGAG<br>CGATTGTATGAAGCTGTTAATGTTCTTCTTTACTTACAGAGTCCTATTACTGGAGGTTTTGCTATTTGGGAGC<br>CACCTGTCCCGCAACCATATTTACAAATGTTGAATCCTTCAGAACTTTTCGCAGATATTGTTGTTGAGAAAGA<br>GCATGTCGAGTGTACAGCATCAATTATTCAAGCACTTTTAGCCTTCAAACGACTGCACCCAGGGCACAGGGAG<br>AAAGAAATTGAAATTTCCGTGGCAAAAGCAGTTACATTTTTGGAGGGAAAACAACAGCATGATGGTTCATGGT<br>ATGGTTATTGGGGAATATGCTTTCTATATGGCACATTCTTTGCTTTAGGAGGCTTAGTTTCTGCTGGAAAAAC<br>ATATAATAACAGTGAAGAAATTCGTAAAGCAGTTAATTTTTTCATTTTGACTCAAAATGAAGAAGGCGGTTGG<br>GGAGAAAGTATCAAATCTTGCCCTACTGAAGTATACACACCCTTGGATGGAAACCGAACAAACCTAGTTCAAA<br>CATCATGGGCTATGCTTGCTCTTATGTTAGGTGGACAGGCTGAAAGAGATCCAACACCTTTGCATAAAGCAGC<br>AAAGATATTAATTAATGCACAGATGGATAATGGAGATTTTCCTCAACAAGAGATTACTGGAGTTTACATGAAG<br>AATTGCATGCTACATTATCCAGAATACAGAAACATCTTCCCACTTTGGGCACTTGGCGAATATCGCAAACGAG<br>TTTGGGTCAATTAA |
| >CoOSC<br>4_CDS | ATGTGGGAGTTAAAGATAGCCGAAGGCGATGGCCCTTATTTGTATAGTACCAACAACTTTGTTGGTAGACAAT<br>TCTGGGAATTTAATCCAGATGCTGGAACTTTTGAGGAGAAACAAGAGATTGAAACGGCTCGTGAGAATTATAA<br>AAATAATCGAAGAAATGGAGGATTTCATGCTTGTGGCGACCTTCTCATGCGGAGGCAGTTGATAAAGGAAAAT<br>GGAATAGATCTTACAAGCATAGCTCCTGTGAGAGTAAAGGAGCACGAACACGTCAACTTTGAAGGTGTAACAA<br>CTGCAGTTAGGAAAGCGGTTAGATTACAACGATCCATCCAAGCAAAAGACGGTCATTGGCCAGCTGAAAATGC<br>TGGCCCTATGTTTTTCACTCCTCCACTTGTAATTGCTTTATACATTAGCGGTACGATTAATATAATCTTGACC<br>GAAGAGCACAGGAAAGAGATGATAAGGTACTTCTACAATCATCAGAACGAAGATGGGGGATGGGGGTTTTATA<br>TCGAGGGACATAGCACGATGATTGGATCCGCATTGAGTTATGTGGCCTTACGAATACTAGGAGAGAGAGAAGA<br>TGGCGCCATTTCAAGAGGCCGCAAGTGGATACTTGATCATGGTGGTGCAACCTCTATTCCCTCTTGGGGAAAG<br>GTTTATCTTTCGGTCCTTGGAGTGTATGAATGGGAAGGCTGCAACCCATTGCCACCAGAATTTTGGCTATTCC<br>CATCCGCGTTCCCTTTTCATCCCGCCGAGATGTGGTGCTACTGTCGGACAACCTACATGCCCATGTCATATTT<br>ATACGGAAAAAGAATCCAAGGACCAATCACAAATCTTGTTTTATCATTACGAAAAGAAATCCACCCCACCCCT<br>TTTGAGGACATTAGTTGGAATAAACAAAGGAATAACTGTTGCAAGGAGGACTTCTACTACCCACATTCATTTC<br>TTCAAGATGCATTGTGGCATAGCCTACACCACCTTACTGAGCCGGTTCTCAAACATTGGCCATTTTCCAAACT<br>ACGAAACAGAGCAATTGACAGAGTCGTTGAACTGATGCGTTATGAATCACAAGAGACAAGATACATGACCATT<br>GGATGCGTTGAAAAAAGTTTACAAATGATGTGTTGGTGGGCAGAGAATCCAAATGGGGATGAGTTCAAATATC<br>ACCTTGCAAGAGTACCCGATTATTTGTGGATTGCAGAAGATGGTATGACGATGCATAGTTTTGGTAGTCAAGT<br>GTGGGATTGTTCTCTTGCAATTCAAGCAATTCTTGCAAGTAATATGGTAGAGGAATATGGTGATTGTCTCAAA<br>AAGGCACACTCGTACTTAAGAGAATCACAGGTAAAAGAAAATCCTTCAGGCGATTTTACTCGGATGTGTCGAC |

| | |
|---|---|
| | AGTTCACTAAAGGATCATGGACTTTCTCGGATCAAGATCATGGATGGACTGTCTCAGATTGTACAGCTGAAGC<br>ACTAAAGTGTCTATTGTTATTATCAAACATGCCGAAAGAAATAGCTGGAAACAAAGACAACAGCTCCCGACTC<br>TATGATGCAGTGAATGTGCTTCTTTACATGCAGAGTCCTATAAGTGGAGGATTTGCTGTTTGGGAGCCACCAA<br>TTCCGAAGCCATTTCTACAATTACTTAATCCTTCAGAGATTTTTGCAGACATTGTGGTTGAGAAAGAGCATGT<br>GGAGACCACATCTTCCATTATTGGAGTTCTAATGGAGTTCAATAGCCTCCACCCAAGACACCGAAAGGAAGAA<br>ATACAACTTTCGATTACAAAAGGAATACGCTATCTTGAGGAAACACAATGGCATGATGGTTCATGGTATGGCT<br>ATTGGGGAGTATGTTTCATATATGGAACATTTTTTGCTCTAAGGGGTTTAACTTGTGTCGGAAAAACATATGA<br>AAACAACGAAGCAGTATGTAAAGGTGTTGAGTTCTTACTTTCAATACAGAACGAAGAAGGGGGTTGGGGGGAA<br>AGCCTCTTATCTTGCCCAACCGAGGTATATACACCGCTGGATGGAAACCGAACAAATTTGGTGCAAACCTCAT<br>GGGCTATGCTTGGTCTTATGTTTGCTGGACAAGTGCAAAGAGACCCGAAGCCGTTGCATAAAGCAGCAAAATT<br>ATTGATCAACGCCCAGATGGATAATGGAGATTTTCCTCAGCAGGAAATTACAGGAGTCTACATGAAGAACTGC<br>TTACTCCTTTACGCACAATACAGGAACATATTTCCACTTTGGGCACTTGGGGAGTACCGTAAACGTGTATGGT |
| >CoOSC5_CDS | ATGTGGAAGCTGAAGATAGCAGAAGGAAACGATCCTTACTTGTTTAGCACTAACAACTTTGTTGGGCGTCAGA<br>CTTGGGAGTTTGATCCCGATGCTGGTACTCCCCAAGATGTTGAAAATGCACGCCAATATTTCCTCACCAGGCA<br>AAAGGAAGGTTTTCAAGCATGTAGTGATTTGCTCATGCAGATTCAGTTGACCAAGGAGAACGGGGTTGACTTA<br>TTGACCATACCACCAGCAAGGTTGAGAGATGATGAAGAAGTGAATTATGAAGCGGTGACGACCGCCGTGAAAA<br>AAGCAATCCGATTTCAACGGGCCATACAAGCAAAAGATGGTCATTGGCCAGCTGGACATGTGGGCCCTTTGTT<br>CTTCACTCCACCATTTATCATTGTGTTATACATCAGCGGGACCATCAACACACAATTAACAAAAGAACACAAA<br>AAGGAGATGAAACGGTATATCTACAACCACCAAAATGAAGATGGAGGGTGGGGATTTCATATTGAGGGACATA<br>GCACCATGATGGGATCTGCATTGAACTACATCGCCATACGGTTGTTAGGAGAGGGACCCGATGATGGAAATGG<br>TGCGGTTGACCGAGCTAGAAAGTGGATACTTGACCATGGCGGTGCAACCGCTATTCCATTTTGGGGCAAAATT<br>TATCTCTCAGTGCTAGGAGTGTACGAATGGGAGGGGTGCAACCCAATACCACCAGAATTATGGATTTTTCCCG<br>AAACGTTTCCTCTTCATCCATCGAAAATGTGGTGCCATACTCGGACGGCTTATTTGCCAATGTCGTATTTATA<br>CGGTAGAAAATACCATGGTCCAATCACCGATCTGATTCTCGACTTGAGACAAGAAATTTATCCTATTCCTTAT<br>GACAAGATAATCTGGAATAAACATCGCCACAATTGTTGCAAGGAAGATCTCTTTTTTCATTCAACAATTCAAG<br>ATATGTTGTGGGATGGCCTTCACTATTTTTGTGAACCATTTTTCAAATATTGGCCTTTCACCAAACTTAGAGA<br>AAGAGCACTCAACAGAGTAGTTGAACTGACACGTTATTGTGCACACGAGTCTAGATACATTACAACGGCATCT<br>ATTGAAAAGAGTTTGCAAATGATGTGTTGGTGGGCAGAGAAGCCAAATGGGGATGAATTTAAGCATCATCTTG<br>CCAGAATACCAGATTTCTTATGGTTAGCAGAAGATGGAATGACGATACAAACATTTGGTAGTCAATTATGGGA<br>TTGCACATTTGCAACCCAAGCAATAATCTCAACTAATATGCCCCAAGAATACGGGGATTCACTTAAAAAAGCA<br>AATTTTTACATCAAAGAATCTCAAATCAAGAATAATCCGAGTGGAGATTTTAGTTCAATGTGTAGACAAATTA<br>GCAAAGGTGCGTGGACATTTAGCGACCAAGATCAGGGTTGGCCCGTCTCAGATTGCACAGCTGAAGCGTTAAA<br>ATGTCTTCTTTTACTATCTCAAATGCCAGAGGAAATTACAGGAAAAAAGATTGATAAAGAACAATTATATGAA<br>GCTATTAATTTCTTATTTCATGTGCAGTCTCCTACCAGTGGAGGTTTTGCTTGTTGGGAGCGACCGATCCCAC<br>AACCATATTTAGAGAAGCTGAGTCTTTCAGAAATGTTTGCTGACGTTGTTCTTGAGAAAGAGCATCTCGAGGT<br>TACAGCTTCAATAATTCAAGCTCTAATAGCCTTTAAATGTGCACACCCAACACATAGGGCAAAAGAAATAGAT<br>ATTTCTGTCGCAAATGGAGTGCATTATCTTGAAAATAAACAACTTCCTAACGGTTCATGGTACGGATTTTGGG<br>GAATATGTTTTACATACGGCACGTGCTTTGCATTAGGAGGCTTGGAAGCTGTTGGAAAAACATATAACAACAG<br>TGAATCGGTGCGTAAAGGAGTACAATTTCTCCTTGCAGCACAAAATCAAGAGGGCGGGTGGGGAGAGAGCTAC<br>AAATCTTGCAATACCGAAGTATACACACCGTTGATCGATAATCGATCAACGGTTGTTCAAACCGCGTGGGCCA<br>TGCTTGGGCTTATGTCGGGTGGACAAGCTGAGAGAGATCCGACACCATTACATAAAGCAGCAAAACTTTTGAT<br>TAATGCACAAATGGATAATGGCGATTTCCTGCAACAGGAGTTTACGGGTGCTTCTATGAGGAACGCTACGTTG<br>CATTATCCGTTATATAGGAATTCTTTCACGCTACGGGCCCTTGCGGAATATCGAAAACGTCTTTGGGGGATAA<br>AGTGA |
| >CoOSC6_CDS | ATGTGGAAGTTGAAGATAGCAGAAGGGAATGATCCTTACTTATTTAGTACCAACAATTTCATCGGTAGGCAAT<br>TTTGGGAGTTTGATCCGGATGCCGGTACTCCCGAAGAAAAACAACAAGTCGAAAACGCTCGTCAACATTTTCT<br>AAATAGGCAAAAGGAAGGTTATCAAACATCTAGTGATTTGCTCATGCGGATGCAGTTGACCAAGGAGAATGGG<br>GTTGACTTGTTGAGCATACCACCGGCAAGGTTGAGAGATGACGAAGAAGTGGATTATGAAGCGGTGACGACTA<br>CCGTGAGAAAAGCAGTTCGATATCAACGTGCGATACAAGCAAAAGATGGTCATTGGCCTGCTGAAAATGCGGG<br>CCCTTTGTTCTTCGCTCCACCACTTATAATTGTGTTATACATCAGCGGGACCATCAATACACAATTAACAGAA<br>GAACACAAAAAGGAGCTGAAACGGTATATCTACAACCACCAGAAGGAAGATGGAGGGTGGGGATTTCATATTG<br>AAGGACATAGCACCATGATGAGTTCTGCATTGAACTACATCGCCCTACGGTTGTTAGGAGAAGGACCCGATGA<br>TGGAAATGGTGCAATTGACCGAGCCAGAAAGTGGATAGTTCACCATGGTGGTGCAACCGGAATTCCATCTTGG<br>GGGAAAGCTTATCTATCGGTTCTAGGAGTGTACGAATGGGAGGGATGCAACCCAATGCCACCTGAATTCTGGC<br>TTTTTCCTGAATTATTTCCTTTTCATCCCGCGAAAATGTGGTGCTATTGTCGGTTGACCTATACGCAAATGTC<br>GTATTTATACGGTAGAAAATACCATGGTCCGATCACTGATCTAGTTGTTCATCTAAGACAAGAGATTCATTCT<br>ATTCCTTATGACAAGATAAATTGGAATAAACAACGTCATAATTGCTGCAAGGAAGATCTGATATACCCTCATA<br>CAACAATTCAAGATCTGCTGTGGGATGGTCTACAGTATTTCTGTGAACCACTTTTCAAATATTGGCCGTTCAC<br>CAAACTGAGAGAAAGAGCGCTCAAAAGAGCACTTGAATTGATACGTTATAGCGCACGAGAGAGTAGATACATT<br>ACCATGGCATGTGTTGAAAAGTGTTTGCAATTGATGACTTGGTGGGCAGAGGACCCGAATGGTGACGAGTTTA<br>AGCGCCACCTGGCTCGAGTCCCTGATTACTTGTGGCTTGGAGAAGATGGAATGAAGATGCAAACATATGCTAG |

| | |
|---|---|
| | CCAAGTTTGGGATTGCACAATTGCAACTCAAGCAATAATTTCAACAAATATGCCCCAAGAATACGGGGATTCG<br>CTAAAAAGAGCCAATTTTTTCATCAAAGAATCTCAAATCAAGAACGATCCTCGTGGAGATCTAAGTAAAATAT<br>GTCGACAGTTTAGTAAAGGCGCGTGGCCGTTCACGGATCAAGATCATGGGTGGCCTGTCTCAGACTGCACAGC<br>TGAAGCCGTGAAATGTCTTCTCTTGCTATCTCAAATGCCAGAGGAAATTTCAGGAGAAAAAGCTGATGTGGAG<br>CGACTGTATGACGCTATCACGTTCCTCCTTCATGTACAGTCTCCTATCAGTGGAGGTTTTGCTATTTGGGAAC<br>GACCGATCCCGCAACCGTATTTAGAGAAGTTGAATCCTTCGGAAGTGTTTGCCGACATTGTTCTTGAGAAAGA<br>ACATATGGAGAACACAGTGTCGATAATCCAAGCTTTTGTAGCCTTCAAACGTTTGCATCCAACACATAGGGCA<br>AAAGAATTAGAAAATTCATTGGCAAAAGCCGTGCGTTTTGTTGAGAAAAGACAATTGCCTAACGGTTCATGGT<br>ATGGGTATTGGGGAATATGTTATATATATGGCACGTTATTCGCATTAGGAGGCTTAGAAGCTGTTGGAAAAAC<br>ATACGACAACAGTGAATCGGTTCGCAAAGGAGTAAAGTTTCTCCTCTCGACACAAAATGAAGAAGGCGGGTGG<br>GGGGAGAGCTACAAATCGTGCCTAACCGAAGTGTTCACACCGTTGGTTGAGAACCGAACAAACTTGGTCCAAA<br>CCGCGTGGGCCATGCTTGGGCTTATGTCGGGTGGGCAAGCTGAGAGAGATCCAACACCTTTACATAAAGCAGC<br>AAAACTCTTGATTAATGCACAAATGGATAACGGCGATTTCCCGCAACAGGAGTTAACTGGAGCTTCTTTGAGG<br>AACTGCATGTTACTTTATCCATTGTACAGGAATTCTTTCACGCTATTGGCCCTTGCTGAGTACCGAAAACGTC<br>TTTGGGCTATCAACTAA |
| >CoOSC7_CDS(gDNA) | TCTCGCCTAAGAAAGGAGAACCCGATAAACGAAATACCAGAAGCAATAAAGCTTAATGAAACTGAAGAATTAA<br>CAAATGAAGCAGTCACAACTACACTGAGAAGAGCCATCAGCTTCTACTCAACCATTCAAGCTCATGATGGTCA<br>CTGGCCTGCTGAGTCTGCTGGCCCTTTATTCTTCCTTCCTCCACTGGTAATAGCACTGTATGTGACAGGAACC<br>ATGAATGCTATTCTAACACTTGCACATCAAGTAGAAATCAAACGCTACTTGTATAATCATCAGAATGAAGATG<br>GAGGTTGGGGATTACATATAGAGGGTGACAGCACGATGTTTGGGTCAGTACTTAGTTACATCACCTTGAGATT<br>GCTTGGAGAAGAAGCCGAGGATATGGCGGTGGTCAAGGGCCGTAAATGGATCCTTGACCACGGTGGTGCAATT<br>GGAACTCCTTCATGGGGAAAATTTTGGCTCACGGTTCTTGGAGTATATGAATGGGAAGGTTGTAACCCTATGC<br>CACCAGAATTTTGGCTCCTTCCTAAAATTTTCCCAATTCATCCAGGAAAAATGCTGTGTTATTGTCGCTTAGT<br>TTACATGCCGATGTCGTACCTATATGGAAAAAGATTCGCGGGAAAAATAACTGGATTGGTTCAAGAACTAAGG<br>CAAGAGCTTTACACGGCTCCTTATCATGAGATTAATTGGAATAAAGGGCGAAATACTTGTGCAAAGGAAGATC<br>TCTATTACCCACATCCTCTTGTTCAAGATATGTTATGGGGCGTGCTTCATAACATTGCTGAACCTATTTTAAC<br>ACGTTGGCCGTTTTCCAAACTACGAGAAAAGGCTCTAAAAGTTGCAATGGAGCATGTTCACTATGAAGATCAG<br>AGCAGTAGATATCTTTGCATTGGGTGTGTAGAAAAGGTGTTGTGCTTGATTGCGACCTGGGTGGAAGATCCAA<br>ACGGGGACGCATTTAAGCGCCATCTAGCACGAATTCCTGACTACTTTTGGGTGGCTGAAGATGGGATGAAAAT<br>GCAGAGTTTTGGGTGTCAAATGTGGGATGCATCCTTTGCTATTCAAGCGATTATGTCAAGTAATCTAGCAGAA<br>GAATACGGACCCACTCTTAAAAAAGCACACGAATTTGTAAAGTCGTCGCAGGTTCGCGATAATCCTCCTGGAG<br>ATTTCGGTAAAATGTACAGACACATGTCTAAAGGTGCGTGGACGTTTTCAATGCAAGACCATGGTTGGCAAGT<br>GTCAGATTGCACCGCAGAAGGCTTGAAGGTTGCACTTTTATATTCCCAAATGAGCCCAGAACTTGTTGGTGAA<br>AAACTTGAAAATGAGCATCTCTACGATGCTGTCAATATTATTCTTTCCTTACAAAGTAAAAACGGCGGTTTTC<br>CTGCTTGGGAACCACAAAGGGCATATTCTTGGCTGGAGAAGTTCAATCCGACGGAATTTTTTGAAGATGTGAT<br>GATTGAACGAGAGTATGTGGAATGCACTTCATCCGCCGTCCAAAGTTTAGAACTCTTCAAGAAATTGCACCCT<br>AGACACAGAACTAAGGAGATCGAACATTGCATTTCAAGAGCGTTGAAGTACATTGAAGATATGCAAAACCCTG<br>ATGGTTCATGGTATGGTTGTTGGGGAATATGTTACACATATGGTACATGGTTTGCAGTAGATGCACTAGTAGC<br>TTGTGGGAAGAACTATCATAACTGTCCCGCCCTTCGAAAAGCTTGCCAATTTCTAATATCAAAACAACTTCCT<br>GATGGAGGATGGGGTGAGAGTTATCTTTCCAGTGCTAATAAGGAATATACAAATTTGGATGGAGATCGGTCGA<br>ATTTAGTGCAAACATCTTGGGCTTTAATCTCACTTATTAAAGCTGGACAGGCTGGAATTAATGTTACACCAAT<br>AGATCGTGGAATACGACTTCTAATAAATTCACAAACCGAAGATGGAGACTTCCCTCAACAGGAAATCACAGGA<br>GTTTTTATGAAGAATTGCACCCTCAATTACTCATCGTTTCGAAACATTTTCCCCATATGGGCTCTTGGCGAGT<br>ATCGTCGTATAGTTCAGTTCGGATGA |
| >CoOSC8_CDS | ATGTGGAGGTTGAAGATAGCAGAAGGTGTCAATGATCCACTCCTGCATAGCACAAACAACTTTGTTGGTAGGC<br>AAACATGGGAGTTTGATCCGAATTATGGAACTCCCGAAGAAATTGATGAAGTTGAAAAGGCTCGGCTTCATTA<br>TTGGAATCATCGACACCAAGTTGAGCCATGCAGTGACGTGTTATGGCGTATGCAGTTTTTAAGAGAGAAGAAG<br>TTCAAACAAACCATACCCCAGGTGAAGATTGAGGATGGTGAAGAGATATCTTATGAAAAAGTAACTGCCACAT<br>TGAGGAGGTCTGTTCACTTATTTGCAGCCTTGCAAGCTGAAGATGGTCACTGGCCAGCCGAAAATGCTGGTCC<br>CATGTATTTCATGCAGCCACTGGTGATATGCCTATACATCACAGGGCATCTGAACGGTGTGTTTCCAGCCGAA<br>CACAGAAAAGAAATTCTTCGATACTTATATTGTCATCAGAATGAAGATGGTGGTTGGGGGTTTTACATGGAAG<br>GGCAGAGCACAATGTTTGGCACAACTCTGAGTTACATTTGTATGCGTCTTCTAGGAGAAGGACCTGATAACGG<br>CGCATGCACCAAAGCACGGAAATGGATTCTAGATCATGGAAGTGCGACAGTTATTCCCTCTTGGGGGAAAACG<br>TGGCTTTCGGTCCTTGGTGTTCGTGAGTGGGTAGGAAGCAACCCAATGCCCCCAGAATTCTGGATCCTTCCTT<br>CTTTTCTTCCCATGCATCCAGGAAAAATGTGGTGTTACTGTAGGCTGGTATACATGCCGATGTCATACCTATA<br>CGGGAAACGATTTGTGGGTCCCATCACTCCTCTGGTTTTACAACTCAGGGAGGAGCTTTATTCACAACCATAC<br>GATGAAATTAACTGGAAGAGTACAAGACACGTGTGTGCCAAGGAGGACCTATACTACCCGAGCCCTTTTTTAC<br>AAGATTTATTATGGGACAGTCTCTACATTTTTACCGAGCCTCTCCTAACTCGTTGGCCATTTAACTTGTTGCG<br>CAAGAAAGCACTCAAAACAACCATGAAGCACATCCATTATGAAGACGAGAATAGTCGATATATCACAATTGGT<br>TCTGTTATTAAGTCACTGTGTATGCTTGCGTGCTGGGATGAGGATCCAAATGGGATGTGCTTTAAAAAACACC<br>TTGCCAGAATCCCAGATTATATCTGGATTGCTGAGGATGGAATGAAAATGCAGAGTTTTGGCAGTCAATCATG |

| | |
|---|---|
| | GGATGCAAGTCTTTCAATTCAAGCTTTGTTGGCAACTGATCTTTCCAATGAAATTGGCCCTATGTTAAAGAAA<br>GGACATGACTTCATTAAAGCATCACAGGTTAAGGATAATCCTTCTGGATTATTTAAAAGCATGTATCGCCATA<br>CTTCTAAAGGTTCATGGACCTTCTCAGATCAAGATCATGGATGGCAACTTTCAGACTGTACTGCTGAAGGATT<br>GAGGTGTTGCCTCCTGTTTTCAACGATGCCCCCAGAAATTGTTGGCAAAAAGATGCAGCCTGAGCAATTTTTT<br>GATGCTGTCAACATAATACTTTCCTTACAGAGTAAAAATGGCGGCCTATCAGGATGGGAACCAGCAAGATCAT<br>CAAAATGGCTGGAGATTCTCAATCCTACCGAATTTTTTGAGGACATTGTGATCGAGCATGAGTACGTGGAATG<br>CACATCATCAGGAATACAGACCCTTGTCTTGTTTAACAAGTTATACCCCGAATATAAGACGAAAGAAATACAA<br>CGTTTCCTTACAAATGCAGGTGGATATTTGGAGAAAACTCAAACGCCAGATGGTTCTTGGTACGGAGAGTGGG<br>GTATGTGTTTTACATATGCTACCTGGTTTGCTCTTGGAGGTCTGGAAGCAATTGGAAAGACGTATGAAAACTG<br>TGAAGCAATCCGTAAAGCTGTTAATTTTCTGTTGAAAATACAACGGGAAGATGGTGGATGGGGAGAAAGCCAC<br>CGATCTTCCCCTGAAAAGAAATATATACCTTTAGAAGGCAATCGTCAGTCAAACTTGGTAAATACTGCATGGG<br>CTATGATGGGATTGATTCATTCTGGACAGGCAGAAAGAGACCCAACACCTTTGCACAGAGGAGCTAAGTTGTT<br>GATCAATTCTCAGATGGAAAATGGTGATTTTCCGCAACAGGAAACAACTGGAGTTTTCAAGAAGAATTGCTTG<br>TTGCACTTTTCACTATTCAGGGATATATTCCCAATGTGGGCCCTAGCTAGCTACCGTAAAAGTGTGCTACCTC<br>AGCCCACAAGCATCTAG |
| >CoOSC<br>9_CDS | ATGTGGAGATTAAGACTAGGAGAAGGGGCTGACGATCCATACTTGTTTAGCACAAACAACTTTGTTGGAAGAC<br>AAACATGGGAATTTGATCCCAATTATGGAACTCTAGAAGAACGTGGTGAGGTGGAACAAGCCCGTACCGATTT<br>CTGGAATAACCGACACAAAGTTAAACCTTGTAACGATGCCATTTGGCGCATGCAGTTTCTAAGAGAGAAGAAA<br>TTTAAACAAACAATACCTCAAGTGAAAATAGAGGATGGTGAAGAAATATCATATGAAAAGGCTACAACAACAT<br>TGAAAAGGTCGGTTAACTACATGGCAGCTTTGCAATCTGATCATGGTCATTGGCCAGCTGAAATTGCGGGCCC<br>CCAATATTTCATGCAACCCCTGGTCTTTTGCTTGTATATTAGCGGGCACCTTAACACCGTTTTCCCAGCAGAA<br>CATCAAAAAGAAATCTTACGGAGTTTATATACTCATCAGAATGAAGATGGTGGTTGGGGACTTCACATAGAAG<br>GTCATAGCACGATGTTTTGTACCACTTTAAGTTACATTTGTATGCGTATGTTAGGAGAAGGCCCCGATGGTGG<br>GCTAAATGGCGCCTGCACAAGAGCCCGAAAATGGATTCTTGATCATGGTAGTGTCACACACATTCCATCATGG<br>GGCAAAACATGGCTTTCAATACTTGGTCTATGCGAATGGGCTGGAACCAACCCAATGCCCCCGGAGTTCTGGA<br>TCCTTCCTTCTTTCCTACCTATGCATCCAGCAAAAATGTGGTGCTATTGTCGACTGGTATACATGCCAATGTC<br>ATACTTATATGGAAAAAGATTTGTGGGTCCAATTACTCCTTTGATTCTGCAACTTAGAGATGAACTTTATTTA<br>CAGCCTTACAATGAGATCAAGTGGGGAAGCATAAGACATTTGTGTGCAAAGGAAGATTTATATTATCCTCATC<br>CTTTACTACAAGATTTAATATGGGATAGTCTTTATGTTCTTACTGAGCCTTTATTAACTCGTTGGCCGTTTAA<br>TAAATTGCGCGAAAAAGCACTCCAAACAACAATGAAACATATCCATTATGAAGACGAGAATAGTCGATATATC<br>ACGATTGGTTGTGTAGAAAAGGCACTTTGTATGCTTGCATGTTGGGTTGAGGATCCAAATGGCGATGCTTACA<br>AAAAGCATCTAGCTCGAGTGCCGGATTATCTCTGGATTGCTGAAGATGGAATGAAAATGCAGAGTTTTGGAAG<br>TCAAGAATGGGATGCTGGTTTTGCCATTCAAGCTCTATTAGCTGCTGATTTTACTGATGAAATTGCATCTACC<br>TTAAAGAAAGGGCATGAATTTATTAAAGCTTCACAGGTTAAGGATAATCCTTCTGGTGATTTTAAAAGCATGC<br>ACCGTCATATTTCAAAGGGATCATGGACTTTTTCGGATCAAGATCATGGATGGCAAGTTTCTGATTGTACTGC<br>TGAAGGACTAAAGTGTTGCCTCCTATTCTCAACCATGCCACCAGAAATTGTTGGTGAGCAGATGCCAGCTGAG<br>CAATTGAACAATGCTGTCAACGTAATACTTTCCTTACAGAGCAAAAATGGTGGACTAGCTGCATGGGAGCCAG<br>CAGGAGCTGCAGAATGGTTGGAGATTCTGAATCCTACAGAGTTTTTTGCTGATATCGTCATTGAGCACGAGTA<br>TGTTGAGTGCACAGGATCAGCAATGCATGCCCTTGTTCTGTTTAAGAAGTTATACCCAGGACATAGAAGGAAA<br>GAGATTGAAAATTTCCTTATAAATGCTACTAAATACCTCGAGAATATACAAATGCCAGATGGCTCATGGTATG<br>GTAACTGGGGAGTGTGTTTTACATATGGTACCTGGTTTGCTCTTGGAGGATTGGCAGCTGTTGGAAAAACATA<br>TGAAAATTGTGCAGCAATCCGTAAAGCGGTTAATTTTCTGTTGGAAACACAGCTGAAAGATGGCGGTTGGGGA<br>GAAAGTTATAAATCATGCCCAGAAAAGAGATATATACCTCTAGAAGGAGGCAGGTCCAATTTGGTGCACACTG<br>CATGGGCTTTGATGGGTCTAATTCATTCTCGCCAGATGGATAGAGATGCAACACCCCTACACAGAGCAGCCAA<br>GTTGTTGATCAACTCACAGTTGGAAACTGGTGATTTTCCTCAACAGGAAATAACTGGAGTGTTTATGAAGAAC<br>TGCATGCTACACTATCCAATGTACAAAAATATATACCCAATGTGGGCCCTAGCTGAGTATAGAAAGCATGTTC<br>TACCACTGCTCAAACCATCGATCAAGTGA |
| >CoOSC<br>10_CDS | ATGTGGAGATTGAAACTAGGAGAAGGGGCCGACGATCCATACTTGTTTAGCACAAACAACTTTGTTGGAAGAC<br>AAACATGGGAATTCGATCCAAATTATGGAACTCCCGAAGAACGTGCTGAGGTGGAACAAGCTCGTACCGATTT<br>CTGGAATAACCGACACAAAGTTAAACCTTGCAATGATGCCATTTGGCGCATGCAGTTTCTAAGAGAGAAGAAA<br>TTTAAACAAACAATACCTCAAGTGAAAATAGAGGATGGTGAAGAAATATCATATGAAAAAGCTACAACGACAT<br>TGAAAAGGTCGGTTAATTACATGGCAGCTTTGCAGTCTGATCATGGTCATTGGCCAGCTGAAATTGCGGGTCC<br>CCAATATTTCATGCAACCATTGGTCTTTTGCTTGTATATTAGCGGGCATCTTAACACCGTTTTCCCAGCAGAA<br>CACCAAAAGGAAATCTTACGGAGTTTATATACTCATCAGAATGAAGATGGTGGTTGGGGACTTCACATAGAAG<br>GACATAGCACGATGTTTTGTACGACTTTAAGTTACATTTGCATGCGTATGTTAGGAGAAGGGCCTGATGGTGG<br>GCTAAATGGCGCATGCACAAAAGCCAGAAAATGGATTCTTGATCATGGTAGTGTCACACACATTCCATCATGG<br>GGCAAAACATGGCTTTCTATACTCGGTCTATGTGAATGGGCTGGAACCAACCCAATGCCTCCCGAGTTCTGGA<br>TCCTTCCGTCTTTCCTACCTATGTATCCAGCAAAAATGTGGTGCTATTGTCGACTTGTATACATGCCAATGTC<br>GTACCTATATGGAAAAAGATTTGTGGGCCCCATTACTCCATTGATTCTTCAACTTAGAGATGAACTTTATTTA<br>CAGCCTTACAATGAAATCAAGTGGGGAAGCATAAGACATTTGTGTGCAAAGGAAGATTTATACTATCCTCATC<br>CTTTACTACAAGATTTAATATGGGATAGTCTTTATGTTCTTACTGAGCCTTTACTAACTCGTTGGCCGTTTAA |

| | |
|---|---|
| | CAAATTGCGCGAAAAAGCACTCCAAACAACAATGAAACATATCCATTATGAAGACGAGAATAGTCGATATATC<br>ACTATCGGTTGTGTAGAAAAGGCACTTTGTATGCTTGCATGTTGGGTTGAGGATCCGAATGGTGATGCTTACA<br>AAAAGCATTTAGCTCGAGTGCCAGATTATCTCTGGATTGCTGAAGATGGAATGAAAATGCAGAGTTTTGGAAG<br>TCAAGAATGGGATGCTGGTTTTGCCATTCAAGCTCTATTGGCTGCTGATTTTACTGATGAAATTGCATCTACC<br>TTAAAGAAAGGACATGAATTCATTAAAGCTTCACAGGTTAAGGATAATCCTTCTGGTGATTTTAAAAGCATGC<br>ACCGTCATATTTCAAAGGGATCATGGACCTTTTCAGATCAAGATCATGGATGGCAAGTTTCTGATTGTACTGC<br>TGAAGGACTAAAGTGTTGCCTCCTATTCTCAACCATGCCACCAGAAATTGTTGGTGAGCAGATGCCAGCTGAG<br>CAATTGAACAATGCTGTAAATGTAATACTTTCCTTACAGAGCAAAAATGGTGGGCTAGCAGCATGGGAGCCAG<br>CAGGAGCTGCAGAATGGTTGGAGATTCTGAATCCTACAGAGTTTTTTGCTGATATCGTCATTGAGCACGAGTA<br>CGTTGAGTGCACAGGATCAGCAATGCATGCCCTTGTTTTGTTTAAAAAGTTATACCCGGGTCACAGAAGGAAA<br>GAGATTGAAAATTTCCTTATAAATGCAACTAAATACCTCGAGAGTATACAGATGCCAGATGGCTCATGGTATG<br>GTAACTGGGGAGTGTGTTTTACATATGGTACCTGGTTTGCTCTTGGAGGATTGGCAGCTGTTGGAAAAACATA<br>TGAAAACTGTGCAGCAATCCGAAAAGCGGTTAATTTTCTGTTGGGAACACAGCTGAAAGATGGCGGTTGGGGA<br>GAAAGTTATAAATCCTGCCCAGAAAAGAGATATATACCTTTGGAAGGAGGCAGGTCCAACTTGGTGCACACTG<br>CATGGGCTCTGATGGGTCTAATTCATTCTCGCCAGATGGAGAGAGATGCAACACCTCTACACAGAGCAGCCAA<br>GTTGTTGATAAACTCACAGTTGGAAACTGGTGATTTTCCACAACAGGAAATAACTGGAGTGTTTATGAAGAAT<br>TGCATGCTTCACTATCCAATGTACAAAAATATATACCCAATGTGGGCCCTAGCTGAGTATAGAAAGCATGTTC<br>TACCACTGCTCAAACCATCCATCAAGTGA |
| >CoOSC11_CDS | ATGTGGAGATTAAAAATAGCAAAAGGAGGCGACAATCCGTACTTGTACAGTACAAACAACTTTGTTGGAAGAC<br>AAATATGGGAATTTGACCCAAATTATGGGACTCCAGAAGAACATGCTGACGTGGAAAATGCTCGTGTTGGTTT<br>TTGGAATAATCGACATGATATTAAAACTAGCGGTGATGTTCTTTGGCGGATGCAGTTTCTAAGAGAGAATGAA<br>TTTAAACAAACAATACCACAAGTGAAAATAGAAGACGGTGAAAAAATATCAAATGAAAAAGCTACAACAACAT<br>TGAAAAGATCGGTTAACTTCTTTTCAGCTTTGCAAGCTGATGATGGTCATTGGCCCGCTGAAAATGCGGGCCC<br>GTTGTATTTCATGCAACCCTTGGTCATGTGCTTGTATATAACCGGCCATCTCAACACTGTTTTCCCAACAGAA<br>TACCGAAAAGAAATCCTACGGTATATATACTGTCATCAGAATGAAGATGGTGGTTGGGGACTTCACATAGAAG<br>GTCATAGTACGATGTTTTGTACAACTTTAAGTTACATTTGTATGCGTATGTTAGGAGAAGGGCCTGATGGTGG<br>GCTAAATGGCGCATGCACAAAAGCCAGAAAATGGATTCTTGATCATGGTAGTGTCACACACATTCCATCATGG<br>GGCAAAACATGGCTTTCGATACTTGGACTATGTGAATGGGCTGGAACCAACCCAATGCCTCCCGAGTTCTGGA<br>TCCTTCCTTCTTTCCTACCTATGTATCCTGCGAAAATGTGGTGCTATTGTCGACTTGTATACATGCCAATGTC<br>GTACCTATATGGAAAAAGATTTGTGGGTCCCATTACTCCGTTGATTCTTCAACTTAGAGATGAACTTTATTTA<br>CAGCCTTACGATGAAATTAACTGGAAAAGTGTACGACACTTATGCGCAAAGGAAGATTTATATTATCCGCATC<br>CGTTACTACAAGATTTAATGTGGGATAGTCTTTATGTTTTTACTGAGCCTTTTTTAAACCACTGGCCATTTAA<br>TAAATTGCGTGAAAAAGCACTCCAAACAACAATGAAACATATCCATTATGAAGATGAGAATAGTCGATATATC<br>ACTATTGGTTCGGTAGAAAAGGCATTGTGTATGCTTGCATGTTGGGTTGAGGATCCAAATGGCGTTTGTTTTA<br>AAAAGCATTTAGCCCGAATCCCAGATTATTTATGGGTTGCTGAAGATGGAATGAAAATGCAGAGTTTTGGAAG<br>CCAACAGTGGGATGCTGGTTTTGCTATTCAAGCTTTATTGGCTACGGATCTTACAGACAAAATTGGGTCTACA<br>TTGATGAAAGGACATGAATTTATTAAAGTTTCACAGGTTAAGGATAATCCTTCTGGTGATTTTAAAAGCATGC<br>ACCGTCATATTTCAAAGGGATCATGGACCTTTTCAGATCAAGATCATGGATGGCAAGTTTCAGACTGTACTGC<br>TGAAGGACTAAAGTGTTGCCTCCTATTCTCAACCATGCCACCAGAAATTGTCGGTGAGAAAATGAAACCCGAG<br>CAACTAAACGATGCTGTCAACATAATACTTTCCTTACAGAGCAAAAACGGTGGGCTGGCAGCATGGGAACCAG<br>CAGGATCATCAGAATGGCTAGAGGTTCTTAATCCTACAGAGTTCTTTGCAGATATTGTAATTGAACATGAGTA<br>TGTGGAGTGTACATCATCAGCAATGCAAGCCCTAGTTTTATTTAAAGAGTTATTCCCAGGGCATAGAAGGAAA<br>GAGATTGAAAATTTCCTAACTGCTGCAAGTGGATATCTGGAGAACATACAAATGCCAGATGGCTCATGGTATG<br>GAAATTGGGGAGTGTGTTTTACATACGGTACCTGGTTTGCTCTTGGAGGATTGACAGCTGTTGGGAAAACGTA<br>TGAAAATTGTGCAGCAATTCGAAAAGCTGTTAAATTTCTGTTGGAAACACAGCTGGAAGATGGTGGTTGGGGA<br>GAAAGTTATAAATCCTGCCCAGAAAAGAGATATATACCTCTGGAAGGAGGCAAGTCCAACTTGGTCCATACTT<br>CATGGGCTTTGATGGGCCTGATTCATTCTCGCCAGATGGAAAGAGATGCAACACCCCTACACAGAGCAGCCAA<br>ATTGTTAATCAACTCACAATTGGAAAATGGTGATTTTCCACAACAGGAAATAGCTGGAGTGTTTATGAAGAAC<br>TGTATGCTGCATTATCCATTATACAGAAATATATACCCAATGTGGGCCCTAGCTGATTACAGCAAGCAAGTTC<br>TACCATCGAACAGCTGA |
| >CoOSC12_CDS | ATGTGGAAATTGAAGATCGCAGAGGGAGGGAGTCCATGGCTCCGTTCGAATAACGATCACGTCGGCCGGCAAA<br>TTTGGGAGTTTGATCCGACGCTAGGTTCCCTTGAGGAACTTGCCGATATCGAGAAAGTCCGGCAGGCATTCCA<br>TGACAATCGGTTCGAGAAGAAACACAGTTCAGATCTGCTTATGCGTAGTCAGTTTGCGAAGGTGAAGCCACAA<br>TCTGTTTTTCCACCAAAAGTGAATATAAAAGATGCTGAAGATATTACAGAAAATAAAGTAACAGATGTATTAA<br>GAAGAGCAATTAGTTTCCATTCAACTCTTCAAGCAGATGATGGACATTGGCCAGGAGATTATGGTGGTCCCAT<br>GTTTTTATTGCCTGGTTTGGTTATTACTCTAACTATTACTGGAGCATTGAATGCGGTCTTATCCAAAGAACAC<br>AAGCGGGAGATGGTTCGTTACCTTTACAATCATCAGAATATAGATGGTGGGTGGGGCCTACACATTGAGGGTC<br>ATAGCACCATGTTTGGTAGTGCATTAAACTATGTTACTTTGAGATTACTTGGGGAAGGGGCTAATGATGGAGA<br>AGGGGCAATGGAAAAAGGGCGAAAATGGATTTTGGATCATGGTGGGGCCACTTCAATAACATCATGGGGAAAA<br>TTCTGGCTTTCAGTACTTGGTGTATTTGAATGGTCGGGAAATAATCCGTTGCCCCCTGAGATGTGGATACTTC<br>CATATTTCCTTCCGATACATCCAGGTAGGATGTGGTGTCACTGTAGAATGGTGTACCTACCTATGTCATATTT |

| | |
|---|---|
| | ATATGGGAAAAGATTTGTGGGACCCATCACATCAACAGTTCTAGCCTTGAGAAAAGAACTGTTCACTGTTCCT<br>TATCATGATATAGATTGGAATGTTGCAAGAAATCTATGTGCAAAGGAAGATCTTTACTACCCACATCCACTTG<br>TTCAAGACATACTTTGGGCTACTCTTGACAAATTTGTAGAGCCCGTGTTAATGAGATGGCCCGGAAAGAAGTT<br>GAGAGCAAAAGCTCTTCGTACTGCAATGGAACATATTCATTATGAGGATGAAAATACTCGATATATTTGCATC<br>GGGCCAGTAAACAAGGTTTTAAATATGCTCTGCTGTTGGGCAGAAGACCCAAACTCAGAGGCATTTAAGTTAC<br>ACCTACCAAGAATACATGATTATTTGTGGATTGCTGAAGATGGCATGAAGATGCAGGGTTATAATGGTTGTCA<br>ACTATGGGATACTGCATTCGCTGTACAAGCAATTATTTCCGCAAACCTCATTGAAGAATTTGGTTCAACACTG<br>AAAAAGGGACACATGTATATAAAGAATTCACAGGTGTTAGATAACTGCACTGGTGATCTTGATTATTGGTATC<br>GTCATATTTCAAAAGGTGCTTGGCCTTTTTCAACTGCAGATCACGGCTGGCCCATTTCAGATTGTACTGCAGA<br>AGGCCTAAAGGCTGCACTACTGCTCTCAAAATTGCCAACAGAGATCGTTGATGAACCAGTGGATGAAAATCGG<br>CTGTACGATTCCGTCAATGTTATTCTATCTTTACAGAATGGTGATGGTGGTTTTGCAACTTATGAACTAACAA<br>GATCCTACAGATGGTTAGAGTTGGTCAATCCTGCTGAAACTTTTGGTGACATTGTTATCGACTATCCATATGT<br>AGAGTGTACTTCAGCTGCAATACAAGCTCTGGTGGCTTTTAAGAAACTGTACCCTGGACATAGGAGAAAAGAA<br>GTAGAACGTTGTATTGACAAAGCTGCTGCATTCATTGAAAGAATACAAGAATCAGATGGTTCATGGTATGGAT<br>CATGGGCGGTTTGTTTCACTTATGGGACATGGTTTGGAATCAAAGGTTTAGTGGCTGCTGGGAAGACATACAG<br>TAACTGTAGTAGCATTCGCAAGGCTTGTAACTTTTTGTTGTCTAAACAACTTGCTTCTGGTGGTTGGGGAGAG<br>AGTTATCTCTCCTGTCAGGATAAGGTTTATACGAATCTTGAGGGAAATCGGTCTCATGTGGTAAATACAGGAT<br>GGGCTATGCTATCTCTCATTGATGCTGAGCAGGCGAAGAGAGATCCGACACCATTGCACCGTGCAGCAAGAGT<br>GCTGATTAATTCTCAGATGGAAAATGGAGATTTCCCACAACAGGAGATAATGGGGGTGTTTAACAGGAACTGC<br>ATGATAACATATGCTGCATACAGAAACATCTTCCCTATTTGGGCCTTAGGAGAATACCGTTGCAGAGTACTTA<br>CTTAA |
| >CoOSC<br>13_CDS | ATGTGGAAATTGAAGATCGCAGAGGGAGGGAGTCCATGGCTCCGTTCGACTAACGATCACGTCGGCCGTCAAA<br>TTTGGGAGTTTGATCCTACGCTTGGGTCCCTTGAGGAACTCGCCGATATCGAGAAAGTCCGGCAGTCATTTCA<br>TGACAATCGGTTTGAAAAGAAACACAGTTCAGATCTGATTATGCGTAGTCAGTTTGCGAAGGTGAAGTCACAA<br>TCTGTTTTTCCACCAAAAGTGAATATAAAAGATGCTGAAGATATTACAGAGGATAAAGTAACAGATGTATTAA<br>GAAGAGCACTTGGTTTCTATTCAACCCTCCAAGCAGATGATGGACATTGGCCAGGAGATTATGGAGGCCCCAT<br>GTTTTTATTGCCTGGCTTGGTTATTACTCTAACTATTACTGGGGCACTGAATGCAGTCTTATCCAAAGAGCAC<br>AAAAGGGAGATGGTTCGTTACCTTTACAATCATCAGAATATAGATGGCGGGTGGGGCCTACACATTGAGGGTC<br>ATAGCACCATGTTTGGTAGTGCATTAAACTATGTGACTTTGAGATTACTTGGGGAAGGGGCTAATGATGGACA<br>AGGGGCAATGGAAAAGGGACGACAATGGATTTTGGATCATGGTGGGGCCACTTTCATAACATCATGGGGAAAA<br>TTCTGGCTTTCTGTACTTGGTGTGTTTGATTGGTCTGGAAATAATCCATTACCCCCTGAAATGTGGCTCTTTC<br>CATATTTCCTCCCGATACATCCAGGTAGGATGTGGTGTCATTGTAGAATGGTTTACCTACCTATGTCTTACTT<br>ATATGGAAAGAGGTTTGTGGGACCCATCACATCAACAATTCTTGCCTTGAGAAAAGAGCTGTTTACTGTTCCT<br>TATCATGACATTGATTGGAATGAAGCAAGAAATCTTTGTGCAAAGGAAGATCTTTACTACCCACATCCACTTG<br>TGCAAGACATACTTTGGGCCACTCTTGACAAGTTTGTGGAGCCCGTTTTAATGAGATGGCCCGGAAAGAAGTT<br>GAGAGAGAAAGCTCTTTGTACTGCAATGGAACATATTCATTATGAGGATGAAAATACTCGATATATTTGCATT<br>GGCCCAGTAAACAAGGTGCTAAATATGCTCTGCTGCTGGGTAGAAGACCCAAACTCAGAGGCTTTCAAGTTAC<br>ACCTCCCAAGAATACAAGATTACCTATGGCTCGCTGAAGATGGCATGAAAATGCAGGGTTATAATGGTAGTCA<br>ACTATGGGATACTGCATTTGCTGTACAAGCAGTTATTTCTACAAACCTCATTGAAGAATTTGGTCCAACATTG<br>AAAAAAGGACACATGTATATAAAGAATTCACAGGTGCTAGAGAACTGTACTGGTGATCTTGATTATTGGTATC<br>GTCATATTTCAAAAGGGGCTTGGCCTTTTTCAACTGCTGATCACGGCTGGCCCATTTCAGATTGTACTGCAGA<br>AGGCTTAAAGGCTGCACTCCTGCTTTCAAAATTGCCAAAACAAATTGTTGATGAACCAGTGGATGAAAATCGG<br>CTGTATGATTCCGTCAATGTTATTCTATCTTTACAGAATGGTAATGGTAGTTATGCAACATATGAACTCACAA<br>GATCCTACAGCTGGTTAGAGTTGGTCAATCCTGCTGAAACTTTCGGTGACATTGTTATCGACTACCCATATGT<br>AGAGTGTACTTCGGCTGCAATTCAAGCTCTAGTAGCTTTTAAAAAATTATACCCTGGGCATAGGAGAGAGGAG<br>GTACAACGTTGTATTGATAAAGCTGCTGCATTCATTGAAACAATCCAAGAATCAGACGGTTCATGGTATGGAT<br>CATGGGCAGTTTGTTTCACTTATGGTACATGGTTTGGAATCAAAGGTTTAGTGGCTGCTGGAAAGAGTTACAG<br>TAACTGCTCCTCCATTCGCAAGGCTTGTGACTTTTTGTTGTCTAAACAACTTGCTTCTGGTGGCTGGGGAGAG<br>AGTTACCTCTCATGTCAGGACAAGGTTTATACCAATCTTGAGGGAAATCGTTCTCATGTGGTAAATACAGGGT<br>GGGCTATGCTAGCTCTCATTGATGCTGAACAGGCTAAGAGAGATCCAAGACCATTGCACCTTGCTGCAAGAGT<br>ATTAATTAATTCTCAGATGGAAAATGGAGACTTCCCACAACAGGAGATCATGGGGGTGTTTAACAGAAACTGC<br>ATGATAACGTATGCTGCATACAGAAACATTTTCCCAATTTGGGCATTGGGAGAATACCGCTGCCGAGTACTTC<br>AGCAGCCACCTTGTTAA |
| >CoOSC<br>14_CDS | ATGTGGAAGTTGAAGATCGCAGAGGGAGGGAGTCCATGGCTCCGTTCGAATAACAATCATGTCGGCCGGCAAA<br>TTTGGGAGTTTGATCCGACGCTAGGGTCCATTGAGGAACTTGCCGATATCGAGAAAATCCGGCAGACATTCCA<br>TGACAATCGGTTCGAGAAGAAACACAGCTCAGATCTGCTTATGCGTAGTCAGTTTGCGAAGGTGAAGTCTCAA<br>TCTGTTTTTCCGCCAAAAGTGAATATAAAAGATGCTGAAGATATAACAGAAGATAAAGTAACAGATGTATTAA<br>GAAGAGCAGTTAGTTTCCATTCAACTCTTCAAGCAGATGATGGACATTGGCCAGGAGATTATGGCGGTCCCAT<br>GTTTTTATTGCCTGGTTTGGTTATCACTCTAACTATTACTGGAGCACTGAATGCGGTCTTATCCAAAGAACAC<br>AAGCGGGAGATGGTTCGTTACCTTTACAATCATCAGAATATAGATGGTGGGTGGGGCCTACACATTGAGGGTC<br>ATAGCACCATGTTTGGTAGTGCATTAAACTATGTTACTTTGAGATTACTTGGGGAAGGGGCTAATGATGGATC |

| | |
|---|---|
| | AGGGGCAATGGAAAAAGGGCGAAAATGGATTTTGGATCATGGTGGTGCCACTTCAATAACATCATGGGGAAAA<br>TTCTGGCTTTCAGTACTTGGTGTATTTGAATGGTCGGGAAATAATCCGTTGCCACCCGAGATGTGGATCCTTC<br>CATATTTCCTCCCGATACATCCAGGTAGGATGTGGTGTCATTGTAGAATGGTTTACCTACCTATGTCATATTT<br>ATATGGGAAAAGATTTGTGGGACCCATCACAGCAACAGTTCTAGCCTTGAGAAAAGAACTGTTTACTGTTCCT<br>TATCATGACATAGATTGGAATGTTGCAAGAAACCTATGTGCAAAGGAAGATCTTTACTACCCACATCCATTTG<br>TTCAAGACATAATTTGGGCTACTCTTGACAAATTTGTGGAGCCCGTTTTAATGAGGTGGCCTGGAAAGAAGTT<br>GAGAGCAAAAGCTCTTCAAACTGCAATGGAACATATTCATTATGAGGATGAAAATACTCGATATATTTGCATC<br>GGGCCAGTAAATAAAGTGCTAAATATGCTCTGTTGTTGGGCAGAAGACCCAAACTCAGAGGCATTTAAGTTAC<br>ATCTACCAAGAATACAGGATTATTTGTGGATTGCTGAAGATGGCATGAAGATGCAGGGTTATAACGGTAGTCA<br>ACTATGGGATACTGCATTCGCTGTACAAGCAATTATTTCCACAAACCTCATTGAAGAATTTGGTCCAACGCTG<br>AAAAAGGGACACATGTATATAAAGAATTCACAGGTGTTAGATAACTGCACTGGTGATCTTGATTATTGGTACC<br>GTCATATTTCAAAAGGCGCTTGGCCTTTTTCAACTGCAGATCACGGTTGGCCCATTTCAGATTGTACTGCAGA<br>AGGCCTAAAAGCTGCACTCCTGCTCTCAAAATTGCCAAGAGAAATCGTTGATGAACCAGTGGATGAAAATCGG<br>CTGTATGATTCCGTCAATGTTATTCTATCTTTACAGAATGGTGATGGCGGTTTTGCAACTTATGAACTCACAA<br>GATCCTACCATTGGTTAGAGTTGGTGAATCCTGCTGAAACTTTTGGTGACATTGTTATCGACTATCCATATGT<br>AGAGTGTACTTCAGCTGCAATTCAAGCTCTGGTGGCTTTTAAGAAATTGTACCCTGAGCATAGGAGAGACGAA<br>GTAGAACATTGTATTGATAAAGCTGCTGCGTTTCTTGAAAGAATACAAGAATCAGACGGTTCATGGTATGGAT<br>CATGGGCGGTTTGTTTCACTTATGGGACATGGTTTGGGATCAAAGGTTTAGTGGCAGCTGGAAAGACGTACAG<br>TAATTGCTGTAGCGTTCGCAAGGCTTGTGACTTTTTGTTGTCTAAACAACTTGCTTCTGGTGGTTGGGGAGAG<br>AGTTATCTCTCGTGTCAGAATAAGGTTTATACGAATCTTGAGGGAAATCGGTCTCATGTGGTAAATACAGGGT<br>GGGCTATGCTAGCTCTCATTGATGCTGAGCAGGCGAAGAGAGATCCAACACCACTGCACCGTGCAGCAAGAGT<br>GTTGATTAATTCTCAGATGGAAAATGGAGATTTCCCACAACAGGAGATAATGGGGGTGTTCAACAGGAACTGC<br>ATGATAACATATGCTGCATACAGAAACATCTTCCCTATTTGGGCCTTAGGAGAATACCGTTGCCGAGTACTTA<br>CTTAA |
| >CoOSC<br>15_CDS | ATGTGGAAATTGAATATCGCAGAGGGAGGGAGTCCATGGCTCCGTTCGACAAATGATCACGTCGGCCGGCAAA<br>TTTGGGAGTTTGATCCGACGCTTGGGTCCCTTGAGGAACTCGCCGATATCGAGAAGGTCCGTCAGACATTCCA<br>TGACAATCGGTTTGAAAAGAAACACAGCTCAGATCTGATTATGCGTAGTCAGTTTGCAAAGGCGAAGTCACAA<br>TCTGTTTTTCCACCAAAAGTGAATATAAAAGATGCTGAAGATATTACAGAGGATAAAGTAACAGATGTATTAA<br>GAAGAGCAGTTGGTTTCTACTCAACCCTCCAAGCAGATGATGGACATTGGCCAGGAGATTATGGAGGCCCCAT<br>GTTTTTATTACCTGGATTGGTTATTACTCTAACTATTACTGGAGCACTCAATGCAGTCTTATCCAGAGAGCAT<br>AAACGGGAGATGGTTCGTTACCTTTACAATCATCAGAATATAGATGGTGGGTGGGGCCTACACATCGAGGGTC<br>ATAGCACCATGTTTGGTAGTGCATTAAACTATGTGACTTTGAGATTACTTGGGGAAGGGGCTAATGATGGACA<br>AGGGGCAATGGAAAAGGGAAGAAAATGGATTTTGGATCATGGTGGTGCCACTTTCATAACATCATGGGGAAAA<br>TTCTGGCTTTCTGTACTTGGTGTGTTTGATTGGTCTGGAAATAATCCAATGCCCCCTGAGATGTGGCTCTTTC<br>CATATTTCCTCCCGATACATCCAGGTAGGATGTGGTGTCATTGTAGAATGGTTTACCTACCTATGTCTTACTT<br>ATATGGAAAGAGGTTTGTGGGACCCATCACATCAACAATTCTTGCCTTGAGAAAAGAGCTGTTTACTGTTCCT<br>TATCATGACATTGATTGGAATGAAGCAAGAAATCTTTGTGCAAAGGAAGATCTTTACTACCCACATCCACTTG<br>TGCAAGACATACTTTGGGCCACTCTTGACAAGTTTGTGGAGCCCATTTTAATGAGATGGCCTGCAAAGAAGTT<br>GAGAGAGAAAGCTCTTTGTACTGCAATGGAACATATTCATTATGAGGATGAAAATACTCGGTATATTTGCATT<br>GGTCCAGTAAACAAGGTGCTAAATATGCTCTGCTGCTGGGTAGAAGACCCAAACTCAGAGGCTTTCAAGTTAC<br>ACCTCCCAAGAATACAAGATTACCTATGGCTTGCTGAAGATGGCATGAAAATGCAGGGTTATAATGGTAGTCA<br>ACTATGGGATACTGCATTTGCTGTACAAGCAATCATATCTACAAACCTCATTGAAGAATTTGGTCCAACATTG<br>AAAAAAGGACACATGTATATAAAGCATTCACAGGTGCTAGATAACTGTACTGGTGATCTTGATTATTGGTATC<br>GTCATATTTCAAAAGGTGCTTGGCCCTTTTCAACTGCTGATCATGGTTGGCCCATTTCAGATTGCACTGCAGA<br>AGGCCTAAAGGCTGCACTCCTGCTTTCAAAATTTCCAACAGAAGTTGTTGATGAAGCAGTGGATGAAAACCGG<br>CTGTATGATTCCGTCAATGTTATTCTATCTTTACAGAATGGTGATGGTAGTTATGCAACATATGAACTCACAA<br>GATCCTACAGCTGGTTAGAGTTGGTTAATCCTGCTGAAACTTTCGGTGACATTGTTATTGACTACCCATATGT<br>AGAGTGTACATCGGCTGCAATTCAAGCTCTTGTGGCTTTTAAAAAGTTATACCCTGGGCATAGGAGAGAGGAA<br>GTAGAACGTTGTATTGATAAAGCTGCTGCATTCATTGAAACAATCCAAGAATCAGACGGTTCATGGTATGGAT<br>CATGGGCAGTTTGTTTCACTTATGGAACATGGTTTGGAATCAAAGGTTTAGTGGCTGCTGGAAAGAGTTACAG<br>TAACTGCTCTTCCATTCGCAAGGCTTGTAACTTTTTGCTGTCTAAACAACTTGCTTCTGGTGGTTGGGGAGAG<br>AGTTATCTCTCGTGTCAGGACAAGGTTTATACCAATCTTGAGGGAAACCGTTCTCATGTGGTAAATACAGGGT<br>GGGCTATGCTAGCTCTCATTGATGCTCAACAGGCTAACAGAGATCCAAGACCATTGCACCTTGCTGCAAGAGT<br>ATTAATTAATTCTCAGATGGAAAATGGAGACTTCCCACAACAGGAGATCATGGGGGTGTTTAACAGAAACTGC<br>ATGATAACGTATGCTGCATACAGAAACATTTTCCCAATTTGGGCATTGGGAGAATACCGCTGCCGAGTCCTTC<br>AGCCACCTTGTTAA |
| >CoOSC<br>16_CDS | ATGTGGAAATTGAAGATCGCAGAGGGAGGGAGTCCATGGCTCCGTTCGAATAACAATCACGTCGGCCGGGAAA<br>TTTGGGGGTTCGATCCGACGCTAGGGTCCCTTGAGGAACTCGCCGATATCGAGAAAGCCCGGCAGACATTCCA<br>TGACAATCGGTTCGAGAATAAACATAGCTCAGATCTGCTTATGCGTAGTCAGTTTGCAAAGGTGAAGTCGCAA<br>TATGTTTTCCCTCCAAAAGTGACTATAAAAGATGCTGAAGATATAACAGAAGATAAAGTAACAGATGTATTAA<br>GAAGAGCAATTGGTTTCCATTCAACTCTTCAAGCAGATGATGGACATTGGCCAGGAGATTATGGTGGCCCCAT |

| | |
|---|---|
| | GTTTTTATTGCCTGGTTTGGTTATCACTCTAACTATTACTGGGGCACTGAATGCCGTCTTATCCAAAGAACAC<br>AAACGGGAGATGGTTCGTTACCTTTACAATCATCAGAATATAGATGGTGGGTGGGGCCTGCACATTGAGGGTC<br>ATAGCACCATGTATAGTAGTGTAATAAACTATGTTACGTTGAGATTACTTGGGGAAGGGGCTAATGATGGAGA<br>AGGGGCAATGGAAAAAGGACGAAAATGGATTTTGGATAATGGTGGGGCTACTTCAATAACATCATTTGGAAAA<br>TTCTGGCTTTCAGTACTTGGTGTATTTGAATGGTCTGGAAATAATCCTCTGCCGCCTGAGATGTGGATCCTTC<br>CATATTTCCTTCCAATACATCCAGGTAGGATGTGGTGTCATTGTAGAATGGTTTTCCTACCTATGTCATATTT<br>ATACGGGAAAAGATTTGTGGGACCCATCACAACACAAGTTCTAGCCTTGAGAAAAGAACTGTTCACTGTTCCT<br>TATCATGACATAGATTGGAATGTTGCAAGAAACCTATATGCAAAGGAAGATCTTTGCTACCCAAATCCACTTG<br>TTGAATACATACTTTGGCCTACTCTTAACAAATTTGTGGAGCCCATGTTAATGCAATGGCCTGGAAAGAAGTT<br>GAGAGAAAAAGCTCTTCTTACTGCAATGGAACATATTCATTATGATGATGAAAATACTCAATATCTCTGCATT<br>GGGCCAGTAAACAAGGCGCTAAATATGCTCTGCTGTTGGGCAGAAGATCCAAACTCAGAGGCATTTAAGTTAC<br>ACCTACCAAGAATACATGATTATCTGTGGATTGCTGAAGATGGCATGAAGATGAAGGGTTATAACGGTACTCA<br>ACTATGGGATACTGCATTCGCTATACAAGCAATTATTTCCACAAACCTCATTGAAGAATTTGGTCCAACACTA<br>AGAAAAGGACACATGTATATAAAGAATTCACAGGTGTTAGATAACAGCACTGGTAATCTTGATTATTGGTATC<br>GTCACATTTCAAAAGGTGCTTGGACTTTTTCAACTGCAGATCAAGGTTGGCCCATTTCAGATGGTACTGCAGA<br>GGGCCTAAAATCTGCACTCCTGCTCTCAAAATTGCCAACAGAAATCGTTGATGAACCACTGGATGAAAATCGA<br>CTGTATGATTCTGTCAATGTTATTTTATCTTTACAGAATGGTGATGGTGGTTTTGCATCTTATGAACTCACAA<br>GATCCTACAGCTGGTTAGAGTTTGTCAATCCTGCTAAAACTTTTGGTGATATGGTTATTGACTATACATATGT<br>AGAGTGTACTTCAGCATCAATTCAAGCTCTGGTGACTTTTAAGAAACTGTACCCTGGGCATAGGAGAGAAGAA<br>GTGCAACATTGTATTGATAAAGCTGCTGCATTCATTGAAAGAATGCAAGAATCAGACGGTTCATGGTATGGAT<br>CATGGGGGGTGTGTTTCACTTATGGGACATGGTTTGGAGTCAAAGGTTTAGTGGCTGCTGGAAAGACTTACAC<br>TAACTGTAGTAGCATTCGCAAGGCTTGCAACTTTTTATTGTCTAAACAACTTGCTTCTGGTGGTTGGGGAGAG<br>AGTCATCTCTCGTGTCAGAAAAAGGTTTATATGAATCTTGAGGGAAATCGGTCTCATGTGGTGAATACGGGGT<br>GGGCTATGCTAGCTCTCATTGATGCTGAGCAGGCGAAGAGAGATCCAACACCATTGCACCGTGCAGCAAGAGA<br>ATTGACTAATTCTCAGATGGAAAATGGAGATTTCCCACAACAGGAGATAATTGGGGTGTTCAACAGGAACTGC<br>ATGATAACATATGCTGCATACAGAAACATCTTCCCTATTTGGGCCTTAGGAGAATACCGTTGCCGAGTACTTC<br>AAATGCTCTCCTCGATCTAG |
| >CoOSC 17_CDS (gDNA) | ATGTGGAAATTGAAAATTGGTGAAAAGAATGGAAAGCTGGAAATTGGTGACGGAAATGGTGATGAATATTTGT<br>ATAGTACCAATAAATTTGTGGGGAGACAAACATGGGAGTTCGATCCAAGTCAGGAAGAACGTGAGCAAATAGA<br>AAGGATTCGAGAGCAATTCTTGAACAATAAGAAGAAACTCGACATCCATTGTTGTGGTGACTTGCTCATGCGA<br>AACCAGATTCAAGCTTGGGACGGTCATTGGCCAGCTGAAAATGCGGGTCCCCTTTTTTTCACCCCTCCTCTGG<br>TAATTGATTCGCGTTGTGGTACATTGGATACAATTTTAACAAAAGAACACCAGAAGGTTCGGTACATGTACAT<br>CCATCAAAATGAAGATGGAGGGTGGGGATTCGGGATGATTGGAAGTGCTTTGAACTATGTGGCTCTAAGACTT<br>GTTGGAGAAGCTTCACCTGATGATGATGCCCTTGCTAGAGGCCATAAATGGATATTTGATCACACATCCATCC<br>CTTCATGGGGCAAGCTTTATCTCGCGGTATATATGGTATATCAATGGGAAGGCTGCAATCCTCTCCCTCCAGA<br>ATTTTGGCTATTTCCATCCGATCTTGTTTTGTCTTTGAGAAAAGAGATTCATACTCTTCCTTACCACCAGATT<br>AACTGGAACAAACAACGACATAACTGTTGCAAGGTGGATCTATACTACCCTCACTCTTCCATACAAGATCTGC<br>TGCATTATTTTTGTGAACCGCTTGTCAGAAAATGGCCTTTCAATAAGCTAAGAGAGAAGGGTGTTGACCTAAT<br>GCGTTATGGATCCGAGGAAGGACGTTACATAGGCATGGGTTGTGTTGACAAGGCTTTACAAATGATGTGTTTT<br>TATCCGAATGGGAATGACTTCAAACGCCACCTGGCTAGAGTCCCTGATTACTTGTGGGTGGCAGAAGATCAAA<br>GTTTCGGTAGTCAGTTGTGGGATTGCACTCTTGTAACTCAAGCAATTATGGCCACTAATATGGTTGATGAGTA<br>TGGAGATTCTCTCAAAAAAGCCCACTTTTACTTAAAACAGTCACAGATTAAAGAGAACCCTAAAGGCGATTTC<br>ACAAAAATGTGTCGTTTATTCACGAAAGGGGCATGGACTTTCTCTGATCAAGATCATGGATGGGTCGTATCAG<br>ATTGCACTGCAGAAGCTTTGATTTGTTTATTGGCATTGTCACAAATGCCACAAGATATTGCTGGCGAAAAGGC<br>TGAAGTTGATAGATTATATGATGCTGTGAATGTCCTCCTTTACCTACAAAGTCCTGAAAGCGGTGGTTTCGCA<br>ATTTGGGAGCCACCAGTTCCAAAACCATATTTACAGATGTTGAACCCTTCGGAACTTTTTGCAGACATAGTGG<br>TTGAGAAAGAGCATGTTGAATGTACTGGATCCATAATCCAAGCATTGAATTCATTCAAAAATCTGCACCCAGG<br>GCACCGTGAGAAAGAAATAGAAGCTGCTATCGAGAAAGGCATACGTTTTTTGGAAAATAAACAACAAAATAAT<br>GGCTCATGGTATGGTTATTGGGGTATTTGTTTTCTCTATGGCACATTTTTTGTGCTGCAAGGTTTAGTATCAT<br>GTGGGAAAACATATGAAAATAGTGAAACTGTTCGGAAAGCTGTCAATTTTTTGCTCTCAACACAGAATTCAGA<br>AGGTGGTTGGGGAGAGAGTTTTGAGTCGTGCCCACAAGAGAAATTCATACCTTTGGAGGAAAATAGAACAAAT<br>TTGGTGCAAACTTCATGGGCGATGCTTGGTCTTCTCTATGGTGGACAGGTTGAAAGGGATGTAACACCGTTAC<br>ACAAGGCAGCAAAATTGCTCATTAATGGACAATTGGATAATGGAGATTTTCCTCAACAGGAAATAACGGGAGT<br>GTACATGAAAAACTGCATGTTACATTATCCAGAATATAGGAACACTTTTCCGTTAGGAGAATACCGCAAGCTT<br>GTTTGGTTGCCAAAACAACAAGTCTAA |
| >CoOSC 18_CDS (gDNA) | ATGTGGAGCTTGAAGATAGCAGAAGGGAATGATCATCCTTACTTGTATAGCACCAACAGTTTTGTTGGTAGGC<br>AAATTTGGGAGTTTGATCCCGATGCCGGTACTCCTCAAGAAAAACAACAAGTCAACGCTCGTCAACATTTTAG<br>AAACAATCTAAGCAAAGGTGTTCATCCATGCAGTGATTTGCTTATGCGGATGCAGTTGATGAAAGAGAATGGA<br>ATCGATTTATTGAGTATAGCACCATCAAGATTAGGAGAGAATGAGGAAGTGAATTATGAAGCAGTAACAAGAG<br>CAGTGAAGAAAGCAATCCGAATGAACCGTGCAATTCAAGCAAAGGACGGTCCCGAAAACGCTGGTCCTATGTT<br>TTTCACTCCTCCCCTCCTTATTGCAATGTACATAAGTGGAAACATCAACACACATTTAACCAAAGAACACAGG |

| | |
|---|---|
| | ACCGAAATGATACGTTATATCTACAATTATCAAAACGAAGATGGAGGGTGGGGATTTTATATCACAGCAGATC<br>GGGTCTGCTTTGAGCTATGTAGGTTACTAGGAGAGGGGCCCAATGATGGAAATGGTGCGGTTGAACGAGGCAG<br>GAAGTGGATACTTGACCATGGTGGTGCAACCTCCATTCCCTCTTGGGGAAAAACTTATCTCTCAGTTCTAGGA<br>GTATATGAATGGGAAGGATGTAATCCACTTCCACCAGAATTTTGGCTTTTCCCTCAAGCTTTACCATATCATC<br>CAAAAATGTGGTGCTATTGTCGGACAACTTATATGCCTATGTCATACTTATACGGGAGAAAATTTCACGGTCC<br>AATCACTGATCTTGTTCTACAAATTCGAAAAGAAATTCATCCGATCCCGTACAATGAGATAAATTGGAATAAA<br>CAACGACATAATTGTTGCAAGGAAGATCTCTACTACCCTCATTCAACGGTACAAGATCTATTGTGGGATGGTC<br>TTCATTACTTAAGTGAGCCAATTTTTAAGTTTTGGCCTTTTACAAAGTTAAGAGAAAGAGGTCTCAAAAGAGC<br>AGTTGAATTAATGTGATATGGTGCTCAACAAAGCAGATATATTACCATTGGATGTGTAGAAAAGAGCTTGCAA<br>ATGATGTGTTGGTGGGCGGAGAATTCAAACGGGGTTGAATTCAAGCGTCATCTTGCTAGAGTGCCCGATTATT<br>TATGGTTAGCAGAAGATGGAATGAAGATGCAAAGTTTTGGAAGCCAAGTATGGGATTGTGCACTCGCAACACA<br>GGCAATAATAGCTAGTAATATGGTTGAAGAATATGGCGATTCACTTAAAAAAGCCAATTTCTACATCAAAAAA<br>TCTCAAATCAAGTATAATACGTGTGGTGATTTTAGTAAAATGTGTCGGCAGTTTACTAAAGGTTCGTGGACTT<br>TCTCTGATCAAGATCATGGCTGGGTTGTATCGGATTGCACAGCTGAAGCATTGAAGTGTCTTTTATTACTATC<br>CCAAATGCCAGAGGAAATTTCGGGAGACAAAGCAGATAATGAGAGATTATATGAAGTTGTTAATGTCCTTCTT<br>TACTTACAGAGTCCTACAAGTGGAGGTTTTGCTATTTGGGAGCCACCTGTCCCACATCCATATATGCAGATGC<br>TGAATCCTTCAGAACTATTTGCAGATATTGTTGTTGAGAAAGAGCATGTTGAGTGCACTGCATCGATTATTCA<br>AGCCATTTTAGCCTTTAAACGGTTGCATCCACAACACAGGGAGAAAGAGATTGAAATTTCTGTGGCAAAATCA<br>GTTACTTTTCTGGAAGGAAAACAACGGGATGATGGTTCGTGGTATGGTTATTGGGGAATATGCTTTTTATATG<br>GCACGTTCTTTGCTATATGAGGGTTAGAAGCGGCTGGAAAAACATATAATAACAGCGAAGCAGTTTGTGAAGC<br>AGTTAATTTTTTTCTTTTGACACAAAATGAAGAAGGTGGTTGGGGAGAAAGCTTCAAATCCTGCCCTAGTGAT<br>TTATACACACCGTTGGATGGAAATCGAACAAATCTAGTTCAAACATCATGGGCTATGCTTGGTCTTATGTTAG<br>GTGGACAAATTGAGAGAGATCCAACACCAATGCATAAAGCAGCAAAGATATTAGTTAATGCACAGACGGATAA<br>TGGAGATTTTCCCCAACAGAAGATTACTGGAGTATACATGAAAAATTGCACGCTACATTATCCGGAATACAGG<br>AACATTTTCCCGCTTTGGGCACTTGGGGAATATCGCAAACGGGTTTGGGTCAACTAA |
| >CoOSC<br>19_CDS<br>(gDNA) | ATGTGGAAGTTGAAGATAGCAGAAGGGAACGATCCTTACTTGTTTAGCACCAACAACTTTGTTGGACGCCAAA<br>CTTGGGAGTTCGATCCCAATGCCGGTACTACTGAAGAAAAACAAGAAATCGAAAACGCTCGCCAATATTTCTT<br>GAATAGGCAAAAGGAAGGTTATCAAGCATCTAGTGATTTGCTCACGCGGATACAGTTGACCAAGGAGAACGGG<br>ATTGACTTATTGAGCATACGACAGGCAAGACTGAGAAATGACGAAGAAGTGAATTATGAAGCTGTGACGACTG<br>CCGTGAAAAAAGCAGTCCGATTCCAACGTGCGATACAAGCAAAAGATGGCCATTGGCCTGCTGAACAAGCCGG<br>CCCTTTGTTCCTCACTCCACCACTTATTATTGTGTTATACATCAGTGGGGCCATCAATACGCATTTAACAAAA<br>GAACACAAGAATGAGATGAAACGTTTTATCTACAATCACCAAAATGAAGATGGAGGGTGGGTATTTCATATTG<br>AGGGACATAGCACCATGATGGGATCTGTATTAAACTACATCTCCCTACGATTGTTAGGAGAAGGACCCGATGA<br>CAAAAACGGTGCAGTAGCTCGAGGAAGGAAGTGGATACTTGACCATGGTGGTGCAACCTCGATTCCGTCTTGC<br>GGCAAAACTTATCTCTCGGTCCTAGGACTGTACGAATGGGAGGGGTGCAACCCAATGCCACCGGAATTCTGGA<br>TTTTTCCAGAAGTGTTTCCTTTTCATCCTGCGAAAATGTGGTGCTATTGTCGGACCGCCTATATGCCAATGTC<br>ATATTTATATGGTAGAAGATACCATGGTCCGATCACTGATCTAGTTCTTCATCTAAGACGAGAGATTTATCGC<br>GATCCTTATGACAAGATAAACTGGAACAAACAACGCCATAATTGTTGCAAGGAAGATCTCATTTACCCTCATT<br>CAACACTTCAAGATTTGTTGTGGGATGGTCTTCACTATTTTTGTGAACCATTTATCAAGTATTGGCCTTTCAC<br>CAAACTACGAGAAAGAGCCCTCAAAAGAGCAATTGATTTGATTCGTTATAGCGCCCAAGAGAGTAGATACATC<br>AACATGGCATGTGTTGAAAAGTGTTTGCAAATGATGTGTTGGTGGGCAGAGAGCCCGAATGGGGATGAGTTCA<br>AACATCACCTCGCTAGAATGCCCGATTTCTTGTGGCTTGGAGAAGACGGAATGAAGATGCAAACATTTGGAGG<br>TCAACTATGGGATTGCACATTAGCAACTCAAGCAATAATATCAAGTAATATGCCAGAAGAATATGGGGATTCA<br>CTTAGAAAGGCCAATTTCTATATCAAAGAAACTCAAATCACGAGTAATCCGAGTGGAGACTACAGTAAAATGT<br>ATCGACAATTAAGTAAAGGGGCATGGGGATTTACTGATAAAGATAACGGTTGGGCTGTCTCAGATTGCACATC<br>TGAAGCACTGAAGTGTCTTCTCTTACTATCTGAAATGCCCGAGGAAATATCGGGAGAAAAAGCTGATAATGAG<br>CGATTGTACGATGCTATTACTTTCCTATTTCATGTACAATCTCCTACTACCGGAGGTTTTGCTATTTGGGAAA<br>AACCGATACCACAACCATATTTAGAGAACTTGAATCCTTCAGAAGTGTTTGCCGACATTGTTCTTGAGAAAGA<br>GCATCTTGAGAATACAGCGTCGATAATCCAAGCTCTAGTAGCCTTCAGACGTTTACACCCATCACATAGGCCC<br>ATAGAAATAGAAAATTCAGTGGCAAAAGCAGTGCATTTTGTTGAGAAAAGACAACTGCCTAACGGTTCATGGT<br>ACGGCTATTGGGGAATATGTTTTATATATGGCACATTCTTTGCATTAGAAGGCTTAGAAGCTGCTGGAAAAAC<br>ATATAACAATAGTGAAACAATTCGTAAAGGAGTAAAGTTTCTCCTTTCGACACAAAATGAAGAGGGTGGGTGG<br>GGAGAGAGCTACAAATCCTGCCTAAGTGAAGTCTTCACACCGTTGATCGAGAATCGAACAAACTTGGTTCAAA<br>CCGCGTGGGCCATGCTCGGGCTTATGTCAGGTGGACAAGCTGAGAGAGATCCAACACCCTTACATAAAGCAGC<br>AAAACTCTTGATTAATGCACAAATGGATAACGGTGATTTCCCACAACAGGAGTTTACTGGAGCTTCGTTAAGA<br>AATTGCATGTTACATTATCCATTGTACAGAAATTCTTTCACGCTGCGGGCACTTGCGGAATATCGAAAGCGAC<br>TTTGGACTATAAACTAA |
| >CoOSC<br>20_CDS<br>(gDNA) | ATGTGGAAACTGAAGATAGCAGAAGGGAAAGATCCTTACTTGTTTAGCACTAACAACTTTGTTGGGCGTCAAA<br>CTTGGGAGTTCGATCCCGATGCCGGTACTCCCGAAGAAAAACAAGATGTCGAAAATGCACGTCAATTTTTCCT<br>CGGAAGGCAAAAGGAAGGTTTTCAAGCATGTAGCGATTCGCTCATGCGGATGCAATTGACCAAGGAGAACGGG<br>GTTGACTTATTGAGCATACCACCGGCAAGGTTGAGAGATGAGGAACAAGTGAATTATGTAGCGGTGACGACTG |

| | |
|---|---|
| | CCGTGAAAAAAGCAGTCCGATTTCAACGGGCCATACAAGCAAAAGATGGTCACTGGCCTGCTGGACATACCGG<br>CCCTTTGTTCTTCACTCCACCACTTATAATTGTGTTATACATCAGTGGGACCATCAATACACAATTAACAGAA<br>GAACACAAAAAGGAGCTGAAACGGTATATTTACAACCACCAAAATGAAGATGGAGGGTGGGGATTTCATATTC<br>ATGGACATAGCACCATGTTGGGTTCTGTGCTGAACTACATCGCCCTACGGTTGTTAGGAGAAGGACCCGATGA<br>TGGAAATGGCGCGGTTGACCGAGCCAGAAAGTGGATACTTGACCATGGTGGTGCAACCGCTATTCCATCTTGG<br>GGAAAATTATATCTCTCGGTGCTAGGAGTGTACGAATGGGAAGGGTGCAACCCTATGCCACCAGAATTCTGGA<br>TCTTTCCCGACGCGTTTCCTTTTCATCCCAAAATGTGGTGCTATTGTCGGACGGCCTATATGCCAATGTCATA<br>TTTATACGGTAGAAAATACCATGGTCCAATCACCGATCTCATTCTCCACCTAAGACAAGAAATTTATTCTATT<br>CCTTATGACAAGATAATTTGGAATAAACATCGCCACAATTGTTGCAAGGAAGATCTCTTTTACCTTCATTCAA<br>CAATTCAAGATATGTTGTGGGATGGTCTTCATTATTTTTGTGAACCATTTTTCAAGTTTTGGCCTTTCACCAA<br>ACTTAGAGAAAGAGCACTCAAAAGAGCCGTTGAATTGATACGTTATAGTGCCCATGAGACTAGATACATCACA<br>ATGGCATCTATTGAAAAGAGTTTGCAAATGATGTGTTGGTGGGCAGAGAACCCAAATGGGGATGAATTTAAGC<br>ATCATCTTGCAAGAGTACCAGATTTCTTATGGTTAGGAGAAGATGGAATGGTGATGCAAACATTTGGTAGTCA<br>ATTATGGGATTGCACATTTGCAACCCAGGCAATAATTTCAAGTAATATGCCCCAAGAATTCGGGGATTCACTT<br>AAAAAGGCGAATTTTTATATCAAAGAATCTCAAATCAAGAATAACCCGAGTGGAGATTTTAGTAAAATGTGTC<br>GCCAATTTAGTAAAGGTGCATGGACCTTTAGCGACCAAGATCAGGGTTGGCCAGTCTCAGATTGCACAGCTGA<br>AGCCTTGAAATGTCTTCTTTTATTATCTCAACTGCCAGAGGAAATTACAGGAAAAAAGATTGATAAAGAACAA<br>TTATATGAAGCTATTAATTTCTTATTTCATGTACAGTCTCCTACCAGTGGAGGTTTTGCTTGTTGGGAACGAC<br>CGATCCCACAACCATATTTAGAGAAGTTGAATCCTTCAGAAGTGTTTGCTGACATCGTTCTTGAGAAAGAGCA<br>TCTTGAGATTACAGTGTCGATAATTCAAGCTCTAATAGCGTTTAAATGTTTGCACCCAACACATAGGGCGAAA<br>GAAATAGATATTTCTGTCGCAAATGGAGTGCGTTATCTTGAAAATAAACAATTTACTAACGGTTCATGGTACG<br>GGTATTGGGGAATATGTTTTATATACGGCACATTCTTTGCGTTAGGAGGCTTAGAAGCTGCTGGAAAAACATA<br>TAACAACAGTGAATCGGTTCGTAAAGGAGTACAATTTCTCCTTTCGACACAAACCAAAGGGGGCGGGTGGGGA<br>GAGAGCTACAAATCTTGCAATACCGAAGTGTACACACCGTTGACCGATAACCGAACAACGGTTGTTCAAACCG<br>CGTGGGCCATGCTTGGGCTTATGTCGGGTGGACAAGCTGAGAGAGATCCAACACCATTACATAAAGCCGCAAA<br>ACTTTTGATTAATGCACAAATGGATAATGGCGATTTTCCGCAAGAGGAGTTTACTGGTGCTTCTATGAGGAAC<br>TGTATGTTACATTATCCGTTATACAGGAATTCTTTCACGCTTCGGGCCCTTGCGGAATACCGAAACCGTATTT<br>GG |
| >CoOSC21_CDS (gDNA) | ATGTGGCGATTAAAGATAGGAGAAGGAGGAAACGATCCACACCTTTACAGCACAAACAACTTTGTTGGTAGGC<br>AAACATGGGAATTTGATCCAAATTATGGAACACCCGAAGAAATTGATGAAGTTGAAGAAGCTCGTCTTCATTT<br>CTGGAATAACCGTCACCAAGTTAAGCCAAGTAGCGATCTCCTTTGGCGCATGCAGTATTTAAGAGAAAAGAAG<br>TTCAAACAAACTATACCCCAGGTGAAGATTGAGGATGGTGAAGAGATATCTTATGAAAAAGTAACTGCCACAT<br>TGAGGAGGTCTGTTCACTTATTTGCTGCTTTGCAAGCTGAAGATGGTCACTGGCCTGCTGAAAATGCTGGTCC<br>CATGTATTTCATCCAACCATTGGTTATAAGCTTGTATATCACAGGGCATCTTAACAGTGTGTTCCCATCAGAA<br>CACCGAAAAGAAATTCTTCGATACTTGTATTGTCATCAGAACGAAGATGGAGGGTGGGGATTGCACATGGAAG<br>GTCATAGCATAATGTTTGGCACAACTTTGAGTTACATAACCATGCGTCTTTTAGGAGAAGGAGCTGATAATGG<br>TGCATGCACCAAAGCAAGAAAATGGATACTTGATCATGGCAGTGTGACTACCATTCCCACTTGGGGGGAAATA<br>TGGCTTTCAATATTTGGTGTTTATGAGTGGATAGGATGCAACCCAATTCCCCCAGAGTTCTGGATCCTCCCTT<br>CCTTTCTTCCCATGCATCCAGGTAAAATGTGGTGTTACTGTAGGCTGGTATACATGCCGATGTCATACCTTTA<br>CGGGAAACGATTTGTGGGCCCCATCACTCCTCTTGTGTTACAACTTCGGGATGAACTTTACACACAATCCTAC<br>AATGAAATCAATTGGAAGAGTATACGGAATTTGTGTTGTAAGGAGGACCTATACTACCCTCATCCTGTGATAC<br>AAGATCTCATATGGGATGGTCTTTACTATTTTACCGAGCCTCTTTTATGCCGATGGCCATTTAACAAATTGCG<br>AGAGAAAGCACTTGAAACAGCCATGAAGCATATCCATTATGAAGAAGAGAATACCCGATATAACACAATAGGA<br>TGTGTCGAGAAGGTACTATGTATGCTTGCAAGTTGGGTTGAAGATCCAAATGGGGTCCACTTTAAGAAGCACC<br>TCGCCCGAATCCCAGATTATATCTGGGTTGCTGAAGATGGTATGAAAATGCAGAGTTGTGCTGGATCTCAAGA<br>ATGGGATACCGGTTTGGCCATCCAAGCTCTTTTAGCTACTAATCTTACCGACGAAATTGGCCCTACATTGAAG<br>AAAGGACATGACTTTATTAAAGCATCACAGGTTAAGGACAATCCTTCTGGTGACTTTAAAAGCATGTATCGTG<br>CTATGTCCAAAGGAGCCTGGACTTTTGCTAGTCAAGATCACGGATGGCAAGTTACAGATTGTACCGCTGATGG<br>ATTGAAGTGTTGCCTCCTATTGTCAATGATGCCGCCAGAAATTGTAGGCAAGAAGATGGAGGCCGAGAAACTG<br>TATGATGCTGTCAACATATTACTTTCCATGCAAAGCAAAAATGGCGGCATGTCAGCATGGGAACCAGCAGGAT<br>CATCAAAATGGTTGGAGATGCTCAATCCTACAGAGTTATTTGCTGACATAGTCATTGAGCATGAGTATGTTGA<br>ATGCACATCATCAGTAATGCAAAGTCTTGTTCTGTTTAAGAAGATGTACCCTCAACATAGGAGGCAAGAGATA<br>GGAAGCCTCCTCACACGTGCGACTACATACTTGGAAGCAACGCAGATGCCAGATGGTTCATGGTACGGAGAGT<br>GGGCTGTGTGTTTTATATATGGCACCTCGTTTGCACTCGAAGGACTGGCACCAATTGGAAAGACGTTTGAAAA<br>TTGTCTTGCAATTCGTAAAGATGTGAATTTTCTGCTTAACACATAAAAGGAGGATGGTGGTTGGGGAGAAAGC<br>TATTTATCATGTTCTGACAAGAAATATGTAGCTCTAGAAGGAGGTCGGTCCAACTTGGTACAAACTGCATGGG<br>CTATGATGGGATTGATACATTCTAAACAAGCGGAGAGAGACTCTACACCCTTACATAGAGCTGCCAAGTTGTT<br>GATCAATTCCCAAACGAAACATGGTCATTTTCCACAACAGGAAGTAAGTGGAGCTTTCAAGAAGACATCTCTG<br>TTGCACTATCCATGCTACAGGAATCACTTCCCAATTTGGGCTCTAGCTGAATATCGTAACCAAGTGCTGCCAA<br>AACCCACATGCATCTAG |

| | |
|---|---|
| >CoOSC22_CDS(gDNA) | ATGTGGAGATTAAAGATAGGAGAAGGAGGAAATAATCCACATCTTTACAGCACAAATAACTTTGTCGGTAGGC<br>AAACATGGGAATTTGATCCAAATTATGGAACACCCGAAGAAATTGATGAAGTTGAAGAAGCTCGTCTTCATTT<br>CTGGAATAACCGTCACCAAATTAAGCCCAGTAGCGATCTCCTTTGGCGCATGCAGTTTTTAAGAGAGAAAGAA<br>TTCAAACAAAGCATACCACAGGTGAAGATTGAGGATGGTGAAGAGATATCTTATGAAAAAGTAACGGCCACAT<br>TGAGGAGGTCTGTTCACTTATTTGCTGCTTTGCAAGCTGAAGATGGTCACTGGCCAGCCGAAAATGCTGGTCC<br>CATGTATTTCATCCAACCATTGGTCATATGCTTGTATATCACACGACATCTTAACAGTGTGTTCCCATCAGAA<br>CACCGAAAAGAAATTCTTCGATACTTGTATTGTCATCAGAACGAAGATGGAGGGTGGGGATTGCACATGGAAG<br>GTCATAGCATAATGTTTGGCACAACTCTGAGTTACATTTGCATGCGTCTTTTAGGAGAAGGACCTGATAATGG<br>TGCATGTACCAAAGCAAGAAAATGGATACTTGATCATGGCAGTGTCACCACTATTCCCACTTGGGGGAAAATA<br>TGGCTTTCAATATTTGGAGTTTATGAGTGGATAGGATGCAACCCAATTCCCCCAGAGTTCTGGATCCTCCCTT<br>CCTTTCTTCCCATGCATCCAAAAATGTGGTGTTACTGTAGGCTGGTATACATGCCGATGTCATACCTTTACGG<br>GAAACGATTTGTGGGTCCAATCACTCCTCTTGTGTTACAACTCAGGGAGGAACTATACACACAACCATATAGT<br>GAAATCAAGTGGAAGAGTATAAGGAATTTGTGTTGTAAGGAGGACCTATACTACCCACACCCTTTTATACAAG<br>ATCTAATATGGGATGGTCTTTACTATTTGACCGAGCCTCTTTTATGCCGATGGCCATTTAACAAATTGCGACA<br>GAAAGCACTTGAAACAGCCATGAAACATATCCATTATGAAGAAGAGAATACTCGATATATCACAATAGGATGT<br>GTGGAGAAGGTATTATGTATGCTTGCAAGCTGGGTTGAGGATCCAAATGGGGTCCACTTTAAAAAGCACCTCG<br>CTCGAATCCCAGATTATATCTGGGTTGCTGAAGATGGAATGAAAATGCAGAGTTGTACTGGCTCTCAAGATTG<br>GGATGCCGGTTTGGCCATCCAAGCTCTTTTAGCTACTAATCTTACCGACGAAATTGGGCCTACACTGAAGAAA<br>GGACATGACTTTATTAAAGCCTCACAGGTTAAGGACAATCCTTCTGGTGACTTTAAAAGCATGTATCGCCCTA<br>TATCCAAAGGAGCCTGGACCTTTCCTAGTCAAGATCATGGATGGCAAGTTTCAGATTGTACTGCTTATGGATT<br>GAAGTGTTGCCTTCTATTCTCAATGATGCCGCCAGAAATTGTAGGCAAGAAAATGGAGGCCGAGAAACTATAT<br>GATGCTGTCAACGTATTACTTTCCATGCAAAGCAAAAATGGCGGGATGTCAGCATGGGAACCAGCAGGAACAT<br>CAAAATGGTTGGAGATGCTCAATCCTACAGAGTTATTTGCTGACATAGTCATTGAGCATGAGTATATTGAATG<br>CACATCATCAGTAATGCAGAGTCTTGTTTTGTTTAAGAAGATGTACCCTCAACACAGGAGGCAAGAGATAGAA<br>AGCCTACTCACACGTGCGACTACATACTTGGAAACAATGCAGATGCCAGATGGATCATGGTACGGAGAGTGGG<br>GTGTGTGTTTTATATATAGCACCTCGTTTGCACTTGAAGGACTGGCAGTAGTTGGAAAGACGTTTGAAAATTG<br>TCTTGCAATTCGTAAAGGTGTGAATTTTCTGCTCAACACACATATGGAAGATGGCGGTTGGGGAGAAAGTTAT<br>TTATCATGTTCCGAGAAGAAATATGTAGCTCTAGAAGGAGGTCGGTCCAACTTGGTACAAACTGCATGGGCTA<br>TGCTGGGGTTAATACAGTCTAAACAGGCGGAAAGAGACCCAACGCCCTTACATAGAGCTGCCAAGTTGTTGAT<br>CAATTCCCAAACGAAAAATGGTGATTTTCCACAACAGGAAGTAAGTGGAGCTTTCAAAAAGACATCTCTGTTG<br>CACTATGCATGCTACAGGGATTACTTCCCAATTTGGGCTCTAGCTCAATATCGTAACCAATTGCTGCCACAAC<br>TCACAACAAATTAA |
| >CoOSC23_startCDS(gDNA) | ATGTGGAAGCTGAAGATAGCAGAAGGAAACGATCCTTACTTGTTTAGCACCAACAACTTTGTTGGGCGTCAAA<br>CTTGGGAGTTCGATTCTGATGCTGGTACTCCCGAAGAAAAACAAGATGTCGAAAATGCACGTCAATATTTCAT<br>CAGCAGGCAAAAGGAAGGTTTTCAAGCATGTAGCGATTTGCTCATACGGATTCAGTTGACCAAGGAGAACGGG<br>GTTGACTTATTTAGCATACCACCGCCAAGGTTGAGAGATGATGAAGTAAATTATGAAGCGGTGACGACTGCCG<br>TGAAAAAAGCAGTCCGATTTCAACGGGCCATACAAGCAAAAGATGGGCATTGGCCTGCTGGACATGTGGGCCC<br>ATTGTTCTTCACTCCACCATTTATAATTGTGTTATACATCAGCGGGACCATCAATACACAATTAACAAAAGAA<br>CACAAAAAGGAGATGAAACGGTATATCTACAACCACCAAAATGAAGATGGAGGGTGGGGATTTCATATTGAGG<br>GACATAGCACCATGATGGGATCTGCATTGAACTACATCACCTTACGTTTGTTATGAGAGGGACCCGATGATGG<br>AAATGGCACGGTTGACCGGGCCAGAAAGTGGATACTTGACCATGGTGGTGCAACCGCTGTTCCATCTTGGGGC<br>AAAACTTATCTCTCGGTGCTAGGAGTGTACGAATGGGAGGGGTGCAACCCAATACCACCAGAATTATGGATTT<br>TTCCCGAAGCGTTTCCTCTTCATCCA |
| >CoOSC23_endCDS(gDNA) | TTGATGAAAGAGAATGGAATAGATTTATTGAGCATACCACCAGTAAGGTTAGGAGAGAATGAGCAAGTGAATT<br>ATGAAGCAGTGACGATATCAGTGACGAAAGCAATTCGATTGAACCGTGCCATTCAAGCAAAGGATGGTCACTG<br>GCCTGCTGAAGCGATGTTTTTCACTCCTCCCCTCCTTATTGCTATGTACATCAGTGGAGCCATTGACACACAT<br>TTAACCAAAGAACACAAGACCGAAATGATACGTTATATCTACAACCATCAAATTAGACAAGAAATTCATGTGA<br>TCCCTTACAATGAGATAAATTGGAATAAACAACGGCATAACTGTTGTAAGGAAGATTTGTATTACCCTCATTC<br>AACGGTACAAGATCTTTTGTGGGATGGTCTTCAGTACTTAAGTGAACCGATTCTTAAATATTGGCCTTTTACA<br>AAGTTAAGAGAAAGAGGTCTCAAAAGAGCAGTTGAATTAATGCGATATGGTGCTCAAGAAAGCAGATATATGA<br>CCATTGGATGTGTTGAAAAGAGCTTGCAAATGATGTGTTGGTGGGCAGAGAACCCTAATGGGGATGAATTTAA<br>GCATCATCTTGCTAGAGTGCCTGATTATTTGTGGTTAGCAGAAGATGGAATGAAAATGCAAAGTTTTGGGAGC<br>CAATTATGGGATTGTGTACTTGCAACTCAAGCAATAATAGCTACTGATATGGTTGAAGAATACGGCGATTCGC<br>TTAAAAAAGCTCATTTTTATATCAGAGAATCTCAAGTTAAACAAAATCCTACAGGTGATTTTAGTAAAATGTG<br>TAGACAGTTTACTAAAGGGTCATGGACTTTTTCTGACCAAGATCAAGGTTGGGTTGTCTCAGATTGCACAGCT<br>GAAGCATTAAAGTGTCTTTTGTTACTATCCCAAATGCCAGAGGAAATTTCGGGAGAAGAAGCTGATAATGAGA<br>GATTGTATGAAGCTGTGAATGTTCTTCTTTACTTACAGAGTCCTATAACAGGAGGTTTTGCTATTTGGGAGCC<br>ACCTGTCCCACAACCATATTTACAAATGTTGAATCCTTCAGAACTTTTCGCAGATATTGTTGTTGAGAAAGAG<br>CATGTTGAGTGTACAGCATCAATTATTCAAGCACTTTTAGCCTTCAAACGGTTGCACCCAGGGCACAGGGAGA<br>AAGAAATTGAAATTTCCTTGGCAAAAGCAGTTACTTTTGTGGAGGGAAATCAACGGCATGATGGTTCATGGTA<br>TGGTTATTGGGGAATATGCTTTCTATATGGCACATTCTTTGCTGTAGGAGGCTTAGTTTCTGCTTGGAAAACA |

| | |
|---|---|
| | TATAATAACAGTGAATCAATTCGTAAAGCAGTTAATTTTTTTATTTTGACACAAAATGAACAAGGCGGTTGGG GAGAAAGTATCAAATCTTGTCCCACTGAAGTATACACACCGTTGGATGGCAATCGAACAAACCTAGTTCAAAC ATCATGGGCTATGCTTGGTCTTATGTTAGGTGGACAGGTTGAAAGAGATCCAACACCTTTGCATAAAGCAGCA AAGATATTAATTAATGTACAGATGGATAATGGAGATTTTCCTCAACAGGAGATTACTGGAGTTTACAATAAGA ATTGCATGCTACATTATCCAGAATACAGAAACATCTTCCCACTTTGGGCACTTGGGGAATATCGCAAACGAGT TTGGATCAATTAA |
| >CoOSC 24_CDS (gDNA) | ATGTGGGAGTTAAAGATAGCTGAAGGGGATGGCCCTTATTTGTATAGTACCAACAACTTTGTCGGTAGACAAT TTTGGGAATTTAATCCAGATGCTGGAACTTTAGAGGAGAAACAAGAGATTGAAATGGCTCGCCAGAAGTATAA AAATAATCGAAGAAATGGAGGATTTCATGCTTGTGGCGACCTTCTCATGCGGAGGCAGATGATAAAGGAAAAC GGAATAGATCTTACGAGCATAGCTCCTATGAGAGTAAAGGAACACGAACATGTCAACTTTGAAGCTGTAACAA CCGCAGTTAGGAAAGCGGTTAGATTACAACGAGCCATCCAAGCAAAAGATGGCCCTATGTTTTTCACTCCTCC ACTTGTAATTGCTTTATACATTAGCGGTATGATTAATATAATCTTGAGTGAAGAACATAAGAAAGAGATGATA AGGTTCTTCTACAATCATCAGAATGAAGATGGGGGTACGATGATTGGATCTGCATTGAGTTACATACTAGGAG AGGCAGAAGATGGGGCCATTTCAAGAGGCCGCAAGTGGATACTTGATCATGGTGGTGCAACCTCTATTCCCTC TTGGGGAAAGGTTTATCTTTCGGTCCTTGGAGTGTATGAATGGGACCCACTGCCACCAGAATTTTGGCTATTT CCATCGGCGTTCCCTTTTCATCCAGAGATGTGGTGCTACTGTTACATGCCAATGTCATATTTATACGGAAAAA GAATCCAAGGACCAATCACAAATCACCCCACCCCTTTTGAGGATATTAGTTGGAATAAACAACGGAATAACTG TTGCAAGGAGGACTTCTACTATCCACATTCGTTTCTTCAAGATGCATTGTGGCATAGCCTTACTGAGCCGGTT CTCAAACATTGGCCATTTTCTAAACTACGAAACAGAGCTGTTGACAGAGTTGTTGAACTGATGCGCTATGAAA CAAGATACATGACCATTGAAAAAAGTTTACAAATGATGTGTTGGTGGGCAGAGAATCCAAATGGGGATGAGTT CAAATATCACCTTGCAAGAGTACCGGATTACTTGTGGATTGCAGAAGATGGGATGACAATGCACAGTTTTGGC AGTCAAGTGTGGGATTGCTCTCTTGCAATTCAAGCAATTGTTGCAAGTAATATGGCTGACGAATACGGTGATT GTCTCAAAAAGGCACACTTTTACTTAAGAGAATCACAGGTAAAAGAAAATCCTTCAGGCGATTTTACTCGGAT GTGCCGACAGTTCACTAAAGGATCATGGATGGACTGTCTACAGTTATTTTGTCCAATAGTGTTATTATCAAAC ATGCCGAAAGAAATAGCTGGAAACAAAGACAACACCGCCCGACTCTATGATGCAGTGAACAGTGGAGGATTTG CGGCACCAATTCCAAAGCCATTTCTACAGTTACTTAATCCTTCCGAGATTTTTGCAGACATTGTGGTTGAGAA AGAGATGATTGTGGAGACCACATCTTCCATTATCGGAGTTCTAATGGAGTTCAATCGCACCCACCCAGAAAGG AAGAAATTGATTTCAAAAGGAATACGCTATCTTGAGGAAACATGGGTTGATAGGTATGGGTACTGGGGAGTAT GCTTCATATATGGAACATTCAGGGGTTTAACTTGTGTCGGAAAAACATATGACAACAACGAAGCAGTGTGTAA GTTCTTACTTTCAATACAGAATGAAGAAGGGGGTTGGGGGGAGAGCCTCTTATCTTGCCCTACCGAGGTATAT ACACCGCTAGATGGAAACCGAACAAACTTGGTGCAAACCTCATGGGCTATGCTTGGTCTTATGTTTACGGTGC AAAGAGATCCGATGCCATTGCATAAAGCGGCAAAATTGTTGATTAACGCCCAGATGGATAACGGAGATTTTCC TCAACAGGAAATTACAGGACTCTACATGAAGAACTGCTTGCTCCTCTACGCACAATACAGGAACATATTTCCA CTTTGGGCACTTGGGGAGGAGTACCGTAAACGTGTTTGG |
| >CoOSC 25_CDS (gDNA) | CTAAGAAAGGAGAAGCAGATAGATAAAATACCCGAAGCAAACGAAACAGAAGAATTAACAAATGAAGCAGTTG CAACTACAGTCAGAAGAGCCATCAGCTTCTACTCAACCATTCAAGCTCATGATGGTGCTGAATCTGCTGGCCC TTTATTCTTCCTTCCTCCACTGGATAGCACAGGAACCATGAATGTTATTCTAACTCCTGCACATCAAGTAGAA ATCAAACGCTACTTGTATAATCATCAGAATAAAGATGGAGGTTGGGGATTACATATAGAGGGCCACAGCACGA TGTTTGGGTCAGTTTTTAGTTACATCACCTTGAGGTTGCTTGGAGAAGAAGCCGATAGCGGTGGCGGTGGTTG TAAATGGATCCTTGACCACGGTGGTGCAATTGGGACTCCTTCATGGGGAAAGTTTTGGCTCACAGTTCTTGGA GTATATGAATGGGAAGGTTGTAATCCTATGCCACCGGAATTTTGGCTCCTTCCTAAAATTTTCCCAATTCATC CAAAAATGTTATGTTATTGTCGCTTAGTTTACATGTACTTATACGGCAAAAGATTCGTCGGGAGAATAACTGG ATTAGTTCAAGCACTAAGGCAAGAGCTTTATACGGGTCCTTATCATGAGATTAACTGGAATAAAGGGCGAAAT ACTTGTGCAAAGGAAGATCTGTATTACCCGCATCCTCTAGTTCAAGATATGTTATGGGGCGTTCTTCACAACA TTGGTGAACCTATCTTAACACGTTGGCCATTTTCCAAACTACGGGAAAAGGCCCTAAAAGTTGCAATGGAGCA TGTTCACTATGAAGATCAGAGCAGTAGATATCTTTGCATTGGCTGTGTAGAAAAGGTGTTATGCTTGATTGCG ACCTGGGTGGAAGATCCAAAAGGAAACGCATTTAAACGCCATCTTGCACGAATTCCTGACTACTTTTGGGTGG CTGAAGATGGGATGAAAATGCAGAGTTTTGGCTGTCAAATGTGGGATGCAGCATTTGCTATTCAAGCTAGTAA TCTAACAGAAGAATACGGGCCGACTCTTAAAAAAGCACATGAATTTGTGAAGGCCTCACAGGTATTTTCAGGT TCGCCCCGTGGAGATTTCGGTAAAATGTACAGACACATGTCTAAAGGTGCGTGGACGTTTTCAATGCAAGACC ATGGTTGGCAAGTGTCAGATTGTACCGCAGAAGGCTTGAAGGTTGCACTTTTATATTCCCAAATGAGCCCAGA ACTTGTTGGTGAAAAACTTGAAAATGAGCATCTCTACGACGCTGTCAATGTTATTCTTTCCTTACAAAGTAAA AATGGCGGTTTTCCTGGTTGGGAACCACAAAGGGCATATTCTTGGCTGGAGAAGTTCAATCCGACGGAATTTT TTGAAGATGTGATGATTGAACGAGAGTATGTGGAATGCACTTCCTCTGCCATCCAAAGTTTGGCGCTCTTCAA GAAATTGCACCCTGGACACAGGACGATGAAGATCGAACATTGCATTTCAAAAGCGGTCAAGTACATCGTAGAC ATGCAAAACACTGATGGTTCATGGTATGGTTGTTGGGGAATATGTTACACCTATGGTACATGGTTTGCAGTAG ATGCACTAGTAGCTTGTGGGAAGAACTATCATAACAGTCCCTCCCTTCGAAAAGCTTGCCAATTTCTGCTATC AAAACAACTTCCAGATGGAGGATGGGGTGAGAGTTATCTTTCGAGTGCAAATAAGGAATATACAAATTTGGAT GGAGACCGGTCGAATTTAGTGCAAACATCTTGGGCTTTACTCTCACTTATTAAAGCTGGACAGGCTGGAGTTA ATCTCACACCAATAGATCGTCTTGTAATAAATTCACAAACTGAAGACGGAGACTTCCCTCAACAGGAAATCAC |

| | |
|---|---|
| | AGGAGTTTTTATGAAGAATTGCACCCTCAATTACTCATCGTTTCGAAATATTTTCCCCATATGGGCTCTTGGT<br>GAGTATCGGCGTATTCGCATGAAACTC |
| >CoOSC 26_CDS (gDNA) | ATGTGGAAGCTAAAGCTATCGAAAGGCGATGATGATCCTGGTGTTAGAAGCGTAAATAATCATATCGGTAGAC<br>AATATTGGGAATTCGATCCACTTGCAGGAACTCCTGAGGAACAATCTCAAATCAACAACATGCGAGAAGAATT<br>CACCAGAAACAAGGTGAAGGTGAAGCATAGTTCTGATCTTCTGATGAGATTCCAGTTTGCAAGCGAGATTAAT<br>GGTAGTGAAATGAAAAAATCACAAGTGATTGAAGCAAAAGATGAAGATGGAGAAGAAGTTGTGGTGAAGACTT<br>TGAAGAAAGGTTTGAAGTTTTACTCAAGTCTTCAAGGGGAGGATGGGAGTTGGCCTGCTGATTATGGTGGTCC<br>GTTGTTTCTCCTTCCTGGCCTGATCATTGCGTTGCATATGATGGGGGCGAAGGACAGAGCTTTATCCGTAGAA<br>CACCAAAGGGAAATCCGTCGCTATCTTTACAACCATCAGAATGTTGATGGTGGTTGGGGTTTACATATAGAAG<br>GTCACAGCACCATGTTTTGTACAGCTTTAAACTATGTGAGCCTAAGATTACTTGGAGAAAGAATGGATGGTGG<br>TGAAGGAGCCATGACCAAAGCAAGAAAATGGATTCTTGATCATGGTAGTGTTACACACATACCCTCATGGGGC<br>AAATTTTGGCTTTCCGTACTAGGGGTGTATGAATGGAGTGGCAACAATCCTTTGCCACCAGAAATGTGGCTTC<br>TCCCGTACTTTGTTCCTTTACACCCAGGTAGGATGTGGTGCCACACAAGGATGGTATACTTGCCAATGTCGTA<br>TATATATGGTAAACGATTCGTCGGACCGATCAATTCCATTGTTTTGTCACTAAGAAGAGAGTTGTATAATACT<br>CATTATTACCAAGTTAATTTGATTATTACTGAAATTTTAACGTTAAGAATATCAAAATACTCAATATTATTAA<br>TTTATTTGATGATTAATGATATTTTAACCATCATACTACTAAAGTATTATTTAAGTATACTAAAAAAAAAAAA<br>GAAAAGAAAAGAGTTATGGCCGAAAAACGAAGCGAGGGGAACAAACCCACCAAATCACAAGGTGACATGTGGC<br>ACAATGAGAGAGAAAGTACTCAACATGTTATGCTGTTGGGTAGAGGATCCAATGTCACATGTAAATAAATTGC<br>ATCTCTCTAGAATTGAAGACTATCTTTGGATAGCTGAGGATGGAATGAAGATGCAGGGATATAACGGGTCGCA<br>ATTGTGGGATTCTGTTTTTGCGGTTCAAGCAATATTAGCAACGAATCTTGTAGACGAATATGGTTCCATGCTT<br>AGAAAAGCCCACAGCTTCATAAAAAACTCACAAATTAGAGAAAATAGCTCTGGTAATATTCAGTCATGGTATC<br>GTCACATAATACGCGGTGGATGGCCATTTTCTACTCCAGATAATGGTTGGCCGGTATCAGATTGCACTGCAGA<br>AGCTTTGAAGACAGTGTTGATGCTATCGCAAATGCCAAACGACATCGTTGGAGAAGCAATTGCACCAGAATGT<br>TTATATGACGCTGGTCATTTACTTCTAACTCTTCAGAATGGAAATGGTGGATTTTCGTCATATGAGCCCATGA<br>GGTCCTACCCATGGTTAGAGGTGTTCAATCCAGCTGAAACATTTGGAGACATCATTGTGGATTACTATGTCGA<br>GTGTACGTCAGCTGTAGTGCAAGGACTAAGTTCGTTTATGAAGCTATATCCAAGCCACCGAAGGGACGAAATA<br>GAAGCATGTGTCGATAAAGCCATATCGTTCATCGAGAACGTGCAATTGCACGATGGTTCATGGTATGGATCAT<br>GGGGAATTTGCTACACTTATGGAACATGGTTTGGTATAAAAGGGTTGGTTGCAACCGGTGAAACATATGAAAC<br>TAATTATAGCATAAGAAAGGCGTGTGCGTTCTTACTCTCGAAGCAACTTCCGTCCGGTGGTTGGGGCGAGAGT<br>TACATTTCTTGCGAGCAAAAGAATTACACAAATATTGCAGGGAACAAATCCCACATTGCAAACACTTCATGGG<br>CTTTGCTAGCCTTGATTGAAGCCGGACAAGCTAAACGAGATAAAATGCCACTCCATCGTGCTGCAAAAGTACT<br>TATTGATCACCAAATGGAGAATGGAGATTTTCCTCAACAGGAAATTATTGGGGTGTTCAACAAGAACTGTATG<br>ATAAGTTACTCATCTTACCGAAACATATTTCCTATTTGGGCTCTTGGAGAGTACCTCAATCGCGTAATCAGGA<br>ACAATTTACCCTCTTGA |
| >CoOSC 27_CDS (gDNA) | CTAAAGCTATCGAAAGGCGACGATGATCCTAGTGTTAGAAGCGTGAATAATCATATCGGTAGACAATATTGGG<br>AATTCGATCCATGTGCAGGAACTCCCGAGGAACGATCTCAAATCGACAACATGCGAGAAGAATTTACCATAAA<br>CAAGGCGAAGGTGAAGCATAGTTCTGATCTTTTCATGAGATTCCAGTTTGGAAGGGAGAATTGCAGTGAAATT<br>ATGGAAAAATCACAAGTGACTGAAGATGGAGATGAAGTTGTGGTGAAGACTTTGAAGAAAGCTTTAAGGTTTT<br>ACTCAACTCTTCAAGGTGAGGATGGGAGTTGGCCTGCTGATTATGGTGGTCCTTTGTTTCTCCTTCCTGGCCT<br>GATCATTGCGTTGCATATGATGGGGGCGATGGACATAGCTTTGTCCGTAGAACACCAAAGGGAAATCCGTCGC<br>TATCTTTACAACCATCAGAATGTTGATGGTGGTTGGGGTTTACATATAGAAGGTCAGAGCACCATGTTTTGTA<br>CAGCTTTGAACTATGTGAGCCTAAGATTACTTGGAGAAAGAATGGATGGTGGTCAAGGAGCCATGGCCAAAGC<br>AAGAAAATGGATTCTTGATCATGGTAGTGTTACACACATACCCTCATGGGGCAAATTTTGGCTTTCTGTACTA<br>GGGGTGTATGAATGGAGTGGCAACAATCCTTTGCCACCAGAAATGTGGCTTCTCCCATACTTTGTTCCTTTAC<br>ACCCAAGGATGTGGTGCCACACAAGGATGGTATACTTGCCAATGTCGTATTTATATGGTAAACGATTCGTTGG<br>GCCAATCAATTCCATAGTTTTGTCACTAAGAAGAGAGTTGTATAATACTCTTTATAACCAAGTTAATTGGGAC<br>TTGGCAAGAAATCAATGTGCCAAGGAAGATTTGTATTACCCTCATCCTATGATACAAGACATTTTATGGGGTG<br>GTTTGAACAAGATTGCCGAACCACTTTTGATGCAATGGCCATTCTCAAAGCTTAGAAAGAAAGCATTAACCAC<br>GGTTATGCAACATGTTCATTATGAGGACAAAAACACTCAGTATATTTGCATTGGACCTGTTAATAAGGTGCTC<br>AACATGTTATGCTGTTGGGTAGAGGATCCAATGTCTCATGTAAATAAATTGCATCTCTCAAGAATCAAAGACT<br>ATCTTTGGATAGCAGAGGATGGAATGAAGATGCAGGGATACAACGGATCACAATTGTGGGATTCTGTATTCGC<br>GGTTCAAGCAATATTAGCAACGAATCTGGTAGACGAATATGGTTCCATGCTTAGAAAAGCCCACAGTTTCATA<br>AGAAACTCACAAATTAGAGAAAATAGCTCTGGTAATATTCAGTCATGGTATCGTCACATAATACGCGGTGGAT<br>GGCCGTTTTCGACTCCAGATAATGGTTGGCCGGTATCAGATTGCACTGCAGAAGCTTTGAAGACAGTGTTGAT<br>GCTATCACAAATGCCAAAAGACATAGTTGGAGAAGCAGTTGCACCAGAATGTTTATATGATGCTGTTCATTTA<br>CTTCTAACTCTTCAGCAGAATGGAAATGGTGGATTTTCGTCATATGAGCCCATGAGGTCCTACCCATGGTTAG<br>AGGTGTTCAATCCAGCTGAAACATTTGGAGACATCATTGTCGATTACTATGTCGAGTGTACATCAGCTGTAGT<br>GCAAGGACTAAGTTCGTTTATGAAGCTATATCCAAGCCACTGTAGGGATGAAATAGAAGCATGTGTCGATAAA<br>GCCATAACTTTCATCCAGAACGTGCAATTGCACGATGGTTCATGGTATGGATCATGGGGAATTTGCTACACTT<br>ACGGGACATGGTTTGGTATAAAAGGGTTGGTCGCAGCGGGTGCGACGTATGAAACTAACCATAACATAAGAAA<br>GGCGTGTGCGTTTTTACTCTCGAAGCAGCTTCCCTCTGGTGGTTGGGGCGAGAGCTACGTTTCTTGCGAGCAA |

| | |
|---|---|
| | AAGAATTACACGAATATCGCATGGAACAAATCCCACATTACAAACACTTCATGGGCTCTGCTAGCCCTGATTG<br>AAGCAGGACAAGCTAAACGAGATAAAGTGCCACTCCATCGTGCTGCGAAAGTACTAATTGATCACCAAATGGA<br>GAATGGAGATTTTCCTCAGCAGGAAATTATTGGGGTGTTCAACAAGAACTGTATGATAAGTTACTCATCTTAC<br>CGAAACATATTTCCTATTTGGGCTCTTGGAGAGTACCTAAATCGCGTGTGGGTGGTGTTACCAAAGTGTAATG<br>TATAG |
| >CoOSC<br>28_CDS<br>(gDNA) | ATGTGGAGCTTGAAGATAGCAGAAGGGAATGGTCATCCTTACTTGTTTAGCACCAACAATTTTGTTGGTAGGC<br>AAATTTGGGAGTTTGATCCTGATGCCGGTACTCCCGAAGAAAAACAACAAGTCGAAAACGCTCGTCAACATTT<br>TACAAACAATCGAAGCAAAGGTGTTCATCCATGCAGTGATTTGCTTATGCGGATGCAGTTGATCAAAGAGAAT<br>GGAATGGATTTATTGAATATACCACCTTCAAGATTAGGAGAAAATGAGCAAGTGAATTATGAAGCGGTAACGA<br>CAGCAGTGAAGAAAGCAATCCGATTGAACCGTGCGATTCAAGCAAAGGATGGTCATTGGCCTGCTGAAAATGC<br>AGGCCCCATGTTTTTCACTCCTCCCCTCCTTATTGCGATGTATATAAGTGGAGCCATCAACACACATTTAACC<br>AAAGAACACAAGTCCGAAATGATACGTTATATCTACAATCATCAAAACGAAGATGGAGGATGGGGATTTTATA<br>TCGGGGGAGACAGCACCATGATCGGGTCTGCTTTGAGCTATGTTTTCCTGAGATTACTAGGAGAGGGGCCCGA<br>TGATGGAAATGGTGCGGTTGGCCAAGCCAGGAGGTGGATACTTGACCATGGTGGTGCAACCATTCCCTCATGG<br>GGAAAAACTTATCTCTCGGTTCTTGGAGTATATGAATGGGAAGGATGCAATCCACTTCCACCAGAATTTTGGC<br>TTTTCCCTCGAGCATTACCATATCATCCAAAAATGTGGTGCTATTGTCGGACAACTTATATGCCTATGTCATA<br>CTTATACGGGAGAAAATTTCATGGTCCACTCACTGAACTTGTTCTACAAATTCGACAAGAAATTCATTCGATC<br>CCGTACAATGAGATAAATTGGAATAAACAGCGACACAACTGTTGCAAGGAAGATCTCTACTACCCTCATTCAA<br>CGGTACAAGATCTGTTGTGGGATGGTCTTCATTACTTAAGTGAGCCAATTCTTAAGTTTTGGCCTTTTACAAA<br>GTTAAGAGAAAGAGGTCTCAAAAGAGCAGTTGAATTAATGCGATATGGTGCTCAAGAAAGCAGATATATCACC<br>ACTGGATATATCTCAAAGAGCTTGCAAATAATGTGTTGGTGGGCGGAGAACCCAAACGGGGTTGAATTCAAGC<br>GTCATCTTGCTAGAGTGCCTGATTATTTGTGGTTAGCAGAAGATGGAATGAAGATGCAGAGTTTTGGAAGCCA<br>GTTATGGGATTGTGTACTCGCAACGCAGGCAATAATAGCTAGTGATATGGTCGAAGATTATGGTGATTCACTT<br>AAAAAAGCCAATTTCTACATCAAAAAATCTCAAATCAAGACTAATCCATGTGGTGATTTTAGTAAAATGTGTC<br>GGCAATTTACTAAAGGTTCGTGGACTTTCTCTGACCAAGATCAAGGTTGGGTTGTATCGGATTGCACAGCTGA<br>AGCATTGAAGTGTCTTTTGTTACTATCCCAAATGCCAGAGGAAATTTCGGGAGCAAAAGCCGATAATGAGCGA<br>TTATATGAAGCTGTTAATGTCCTTCTTTACTTACAGAGCCCTACAAGTGGAGGTTTTGCTATTTGGGAGCCAC<br>CTGTCCCACAACCATATTTGCAGATGCTGAATCCTTCAGAACTATTTGCAGATATTGTTGTTGAAAAAGAGCA<br>TGTTGAGTGCACAGCATCAATTATTAAAGCTCTTTTAGCCTTCAAACGGTTGCATCCACAACATAGGGAGAAA<br>GAGATTGAAATTTCTGTCACAAAAGCAGTTACTTTTCTAGAAGGAAAACAACAGGGTGATGGCTCGTGGAATG<br>GTTATTGGGGAATATGCTTTTTATATGGCACATTCTTCGCCATAGGAGGGTTAGAAGCTGCTGGAAAAACATA<br>CAAAAACAGTGAAACAATTCGTAAAGCGGTTAATTTTTTTCATTTGACACAAAATGAAGAAGGTGGTTGGGGA<br>GAAAGCATCAAATCCTGCCCTAGTGAAGTATACACACCGTTGGATGAAAATCGAACAAATCTAGTTCAAACAT<br>CATGGGCTATGCTTGGTCTTATGTTAGGTGGACAGATTGAGAGAGATCCAACACCCTTGCATAAAGCCGCAAA<br>GATATTAATTAATGCACAGATGGATAATGGAGATTTTCCTCAACAAGAGATCACTGGAGTCTACATGAAAAAT<br>TGCATGCTACATTATGCGGAATACAGGAACATTTTCCCGCTTTGGGCACTTGGGGAATATCGCAAACGGGTTT<br>GGGTCAACTAA |
| >CoOSC<br>1_(CoT<br>XSS) | MWKLKIGEKNGKLEIGDGNGDEYLYSTNNFVGRQTWEFDPSAGTQEERDQIERIREQFLNNKKKLDIHCCGDL<br>LMRTQLIKESQIDLTSELLVRLKDEEDVNYEAVTMAVKKAVLLNRAIQAWDGHWPAENAGPLFFTPPLIIALY<br>ISGTVDTILTKEHQKEMIRYMYIHQNEDGGWGFYISGKSTMIGSALNYVALRLLGEASPTDDDDIGALARGHK<br>WIFDHGGATSIPSWGKVYLAVLGVYEWEGCNPLPPEFWLFPSFLPYHPAKMWCYCRTTYMPMSHLYGKSYHGP<br>ITDLVLSLRKEIHTLPYHQINWNKQRHNCCKLDLYYPHSSIQDMLWDSLHYFCEPLVKKWPLNKLREKGLKRV<br>VDLMRYGSEEGRYIGMGCVDKALQMMCFYAEDPNGIDFKHHLARVPDYLWLAEDGMKMQSFGSQLWDCTLVTQ<br>AIMASNMVDEYGDSLKKAHFYLKQSQIKENPKGDFTKMCRLFTKGAWTFSDQDHGWVVSDCTAEALICLLALS<br>QMPQDIAGEKAEVDRLYDAVNVLLYLQSPESGGFAIWEPPVPKPYLQMLNPSELFADIVVEKEHVECTGSIIQ<br>ALNSFKNLHPRHREKEIEAAIEKGIRFLENKQQNDGSWYGYWGICFLYGTFFVLQGLVSCGKTYENSETVRKA<br>VNFLLSTQNSEGGWGESFESCPQEKFIPLEGNRTNLVQTSWAMLGLLYGGQVERDVTPLHKAAKLLINGQLDN<br>GDFPQQEITGVYMKNCMLHYPEYRNTFPLWALGEYRNRVWLPKQQV |
| >CoOSC<br>2 | MWSLKIAEGNDHPYLFSTNNFVGRQIWEFDPDAGTSEEKQQVKNVRQHFRSNLRGGVHPCSDLLMRMQLIKEN<br>GIDLLSIRPARLGENEEVNYEAITTAVKKAVRLNRAIQAKDGHWPAENAGSMISTPALLIAMYISGTINTHLT<br>KEHKTEMIRYIYNHQNEDGGWGFYIGGQSIMIGSALSYVALRLLGEGPDDGNGAVGRARKWILDHGGATAIPF<br>LGKIYLSVLGVYEWEGCNPLPPEIWLSPQALPYHPSKMLSHFRTTYMPMSYLYGRKIHGPITDLVQQLRNEIH<br>VIPYNNINWNKQRHNCCKEDLYYPHSTVQDLLWDGLHYLSEPILKFWPFTKLRERGLKRAVELMRYGAQESRY<br>ITTGYISKSLQIMCWWAENPNGDEFKRHLARVPDYLWLAEDGMKMQSFGSQLWDCALATQAIIASDMVDEYGD<br>SLKKANFYLKESQIKNNPSGDFSKMCRQFTKGSWTLSDQDQGWVVSDCTAEAMKCLLLLSQMPDEITGEKAET<br>ERLYEAVNVLLYLQSPISGGFAVWEPPVPQPYLQMLNPSELFADIIVEKEHVECTASVIQALLAFNRLHPQHR<br>EKEIEISVAKAVSFLEEKQQHDGSWYGYWGVCFLYGTFFAIGGLEAAGKTYKNSETIRKAVNFFLLTQNEEGG<br>WGESIKSCPSEVYSPLDGNRTNLVQTSWAMLGLMLGGQIERDPTPLHKAAKILINAQMDNGDFPQQEITGVYM<br>KNCMLHYAEYRNIFPLWALGEYRKRVWVN |

| | |
|---|---|
| >CoOSC3_(CoMAS) | MWKLKIAEGNDPYLFSTNNFVGRQIWEFDPDAGTPEEKQEVENARQHYRNNRKEDVHPCSDLLMRMQLIKENG<br>IDLLSIPPARLGDNEQVNYEAVTRSVTKAVRLNRAIQAKDGHWPAENAGPMFFTPPLLIAMYISGAIATHLTK<br>EHKTEMIRYIYNHQNDDGGWGFYIEGHSTMIGSALSYVALRLLGEGPHDGNGAVDRARKWIIDHGGATSIPSW<br>GKTYLSVLGVYEWEGCNPLPPEFWIFPETLPYHPAKMWCYCRTTYMPMSYLYGTKFHGPITDLVLQLRQEIHA<br>IPYNEINWNKQRHNCCKEDLYYPHSTLQDLLWDGLHYLSEPFLKYWPFKKLRERGLKRAVELMRYGAQQSRYM<br>TIGCVEKSLQMMCWWAENPNGDEFKHHLARVPDYLWLAEDGMKMQSFGSQLWDCTLATQAIIATDMVEEYGDS<br>LKKAHFYIKESQVKQNPSGDFSQMCRQFTIGSWTFSDQDHGWVVSDCTAEALKCLLLLSQMPEEIVGEKADNE<br>RLYEAVNVLLYLQSPITGGFAIWEPPVPQPYLQMLNPSELFADIVVEKEHVECTASIIQALLAFKRLHPGHRE<br>KEIEISVAKAVTFLEGKQQHDGSWYGYWGICFLYGTFFALGGLVSAGKTYNNSEEIRKAVNFFILTQNEEGGW<br>GESIKSCPTEVYTPLDGNRTNLVQTSWAMLALMLGGQAERDPTPLHKAAKILINAQMDNGDFPQQEITGVYMK<br>NCMLHYPEYRNIFPLWALGEYRKRVWVN |
| >CoOSC4 | MWELKIAEGDGPYLYSTNNFVGRQFWEFNPDAGTFEEKQEIETARENYKNNRRNGGFHACGDLLMRRQLIKEN<br>GIDLTSIAPVRVKEHEHVNFEGVTTAVRKAVRLQRSIQAKDGHWPAENAGPMFFTPPLVIALYISGTINIILT<br>EEHRKEMIRYFYNHQNEDGGWGFYIEGHSTMIGSALSYVALRILGEREDGAISRGRKWILDHGGATSIPSWGK<br>VYLSVLGVYEWEGCNPLPPEFWLFPSAFPFHPAEMWCYCRTTYMPMSYLYGKRIQGPITNLVLSLRKEIHPTP<br>FEDISWNKQRNNCCKEDFYYPHSFLQDALWHSLHHLTEPVLKHWPFSKLRNRAIDRVVELMRYESQETRYMTI<br>GCVEKSLQMMCWWAENPNGDEFKYHLARVPDYLWIAEDGMTMHSFGSQVWDCSLAIQAILASNMVEEYGDCLK<br>KAHSYLRESQVKENPSGDFTRMCRQFTKGSWTFSDQDHGWTVSDCTAEALKCLLLLSNMPKEIAGNKDNSSRL<br>YDAVNVLLYMQSPISGGFAVWEPPIPKPFLQLLNPSEIFADIVVEKEHVETTSSIIGVLMEFNSLHPRHRKEE<br>IQLSITKGIRYLEETQWHDGSWYGYWGVCFIYGTFFALRGLTCVGKTYENNEAVCKGVEFLLSIQNEEGGWGE<br>SLLSCPTEVYTPLDGNRTNLVQTSWAMLGLMFAGQVQRDPKPLHKAAKLLINAQMDNGDFPQQEITGVYMKNC<br>LLLYAQYRNIFPLWALGEYRKRVW |
| >CoOSC5 | MWKLKIAEGNDPYLFSTNNFVGRQTWEFDPDAGTPQDVENARQYFLTRQKEGFQACSDLLMQIQLTKENGVDL<br>LTIPPARLRDDEEVNYEAVTTAVKKAIRFQRAIQAKDGHWPAGHVGPLFFTPPFIIVLYISGTINTQLTKEHK<br>KEMKRYIYNHQNEDGGWGFHIEGHSTMMGSALNYIAIRLLGEGPDDGNGAVDRARKWILDHGGATAIPFWGKI<br>YLSVLGVYEWEGCNPIPPELWIFPETFPLHPSKMWCHTRTAYLPMSYLYGRKYHGPITDLILDLRQEIYPIPY<br>DKIIWNKHRHNCCKEDLFFHSTIQDMLWDGLHYFCEPFFKYWPFTKLRERALNRVVELTRYCAHESRYITTAS<br>IEKSLQMMCWWAEKPNGDEFKHHLARIPDFLWLAEDGMTIQTFGSQLWDCTFATQAIISTNMPQEYGDSLKKA<br>NFYIKESQIKNNPSGDFSSMCRQISKGAWTFSDQDQGWPVSDCTAEALKCLLLLSQMPEEITGKKIDKEQLYE<br>AINFLFHVQSPTSGGFACWERPIPQPYLEKLSLSEMFADVVLEKEHLEVTASIIQALIAFKCAHPTHRAKEID<br>ISVANGVHYLENKQLPNGSWYGFWGICFTYGTCFALGGLEAVGKTYNNSESVRKGVQFLLAAQNQEGGWGESY<br>KSCNTEVYTPLIDNRSTVVQTAWAMLGLMSGGQAERDPTPLHKAAKLLINAQMDNGDFLQQEFTGASMRNATL<br>HYPLYRNSFTLRALAEYRKRLWGIK |
| >CoOSC6 | MWKLKIAEGNDPYLFSTNNFIGRQFWEFDPDAGTPEEKQQVENARQHFLNRQKEGYQTSSDLLMRMQLTKENG<br>VDLLSIPPARLRDDEEVDYEAVTTTVRKAVRYQRAIQAKDGHWPAENAGPLFFAPPLIIVLYISGTINTQLTE<br>EHKKELKRYIYNHQKEDGGWGFHIEGHSTMMSSALNYIALRLLGEGPDDGNGAIDRARKWIVHHGGATGIPSW<br>GKAYLSVLGVYEWEGCNPMPPEFWLFPELFPFHPAKMWCYCRLTYTQMSYLYGRKYHGPITDLVVHLRQEIHS<br>IPYDKINWNKQRHNCCKEDLIYPHTTIQDLLWDGLQYFCEPLFKYWPFTKLRERALKRALELIRYSARESRYI<br>TMACVEKCLQLMTWWAEDPNGDEFKRHLARVPDYLWLGEDGMKMQTYASQVWDCTIATQAIISTNMPQEYGDS<br>LKRANFFIKESQIKNDPRGDLSKICRQFSKGAWPFTDQDHGWPVSDCTAEAVKCLLLLSQMPEEISGEKADVE<br>RLYDAITFLLHVQSPISGGFAIWERPIPQPYLEKLNPSEVFADIVLEKEHMENTVSIIQAFVAFKRLHPTHRA<br>KELENSLAKAVRFVEKRQLPNGSWYGYWGICYIYGTLFALGGLEAVGKTYDNSESVRKGVKFLLSTQNEEGGW<br>GESYKSCLTEVFTPLVENRTNLVQTAWAMLGLMSGGQAERDPTPLHKAAKLLINAQMDNGDFPQQELTGASLR<br>NCMLLYPLYRNSFTLLALAEYRKRLWAIN |
| >CoOSC7(partial) | SRLRKENPINEIPEAIKLNETEELTNEAVTTTLRRAISFYSTIQAHDGHWPAESAGPLFFLPPLVIALYVTGT<br>MNAILTLAHQVEIKRYLYNHQNEDGGWGLHIEGDSTMFGSVLSYITLRLLGEEAEDMAVVKGRKWILDHGGAI<br>GTPSWGKFWLTVLGVYEWEGCNPMPPEFWLLPKIFPIHPGKMLCYCRLVYMPMSYLYGKRFAGKITGLVQELR<br>QELYTAPYHEINWNKGRNTCAKEDLYYPHPLVQDMLWGVLHNIAEPILTRWPFSKLREKALKVAMEHVHYEDQ<br>SSRYLCIGCVEKVLCLIATWVEDPNGDAFKRHLARIPDYFWVAEDGMKMQSFGCQMWDASFAIQAIMSSNLAE<br>EYGPTLKKAHEFVKSSQVRDNPPGDFGKMYRHMSKGAWTFSMQDHGWQVSDCTAEGLKVALLYSQMSPELVGE<br>KLENEHLYDAVNIILSLQSKNGGFPAWEPQRAYSWLEKFNPTEFFEDVMIEREYVECTSSAVQSLELFKKLHP<br>RHRTKEIEHCISRALKYIEDMQNPDGSWYGCWGICYTYGTWFAVDALVACGKNYHNCPALRKACQFLISKQLP<br>DGGWGESYLSSANKEYTNLDGDRSNLVQTSWALISLIKAGQAGINVTPIDRGIRLLINSQTEDGDFPQQEITG<br>VFMKNCTLNYSSFRNIFPIWALGEYRRIV |
| >CoOSC8 | MWRLKIAEGVNDPLLHSTNNFVGRQTWEFDPNYGTPEEIDEVEKARLHYWNHRHQVEPCSDVLWRMQFLREKK<br>FKQTIPQVKIEDGEEISYEKVTATLRRSVHLFAALQAEDGHWPAENAGPMYFMQPLVICLYITGHLNGVFPAE<br>HRKEILRYLYCHQNEDGGWGFYMEGQSTMFGTTLSYICMRLLGEGPDNGACTKARKWILDHGSATVIPSWGKT<br>WLSVLGVREWVGSNPMPPEFWILPSFLPMHPGKMWCYCRLVYMPMSYLYGKRFVGPITPLVLQLREELYSQPY<br>DEINWKSTRHVCAKEDLYYPSPFLQDLLWDSLYIFTEPLLTRWPFNLLRKKALKTTMKHIHYEDENSRYITIG<br>SVIKSLCMLACWDEDPNGMCFKKHLARIPDYIWIAEDGMKMQSFGSQSWDASLSIQALLATDLSNEIGPMLKK<br>GHDFIKASQVKDNPSGLFKSMYRHTSKGSWTFSDQDHGWQLSDCTAEGLRCCLLFSTMPPEIVGKKMQPEQFF |

| | |
|---|---|
| | DAVNIILSLQSKNGGLSGWEPARSSKWLEILNPTEFFEDIVIEHEYVECTSSGIQTLVLFNKLYPEYKTKEIQ<br>RFLTNAGGYLEKTQTPDGSWYGEWGMCFTYATWFALGGLEAIGKTYENCEAIRKAVNFLLKIQREDGGWGESH<br>RSSPEKKYIPLEGNRQSNLVNTAWAMMGLIHSGQAERDPTPLHRGAKLLINSQMENGDFPQQETTGVFKKNCL<br>LHFSLFRDIFPMWALASYRKSVLPQPTSI |
| >CoOSC 9 | MWRLRLGEGADDPYLFSTNNFVGRQTWEFDPNYGTLEERGEVEQARTDFWNNRHKVKPCNDAIWRMQFLREKK<br>FKQTIPQVKIEDGEEISYEKATTTLKRSVNYMAALQSDHGHWPAEIAGPQYFMQPLVFCLYISGHLNTVFPAE<br>HQKEILRSLYTHQNEDGGWGLHIEGHSTMFCTTLSYICMRMLGEGPDGGLNGACTRARKWILDHGSVTHIPSW<br>GKTWLSILGLCEWAGTNPMPPEFWILPSFLPMHPAKMWCYCRLVYMPMSYLYGKRFVGPITPLILQLRDELYL<br>QPYNEIKWGSIRHLCAKEDLYYPHPLLQDLIWDSLYVLTEPLLTRWPFNKLREKALQTTMKHIHYEDENSRYI<br>TIGCVEKALCMLACWVEDPNGDAYKKHLARVPDYLWIAEDGMKMQSFGSQEWDAGFAIQALLAADFTDEIAST<br>LKKGHEFIKASQVKDNPSGDFKSMHRHISKGSWTFSDQDHGWQVSDCTAEGLKCCLLFSTMPPEIVGEQMPAE<br>QLNNAVNVILSLQSKNGGLAAWEPAGAAEWLEILNPTEFFADIVIEHEYVECTGSAMHALVLFKKLYPGHRRK<br>EIENFLINATKYLENIQMPDGSWYGNWGVCFTYGTWFALGGLAAVGKTYENCAAIRKAVNFLLETQLKDGGWG<br>ESYKSCPEKRYIPLEGGRSNLVHTAWALMGLIHSRQMDRDATPLHRAAKLLINSQLETGDFPQQEITGVFMKN<br>CMLHYPMYKNIYPMWALAEYRKHVLPLLKPSIK |
| >CoOSC 10 | MWRLKLGEGADDPYLFSTNNFVGRQTWEFDPNYGTPEERAEVEQARTDFWNNRHKVKPCNDAIWRMQFLREKK<br>FKQTIPQVKIEDGEEISYEKATTTLKRSVNYMAALQSDHGHWPAEIAGPQYFMQPLVFCLYISGHLNTVFPAE<br>HQKEILRSLYTHQNEDGGWGLHIEGHSTMFCTTLSYICMRMLGEGPDGGLNGACTKARKWILDHGSVTHIPSW<br>GKTWLSILGLCEWAGTNPMPPEFWILPSFLPMYPAKMWCYCRLVYMPMSYLYGKRFVGPITPLILQLRDELYL<br>QPYNEIKWGSIRHLCAKEDLYYPHPLLQDLIWDSLYVLTEPLLTRWPFNKLREKALQTTMKHIHYEDENSRYI<br>TIGCVEKALCMLACWVEDPNGDAYKKHLARVPDYLWIAEDGMKMQSFGSQEWDAGFAIQALLAADFTDEIAST<br>LKKGHEFIKASQVKDNPSGDFKSMHRHISKGSWTFSDQDHGWQVSDCTAEGLKCCLLFSTMPPEIVGEQMPAE<br>QLNNAVNVILSLQSKNGGLAAWEPAGAAEWLEILNPTEFFADIVIEHEYVECTGSAMHALVLFKKLYPGHRRK<br>EIENFLINATKYLESIQMPDGSWYGNWGVCFTYGTWFALGGLAAVGKTYENCAAIRKAVNFLLGTQLKDGGWG<br>ESYKSCPEKRYIPLEGGRSNLVHTAWALMGLIHSRQMERDATPLHRAAKLLINSQLETGDFPQQEITGVFMKN<br>CMLHYPMYKNIYPMWALAEYRKHVLPLLKPSIK |
| >CoOSC 11 | MWRLKIAKGGDNPYLYSTNNFVGRQIWEFDPNYGTPEEHADVENARVGFWNNRHDIKTSGDVLWRMQFLRENE<br>FKQTIPQVKIEDGEKISNEKATTTLKRSVNFFSALQADDGHWPAENAGPLYFMQPLVMCLYITGHLNTVFPTE<br>YRKEILRYIYCHQNEDGGWGLHIEGHSTMFCTTLSYICMRMLGEGPDGGLNGACTKARKWILDHGSVTHIPSW<br>GKTWLSILGLCEWAGTNPMPPEFWILPSFLPMYPAKMWCYCRLVYMPMSYLYGKRFVGPITPLILQLRDELYL<br>QPYDEINWKSVRHLCAKEDLYYPHPLLQDLMWDSLYVFTEPFLNHWPFNKLREKALQTTMKHIHYEDENSRYI<br>TIGSVEKALCMLACWVEDPNGVCFKKHLARIPDYLWVAEDGMKMQSFGSQQWDAGFAIQALLATDLTDKIGST<br>LMKGHEFIKVSQVKDNPSGDFKSMHRHISKGSWTFSDQDHGWQVSDCTAEGLKCCLLFSTMPPEIVGEKMKPE<br>QLNDAVNIILSLQSKNGGLAAWEPAGSSEWLEVLNPTEFFADIVIEHEYVECTSSAMQALVLFKELFPGHRRK<br>EIENFLTAASGYLENIQMPDGSWYGNWGVCFTYGTWFALGGLTAVGKTYENCAAIRKAVKFLLETQLEDGGWG<br>ESYKSCPEKRYIPLEGGKSNLVHTSWALMGLIHSRQMERDATPLHRAAKLLINSQLENGDFPQQEIAGVFMKN<br>CMLHYPLYRNIYPMWALADYSKQVLPSNS |
| >CoOSC 12 | MWKLKIAEGGSPWLRSNNDHVGRQIWEFDPTLGSLEELADIEKVRQAFHDNRFEKKHSSDLLMRSQFAKVKPQ<br>SVFPPKVNIKDAEDITENKVTDVLRRAISFHSTLQADDGHWPGDYGGPMFLLPGLVITLTITGALNAVLSKEH<br>KREMVRYLYNHQNIDGGWGLHIEGHSTMFGSALNYVTLRLLGEGANDGEGAMEKGRKWILDHGGATSITSWGK<br>FWLSVLGVFEWSGNNPLPPEMWILPYFLPIHPGRMWCHCRMVYLPMSYLYGKRFVGPITSTVLALRKELFTVP<br>YHDIDWNVARNLCAKEDLYYPHPLVQDILWATLDKFVEPVLMRWPGKKLRAKALRTAMEHIHYEDENTRYICI<br>GPVNKVLNMLCCWAEDPNSEAFKLHLPRIHDYLWIAEDGMKMQGYNGCQLWDTAFAVQAIISANLIEEFGSTL<br>KKGHMYIKNSQVLDNCTGDLDYWYRHISKGAWPFSTADHGWPISDCTAEGLKAALLLSKLPTEIVDEPVDENR<br>LYDSVNVILSLQNGDGGFATYELTRSYRWLELVNPAETFGDIVIDYPYVECTSAAIQALVAFKKLYPGHRRKE<br>VERCIDKAAAFIERIQESDGSWYGSWAVCFTYGTWFGIKGLVAAGKTYSNCSSIRKACNFLLSKQLASGGWGE<br>SYLSCQDKVYTNLEGNRSHVVNTGWAMLSLIDAEQAKRDPTPLHRAARVLINSQMENGDFPQQEIMGVFNRNC<br>MITYAAYRNIFPIWALGEYRCRVLT |
| >CoOSC 13 | MWKLKIAEGGSPWLRSTNDHVGRQIWEFDPTLGSLEELADIEKVRQSFHDNRFEKKHSSDLIMRSQFAKVKSQ<br>SVFPPKVNIKDAEDITEDKVTDVLRRALGFYSTLQADDGHWPGDYGGPMFLLPGLVITLTITGALNAVLSKEH<br>KREMVRYLYNHQNIDGGWGLHIEGHSTMFGSALNYVTLRLLGEGANDGQGAMEKGRQWILDHGGATFITSWGK<br>FWLSVLGVFDWSGNNPLPPEMWLFPYFLPIHPGRMWCHCRMVYLPMSYLYGKRFVGPITSTILALRKELFTVP<br>YHDIDWNEARNLCAKEDLYYPHPLVQDILWATLDKFVEPVLMRWPGKKLREKALCTAMEHIHYEDENTRYICI<br>GPVNKVLNMLCCWVEDPNSEAFKLHLPRIQDYLWLAEDGMKMQGYNGSQLWDTAFAVQAVISTNLIEEFGPTL<br>KKGHMYIKNSQVLENCTGDLDYWYRHISKGAWPFSTADHGWPISDCTAEGLKAALLLSKLPKQIVDEPVDENR<br>LYDSVNVILSLQNGNGSYATYELTRSYSWLELVNPAETFGDIVIDYPYVECTSAAIQALVAFKKLYPGHRREE<br>VQRCIDKAAAFIETIQESDGSWYGSWAVCFTYGTWFGIKGLVAAGKSYSNCSSIRKACDFLLSKQLASGGWGE<br>SYLSCQDKVYTNLEGNRSHVVNTGWAMLALIDAEQAKRDPRPLHLAARVLINSQMENGDFPQQEIMGVFNRNC<br>MITYAAYRNIFPIWALGEYRCRVLQQPPC |
| >CoOSC 14 | MWKLKIAEGGSPWLRSNNNHVGRQIWEFDPTLGSIEELADIEKIRQTFHDNRFEKKHSSDLLMRSQFAKVKSQ<br>SVFPPKVNIKDAEDITEDKVTDVLRRAVSFHSTLQADDGHWPGDYGGPMFLLPGLVITLTITGALNAVLSKEH |

| | |
|---|---|
| | KREMVRYLYNHQNIDGGWGLHIEGHSTMFGSALNYVTLRLLGEGANDGSGAMEKGRKWILDHGGATSITSWGK<br>FWLSVLGVFEWSGNNPLPPEMWILPYFLPIHPGRMWCHCRMVYLPMSYLYGKRFVGPITATVLALRKELFTVP<br>YHDIDWNVARNLCAKEDLYYPHPFVQDIIWATLDKFVEPVLMRWPGKKLRAKALQTAMEHIHYEDENTRYICI<br>GPVNKVLNMLCCWAEDPNSEAFKLHLPRIQDYLWIAEDGMKMQGYNGSQLWDTAFAVQAIISTNLIEEFGPTL<br>KKGHMYIKNSQVLDNCTGDLDYWYRHISKGAWPFSTADHGWPISDCTAEGLKAALLLSKLPREIVDEPVDENR<br>LYDSVNVILSLQNGDGGFATYELTRSYHWLELVNPAETFGDIVIDYPYVECTSAAIQALVAFKKLYPEHRRDE<br>VEHCIDKAAAFLERIQESDGSWYGSWAVCFTYGTWFGIKGLVAAGKTYSNCCSVRKACDFLLSKQLASGGWGE<br>SYLSCQNKVYTNLEGNRSHVVNTGWAMLALIDAEQAKRDPTPLHRAARVLINSQMENGDFPQQEIMGVFNRNC<br>MITYAAYRNIFPIWALGEYRCRVLT |
| >CoOSC<br>15 | MWKLNIAEGGSPWLRSTNDHVGRQIWEFDPTLGSLEELADIEKVRQTFHDNRFEKKHSSDLIMRSQFAKAKSQ<br>SVFPPKVNIKDAEDITEDKVTDVLRRAVGFYSTLQADDGHWPGDYGGPMFLLPGLVITLTITGALNAVLSREH<br>KREMVRYLYNHQNIDGGWGLHIEGHSTMFGSALNYVTLRLLGEGANDGQGAMEKGRKWILDHGGATFITSWGK<br>FWLSVLGVFDWSGNNPMPPEMWLFPYFLPIHPGRMWCHCRMVYLPMSYLYGKRFVGPITSTILALRKELFTVP<br>YHDIDWNEARNLCAKEDLYYPHPLVQDILWATLDKFVEPILMRWPAKKLREKALCTAMEHIHYEDENTRYICI<br>GPVNKVLNMLCCWVEDPNSEAFKLHLPRIQDYLWLAEDGMKMQGYNGSQLWDTAFAVQAIISTNLIEEFGPTL<br>KKGHMYIKHSQVLDNCTGDLDYWYRHISKGAWPFSTADHGWPISDCTAEGLKAALLLSKFPTEVVDEAVDENR<br>LYDSVNVILSLQNGDGSYATYELTRSYSWLELVNPAETFGDIVIDYPYVECTSAAIQALVAFKKLYPGHRREE<br>VERCIDKAAAFIETIQESDGSWYGSWAVCFTYGTWFGIKGLVAAGKSYSNCSSIRKACNFLLSKQLASGGWGE<br>SYLSCQDKVYTNLEGNRSHVVNTGWAMLALIDAQQANRDPRPLHLAARVLINSQMENGDFPQQEIMGVFNRNC<br>MITYAAYRNIFPIWALGEYRCRVLQPPC |
| >CoOSC<br>16 | MWKLKIAEGGSPWLRSNNNHVGREIWGFDPTLGSLEELADIEKARQTFHDNRFENKHSSDLLMRSQFAKVKSQ<br>YVFPPKVTIKDAEDITEDKVTDVLRRAIGFHSTLQADDGHWPGDYGGPMFLLPGLVITLTITGALNAVLSKEH<br>KREMVRYLYNHQNIDGGWGLHIEGHSTMYSSVINYVTLRLLGEGANDGEGAMEKGRKWILDNGGATSITSFGK<br>FWLSVLGVFEWSGNNPLPPEMWILPYFLPIHPGRMWCHCRMVFLPMSYLYGKRFVGPITTQVLALRKELFTVP<br>YHDIDWNVARNLYAKEDLCYPNPLVEYILWPTLNKFVEPMLMQWPGKKLREKALLTAMEHIHYDDENTQYLCI<br>GPVNKALNMLCCWAEDPNSEAFKLHLPRIHDYLWIAEDGMKMKGYNGTQLWDTAFAIQAIISTNLIEEFGPTL<br>RKGHMYIKNSQVLDNSTGNLDYWYRHISKGAWTFSTADQGWPISDGTAEGLKSALLLSKLPTEIVDEPLDENR<br>LYDSVNVILSLQNGDGGFASYELTRSYSWLEFVNPAKTFGDMVIDYTYVECTSASIQALVTFKKLYPGHRREE<br>VQHCIDKAAAFIERMQESDGSWYGSWGVCFTYGTWFGVKGLVAAGKTYTNCSSIRKACNFLLSKQLASGGWGE<br>SHLSCQKKVYMNLEGNRSHVVNTGWAMLALIDAEQAKRDPTPLHRAARELTNSQMENGDFPQQEIIGVFNRNC<br>MITYAAYRNIFPIWALGEYRCRVLQMLSSI |
| >CoOSC<br>17 | MWKLKIGEKNGKLEIGDGNGDEYLYSTNKFVGRQTWEFDPSQEEREQIERIREQFLNNKKKLDIHCCGDLLMR<br>NQIQAWDGHWPAENAGPLFFTPPLVIDSRCGTLDTILTKEHQKVRYMYIHQNEDGGWGFGMIGSALNYVALRL<br>VGEASPDDDALARGHKWIFDHTSIPSWGKLYLAVYMVYQWEGCNPLPPEFWLFPSDLVLSLRKEIHTLPYHQI<br>NWNKQRHNCCKVDLYYPHSSIQDLLHYFCEPLVRKWPFNKLREKGVDLMRYGSEEGRYIGMGCVDKALQMMCF<br>YPNGNDFKRHLARVPDYLWVAEDQSFGSQLWDCTLVTQAIMATNMVDEYGDSLKKAHFYLKQSQIKENPKGDF<br>TKMCRLFTKGAWTFSDQDHGWVVSDCTAEALICLLALSQMPQDIAGEKAEVDRLYDAVNVLLYLQSPESGGFA<br>IWEPPVPKPYLQMLNPSELFADIVVEKEHVECTGSIIQALNSFKNLHPGHREKEIEAAIEKGIRFLENKQQNN<br>GSWYGYWGICFLYGTFFVLQGLVSCGKTYENSETVRKAVNFLLSTQNSEGGWGESFESCPQEKFIPLEENRTN<br>LVQTSWAMLGLLYGGQVERDVTPLHKAAKLLINGQLDNGDFPQQEITGVYMKNCMLHYPEYRNTFPLGEYRKL<br>VWLPKQQV |
| >CoOSC<br>18 | MWSLKIAEGNDHPYLYSTNSFVGRQIWEFDPDAGTPQEKQQVNARQHFRNNLSKGVHPCSDLLMRMQLMKENG<br>IDLLSIAPSRLGENEEVNYEAVTRAVKKAIRMNRAIQAKDGPENAGPMFFTPPLLIAMYISGNINTHLTKEHR<br>TEMIRYIYNYQNEDGGWGFYITADRVCFELCRLLGEGPNDGNGAVERGRKWILDHGGATSIPSWGKTYLSVLG<br>VYEWEGCNPLPPEFWLFPQALPYHPKMWCYCRTTYMPMSYLYGRKFHGPITDLVLQIRKEIHPIPYNEINWNK<br>QRHNCCKEDLYYPHSTVQDLLWDGLHYLSEPIFKFWPFTKLRERGLKRAVELMYGAQQSRYITIGCVEKSLQM<br>MCWWAENSNGVEFKRHLARVPDYLWLAEDGMKMQSFGSQVWDCALATQAIIASNMVEEYGDSLKKANFYIKKS<br>QIKYNTCGDFSKMCRQFTKGSWTFSDQDHGWVVSDCTAEALKCLLLLSQMPEEISGDKADNERLYEVVNVLLY<br>LQSPTSGGFAIWEPPVPHPYMQMLNPSELFADIVVEKEHVECTASIIQAILAFKRLHPQHREKEIEISVAKSV<br>TFLEGKQRDDGSWYGYWGICFLYGTFFAIGLEAAGKTYNNSEAVCEAVNFFLLTQNEEGGWGESFKSCPSDLY<br>TPLDGNRTNLVQTSWAMLGLMLGGQIERDPTPMHKAAKILVNAQTDNGDFPQQKITGVYMKNCTLHYPEYRNI<br>FPLWALGEYRKRVWVN |
| >CoOSC<br>19 | MWKLKIAEGNDPYLFSTNNFVGRQTWEFDPNAGTTEEKQEIENARQYFLNRQKEGYQASSDLLTRIQLTKENG<br>IDLLSIRQARLRNDEEVNYEAVTTAVKKAVRFQRAIQAKDGHWPAEQAGPLFLTPPLIIVLYISGAINTHLTK<br>EHKNEMKRFIYNHQNEDGGWVFHIEGHSTMMGSVLNYISLRLLGEGPDDKNGAVARGRKWILDHGGATSIPSC<br>GKTYLSVLGLYEWEGCNPMPPEFWIFPEVFPFHPAKMWCYCRTAYMPMSYLYGRRYHGPITDLVLHLRREIYR<br>DPYDKINWNKQRHNCCKEDLIYPHSTLQDLLWDGLHYFCEPFIKYWPFTKLRERALKRAIDLIRYSAQESRYI<br>NMACVEKCLQMMCWWAESPNGDEFKHHLARMPDFLWLGEDGMKMQTFGGQLWDCTLATQAIISSNMPEEYGDS<br>LRKANFYIKETQITSNPSGDYSKMYRQLSKGAWGFTDKDNGWAVSDCTSEALKCLLLLSEMPEEISGEKADNE<br>RLYDAITFLFHVQSPTTGGFAIWEKPIPQPYLENLNPSEVFADIVLEKEHLENTASIIQALVAFRRLHPSHRP |

| | |
|---|---|
| | IEIENSVAKAVHFVEKRQLPNGSWYGYWGICFIYGTFFALEGLEAAGKTYNNSETIRKGVKFLLSTQNEEGGW<br>GESYKSCLSEVFTPLIENRTNLVQTAWAMLGLMSGGQAERDPTPLHKAAKLLINAQMDNGDFPQQEFTGASLR<br>NCMLHYPLYRNSFTLRALAEYRKRLWTIN |
| >CoOSC 20 | MWKLKIAEGKDPYLFSTNNFVGRQTWEFDPDAGTPEEKQDVENARQFFLGRQKEGFQACSDSLMRMQLTKENG<br>VDLLSIPPARLRDEEQVNYVAVTTAVKKAVRFQRAIQAKDGHWPAGHTGPLFFTPPLIIVLYISGTINTQLTE<br>EHKKELKRYIYNHQNEDGGWGFHIHGHSTMLGSVLNYIALRLLGEGPDDGNGAVDRARKWILDHGGATAIPSW<br>GKLYLSVLGVYEWEGCNPMPPEFWIFPDAFPFHPKMWCYCRTAYMPMSYLYGRKYHGPITDLILHLRQEIYSI<br>PYDKIIWNKHRHNCCKEDLFYLHSTIQDMLWDGLHYFCEPFFKFWPFTKLRERALKRAVELIRYSAHETRYIT<br>MASIEKSLQMMCWWAENPNGDEFKHHLARVPDFLWLGEDGMVMQTFGSQLWDCTFATQAIISSNMPQEFGDSL<br>KKANFYIKESQIKNNPSGDFSKMCRQFSKGAWTFSDQDQGWPVSDCTAEALKCLLLLSQLPEEITGKKIDKEQ<br>LYEAINFLFHVQSPTSGGFACWERPIPQPYLEKLNPSEVFADIVLEKEHLEITVSIIQALIAFKCLHPTHRAK<br>EIDISVANGVRYLENKQFTNGSWYGYWGICFIYGTFFALGGLEAAGKTYNNSESVRKGVQFLLSTQTKGGGWG<br>ESYKSCNTEVYTPLTDNRTTVVQTAWAMLGLMSGGQAERDPTPLHKAAKLLINAQMDNGDFPQEEFTGASMRN<br>CMLHYPLYRNSFTLRALAEYRNRIW |
| >CoOSC 21 | MWRLKIGEGGNDPHLYSTNNFVGRQTWEFDPNYGTPEEIDEVEEARLHFWNNRHQVKPSSDLLWRMQYLREKK<br>FKQTIPQVKIEDGEEISYEKVTATLRRSVHLFAALQAEDGHWPAENAGPMYFIQPLVISLYITGHLNSVFPSE<br>HRKEILRYLYCHQNEDGGWGLHMEGHSIMFGTTLSYITMRLLGEGADNGACTKARKWILDHGSVTTIPTWGEI<br>WLSIFGVYEWIGCNPIPPEFWILPSFLPMHPGKMWCYCRLVYMPMSYLYGKRFVGPITPLVLQLRDELYTQSY<br>NEINWKSIRNLCCKEDLYYPHPVIQDLIWDGLYYFTEPLLCRWPFNKLREKALETAMKHIHYEEENTRYNTIG<br>CVEKVLCMLASWVEDPNGVHFKKHLARIPDYIWVAEDGMKMQSCAGSQEWDTGLAIQALLATNLTDEIGPTLK<br>KGHDFIKASQVKDNPSGDFKSMYRAMSKGAWTFASQDHGWQVTDCTADGLKCCLLLSMMPPEIVGKKMEAEKL<br>YDAVNILLSMQSKNGGMSAWEPAGSSKWLEMLNPTELFADIVIEHEYVECTSSVMQSLVLFKKMYPQHRRQEI<br>GSLLTRATTYLEATQMPDGSWYGEWAVCFIYGTSFALEGLAPIGKTFENCLAIRKDVNFLLNTKEDGGWGESY<br>LSCSDKKYVALEGGRSNLVQTAWAMMGLIHSKQAERDSTPLHRAAKLLINSQTKHGHFPQQEVSGAFKKTSLL<br>HYPCYRNHFPIWALAEYRNQVLPKPTCI |
| >CoOSC 22 | MWRLKIGEGGNNPHLYSTNNFVGRQTWEFDPNYGTPEEIDEVEEARLHFWNNRHQIKPSSDLLWRMQFLREKE<br>FKQSIPQVKIEDGEEISYEKVTATLRRSVHLFAALQAEDGHWPAENAGPMYFIQPLVICLYITRHLNSVFPSE<br>HRKEILRYLYCHQNEDGGWGLHMEGHSIMFGTTLSYICMRLLGEGPDNGACTKARKWILDHGSVTTIPTWGKI<br>WLSIFGVYEWIGCNPIPPEFWILPSFLPMHPKMWCYCRLVYMPMSYLYGKRFVGPITPLVLQLREELYTQPYS<br>EIKWKSIRNLCCKEDLYYPHPFIQDLIWDGLYYLTEPLLCRWPFNKLRQKALETAMKHIHYEEENTRYITIGC<br>VEKVLCMLASWVEDPNGVHFKKHLARIPDYIWVAEDGMKMQSCTGSQDWDAGLAIQALLATNLTDEIGPTLKK<br>GHDFIKASQVKDNPSGDFKSMYRPISKGAWTFPSQDHGWQVSDCTAYGLKCCLLFSMMPPEIVGKKMEAEKLY<br>DAVNVLLSMQSKNGGMSAWEPAGTSKWLEMLNPTELFADIVIEHEYIECTSSVMQSLVLFKKMYPQHRRQEIE<br>SLLTRATTYLETMQMPDGSWYGEWGVCFIYSTSFALEGLAVVGKTFENCLAIRKGVNFLLNTHMEDGGWGESY<br>LSCSEKKYVALEGGRSNLVQTAWAMLGLIQSKQAERDPTPLHRAAKLLINSQTKNGDFPQQEVSGAFKKTSLL<br>HYACYRDYFPIWALAQYRNQLLPQLTTN |
| >CoOSC 23 | MMKENGIDLLSIPPVRLGENEQVNYEAVTISVTKAIRLNRAIQAKDGHWPAEAMFFTPPLLIAMYISGAIDTH<br>LTKEHKTEMIRYIYNHQIRQEIHVIPYNEINWNKQRHNCCKEDLYYPHSTVQDLLWDGLQYLSEPILKYWPFT<br>KLRERGLKRAVELMRYGAQESRYMTIGCVEKSLQMMCWWAENPNGDEFKHHLARVPDYLWLAEDGMKMQSFGS<br>QLWDCVLATQAIIATDMVEEYGDSLKKAHFYIRESQVKQNPTGDFSKMCRQFTKGSWTFSDQDQGWVVSDCTA<br>EALKCLLLLSQMPEEISGEEADNERLYEAVNVLLYLQSPITGGFAIWEPPVPQPYLQMLNPSELFADIVVEKE<br>HVECTASIIQALLAFKRLHPGHREKEIEISLAKAVTFVEGNQRHDGSWYGYWGICFLYGTFFAVGGLVSAWKT<br>YNNSESIRKAVNFFILTQNEQGGWGESIKSCPTEVYTPLDGNRTNLVQTSWAMLGLMLGGQVERDPTPLHKAA<br>KILINVQMDNGDFPQQEITGVYNKNCMLHYPEYRNIFPLWALGEYRKRVWIN |
| >CoOSC 24 | MWELKIAEGDGPYLYSTNNFVGRQFWEFNPDAGTLEEKQEIEMARQKYKNNRRNGGFHACGDLLMRRQMIKEN<br>GIDLTSIAPMRVKEHEHVNFEAVTTAVRKAVRLQRAIQAKDGPMFFTPPLVIALYISGMINIILSEEHKKEMI<br>RFFYNHQNEDGGTMIGSALSYILGEAEDGAISRGRKWILDHGGATSIPSWGKVYLSVLGVYEWDPLPPEFWLF<br>PSAFPFHPEMWCYCYMPMSYLYGKRIQGPITNHPTPFEDISWNKQRNNCCKEDFYYPHSFLQDALWHSLTEPV<br>LKHWPFSKLRNRAVDRVVELMRYETRYMTIEKSLQMMCWWAENPNGDEFKYHLARVPDYLWIAEDGMTMHSFG<br>SQVWDCSLAIQAIVASNMADEYGDCLKKAHFYLRESQVKENPSGDFTRMCRQFTKGSWMDCLQLFCPIVLLSN<br>MPKEIAGNKDNTARLYDAVNSGGFAAPIPKPFLQLLNPSEIFADIVVEKEMIVETTSSIIGVLMEFNRTHPER<br>KKLISKGIRYLEETWVDRYGYWGVCFIYGTFRGLTCVGKTYDNNEAVCKFLLSIQNEEGGWGESLLSCPTEVY<br>TPLDGNRTNLVQTSWAMLGLMFTVQRDPMPLHKAAKLLINAQMDNGDFPQQEITGLYMKNCLLLYAQYRNIFP<br>LWALGEEYRKRVW |
| >CoOSC 25 | LRKEKQIDKIPEANETEELTNEAVATTVRRAISFYSTIQAHDGAESAGPLFFLPPLDSTGTMNVILTPAHQVE<br>IKRYLYNHQNKDGGWGLHIEGHSTMFGSVFSYITLRLLGEEADSGGGGCKWILDHGGAIGTPSWGKFWLTVLG<br>VYEWEGCNPMPPEFWLLPKIFPIHPKMLCYCRLVYMYLYGKRFVGRITGLVQALRQELYTGPYHEINWNKGRN<br>TCAKEDLYYPHPLVQDMLWGVLHNIGEPILTRWPFSKLREKALKVAMEHVHYEDQSSRYLCIGCVEKVLCLIA<br>TWVEDPKGNAFKRHLARIPDYFWVAEDGMKMQSFGCQMWDAAFAIQASNLTEEYGPTLKKAHEFVKASQVFSG<br>SPRGDFGKMYRHMSKGAWTFSMQDHGWQVSDCTAEGLKVALLYSQMSPELVGEKLENEHLYDAVNVILSLQSK<br>NGGFPGWEPQRAYSWLEKFNPTEFFEDVMIEREYVECTSSAIQSLALFKKLHPGHRTMKIEHCISKAVKYIVD |

| | |
|---|---|
| | MQNTDGSWYGCWGICYTYGTWFAVDALVACGKNYHNSPSLRKACQFLLSKQLPDGGWGESYLSSANKEYTNLD GDRSNLVQTSWALLSLIKAGQAGVNLTPIDRLVINSQTEDGDFPQQEITGVFMKNCTLNYSSFRNIFPIWALG EYRRIRMKL |
| >CoOSC 26 | MWKLKLSKGDDDPGVRSVNNHIGRQYWEFDPLAGTPEEQSQINNMREEFTRNKVKVKHSSDLLMRFQFASEIN GSEMKKSQVIEAKDEDGEEVVVKTLKKGLKFYSSLQGEDGSWPADYGGPLFLLPGLIIALHMMGAKDRALSVE HQREIRRYLYNHQNVDGGWGLHIEGHSTMFCTALNYVSLRLLGERMDGGEGAMTKARKWILDHGSVTHIPSWG KFWLSVLGVYEWSGNNPLPPEMWLLPYFVPLHPGRMWCHTRMVYLPMSYIYGKRFVGPINSIVLSLRRELYNT HYYQVNLIITEILTLRISKYSILLIYLMINDILTIILLKYYLSILKKKKKRKELWPKNEARGTNPPNHKVTCG TMREKVLNMLCCWVEDPMSHVNKLHLSRIEDYLWIAEDGMKMQGYNGSQLWDSVFAVQAILATNLVDEYGSML RKAHSFIKNSQIRENSSGNIQSWYRHIIRGGWPFSTPDNGWPVSDCTAEALKTVLMLSQMPNDIVGEAIAPEC LYDAGHLLLTLQNGNGGFSSYEPMRSYPWLEVFNPAETFGDIIVDYYVECTSAVVQGLSSFMKLYPSHRRDEI EACVDKAISFIENVQLHDGSWYGSWGICYTYGTWFGIKGLVATGETYETNYSIRKACAFLLSKQLPSGGWGES YISCEQKNYTNIAGNKSHIANTSWALLALIEAGQAKRDKMPLHRAAKVLIDHQMENGDFPQQEIIGVFNKNCM ISYSSYRNIFPIWALGEYLNRVIRNNLPS |
| >CoOSC 27 | LKLSKGDDDPSVRSVNNHIGRQYWEFDPCAGTPEERSQIDNMREEFTINKAKVKHSSDLFMRFQFGRENCSEI MEKSQVTEDGDEVVVKTLKKALRFYSTLQGEDGSWPADYGGPLFLLPGLIIALHMMGAMDIALSVEHQREIRR YLYNHQNVDGGWGLHIEGQSTMFCTALNYVSLRLLGERMDGGQGAMAKARKWILDHGSVTHIPSWGKFWLSVL GVYEWSGNNPLPPEMWLLPYFVPLHPRMWCHTRMVYLPMSYLYGKRFVGPINSIVLSLRRELYNTLYNQVNWD LARNQCAKEDLYYPHPMIQDILWGGLNKIAEPLLMQWPFSKLRKKALTTVMQHVHYEDKNTQYICIGPVNKVL NMLCCWVEDPMSHVNKLHLSRIKDYLWIAEDGMKMQGYNGSQLWDSVFAVQAILATNLVDEYGSMLRKAHSFI RNSQIRENSSGNIQSWYRHIIRGGWPFSTPDNGWPVSDCTAEALKTVLMLSQMPKDIVGEAVAPECLYDAVHL LLTLQQNGNGGFSSYEPMRSYPWLEVFNPAETFGDIIVDYYVECTSAVVQGLSSFMKLYPSHCRDEIEACVDK AITFIQNVQLHDGSWYGSWGICYTYGTWFGIKGLVAAGATYETNHNIRKACAFLLSKQLPSGGWGESYVSCEQ KNYTNIAWNKSHITNTSWALLALIEAGQAKRDKVPLHRAAKVLIDHQMENGDFPQQEIIGVFNKNCMISYSSY RNIFPIWALGEYLNRVWVVLPKCNV |
| >CoOSC 28 | MWSLKIAEGNGHPYLFSTNNFVGRQIWEFDPDAGTPEEKQQVENARQHFTNNRSKGVHPCSDLLMRMQLIKEN GMDLLNIPPSRLGENEQVNYEAVTTAVKKAIRLNRAIQAKDGHWPAENAGPMFFTPPLLIAMYISGAINTHLT KEHKSEMIRYIYNHQNEDGGWGFYIGGDSTMIGSALSYVFLRLLGEGPDDGNGAVGQARRWILDHGGATIPSW GKTYLSVLGVYEWEGCNPLPPEFWLFPRALPYHPKMWCYCRTTYMPMSYLYGRKFHGPLTELVLQIRQEIHSI PYNEINWNKQRHNCCKEDLYYPHSTVQDLLWDGLHYLSEPILKFWPFTKLRERGLKRAVELMRYGAQESRYIT TGYISKSLQIMCWWAENPNGVEFKRHLARVPDYLWLAEDGMKMQSFGSQLWDCVLATQAIIASDMVEDYGDSL KKANFYIKKSQIKTNPCGDFSKMCRQFTKGSWTFSDQDQGWVVSDCTAEALKCLLLLSQMPEEISGAKADNER LYEAVNVLLYLQSPTSGGFAIWEPPVPQPYLQMLNPSELFADIVVEKEHVECTASIIKALLAFKRLHPQHREK EIEISVTKAVTFLEGKQQGDGSWNGYWGICFLYGTFFAIGGLEAAGKTYKNSETIRKAVNFFHLTQNEEGGWG ESIKSCPSEVYTPLDENRTNLVQTSWAMLGLMLGGQIERDPTPLHKAAKILINAQMDNGDFPQQEITGVYMKN CMLHYAEYRNIFPLWALGEYRKRVWVN |

**Supplementary Table 9: Table of oxidosqualene cyclases (OSCs) with accession numbers**. OSCs used in the phylogenetic analyses in Figure 2.

| **Enzyme name** | **Organism** | **Abbreviation** | **Common abbreviation** | **Accession no Protein** |
|---|---|---|---|---|
| Poaceatapetol synthase | *Oryza sativa v. japonica* | OsPTS | OsPTS1, Os08g12730, OsOSC12 | XM_015795471 |
| Onocerin synthase | *Lycopodium clavatum* | LycONS | LcLCD | XM_015795471 |
| Pre-alpha onocerin synthase | *Lycopodium clavatum* | LycpONS | LcLCC | BAU46472 |
| Cycloartenol synthase | *Adiantum capillus-veneris* | AcvCAS | ACX | BAU46471 |
| Isoarborinol synthase | *Oryza sativa v. japonica* | OsIAS | Os11g35710, OsOSC11 | AK067451 (DNA) |
| β-Amyrin synthase | *Avena strigosa* | AsbAS | AsBAS1 | CAC84558 |
| Achilleol B synthase | *Oryza sativa v. japonica* | OsACS | Os11g18194, OsOSC8 | BAG92014 |
| Mixed amyrin synthase | *Oryza sativa v. japonica* | OsMAS | Os06g28820, OsOSC6 | AGG20314 |
| Lupeol synthase | *Iris tectorum* | ItLUS | ItOSC6 | QQR13797 |
| Multifunctional α-amyrin synthase | *Iris tectorum* | ItMTS | ItOSC2 | QQR13795 |
| Multifunctional triterpene synthase | *Costus speciosus* | CspMTS | CsOSC2, CSV | BAB83254 |
| Cycloartenol synthase | *Costus speciosus* | CspCAS | CsOSC1, CSI | BAB83253 |
| Parkeol synthase | *Oryza sativa v. japonica* | OsPS | Os11g08569, OsOSC7 | BAG89914 |
| Cycloartenol synthase | *Avena strigosa* | AsCAS | AsSC1 | AAT38891 |
| Cycloartenol synthase | *Oryza sativa v. japonica* | OsCAS | Os02g04710, OsOSC2 | BAH00370 |
| Cycloartenol synthase | *Iris tectorum* | ItCAS | ItOSC3 | QQR13796 |
| Cycloartenol synthase | *Paris polyphylla* | PpCAS | pPCAS | QIN54760 |
| Cycloartenol synthase | *Fritillaria thunbergii* | FtCAS | | AEO27878 |
| Cucurbitadienol synthase | *Cucurbita pepo* | CpCDS | CPQ | BAD34645 |
| Cycloartenol synthase | *Betula platyphylla* | BpCAS | BPX | BAB83085 |
| Cucurbitadienol synthase | *Momordica charantia* | McCDS | McCBS | BBN67121 |
| Cycloartenol synthase | *Cucurbita pepo* | CpCAS | CPX | BAD34644 |
| Cycloartenol synthase | *Momordica charantia* | McCAS | - | BBN67123 |
| Cycloartenol synthase | *Betula platyphylla* | BpCAS2 | BPX2 | BAB83086 |
| Cycloartenol synthase | *Lagerstroemia speciosa* | LspCAS | LsOSC5 | AZS32331 |

| | | | | |
|---|---|---|---|---|
| Cycloartenol synthase | *Pisum sativum* | PsCAS | PNX | BAA23533 |
| Cycloartenol synthase | *Lotus japonicus* | LjCAS | OSC5 | BAE53431 |
| Cycloartenol synthase | *Glycyrrhiza glabra* | GgCAS | GgCAS1 | BAA76902 |
| Cycloartenol synthase | *Arabidopsis thaliana* | AtCAS | | AAC04931 |
| Cycloartenol synthase | *Ricinus communis* | RcCAS | | ABB76767 |
| Cycloartenol synthase | *Rhizophora stylosa* | RsCAS | | BAF73929 |
| Cycloartenol synthase | *Kandelia candel* | KcCAS | | BAF73930 |
| Cycloartenol synthase | *Panax ginseng* | PgCAS | PNX | BAA33460 |
| Cycloartenol synthase | *Kalanchoe daigremontiana* | KdCAS | - | ADK35127 |
| Cycloartenol synthase | *Withania somnifera* | WsCAS | WsOSC/CS | ADG60271 |
| Cycloartenol synthase | *Nicotiana tabacum* | NtCAS | | AIY33888 |
| Cycloartenol synthase | *Artemisia annua* | AaCAS | | AJE29378 |
| Predicted cycloartenol synthase | *Calendula officinalis* | CoOSC12 | | |
| Predicted cycloartenol synthase | *Calendula officinalis* | CoOSC13 | | |
| Predicted cycloartenol synthase | *Calendula officinalis* | CoOSC14 | | |
| Predicted cycloartenol synthase | *Calendula officinalis* | CoOSC15 | | |
| Predicted cycloartenol synthase | *Calendula officinalis* | CoOSC16 | | |
| Lanosterol synthase | *Lotus japonicus* | LjLAS | cOSC7 | BAE95410 |
| Lanosterol synthase | *Arabidopsis thaliana* | AtLAS | LAS1 | BAE95408 |
| Lanosterol synthase | *Panax ginseng* | PgLAS | PNZ | BAA33462 |
| Predicted lanosterol synthase | *Calendula officinalis* | CoOSC26 | | |
| Predicted lanosterol synthase | *Calendula officinalis* | CoOSC27 | | |
| Lupeol synthase | *Glycyrrhiza glabra* | GgLUS | LUS1 | BAD08587 |
| Lupeol synthase | *Lotus japonicus* | LjLUS | OSC3 | BAE53430 |
| Lupeol synthase | *Lagerstroemia speciosa* | LspLUS | LsOSC1 | AZS32327 |
| Lupeol synthase | *Betula platyphylla* | BpLUS | BPW | BAB83087 |
| Lupeol synthase | *Withania somnifera* | WsLUS | WsOSC/LS | AGA17939 |
| Onocerin synthase | *Ononis spinosa* | OsONS | OsONS1 | AUD09560 |
| Lupeol synthase | *Olea europaea* | OeLUS | OEW | BAA86930 |

| | | | | |
|---|---|---|---|---|
| Predicted lupeol synthase | *Calendula officinalis* | CoOSC7 | | |
| Predicted lupeol synthase | *Calendula officinalis* | CoOSC25 | | |
| Lupeol synthase | *Artemisia annua* | AaLUS | | AJE29379 |
| Lupeol synthase | *Lactuca sativa* | LsLUS | LsOSC5 | QIB85404 |
| Lupeol synthase | *Taraxacum kok-saghyz* | TkLUS | TkLUP | AXU93517 |
| Lupeol synthase | *Taraxacum officinale* | ToLUS | TRW | BAA86932 |
| Mixed amyrin synthase | *Ilex asprella* | IaMAS | IaAS1 | AIS39793 |
| Mixed amyrin synthase | *Catharanthus roseus* | CrMAS | CrAS | AEX99665 |
| Mixed amyrin synthase | *Ocimum basilicum* | ObMAS | ObAS2 | AFH53506 |
| Mixed amyrin synthase | *Olea europaea* | OeMAS | OEA | BAF63702 |
| Bauerenol synthase | *Lactuca sativa* | LsBAUS | LsOSC2 | QIB85401 |
| Bauerenol synthase | *Taraxacum coreanum* | TcBAUS | TcOSC2 | QBO24612 |
| Bauerenol synthase | *Calendula officinalis* | CoBAUS | | |
| Predicted bauerenol synthase | *Calendula officinalis* | CoOSC24 | | |
| Predicted novel/multifunctional triterpene synthase | *Calendula officinalis* | CoOSC6 | | |
| Predicted novel/multifunctional triterpene synthase | *Calendula officinalis* | CoOSC19 | | |
| Predicted novel/multifunctional triterpene synthase | *Calendula officinalis* | CoOSC5 | | |
| Predicted novel/multifunctional triterpene synthase | *Calendula officinalis* | CoOSC20 | | |
| Multifunctional triterpene synthase | *Artemisia annua* | AaMAS | OSC2 | AHF22084 |
| Predicted mixed-amyrin synthase | *Calendula officinalis* | CoOSC23 | | |
| Mixed amyrin synthase | *Calendula officinalis* | CoMAS | CoOSC3 | |
| Predicted mixed-amyrin synthase | *Calendula officinalis* | CoOSC2 | | |
| Predicted mixed-amyrin synthase | *Calendula officinalis* | CoOSC18 | | |
| Predicted mixed-amyrin synthase | *Calendula officinalis* | CoOSC28 | | |
| Dammarenediol-II synthase | *Panax notoginseng* | PngDDS | PnDS | AGS79229 |
| Dammarenediol-II synthase | *Panax ginseng* | PgDDS | PNA | BAF33291 |
| Dammarenediol-II synthase | *Panax quinquefolius* | PqDDS | - | AED99864 |
| Dammarenediol-II synthase | *Centella asiatica* | CaDDS | | AAS01523 |
| Predicted taraxasterol synthase | *Carthamus tinctorius* | CtTXSS | | LUCG01052148.1 (DNA) |

| | | | | |
|---|---|---|---|---|
| Predicted taraxasterol synthase | *Atractylodes lancea* | AlTXSS | | GEGA01000112.1 (DNA) |
| Taraxasterol synthase | *Cynara cardunculus* | CcTXSS | | XP_024993293 |
| Y | *Tragopogon pratensis* | TprTXSS | | FUPX-2015344 (1KP) |
| Taraxasterol synthase | *Tragopogon dubius* | TdTXSS | | DDRL-2016809 (1KP) |
| Predicted taraxasterol synthase | *Cichorium intybus* | CiTXSS | | GGQG01003518.1 |
| Taraxasterol synthase | *Cichorium endivia* | CeTXSS | | GGQM01030486.1 |
| Multifunctional triterpene synthase | *Taraxacum coreanum* | TcTXSS | TcOSC1 | QBO24611 |
| Taraxasterol synthase | *Taraxacum kok-saghyz* | TkTXSS | TkOSC1 | AXU93516 |
| Taraxasterol synthase | *Lactuca sativa* | LsTXSS | LsOSC1 | QIB85400 |
| Taraxasterol synthase | *Calendula arvensis* | CarTXSS | CaOSC1 | |
| Taraxasterol synthase | *Calendula officinalis* | CoTXSS | CoOSC1 | |
| Predicted taraxasterol synthase | *Calendula officinalis* | CoOSC17 | | |
| Predicted taraxasterol synthase | *Mikania micrantha* | MmTXSS | | KAD1358637.1 |
| Predicted taraxasterol synthase | *Tagetes erecta* | TeTXSS | | GGGQ01031191.1 |
| Predicted taraxasterol synthase | *Helianthus niveus* | HnTXSS | | GEWS01056763.1 |
| Taraxasterol synthase | *Helianthus annuus* | HaTXSS | | XP_022035366 |
| Predicted taraxasterol synthase | *Tanacetum parthenium* | TpaTXSS | | DUQG-2008134 (1KP) |
| Predicted taraxasterol synthase | *Chrysanthemum seticuspe* | CseTXSS | | Cse_sc039633.1_g010 |
| Predicted taraxasterol synthase | *Artemisia annua* | AaTXSS | | PWA47914.1 |
| Multifunctional triterpene synthase | *Tripterygium wilfordii* | TwMTS1 | TwOSC1 | AWK97810 |
| Multifunctional friedelin synthase | *Tripterygium wilfordii* | TwMTS2 | TwOSC3 | AWK97812 |
| β-Amyrin synthase | *Nigella sativa* | NsbAS | NsBAS1 | ACH88048 |
| Multifunctional triterpene synthase | *Rhizophora stylosa* | RsMTS2 | RsM2 | BAF80442 |
| Isomultiflorenol synthase | *Luffa cylindrica* | LcIMS | LcIMS1 | BAB68529 |
| Isomultiflorenol synthase | *Momordica charantia* | McIMS | - | BBN67124 |
| Tirucalladienol synthase | *Citrus sinensis* | CsTCS | CsOSC1 | XP_006468116 |
| Tirucalladienol synthase | *Azadirachta indica* | AiTCS | AiOSC1 | MK803262 |
| Tirucalladienol synthase | *Melia azedarach* | MaTCS | MaOSC1 | MK803261 |
| Mixed amyrin synthase | *Lagerstroemia speciosa* | LspMAS | LsOSC2 | AZS32328 |

| | | | | |
|---|---|---|---|---|
| Lupeol synthase | *Ricinus communis* | RcLUS | | ABB76766 |
| Lupeol synthase | *Bruguiera gymnorhiza* | BgLUS | | BAF80444 |
| Multifunctional triterpene synthase | *Kandelia candel* | KcMTS | KcMS | BAF35580 |
| Lupeol synthase | *Citrus sinensis* | CsLUS | CsOSC3 | XP_015382484 |
| Multifunctional triterpene synthase | *Arabidopsis thaliana* | AtPEN6 | At1g78500, T30F21.16 | Q9SYN1 |
| Marneral synthase | *Arabidopsis thaliana* | AtMARS | MRN1, PEN5 | NP_199074 |
| Tirucalladienol synthase | *Arabidopsis thaliana* | AtPEN3 | At5g36150 | Q9LVY2 |
| Thalianol synthase | *Arabidopsis thaliana* | AtTHAS | At5g48010 | Q9FI37 |
| Arabidiol synthase | *Arabidopsis thaliana* | AtARS | At4g15340, PEN1 | BAF33292 |
| Baurol synthase | *Arabidopsis thaliana* | AtBARS | At4g15370, BARS1, PEN2 | NP_193272 |
| β-Amyrin synthase | *Arabidopsis thaliana* | AtbAS | AtbAS | BAG82628 |
| Camelliol synthase | *Arabidopsis thaliana* | AtCAMS | CAMS1 | NP_683508 |
| Multifunctional triterpene synthase | *Arabidopsis thaliana* | AtLUP5 | At1g66960, F1019.4 | NP_176868 |
| Lupeol synthase | *Arabidopsis thaliana* | AtLUP1 | LUP1 | AAD05032 |
| Multifunctional triterpene synthase | *Arabidopsis thaliana* | AtLUP2 | | NP_178017 |
| β-Amyrin synthase | *Citrus sinensis* | CsbAS | CsOSC2 | XP_024957905.1 |
| β-Amyrin synthase | *Gypsophila vaccaria* | GvbAS | SvBS | ABK76265 |
| β-Amyrin synthase | *Polygala tenuifolia* | PtbAS | PtBS | ABL07607 |
| Multifunctional triterpene synthase | *Lotus japonicus* | LjMTS | LjAMY2 | AAO33580 |
| β-Amyrin synthase | *Medicago truncatula* | MtbAS | MtAMY1 | AAO33578 |
| β-Amyrin synthase | *Pisum sativum* | PsbAS | PSY | BAA97558 |
| β-Amyrin synthase | *Lotus japonicus* | LjbAS | LjAMY1 | AAO33579 |
| Mixed amyrin synthase | *Pisum sativum* | PsMAS | PSM | BAA97559 |
| β-Amyrin synthase | *Glycyrrhiza glabra* | GgbAS | GgbAS1 | BAA89815 |
| β-Amyrin synthase | *Glycyrrhiza uralensis* | GubAS | | ACV21067 |
| β-Amyrin synthase | *Euphorbia tirucalli* | EtbAS | EtAS | BAE43642 |
| β-Amyrin synthase | *Bruguiera gymnorhiza* | BgbAS | | BAF80443 |
| Multifunctional triterpene synthase | *Rhizophora stylosa* | RsMTS1 | RsM1 | BAF80441 |
| β-Amyrin synthase | *Tripterygium wilfordii* | TwbAS | TwOSC2 | AWK97811 |

| | | | | |
|---|---|---|---|---|
| β-Amyrin synthase | *Lagerstroemia speciosa* | LspbAS | LsOSC3 | AZS32329 |
| β-Amyrin synthase | *Lagerstroemia speciosa* | LspbAS2 | LsOSC4 | AZS32330 |
| β-Amyrin synthase | *Betula platyphylla* | BpbAS | BPY | BAB83088 |
| β-Amyrin synthase | *Momordica charantia* | McbAS | McBAS | BBN67122 |
| β-Amyrin synthase | *Platycodon grandiflorus* | PgrbAS | - | ASB17950 |
| Taraxerol synthase | *Kalanchoe daigremontiana* | KdTXS | KdTAS | ADK35123 |
| Friedelin synthase | *Kalanchoe daigremontiana* | KdFRS | - | ADK35125 |
| Glutinol synthase | *Kalanchoe daigremontiana* | KdGLS | - | ADK35124 |
| Lupeol synthase | *Kalanchoe daigremontiana* | KdLUS | - | ADK35126 |
| β-Amyrin synthase | *Gentiana straminea* | GsbAS | GsAS1 | ACO24697 |
| Multifunctional triterpene synthase | *Solanum lycopersicum* | SlMTS | SlTTS2 | ADU52575 |
| β-Amyrin synthase | *Solanum lycopersicum* | SlbAS | SlTTS1 | ADU52574 |
| β-Amyrin synthase | *Withania somnifera* | WsbAS | WsOSC/BS | AGA17940 |
| β-Amyrin synthase | *Ocimum basilicum* | ObbAS | ObAS1 | AHJ10507 |
| β-Amyrin synthase | *Ilex asprella* | IabAS | IaAS2 | AIS39794 |
| Taraxerol synthase | *Lactuca sativa* | LsTXS | LsOSC4 | QIB85403 |
| Taraxerol synthase | *Taraxacum coreanum* | TcTXS | TsOSC4 | QBO24614 |
| β-Amyrin synthase | *Eleutherococcus senticosus* | EsbAS | - | APZ88354 |
| β-Amyrin synthase | *Kalopanax septemlobus* | KsbAS | KsBAS | ALO23119 |
| β-Amyrin synthase | *Panax ginseng* | PgbAS | PNY | BAA33461 |
| β-Amyrin synthase | *Aralia elata* | AebAS | AeAS | ADK12003 |
| Shionone synthase | *Calendula officinalis* | CoSHS | CoOSC8 | |
| Pseudogene (Shionone synthase) | *Calendula officinalis* | CoOSC21 | | |
| Pseudogene (Shionone synthase) | *Calendula officinalis* | CoOSC22 | | |
| Baccharis oxide synthase | *Stevia rebaudiana* | SrBOS | StrBOS | BAH23676 |
| Shionone synthase | *Aster tataricus* | AtaSHS | AtaSHS1 | BAK52535 |
| Predicted β-amyrin-like synthase | *Calendula officinalis* | CoOSC9 | | |
| Predicted β-amyrin-like synthase | *Calendula officinalis* | CoOSC10 | | |
| Predicted β-amyrin synthase | *Calendula officinalis* | CoOSC11 | | |

| | | | | |
|---|---|---|---|---|
| β-Amyrin synthase | *Aster sedifolius* | AsebAS | AsOXA1 | AAX14716 |
| β-Amyrin synthase | *Artemisia annua* | AabAS | AaBAS | ACA13386 |
| β-Amyrin synthase | *Lactuca sativa* | LsbAS | LsOSC3 | QIB85402 |
| β-Amyrin synthase | *Taraxacum coreanum* | TcBAS | TcOSC3 | QBO24613 |
| β-Amyrin synthase | *Taraxacum kok-saghyz* | TkbAS | TkOSC6 | AXU93522 |

**Supplementary Table 10.** Taxa and accession numbers of sequences used for positive selection analysis shown in Figure 3B.

| Abbreviation | Enzyme name | Species | Accession no |
|---|---|---|---|
| AaMAS | Multifunctional triterpene synthase | *Artemisia annua* | AHF22084 |
| CoMAS | Mixed amyrin synthase | *Calendula officinalis* | This study |
| CrMAS | Mixed amyrin synthase | *Catharanthus roseus* | AEX99665 |
| IaMAS | Mixed amyrin synthase | *Ilex asprella* | AIS39793 |
| ObMAS | Mixed amyrin synthase | *Ocimum basilicum* | AFH53506 |
| OeMAS | Mixed amyrin synthase | *Olea europaea* | BAF63702 |
| CcTXSS | Taraxasterol synthase | *Cynara cardunculus* | XP_024993293 |
| CoTXSS | Taraxasterol synthase | *Calendula officinalis* | This study |
| HaTXSS | Taraxasterol synthase | *Helianthus annuus* | XP_022035366 |
| LsTXSS | Taraxasterol synthase | *Lactuca sativa* | QIB85400 |
| TcTXSS | Taraxasterol synthase | *Taraxacum coreanum* | QBO24611 |
| TkTXSS | Taraxasterol synthase | *Taraxacum kok-saghyz* | AXU93516 |
| TdTXSS | Taraxasterol synthase | *Tragopogon dubius* | DDRL-2016809 (1KP) |
| CeTXSS | Taraxasterol synthase | *Cichorium endivia* | GGQM01030486.1 |

**Supplementary Table 11.** Statistical tests used in Figure 3.

| Experiment | Sample | p.adjusted |
|---|---|---|
| TXSS infiltration (N=6)<br>Kruskal Wallis<br>Posthoc pair Wilcoxon test<br>(Benjamini-Hochberg correction)<br>Psi+taraxasterol | CoMAS-CoMAS E371D | 0.0062 |
| | CoMAS-CoMAS I367M | 0.0036 |
| | CoMAS-CoMAS I367M+E371D | 0.0036 |
| | CoMAS-CoTXSS | 0.0036 |
| | CoMAS E371D-CoMAS I367M | 0.0036 |
| | CoMAS E371D-CoMAS I367M E371D | 0.0036 |
| | CoMAS E371D-CoTXSS | 0.0036 |
| | CoMAS I367M-CoMAS I367M E371D | 0.6061 |
| | CoMAS I367M-CoTXSS | 0.8182 |
| | CoMAS I367M E371D-CoTXSS | 0.6542 |
| TXSS infiltration (N=6)<br>Kruskal Wallis<br>Posthoc pair Wilcoxon test<br>(Benjamini-Hochberg correction)<br>alpha + beta amyrin | CoMAS-CoMAS E371D | 0.1797 |
| | CoMAS-CoMAS I367M | 0.0124 |
| | CoMAS-CoMAS I367M+E371D | 0.0036 |
| | CoMAS-CoTXSS | 0.0036 |
| | CoMAS E371D-CoMAS I367M | 0.0457 |
| | CoMAS E371D-CoMAS I367M E371D | 0.0036 |
| | CoMAS E371D-CoTXSS | 0.0036 |
| | CoMAS I367M-CoMAS I367M E371D | 0.0036 |
| | CoMAS I367M-CoTXSS | 0.0036 |
| | CoMAS I367M E371D-CoTXSS | 0.0325 |

**Supplementary Table 12.** Statistical tests used in Figure 4.

| Experiment | Sample | p.adjusted |
|---|---|---|
| TXSS infiltration (N=6)<br>Kruskal Wallis<br>Posthoc pair Wilcoxon test<br>(Benjamini-Hochberg correction)<br>Psi-taraxasterol | CoTXSS-CoTXSS G380T | 0.048 |
| | CoTXSS-CoTXSS D385E | 0.0032 |
| | CoTXSS-CoTXSS H492Q | 0.8182 |
| | CoTXSS-CoTXSS P751A | 0.048 |
| | CoTXSS-CoTXSS D385E+H492Q | 0.0032 |
| | CoTXSS-CoTXSS G380T+D385E+H492Q+P751A | 0.0032 |
| | CoTXSS G380T-CoTXSS D385E | 0.0032 |
| | CoTXSS G380T-CoTXSS H492Q | 0.0341 |
| | CoTXSS G380T-CoTXSS P751A | 0.0121 |
| | CoTXSS G380T-CoTXSS D385E+H492Q | 0.0032 |
| | CoTXSS G380T-CoTXSS G380T+D385E+H492Q+P751A | 0.0032 |
| | CoTXSS D385E-CoTXSS H492Q | 0.0032 |
| | CoTXSS D385E-CoTXSS P751A | 0.0032 |
| | CoTXSS D385E-CoTXSS D385E+H492Q | 0.0032 |
| | CoTXSS D385E-CoTXSS G380T+D385E+H492Q+P751A | 0.0032 |
| | CoTXSS H492Q-CoTXSS P751A | 0.0682 |
| | CoTXSS H492Q-CoTXSS D385E+H492Q | 0.0032 |
| | CoTXSS H492Q-CoTXSS G380T+D385E+H492Q+P751A | 0.0032 |
| | CoTXSS P751A-CoTXSS D385E+H492Q | 0.0032 |

| | | |
|---|---|---|
| | CoTXSS P751A-CoTXSS G380T+D385E+H492Q+P751A | 0.0032 |
| | CoTXSS D385E+H492Q-CoTXSS G380T+D385E+H492Q+P751A | 0.0682 |
| | TkTXSS-TkTXSS T374G | 0.0167 |
| | TkTXSS-TkTXSS E379D | 0.0027 |
| | TkTXSS-TkTXSS Q486H | 0.0027 |
| | TkTXSS-TkTXSS A745P | 0.5887 |
| | TkTXSS-TkTXSS E379D+Q486H | 0.0027 |
| | TkTXSS-TkTXSS T374G+E379D+Q486H+A745P | 0.0027 |
| | TkTXSS T374G-TkTXSS E379D | 0.0027 |
| | TkTXSS T374G-TkTXSS Q486H | 0.0027 |
| | TkTXSS T374G-TkTXSS A745P | 0.0027 |
| | TkTXSS T374G-TkTXSS E379D+Q486H | 0.0027 |
| | TkTXSS T374G-TkTXSS T374G+E379D+Q486H+A745P | 0.0027 |
| | TkTXSS E379D-TkTXSS Q486H | 0.4136 |
| | TkTXSS E379D-TkTXSS A745P | 0.0027 |
| | TkTXSS E379D-TkTXSS E379D+Q486H | 0.0027 |
| | TkTXSS E379D-TkTXSS T374G+E379D+Q486H+A745P | 0.0027 |
| | TkTXSS Q486H-TkTXSS A745P | 0.0027 |
| | TkTXSS Q486H-TkTXSS E379D+Q486H | 0.0027 |
| | TkTXSS Q486H-TkTXSS | 0.0027 |

| | | |
|---|---|---|
| | T374G+E379D+Q486H+A745P | |
| | TkTXSS A745P-TkTXSS E379D+Q486H | 0.0027 |
| | TkTXSS A745P-TkTXSS T374G+E379D+Q486H+A745P | 0.0027 |
| | TkTXSS E379D+Q486H-TkTXSS T374G+E379D+Q486H+A745P | 0.0051 |
| TXSS infiltration (N=6) Kruskal Wallis Posthoc pair Wilcoxon test (Benjamini-Hochberg correction) Taraxasterol | CoTXSS-CoTXSS G380T | 0.4062 |
| | CoTXSS-CoTXSS D385E | 0.0065 |
| | CoTXSS-CoTXSS H492Q | 0.6507 |
| | CoTXSS-CoTXSS P751A | 0.4062 |
| | CoTXSS-CoTXSS D385E+H492Q | 0.4866 |
| | CoTXSS-CoTXSS G380T+D385E+H492Q+P751A | 0.8591 |
| | CoTXSS G380T-CoTXSS D385E | 0.0065 |
| | CoTXSS G380T-CoTXSS H492Q | 0.0065 |
| | CoTXSS G380T-CoTXSS P751A | 0.9372 |
| | CoTXSS G380T-CoTXSS D385E+H492Q | 0.0682 |
| | CoTXSS G380T-CoTXSS G380T+D385E+H492Q+P751A | 0.3144 |
| | CoTXSS D385E-CoTXSS H492Q | 0.0065 |
| | CoTXSS D385E-CoTXSS P751A | 0.0065 |
| | CoTXSS D385E-CoTXSS D385E+H492Q | 0.0065 |
| | CoTXSS D385E-CoTXSS G380T+D385E+H492Q+P751A | 0.0065 |
| | CoTXSS H492Q-CoTXSS P751A | 0.3881 |

| | | |
|---|---|---|
| | CoTXSS H492Q-CoTXSS D385E+H492Q | 0.6507 |
| | CoTXSS H492Q-CoTXSS G380T+D385E+H492Q+P751A | 0.2172 |
| | CoTXSS P751A-CoTXSS D385E+H492Q | 0.3144 |
| | CoTXSS P751A-CoTXSS G380T+D385E+H492Q+P751A | 0.4062 |
| | CoTXSS D385E+H492Q-CoTXSS G380T+D385E+H492Q+P751A | 0.3144 |
| | TkTXSS-TkTXSS T374G | 0.1631 |
| | TkTXSS-TkTXSS E379D | 0.0032 |
| | TkTXSS-TkTXSS Q486H | 0.0032 |
| | TkTXSS-TkTXSS A745P | 0.2656 |
| | TkTXSS-TkTXSS E379D+Q486H | 0.0061 |
| | TkTXSS-TkTXSS T374G+E379D+Q486H+A745P | 0.0032 |
| | TkTXSS T374G-TkTXSS E379D | 0.0032 |
| | TkTXSS T374G-TkTXSS Q486H | 0.0032 |
| | TkTXSS T374G-TkTXSS A745P | 0.4848 |
| | TkTXSS T374G-TkTXSS E379D+Q486H | 0.2656 |
| | TkTXSS T374G-TkTXSS T374G+E379D+Q486H+A745P | 0.0032 |
| | TkTXSS E379D-TkTXSS Q486H | 0.0032 |
| | TkTXSS E379D-TkTXSS A745P | 0.0032 |
| | TkTXSS E379D-TkTXSS E379D+Q486H | 0.0032 |
| | TkTXSS E379D-TkTXSS | 0.0032 |

| | | |
|---|---|---|
| | T374G+E379D+Q486H+A745P | |
| | TkTXSS Q486H-TkTXSS A745P | 0.0032 |
| | TkTXSS Q486H-TkTXSS E379D+Q486H | 0.0032 |
| | TkTXSS Q486H-TkTXSS T374G+E379D+Q486H+A745P | 0.0032 |
| | TkTXSS A745P-TkTXSS E379D+Q486H | 0.0114 |
| | TkTXSS A745P-TkTXSS T374G+E379D+Q486H+A745P | 0.0032 |
| | TkTXSS E379D+Q486H-TkTXSS T374G+E379D+Q486H+A745P | 0.0032 |

**Supplementary Table 13.** Statistical tests used in Figure 5.

| Experiment | Sample | p.adjusted |
|---|---|---|
| CoCYP mutants infiltration (N=6) Kruskal Wallis Posthoc pair Wilcoxon test (Benjamini-Hochberg correction) Beta-amyrin to psi-taraxasterol ratio | CoTXSS-CoTXSS + CoCYP716A392 | 0.0054 |
| | CoTXSS-CoTXSS + CoCYP716A392 (A285G) | 0.0054 |
| | CoTXSS-CoTXSS + CoCYP716A392 (A285V) | 0.0077 |
| | CoTXSS-CoTXSS + CoCYP716A392 (A357L) | 0.085 |
| | CoTXSS-CoTXSS + CoCYP716A392 (H424R) | 0.0054 |
| | CoTXSS-CoTXSS + CoCYP716A392 (A285V) | 0.0054 |
| | CoTXSS-CoTXSS + CoCYP716A393 (A285G) | 0.0054 |
| | CoTXSS-CoTXSS + CoCYP716A393 (A285V) | 0.0054 |
| | CoTXSS-CoTXSS + CoCYP716A393 (A357L) | 0.0054 |
| | CoTXSS-CoTXSS + CoCYP716A393 (H424R) | 0.0054 |
| | CoTXSS + CoCYP716A392-CoTXSS + CoCYP716A392 (A285G) | 1 |
| | CoTXSS + CoCYP716A392-CoTXSS + CoCYP716A392 (A285V) | 0.0077 |
| | CoTXSS + CoCYP716A392-CoTXSS + CoCYP716A392 (A357L) | 0.0386 |
| | CoTXSS + CoCYP716A392-CoTXSS + CoCYP716A392 (H424R) | 0.5128 |
| | CoTXSS + CoCYP716A392-CoTXSS + CoCYP716A392 (H424R) | 0.5996 |
| | CoTXSS + CoCYP716A392-CoTXSS + CoCYP716A393 (A285G) | 0.2753 |
| | CoTXSS + CoCYP716A392-CoTXSS + CoCYP716A393 (A285V) | 0.0054 |
| | CoTXSS + CoCYP716A392-CoTXSS + CoCYP716A393 (A357L) | 0.3474 |

| | | |
|---|---|---|
| | CoTXSS + CoCYP716A392-CoTXSS + CoCYP716A393 (H424R) | 0.085 |
| | CoTXSS + CoCYP716A392 (A285G)-CoTXSS + CoCYP716A392 (A285V) | 0.0077 |
| | CoTXSS + CoCYP716A392 (A285G)-CoTXSS + CoCYP716A392 (A357L) | 0.0054 |
| | CoTXSS + CoCYP716A392 (A285G)-CoTXSS + CoCYP716A392 (H424R) | 0.085 |
| | CoTXSS + CoCYP716A392 (A285G)-CoTXSS + CoCYP716A393 | 0.5996 |
| | CoTXSS + CoCYP716A392 (A285G)-CoTXSS + CoCYP716A393 (A285G) | 0.4248 |
| | CoTXSS + CoCYP716A392 (A285G)-CoTXSS + CoCYP716A393 (A285V) | 0.0054 |
| | CoTXSS + CoCYP716A392 (A285G)-CoTXSS + CoCYP716A393 (A357L) | 0.0238 |
| | CoTXSS + CoCYP716A392 (A285G)-CoTXSS + CoCYP716A393 (H424R) | 0.0054 |
| | CoTXSS + CoCYP716A392 (A285V)-CoTXSS + CoCYP716A392 (A357L) | 0.0382 |
| | CoTXSS + CoCYP716A392 (A285V)-CoTXSS + CoCYP716A392 (H424R) | 0.0077 |
| | CoTXSS + CoCYP716A392 (A285V)-CoTXSS + CoCYP716A393 | 0.0077 |
| | CoTXSS + CoCYP716A392 (A285V)-CoTXSS + CoCYP716A393 (A285G) | 0.0077 |
| | CoTXSS + CoCYP716A392 (A285V)-CoTXSS + CoCYP716A393 (A285V) | 0.1113 |
| | CoTXSS + CoCYP716A392 (A285V)-CoTXSS + CoCYP716A393 (A357L) | 0.0077 |

| | | |
|---|---|---|
| | CoTXSS + CoCYP716A392 (A285V)-CoTXSS + CoCYP716A393 (H424R) | 0.0077 |
| | CoTXSS + CoCYP716A392 (A357L)-CoTXSS + CoCYP716A392 (H424R) | 0.0149 |
| | CoTXSS + CoCYP716A392 (A357L)-CoTXSS + CoCYP716A393 | 0.0054 |
| | CoTXSS + CoCYP716A392 (A357L)-CoTXSS + CoCYP716A393 (A285G) | 0.0054 |
| | CoTXSS + CoCYP716A392 (A357L)-CoTXSS + CoCYP716A393 (A285V) | 0.085 |
| | CoTXSS + CoCYP716A392 (A357L)-CoTXSS + CoCYP716A393 (A357L) | 0.0054 |
| | CoTXSS + CoCYP716A392 (A357L)-CoTXSS + CoCYP716A393 (H424R) | 0.1113 |
| | CoTXSS + CoCYP716A392 (H424R)-CoTXSS + CoCYP716A393 | 0.1113 |
| | CoTXSS + CoCYP716A392 (H424R)-CoTXSS + CoCYP716A393 (A285G) | 0.085 |
| | CoTXSS + CoCYP716A392 (H424R)-CoTXSS + CoCYP716A393 (A285V) | 0.0054 |
| | CoTXSS + CoCYP716A392 (H424R)-CoTXSS + CoCYP716A393 (A357L) | 0.4248 |
| | CoTXSS + CoCYP716A392 (H424R)-CoTXSS + CoCYP716A393 (H424R) | 0.1113 |
| | CoTXSS + CoCYP716A393-CoTXSS + CoCYP716A393 (A285G) | 0.2753 |
| | CoTXSS + CoCYP716A393-CoTXSS + CoCYP716A393 (A285V) | 0.0054 |
| | CoTXSS + CoCYP716A393-CoTXSS + CoCYP716A393 (A357L) | 0.0238 |
| | CoTXSS + CoCYP716A393-CoTXSS + CoCYP716A393 (H424R) | 0.0054 |

| | | |
|---|---|---|
| | CoTXSS + CoCYP716A393 (A285G)-CoTXSS + CoCYP716A393 (A285V) | 0.0054 |
| | CoTXSS + CoCYP716A393 (A285G)-CoTXSS + CoCYP716A393 (A357L) | 0.0238 |
| | CoTXSS + CoCYP716A393 (A285G)-CoTXSS + CoCYP716A393 (H424R) | 0.0054 |
| | CoTXSS + CoCYP716A393 (A285V)-CoTXSS + CoCYP716A393 (A357L) | 0.0054 |
| | CoTXSS + CoCYP716A393 (A285V)-CoTXSS + CoCYP716A393 (H424R) | 0.0054 |
| | CoTXSS + CoCYP716A393 (A357L)-CoTXSS + CoCYP716A393 (H424R) | 0.0077 |
| CoCYP mutants infiltration (N=6) Kruskal Wallis Posthoc pair Wilcoxon test (Benjamini-Hochberg correction) Total content of beta-amyrin and psi-taraxasterol | CoTXSS-CoTXSS + CoCYP716A392 | 0.0034 |
| | CoTXSS-CoTXSS + CoCYP716A392 (A285G) | 0.0066 |
| | CoTXSS-CoTXSS + CoCYP716A392 (A285V) | 0.0034 |
| | CoTXSS-CoTXSS + CoCYP716A392 (A357L) | 0.0034 |
| | CoTXSS-CoTXSS + CoCYP716A392 (H424R) | 0.0034 |
| | CoTXSS-CoTXSS + CoCYP716A392 (A285V) | 0.0034 |
| | CoTXSS-CoTXSS + CoCYP716A393 (A285G) | 0.0034 |
| | CoTXSS-CoTXSS + CoCYP716A393 (A285V) | 0.0034 |
| | CoTXSS-CoTXSS + CoCYP716A393 (A357L) | 0.0916 |
| | CoTXSS-CoTXSS + CoCYP716A393 (H424R) | 0.0916 |
| | CoTXSS + CoCYP716A392-CoTXSS + CoCYP716A392 (A285G) | 0.0034 |

| | | |
|---|---|---|
| | CoTXSS + CoCYP716A392-CoTXSS + CoCYP716A392 (A285V) | 0.5674 |
| | CoTXSS + CoCYP716A392-CoTXSS + CoCYP716A392 (A357L) | 0.3073 |
| | CoTXSS + CoCYP716A392-CoTXSS + CoCYP716A392 (H424R) | 0.7255 |
| | CoTXSS + CoCYP716A392-CoTXSS + CoCYP716A392 (H424R) | 0.6476 |
| | CoTXSS + CoCYP716A392-CoTXSS + CoCYP716A393 (A285G) | 0.0034 |
| | CoTXSS + CoCYP716A392-CoTXSS + CoCYP716A393 (A285V) | 0.6476 |
| | CoTXSS + CoCYP716A392-CoTXSS + CoCYP716A393 (A357L) | 0.0034 |
| | CoTXSS + CoCYP716A392-CoTXSS + CoCYP716A393 (H424R) | 0.0034 |
| | CoTXSS + CoCYP716A392 (A285G)-CoTXSS + CoCYP716A392 (A285V) | 0.0034 |
| | CoTXSS + CoCYP716A392 (A285G)-CoTXSS + CoCYP716A392 (A357L) | 0.0034 |
| | CoTXSS + CoCYP716A392 (A285G)-CoTXSS + CoCYP716A392 (H424R) | 0.0034 |
| | CoTXSS + CoCYP716A392 (A285G)-CoTXSS + CoCYP716A393 | 0.0034 |
| | CoTXSS + CoCYP716A392 (A285G)-CoTXSS + CoCYP716A393 (A285G) | 0.0916 |
| | CoTXSS + CoCYP716A392 (A285G)-CoTXSS + CoCYP716A393 (A285V) | 0.0034 |
| | CoTXSS + CoCYP716A392 (A285G)-CoTXSS + CoCYP716A393 (A357L) | 0.0034 |
| | CoTXSS + CoCYP716A392 (A285G)-CoTXSS + CoCYP716A393 (H424R) | 0.0034 |

| | | |
|---|---|---|
| | CoTXSS + CoCYP716A392 (A285V)-CoTXSS + CoCYP716A392 (A357L) | 0.471 |
| | CoTXSS + CoCYP716A392 (A285V)-CoTXSS + CoCYP716A392 (H424R) | 0.6476 |
| | CoTXSS + CoCYP716A392 (A285V)-CoTXSS + CoCYP716A393 | 0.7255 |
| | CoTXSS + CoCYP716A392 (A285V)-CoTXSS + CoCYP716A393 (A285G) | 0.0034 |
| | CoTXSS + CoCYP716A392 (A285V)-CoTXSS + CoCYP716A393 (A285V) | 0.3869 |
| | CoTXSS + CoCYP716A392 (A285V)-CoTXSS + CoCYP716A393 (A357L) | 0.0034 |
| | CoTXSS + CoCYP716A392 (A285V)-CoTXSS + CoCYP716A393 (H424R) | 0.0034 |
| | CoTXSS + CoCYP716A392 (A357L)-CoTXSS + CoCYP716A392 (H424R) | 0.1815 |
| | CoTXSS + CoCYP716A392 (A357L)-CoTXSS + CoCYP716A393 | 0.3073 |
| | CoTXSS + CoCYP716A392 (A357L)-CoTXSS + CoCYP716A393 (A285G) | 0.0034 |
| | CoTXSS + CoCYP716A392 (A357L)-CoTXSS + CoCYP716A393 (A285V) | 0.241 |
| | CoTXSS + CoCYP716A392 (A357L)-CoTXSS + CoCYP716A393 (A357L) | 0.0034 |
| | CoTXSS + CoCYP716A392 (A357L)-CoTXSS + CoCYP716A393 (H424R) | 0.0034 |
| | CoTXSS + CoCYP716A392 (H424R)-CoTXSS + CoCYP716A393 | 1 |
| | CoTXSS + CoCYP716A392 (H424R)-CoTXSS + CoCYP716A393 (A285G) | 0.0034 |

| | | |
|---|---|---|
| | CoTXSS + CoCYP716A392 (H424R)-CoTXSS + CoCYP716A393 (A285V) | 0.8333 |
| | CoTXSS + CoCYP716A392 (H424R)-CoTXSS + CoCYP716A393 (A357L) | 0.0034 |
| | CoTXSS + CoCYP716A392 (H424R)-CoTXSS + CoCYP716A393 (H424R) | 0.0034 |
| | CoTXSS + CoCYP716A393-CoTXSS + CoCYP716A393 (A285G) | 0.0034 |
| | CoTXSS + CoCYP716A393-CoTXSS + CoCYP716A393 (A285V) | 0.471 |
| | CoTXSS + CoCYP716A393-CoTXSS + CoCYP716A393 (A357L) | 0.0034 |
| | CoTXSS + CoCYP716A393-CoTXSS + CoCYP716A393 (H424R) | 0.0034 |
| | CoTXSS + CoCYP716A393 (A285G)-CoTXSS + CoCYP716A393 (A285V) | 0.0034 |
| | CoTXSS + CoCYP716A393 (A285G)-CoTXSS + CoCYP716A393 (A357L) | 0.0034 |
| | CoTXSS + CoCYP716A393 (A285G)-CoTXSS + CoCYP716A393 (H424R) | 0.0034 |
| | CoTXSS + CoCYP716A393 (A285V)-CoTXSS + CoCYP716A393 (A357L) | 0.0034 |
| | CoTXSS + CoCYP716A393 (A285V)-CoTXSS + CoCYP716A393 (H424R) | 0.0034 |
| | CoTXSS + CoCYP716A393 (A357L)-CoTXSS + CoCYP716A393 (H424R) | 0.7255 |

**Supplementary Table 14.** Statistical tests used in Figure 6**.** *=P<0.05, ** P<0.01,*** P<0.0001

| Experiment | Sample | Z-value | p.adjusted (5 dp) |
|---|---|---|---|
| Psitaraxasterol Concentration (Kruskal Wallis (N=6) with Posthoc Dunns Test (Benjamini Hochberg Correction) | CoTXSS – CoTXSS +ACT1 | 2.204541 | 0.03298 (*) |
| | CoTXSS – CoTXSS +ACT2 | 2.204541 | 0.04123 (*) |
| | CoTXSS – CoTXSS +ACT3 | 4.409082 | 0.00006 (***) |
| | CoTXSS + ACT1 – CoTXSS +ACT2 | 0.000000 | 1.00000 |
| | CoTXSS + ACT1 – CoTXSS +ACT3 | 2.204541 | 0.05497 |
| | CoTXSS + ACT2 – CoTXSS +ACT3 | 2.204541 | 0.08246 |
| Taraxasterol Concentration (Kruskal Wallis (N=6) with Posthoc Dunns Test (Benjamini Hochberg Correction) | CoTXSS – CoTXSS +ACT1 | 2.1637159 | 0.06097 |
| | CoTXSS – CoTXSS +ACT2 | 1.3880442 | 0.19815 |
| | CoTXSS – CoTXSS +ACT3 | -0.7756718 | 0.43794 |
| | CoTXSS + ACT1 – CoTXSS +ACT2 | 4.1233077 | 0.00022 (***) |
| | CoTXSS + ACT1 – CoTXSS +ACT3 | 1.9595918 | 0.07051 |
| | CoTXSS + ACT2 – CoTXSS +ACT3 | 2.7352635 | 0.01870 (*) |
| Psitaraxasterol palmitate concentration (Kruskal Wallis (N=6) with Posthoc Dunns Test (Benjamini Hochberg Correction) | CoTXSS – CoTXSS +ACT1 | 1.8923960 | 0.07013 |
| | CoTXSS – CoTXSS +ACT2 | -0.6582247 | 0.05104 |
| | CoTXSS – CoTXSS +ACT3 | -2.5506207 | 0.03226 (*) |
| | CoTXSS + ACT1 – CoTXSS +ACT2 | -2.5506207 | 0.02156 (*) |
| | CoTXSS + ACT1 – CoTXSS +ACT3 | -4.4430167 | 0.00005 (***) |
| | CoTXSS + ACT2 – CoTXSS +ACT3 | -1.8923960 | 0.08766 |
| Psitaraxasterol concentration (Kruskal Wallis (N=6) with Posthoc Dunns Test (Benjamini Hochberg Correction) | CoTXSS+CYP716A382-CoTXSS+CYP716A382+ACT1 | -2.2861904 | 0.04449 (*) |
| | CoTXSS+CYP716A382-CoTXSS+CYP716A382+ACT2 | -0.1632993 | 0.87028 |
| | CoTXSS+CYP716A382-CoTXSS+CYP716A382+ACT3 | 2.1228911 | 0.05064 |
| | CoTXSS+CYP716A382+ACT1 - CoTXSS+CYP716A382+ACT2 | 2.1228911 | 0.04052 (*) |
| | CoTXSS+CYP716A382+ACT1 - CoTXSS+CYP716A382+ACT3 | 4.4090815 | 0.00062 (***) |
| | CoTXSS+CYP716A382+ACT2 - CoTXSS+CYP716A382+ACT3 | 2.2861904 | 0.06673 |
| Taraxasterol Concentration (Kruskal Wallis (N=6) with Posthoc Dunns Test (Benjamini Hochberg Correction) | CoTXSS+CYP716A382-CoTXSS+CYP716A382+ACT1 | -2.4903146 | 0.03829 (*) |
| | CoTXSS+CYP716A382-CoTXSS+CYP716A382+ACT2 | -0.4490731 | 0.65338 |
| | CoTXSS+CYP716A382-CoTXSS+CYP716A382+ACT3 | 1.7962925 | 0.08694 |
| | CoTXSS+CYP716A382+ACT1 - CoTXSS+CYP716A382+ACT2 | 2.0412415 | 0.06184 |

| | | | |
|---|---|---|---|
| | CoTXSS+CYP716A382+ACT1 - CoTXSS+CYP716A382+ACT3 | 4.2866070 | 0.00011 (***) |
| | CoTXSS+CYP716A382+ACT2 - CoTXSS+CYP716A382+ACT3 | 2.2453656 | 0.04949 (*) |
| Faradiol Concentration (Kruskal Wallis (N=6) with Posthoc Dunns Test (Benjamini Hochberg Correction) | CoTXSS+CYP716A382-CoTXSS+CYP716A382+ACT1 | 0.4082483 | 0.68309 |
| | CoTXSS+CYP716A382-CoTXSS+CYP716A382+ACT2 | 1.4288690 | 0.22956 |
| | CoTXSS+CYP716A382-CoTXSS+CYP716A382+ACT3 | 3.5517601 | 0.00230 (**) |
| | CoTXSS+CYP716A382+ACT1 - CoTXSS+CYP716A382+ACT2 | 1.0206207 | 0.36892 |
| | CoTXSS+CYP716A382+ACT1 - CoTXSS+CYP716A382+ACT3 | 3.1435118 | 0.00501 (**) |
| | CoTXSS+CYP716A382+ACT2 - CoTXSS+CYP716A382+ACT3 | 2.1228911 | 0.06753 |
| Psitaraxasterol Palmitate concentration (Kruskal Wallis (N=6) with Posthoc Dunns Test (Benjamini Hochberg Correction) | CoTXSS+CYP716A382-CoTXSS+CYP716A382+ACT1 | -1.239933 | 0.25800 |
| | CoTXSS+CYP716A382-CoTXSS+CYP716A382+ACT2 | -3.017170 | 0.00765 (**) |
| | CoTXSS+CYP716A382-CoTXSS+CYP716A382+ACT3 | -4.174440 | 0.00018 (***) |
| | CoTXSS+CYP716A382+ACT1 - CoTXSS+CYP716A382+ACT2 | -1.777237 | 0.11329 |
| | CoTXSS+CYP716A382+ACT1 - CoTXSS+CYP716A382+ACT3 | -2.934507 | 0.00668 (**) |
| | CoTXSS+CYP716A382+ACT2 - CoTXSS+CYP716A382+ACT3 | -1.157271 | 0.24716 |
| Experiment | Sample | T-value | P-value |
| Faradiol Palmitate concentration (One way ANOVA (N=6) with Posthoc Dunnetts test (CoTXSS+CYP716A382 as reference) | CoTXSS+CYP716A382-CoTXSS+CYP716A382+ACT1 | 2.792 | 0.0295 (*) |
| | CoTXSS+CYP716A382-CoTXSS+CYP716A382+ACT2 | 3.566 | 0.0052 (**) |
| | CoTXSS+CYP716A382-CoTXSS+CYP716A382+ACT3 | -1.642 | 0.2650 |

**Supplementary Table 15.** List of plasmids used in this manuscript.

| **Level 0 Phytobricks** | | | | | | |
|---|---|---|---|---|---|---|
| **Addgene#** | **Code** | **Part Type** | **Description** | **Cloning overhang (top strand)** | | **Source of plasmid** |
| | | | | **5'** | **3'** | |
| N/A | pL0-AstHMGR | CDS | AstHMGR (Avena strigosa truncated 3-hydroxy-3-methylglutaryl-coenzyme A reductase) | AATG | GCTT | A gift from A. Osbourn. Reed et al. (2017) doi: 10.1016/j.ymben.2017.06.012 |
| #50268 | pICH51277 | PROM +5UTR | 35Sshort_TMV (Cauliflower Mosaic Virus 35S promoter + Tobacco Mosaic Virus omega) | GGAG | AATG | Engler et al (2014) doi: 10.1021/sb4001504 |
| #50330 | pICH44022 | CDS | P19 (Tomato Busy Stunt Virus) | AATG | GCTT | Engler et al (2014) doi: 10.1021/sb4001504 |
| #162312 | pEPQD0CM0030 | STOP | stop codon | TTCG | GCTT | Dudley et al (2021) doi: 10.1093/synbio/ysab029 |
| #50337 | pICH41414 | 3UTR + TERM | Cauliflower Mosaic Virus 35S 3' untranslated region and terminator | GCTT | CGCT | Engler et al (2014) doi: 10.1021/sb4001504 |
| #227509 | pUAP41414 | 3UTR + TERM | Cauliflower Mosaic Virus 35S 3' untranslated region and terminator | GCTT | CGCT | This study |
| #227510 | pEPMS0CM0024 | CDS | CoTXSS (Calendula officinalis taraxasterol synthase) | AATG | TTCG | This study |
| #227511 | pEPDG1CB0027 | CDS | CaTXSS (Calendula arvensis taraxasterol synthase) | AATG | GCTT | This study |
| #227512 | pEPMS0CM0033 | CDS | CcTXSS (Cynara cardunculus taraxasterol synthase) | AATG | TTCG | This study |
| #227513 | pEPMS0CM0036 | CDS | HaTXSS (Helianthus annuus taraxasterol synthase) | AATG | TTCG | This study |
| #227514 | pEPMS0CM0034 | CDS | CeTXSS (Cichorium endivia | AATG | TTCG | This study |

| | | | | | | |
|---|---|---|---|---|---|---|
| | | | taraxasterol synthase) | | | |
| #227515 | pEPHS0CM0054 | CDS | TdTXSS (Tragopogon dubius taraxasterol synthase) | AATG | TTCG | This study |
| #227516 | pEPMS0CM0037 | CDS | LsTXSS (Lactuca sativa taraxasterol synthase) | AATG | TTCG | This study |
| #227517 | pEPMS0CM0031 | CDS | TkTXSS (Taraxacum kok-saghyz taraxasterol synthase) | AATG | TTCG | This study |
| #227518 | pEPMS0CM0032 | CDS | TcTXSS (Taraxacum coreanum mixed triterpene synthase) | AATG | TTCG | This study |
| #227519 | pEPMS0CM0026 | CDS | CoMAS (Calendula officinalis mixed-amyrin synthase) | AATG | TTCG | This study |
| #227520 | pEPMS0CM0038 | CDS | CYP716A392 (Calendula officinalis cytochrome P450 1) | AATG | TTCG | This study |
| #227521 | pEPMS0CM0039 | CDS | CYP716A393 (Calendula officinalis cytochrome P450 2) | AATG | TTCG | This study |
| #227522 | pEPMS0CM0040 | CDS | CYP716A429 (Calendula officinalis cytochrome P450 3) | AATG | TTCG | This study |
| #227523 | pEPMS0CM0041 | CDS | CYP716A430 (Calendula officinalis cytochrome P450 4) | AATG | TTCG | This study |
| #227524 | pEPMS0CM0042 | CDS | CYP716A431 (Calendula officinalis cytochrome P450 5) | AATG | TTCG | This study |
| #227525 | pEPDG1CB0028 | CDS | CYP716A392a (Calendula arvensis cytochrome P450 1) | AATG | TTCG | This study |

| | | | | | | |
|---|---|---|---|---|---|---|
| #227526 | pEPDG1CB0024 | CDS | CoACT1 (Calendula officinalis acyltransferase 1) | AATG | TTCG | This study |
| #227527 | pEPDG1CB0025 | CDS | CoACT2 (Calendula officinalis acyltransferase 2) | AATG | TTCG | This study |
| #227528 | pEPDG1CB0026 | CDS | CoACT3 (Calendula officinalis acyltransferase 3) | AATG | TTCG | This study |
| #227529 | pEPCT0CM0207 | CDS | CoACT4 (Calendula officinalis acyltransferase 4) | AATG | TTCG | This study |
| #227530 | pEPCT0CM0208 | CDS | CoACT5 (Calendula officinalis acyltransferase 5) | AATG | TTCG | This study |
| #227531 | pEPCT0CM0209 | CDS | CoACT6 (Calendula officinalis acyltransferase 6) | AATG | TTCG | This study |
| #227532 | pEPCT0CM0210 | CDS | CoACT7 (Calendula officinalis acyltransferase 7) | AATG | TTCG | This study |

| **Level 1 Plasmids** | | | | | | | | |
|---|---|---|---|---|---|---|---|---|
| **Addgene#** | **Code** | **Acceptor (GGAG-CGCT)** | **Description** | **Promoter+ 5'UTR (GGAG-AATG)** | **CDS AATG-TTCG or AATG-GCTT*** | **STOP (TTCG-GCTT)** | **3'UTR + TERM (GCTT-CGCT)** | **Source of plasmid** |
| #48000 | pICH47732 | pICH47732 | MoClo Acceptor Plasmid L1 Position 1 forward | none | none | none | none | Weber et al (2011) doi: 10.1371/journal.pone.0016765 |
| #48001 | pICH47742 | pICH47742 | MoClo Acceptor Plasmid L1 Position 2 forward | none | none | none | none | Weber et al (2011) doi: 10.1371/journal.pone.0016765 |
| #177038 | pEPQD1CB0104 | pICH47781 | 35Sshort_TMV_P19_35S | pICH51277 | pICH44022 | N/A | pICH41414 | Dudley et al (2022) doi: 10.1038/s42003-022-03904-w. |
| #177039 | pEPQD1CB0817 | pICH47732 | 35Sshort_TMV_AstHMGR_35S | pICH51277 | pL0-AstHMGR | N/A | pICH41414 | Dudley et al (2022) doi: 10.1038/s42003-022-03904-w. |
| #227533 | pEPMS1CB0001 | pICH47732 | 35Sshort_TMV_CoTXSS_35S | pICH51277 | pEPMS0CM0024 | pEPQD0CM0030 | pUAP41414 | This study |
| #227534 | pEPMS1CB0010 | pICH47732 | 35Sshort_TMV_CcTXSS_35S | pICH51277 | pEPMS0CM0033 | pEPQD0CM0030 | pUAP41414 | This study |
| #227535 | pEPMS1CB0013 | pICH47732 | 35Sshort_TMV_HaTXSS_35S | pICH51277 | pEPMS0CM0036 | pEPQD0CM0030 | pUAP41414 | This study |
| #227536 | pEPMS1CB0011 | pICH47732 | 35Sshort_TMV_CeTXSS_35S | pICH51277 | pEPMS0CM0034 | pEPQD0CM0030 | pUAP41414 | This study |

| | | | | | | | | |
|---|---|---|---|---|---|---|---|---|
| #227537 | pEPHS1CB0028 | pICH47732 | 35Sshort_TMV_TdTXSS_35S | pICH51277 | pEPHS0CM0054 | pEPQD0CM0030 | pUAP41414 | This study |
| #227538 | pEPMS1CB0014 | pICH47732 | 35Sshort_TMV_LsTXSS_35S | pICH51277 | pEPMS0CM0037 | pEPQD0CM0030 | pUAP41414 | This study |
| #227539 | pEPMS1CB0008 | pICH47732 | 35Sshort_TMV_TkTXSS _35S | pICH51277 | pEPMS0CM0031 | pEPQD0CM0030 | pUAP41414 | This study |
| #227540 | pEPMS1CB0009 | pICH47732 | 35Sshort_TMV_TcTXSS_35S | pICH51277 | pEPMS0CM0032 | pEPQD0CM0030 | pUAP41414 | This study |
| #227541 | pEPMS1CB0003 | pICH47732 | 35Sshort_TMV_CoMAS_35S | pICH51277 | pEPMS0CM0026 | pEPQD0CM0030 | pUAP41414 | This study |
| #227542 | pEPDG1CB0004 | pICH47732 | 35Sshort_TMV_CaTXSS_35S | pICH51277 | pEPDG1CB0027 | pEPQD0CM0030 | pUAP41414 | This study |
| #227543 | pEPMS1CB0018 | pICH47742 | 35Sshort_TMV_CYP716A392_35S | pICH51277 | pEPMS0CM0038 | pEPQD0CM0030 | pUAP41414 | This study |
| #227544 | pEPMS1CB0019 | pICH47742 | 35Sshort_TMV_CYP716A393_35S | pICH51277 | pEPMS0CM0039 | pEPQD0CM0030 | pUAP41414 | This study |
| #227545 | pEPMS1CB0020 | pICH47742 | 35Sshort_TMV_CYP716A429_35S | pICH51277 | pEPMS0CM0040 | pEPQD0CM0030 | pUAP41414 | This study |
| #227546 | pEPMS1CB0021 | pICH47742 | 35Sshort_TMV_CYP716A430_35S | pICH51277 | pEPMS0CM0041 | pEPQD0CM0030 | pUAP41414 | This study |
| #227547 | pEPMS1CB0022 | pICH47742 | 35Sshort_TMV_CYP716A431_35S | pICH51277 | pEPMS0CM0042 | pEPQD0CM0030 | pUAP41414 | This study |
| #227548 | pEPDG1CB0005 | pICH47742 | 35Sshort_TMV_CYP716A392a_35S | pICH51277 | pEPDG1CB0028 | pEPQD0CM0030 | pUAP41414 | This study |
| #227549 | pEPDG1CB0001 | pICH47732 | 35Sshort_TMV_CoACT1_35S | pICH51277 | pEPDG1CB0024 | pEPQD0CM0030 | pUAP41414 | This study |
| #227550 | pEPDG1CB0002 | pICH47732 | 35Sshort_TMV_CoACT2_35S | pICH51277 | pEPDG1CB0025 | pEPQD0CM0030 | pUAP41414 | This study |
| #227551 | pEPDG1CB0003 | pICH47732 | 35Sshort_TMV_CoACT3_35S | pICH51277 | pEPDG1CB0026 | pEPQD0CM0030 | pUAP41414 | This study |
| #227552 | pEPCT1CB0066 | pICH47751 | 35Sshort_TMV_CoACT4_35S | pICH51277 | pEPCT0CM0207 | pEPQD0CM0030 | pUAP41414 | This study |
| #227553 | pEPCT1CB0067 | pICH47751 | 35Sshort_TMV_CoACT5_35S | pICH51277 | pEPCT0CM0208 | pEPQD0CM0030 | pUAP41414 | This study |
| #227554 | pEPCT1CB0068 | pICH47751 | 35Sshort_TMV_CoACT6_35S | pICH51277 | pEPCT0CM0209 | pEPQD0CM0030 | pUAP41414 | This study |
| #227555 | pEPCT1CB0069 | pICH47751 | 35Sshort_TMV_CoACT7_35S | pICH51277 | pEPCT0CM0210 | pEPQD0CM0030 | pUAP41414 | This study |

#### Mutated Level 1 Plasmids

| Addgene# | Code | Acceptor (GGAG-CGCT) | Description | WT plasmid | Mutation | Source of plasmid |
|---|---|---|---|---|---|---|
| #227556 | pEPHS1CB0029 | pICH47732 | 35Sshort_TMV_CoMAS (I367M)_35S | pEPMS1CB0003 | I367M | This study |
| #227557 | pEPHS1CB0030 | pICH47732 | 35Sshort_TMV_CoMAS (E371D)_35S | pEPMS1CB0003 | E371D | This study |
| #227558 | pEPHS1CB0031 | pICH47732 | 35Sshort_TMV_CoMAS (I367M/E371D)_35S | pEPMS1CB0003 | I367M / E371D | This study |
| #227559 | pEPHS1CB0001 | pICH47732 | 35Sshort_TMV_CoTXSS (G380T)_35S | pEPMS1CB0001 | G380T | This study |
| #227560 | pEPHS1CB0002 | pICH47732 | 35Sshort_TMV_CoTXSS (D385E)_35S | pEPMS1CB0001 | D385E | This study |
| #227561 | pEPHS1CB0003 | pICH47732 | 35Sshort_TMV_CoTXSS (H492Q)_35S | pEPMS1CB0001 | H492Q | This study |
| #227562 | pEPHS1CB0004 | pICH47732 | 35Sshort_TMV_CoTXSS (P751A)_35S | pEPMS1CB0001 | P751A | This study |
| #227563 | pEPHS1CB0017 | pICH47732 | 35Sshort_TMV_CoTXSS (D385E/H492Q)_35S | pEPMS1CB0001 | D385E / H492Q | This study |
| #227564 | pEPHS1CB0020 | pICH47732 | 35Sshort_TMV_CoTXSS (G380T/D385E/H492Q/P751A)_35S | pEPMS1CB0001 | G380T / D385E / | This study |

| | | | | | | |
|---|---|---|---|---|---|---|
| | | | | | H492Q / P751A | |
| #227565 | pEPHS1CB0021 | pICH47732 | 35Sshort_TMV_TkTXSS (T374G)_35S | pEPMS1CB0008 | T374G | This study |
| #227566 | pEPHS1CB0010 | pICH47732 | 35Sshort_TMV_TkTXSS (E379D)_35S | pEPMS1CB0008 | E379D | This study |
| #227567 | pEPHS1CB0011 | pICH47732 | 35Sshort_TMV_TkTXSS (Q486H)_35S | pEPMS1CB0008 | Q486H | This study |
| #227568 | pEPHS1CB0012 | pICH47732 | 35Sshort_TMV_TkTXSS (A745P)_35S | pEPMS1CB0008 | A745P | This study |
| #227569 | pEPHS1CB0022 | pICH47732 | 35Sshort_TMV_TkTXSS (E379D/Q486H)_35S | pEPMS1CB0008 | E379D / Q486H | This study |
| #227570 | pEPHS1CB0027 | pICH47732 | 35Sshort_TMV_TkTXSS (T374G/E379D/Q486H/A745P)_35S | pEPMS1CB0008 | T374G / E379D / Q486H / A745P | This study |
| #227571 | pEPDG1CB0007 | pICH47742 | 35Sshort_TMV_CYP716A392 (A285G)_35S | pEPMS1CB0018 | A285G | This study |
| #227572 | pEPDG1CB0009 | pICH47742 | 35Sshort_TMV_CYP716A392 (A357L)_35S | pEPMS1CB0018 | A357L | This study |
| #227573 | pEPDG1CB0010 | pICH47742 | 35Sshort_TMV_CYP716A392 (H424R)_35S | pEPMS1CB0018 | H424R | This study |
| #227574 | pEPDG1CB0018 | pICH47742 | 35Sshort_TMV_CYP716A392 (A285V)_35S | pEPMS1CB0018 | A285V | This study |
| #227575 | pEPDG1CB0012 | pICH47742 | 35Sshort_TMV_CYP716A393 (A285G)_35S | pEPMS1CB0019 | A285G | This study |
| #227576 | pEPDG1CB0014 | pICH47742 | 35Sshort_TMV_CYP716A393 (A357L)_35S | pEPMS1CB0019 | A357L | This study |
| #227577 | pEPDG1CB0015 | pICH47742 | 35Sshort_TMV_CYP716A393 (H424R)_35S | pEPMS1CB0019 | H424R | This study |
| #227578 | pEPDG1CB0021 | pICH47742 | 35Sshort_TMV_ CYP716A393 (A285V)_35S | pEPMS1CB0019 | A285V | This study |

**Supplementary Table 16.** List of primers.

| Primers for mutagenesis of TXSSs | | | |
|---|---|---|---|
| Gene | Mutation | Forward Primer (5' - 3') | Reverse Primer (5' - 3') |
| *CoTXSS* | G380T | TACATAACCATGGGTTGTG TTGACAAGGCTTTAC | ACCCATGGTTATGTAACG TCCTTCTTCGGATCC |
| *CoTXSS* | D385E | GTGTTGAAAAGGCTTTACA AATGATGTGTTTTTATGCCG | AGCCTTTTCAACACAACC CATGCCTATGTAACGTC |
| *CoTXSS* | H492Q | CAAGACCAAGGATGGGTTG TATCAGATTGCACTGCAG | CCATCCTTGGTCTTGATC AGAGAAAGTCCATGCCC |
| *CoTXSS* | P751A | TTACATTATGCAGAATATAGG AACACTTTTCCGTTATGGGC | CCTATATTCTGCATAATGT AACATGCAGTTTTTCATGTACACTC |
| *CoTXSS* | G380T | TACATAACCATGGGTTGTGTT GAAAAGGCTTTAC | ACCCATGGTTATGTAACG TCCTTCTTCGGATCC |
| *CoTXSS* | D385E | GTGTTGAAAAGGCTTTACAAA TGATGTGTTTTTATGCCG | AGCCTTTTCAACACAAC CCATGGTTATGTAACGTC |
| *TkTXSS* | T374G | ATACATAGGATTGGGGTGTGT CGAGAAATCCTTACAAATG | ACCCCAATCCTATGTATC TACCTTCTTCGGCGTTATATTG |
| *TkTXSS* | E379D | TGTGTCGATAAATCCTTACAAA TGATGTGCTTCTCAGC | AGGATTTATCGACACACC CCAATGTTATGTATCTACCTTC |
| *TkTXSS* | Q486H | AGGACCATGGCTGGGTTGTTA GTGATTGCACTG | CAGCCATGGTCCTGGTC GCTAAATGTCCAAGC |
| *TkTXSS* | A745P | CACTACCCCGAATATCGAAAC ACCTTCCCTTTATGG | ATATTCGGGGTAGTGCAGC ATGCAATTCTTCATATAAACC |
| *TkTXSS* | T374G | ATACATAGGATTGGGGTGTGTC GATAAATCCTTACAAATG | ACCCCAATCCTATGTATCTA CCTTCTTCGGCGTTATATTG |
| *TkTXSS* | E379D | TGTGTCGATAAATCCTTACAAA TGATGTGCTTCTCAGC | AGGATTTATCGACACACCC CAATCCTATGTATCTACCTTC |
| *CoMAS* | I367M | GACTATGGGATGTGTTGAAAA GAGCTTGCAAATGATGTGTTGG | TCTTTTCAACACATCCCATA GTCATATATCTGCTTTGTTGAGCAC |
| *CoMAS* | E371D | GACTATTGGATGTGTTGATAA GAGCTTGCAAATGATGTGTTGG | TCTTATCAACACATCCAATA GTCATATATCTGCTTTGTTGAGCAC |
| *CoMAS* | I367M E371D | GACTATGGGATGTGTTGATAA GAGCTTGCAAATGATGTGTTGG | TCTTATCAACACATCCCATA GTCATATATCTGCTTTGTTGAGCAC |
| **Primers for mutagenesis of CYPs** | | | |
| **Gene** | **Mutation** | **Forward Primer (5' - 3')** | **Reverse Primer (5' - 3')** |
| *CoCYP1* | A285G | AGATTCTTGGTTTGTTGAT CGGTGGGCATGAC | AACAAACCAAGAATCTTGC CCGAAATGTCGTG |
| *CoCYP1* | A356L | ACCGCTTCAAGGTGCTTTT AGAGAAGCCC | CTTGAAGCGGTGGGACTAA TCTAAGAACTTCAC |
| *CoCYP1* | H423R | CCCGAGAATGTGTCCCGGAA AAGAGTACG | CACATTCTCGGGCCTCCTC CAAATGGCAC |

| | | | |
|---|---|---|---|
| *CoCYP1* | A284V | AGATTCTTGTGTTGTTGATC GGTGGGCATGAC | CAACAACACAAGAATCTTG CCCGAAATGTCGTG |
| *CoCYP2* | A285G | AGATTCTTGTGTTGTTGATC GGTGGGCATGAC | CAACAACACAAGAATCTTG CCCGAAATGTCGTG |
| *CoCYP2* | A356L | ACCGCTTCAAGGTGCTTTT CGAGAAGCC | CTTGAAGCGGTGGGACTAA TCTAAGAACTTCAC |
| *CoCYP2* | H423R | CCCGAGAATGTGTCCAGGA AAAGAGTATGCC | CACATTCTCGGGCCTCCTC CAAATGGCAC |
| *CoCYP2* | A284V | AGATACTTGTGTTGCTGAT TGGTGGGCATGAC | AGCAACACAAGTATCTTGC CAGAAATGTCGTGTTC |

**Primers for qRT-PCR**

| Gene | Forward Primer (5' - 3') | Reverse Primer (5' - 3') | Amplicon length (bp) |
|---|---|---|---|
| *SAND.2* | TCTTTCAGTTGGAACCCTGCA | CTGCAATATAGCACCAGCAGC | 93 |
| *TXSS* | GGTGACTTGCTCATGCGAAC | TTACCGCCATTGTCACAGCT | 121 |
| *CoCYP1* | TGGCCCATAATCGGGGAAAG | CACACATCACTGCAGCATCC | 145 |
| *CoCYP2* | TTAGCGACGAAGATGGCGAG | CACCAATCAGCAACGCAAGT | 72 |
| *CoACT1* | CGTTTCAAGAGTACGAGGCG | TTTTGCGGCCGAGTAAAACT | 105 |
| *CoACT2* | GTAAAGCCTTCACCCGTTGG | GTGCCCCACATTCATTCGTT | 84 |

**Biosynthesis and Bioactivity of Anti-Inflammatory Triterpenoids in *Calendula officinalis* (pot marigold)**

Golubova D, Salmon M, Su H, Tansley C, Kaithakottil GG, Linsmith G, Schudoma S, Swarbreck D, O'Connell MA, Patron NJ

### Supplementary Methods

1. **Library construction and sequencing**
2. **Genome Assembly**
3. **Functional annotation**

#### 1. Library construction and sequencing

*PacBio HiFi*

A low input library was prepared from a starting input of 984 ng gDNA. The sample was sheared using the Megaruptor 3 instrument (Diagenode, P/N B06010003), 5 ng/µl at a speed setting of 30. The sample underwent AMPure® PB bead (PacBio®, P/N 100-265-900) purification and concentration at a ratio of 1.8X before library construction using the SMRTbell template prep kit 2.0 (PacBio®, P/N 100-983-900).

The HiFi library was prepared from 767 ng of sheared gDNA, according to the instructions in the low input protocol version 05 (PacBio®, P/N 101-730-400). A barcoded adapter (PacBio ®, P/N 101-628-500) was ligated to create the SMRTbell library. The final SMRTbell library was purified with AMPure beads. The final library size was estimated from a smear analysis performed on the FEMTO Pulse® System (Agilent, P/N M5330AA) and quantified by fluorescence (Invitrogen Qubit™ 3.0, P/N Q33216).

Loading calculations for sequencing were made using the PacBio® SMRT®Link Binding Calculator v9.0.0.92017. Sequencing primer v4 was annealed to the adapter sequence of the library. Binding of the library to the sequencing polymerase was completed using Sequel® II Binding Kit v2.0 (PacBio®, P/N 101-842-900). The Sequel® II DNA internal control was spiked into the library complex at the standard concentration prior to sequencing. The sequencing chemistry used was Sequel® II Sequencing Plate 2.0 (PacBio®, P/N 101-820-200) and the Instrument Control Software v9.0.0.92233.

The library was sequenced on four Sequel II SMRT®cells 8M on the Sequel II instrument. The parameters for sequencing per cell were as follows: diffusion loading, CLR sequencing mode, 30-hour movie, 2-hour immobilisation time, 2-hour pre-extension time, 20 pM on plate loading concentration.

A total of 1284 Gb of PacBio data was produced from four SMRT cells. Circular consensus sequencing (CCS) analysis (SMRT Link 9.0.0.92188) yielded 6,098,205 CCS reads with a median length of 13.0 kb at an estimated accuracy greater than Q20 (or 99.9% accuracy). The Q20 CCS reads comprised a total 81Gb of Hifi data (or 58X coverage based on an estimated genome size of 1.4Gb). HiFI reads were processed with CutAdapt v3.2 (http://code.google.com/p/cutadapt/) [--action lowercase -O 20 --errors 8 -b ATCTCTCTCTTTTCCTCCTCCTCCGTTGTTGTTGTTGAGAGAGAT -rc] to remove reads containing SMRT adapter sequences.

*10X Genomics linked reads*

Genomic DNA was quantified using the Qubit dsDNA HS (High Sensitivity) Assay Kit (Q32854, Thermo Fisher Scientific). DNA integrity was checked using the FEMTO Pulse System Genomic DNA 165 kb Kit (FP-1002-0275, Agilent Technologies), which determined that 64.4% of the DNA was greater than 50 kb.The Chromium 10x platform (10x Genomics) micro-fluidic Genome Chip (PN-120216) was used to produce three barcoded linked read libraries, with 1.14ng DNA input, using the Chromium™ Genome Library Kit & Gel bead Kit v2 (120258) following the Chromium Genome Reagent Kits Version 2 User Guide (CG00043). The micro-fluidic chip produces a library of genome gel beads by combining the DNA with the gel beads, reaction mastermix and partitioning oil to create Gel Bead in Emulsions (GEMs). Isothermal incubation of the GEMs was followed by recovery of the barcoded fragments which were then used to create libraries for Illumina sequencing. Library yields were quantified using the Qubit dsDNA HS Assay; insert size was determined with Agilent 2100 Bioanalyzer High Sensitivity DNA chip (5067-4627, Agilent Technologies), and finally checked by qPCR (07960204001, Roche Diagnostics Ltd.) prior to equimolar pooling and sequencing.

*Omni-C*

A sample of fresh leaf was provided to Dovetail Genomics (Scotts Valley, CA, USA).

*Illumina RNA-Seq*

Libraries were constructed using the NEBNext Ultra II RNA Library prep for Illumina kit (NEB#E7760L), NEBNext Poly(A) mRNA Magnetic Isolation Module (NEB#E7490L) and NEBNext Multiplex Oligos for Illumina® (96 Unique Dual Index Primer Pairs) (E6440S/L) at a concentration of 10 µM. Library preparation was performed on the Perkin Elmer (formerly Caliper LS) Sciclone G3 (PerkinElmer PN: CLS145321). mRNA was purified from 1 µg of total RNA using a Poly(A) mRNA Magnetic Isolation Module. Isolated mRNA was fragmented for 12 minutes at 94°C, and first strand cDNA was synthesised. Directionality was retained by adding dUTP during second strand synthesis and subsequent cleavage of the uridine containing strand using USER Enzyme. NEBNext Adaptors were ligated to end-repaired, dA-tailed DNA. The ligated products were purified using Beckman Coulter AMPure XP beads (A63882). Adaptor-ligated DNA was enriched in ten cycles of PCR (30 secs at 98°C, 10 cycles of: 10 secs at 98°C _75 secs at 65°C _5 mins at 65°C). Barcodes (NEBNext Multiplex Oligos for Illumina®) were incorporated during PCR. Library quality was determined using a Perkin Elmer DNA High Sensitivity Reagent Kit (CLS760672) with DNA 1K / 12K / HiSensitivity Assay LabChip (760517) and the concentration measured with a Quant-iT™ dsDNA Assay Kit, high sensitivity (Plate Reader) assay from ThermoFisher (Q-33120). The final libraries were equimolar pooled, verified by qPCR and sequenced on two lanes of a NovaSeq 6000 SP flow cell with 150 bp PE reads, yielding ~100 Gb.

*PacBio IsoSeq*

Libraries were constructed from 300 ng of total RNA. Reverse transcription cDNA synthesis was performed using NEBNext® Single Cell/Low Input cDNA Synthesis & Amplification Module (NEB, E6421). cDNA samples were amplified with barcoded primers for 12 cycles and pooled at equimolar concentration before SMRTbell library construction. The library pool was prepared according to the Iso-Seq protocol version 02 (PacBio, 101-763-800), using SMRTbell express template prep kit 2.0 (PacBio, 102-088-900). The library pool was quantified using a Qubit Fluorometer 3.0 (Invitrogen) and sized using the Bioanalyzer HS DNA chip (Agilent Technologies, Inc.).

The loading calculations were performed using the PacBio SMRTlink Binding Calculator v8.0.0.78409. Sequencing primer v2 was annealed to the Iso-Seq library pool and complexed to the sequencing polymerase with the Sequel II binding kit V2.1 (PacBio, 101-843-000). The sequencing internal control complex 1.0 (PacBio, 101-717-600) was spiked at a standard concentration before sequencing. The Iso-Seq pool was sequenced on the Sequel II instrument with a single Sequel II SMRT®cell 8M cell using Sequel® II Sequencing Plate 2.0 (PacBio®, 101-820-200) and the Instrument Control Software v8.0.0.78867. The parameters for sequencing were diffusion loading, 30 hr movie, 2 hr immobilisation time, 2 hr pre-extension time, 40 pM loading concentration.

PacBio subreads (single SMRT cell) were analysed via the smrtlink-8.0.0.80529 pb_demux_isoseq3 application (default parameters). Processing generated Circular Consensus Sequences (CCS), which were demultiplexed, primers removed, concatemers identified, polyA tails removed and clustered into high quality (predicted accuracy ≥ 0.99) and low quality (predicted accuracy < 0.99) transcripts. In total, 108,882, 79,287 and 94,014 high- and low-quality transcripts were output for three libraries, CoDisc (*Calendula officinalis* Disc), CoLeaf (*C. officinalis* Leaf) and CoRay (*C. officinalis* Ray) and 105,790 high- and low-quality transcripts from *Calendula arvensis.*

#### 2. Genome Assembly

Six assemblies were generated from the HiFI data using four alternative assembly tools Hicanu v2.0 (Nurk et al. 2020), Hifiasm v0.12 (Cheng et al. 2021), WTDBG v2.5 (Ruan and Li 2020) and IPA 1.3.1 (https://github.com/PacificBiosciences/pbipa). The “best” assembly was selected based on contiguity, kmer completeness / coherence against the content of the HiFi data and an assessment of potential misassembles using 10X linked-read information (Jackman et al, 2018). The hifiasm L2 assembly was provided to Dovetail genomics for scaffolding with Omni-C data and the HiRise scaffolding algorithm (see below).

*Repeat identification*

RepeatModeler (v1.0.11 - http://www.repeatmasker.org/RepeatModeler/) was used for *de novo* identification of repetitive elements from the assembled *C. officinalis* genome. High copy protein-coding genes potentially included in the RepeatModeler library were identified and effectively removed by running RepeatMasker v4.0.72 using the *Lactuca sativa* coding genes to hard mask the RepeatModeler library; transposable element genes were first excluded from the *Lactuca sativa* coding-geneset by running TransposonPSI (r08222010). Unclassified repeats were searched in a custom BLAST database of organellar genomes (mitochondrial and chloroplast sequences from eudicotyledons in the NCBI nucleotide division). Any repeat families matching organellar DNA were also hard-masked.

Repeat identification was completed by running RepeatMasker v4.0.72 with a RepBase Viridiplantae library and with the customized RepeatModeler library (i.e. after masking out protein coding genes), both using the -nolow setting.

*Genome quality control and evaluation*

An inspection of the kmer spectrum of HiFi reads was performed using the Kmer Anlaysis Toolkit (KAT) v2.4.2 (Mapleson et al. 2017). A kmer frequency histogram of the full set of Hifi data was produced at a kmer size of 22 leaving all other parameters at default settings. The kmer spectrum shows three clear peaks, consistent with *C. officinalis* having an allotetraploid background with the third peak representing content that is shared by both subgenomes (**Figure 1**). The first peak, representing heterozygous kmers, is small in comparison to the main peak, indicating that Marigold displays a moderate level of heterozygosity.

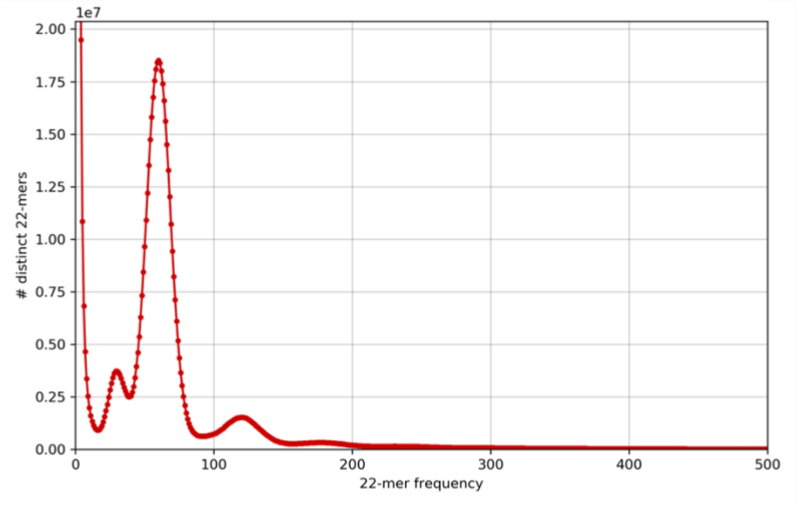

**Figure 1.** Kmer frequency histogram of Hifi data (k = 22).

Alternative draft genome assemblies were generated using 4 tools, Hicanu v2.0 (Nurk et al. 2020), Hifiasm v0.12 (Cheng et al. 2021), WTDBG v2.5 (Ruan and Li 2020) and IPA 1.3.1 (https://github.com/PacificBiosciences/pbipa). For Hifiasm we evaluated alternative parameter options for the merge level (-l parameter) controlling the aggressiveness of haplotype collapsing by hifiasm's internal implementation of Purge Dupes, and the –high-het setting (in beta in version 0.12). Hifiasm and IPA both produce a haplotype collapsed 'primary' assembly representing a single allele per genomic locus and an additional set of 'alternate' haplotigs. Hicanu and WTDBG produce a single haplotype expanded assembly. The Hicanu and WTDBG assemblies were therefore further processed with the Purge Dupes software v0.0.3 (Guan et al. 2020) to produce a primary assembly and associated set of 'alternate' contigs. Purge Dupes was run with default settings to auto detect the coverage threshold at which to collapse haplotigs.

Assemblies produced by the above tools were assessed using a range of assembly QC metrics. The full set of Hifi reads were aligned back to all assemblies using minimap2 (version 2.11) with parameters [-ax asm20]. Read alignments were passed to the software package Asset v1.0.0, to compute read alignment breakpoints, following the protocol described on the Asset github page (https://github.com/dfguan/asset). The count of breakpoints was used as a QC metric. Consensus quality of assemblies was summarised by QV value, computed by comparison of the kmer content of the assemblies to the kmer content of the Hifi reads using the software YAK v0.1 (https://github.com/lh3/yak) (**Table 1**). BUSCO scores were produced for each assembly using the BUSCO pipeline v3.0 (Simão et al. 2015) run with default settings.

**Table 1. Genome assembly QC summary**

| Assembly | Size | N50 | QV | Breakpoints | BUSCO Complete |
|---|---|---|---|---|---|
| **Hifiasm L=1** | 1.3 Gb | 36.6 Mb | 51.5 | 7,245 | 91.7% |
| **Hifiasm L=2** | 1.3 Gb | 36.6 Mb | 51.5 | 7,096 | 91.9% |
| **Hifiasm L=2, Hihet = True** | 1.3 Gb | 27.8 Mb | 52.3 | 6,948 | 91.8% |
| **Hicanu** | 1.1 Gb | 18.75 Mb | 40.4 | 12,657 | 91.2% |
| **IPA** | 1.1 Gb | 6.9 Mb | 42.6 | 1,818 | 91.9% |
| **WTDBG2** | 928 Mb | 258 kb | 38.7 | 14,876 | 89.8% |

To examine assembly coherence against the content of the reads, we used the Kmer Analysis Toolkit (KAT) v2.4.2 (Mapleson et al. 2017) comp function to generate spectra-cn plots (**Figure 2**).

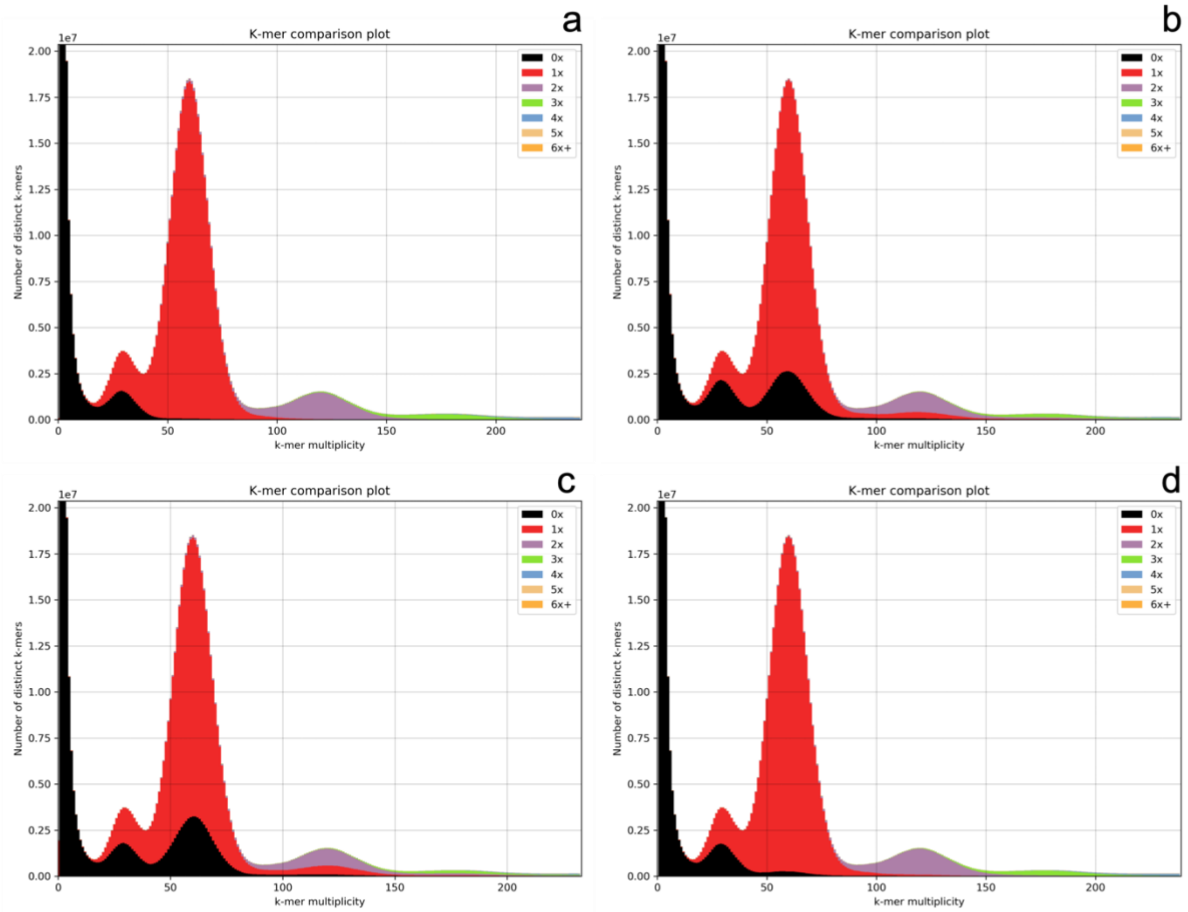

**Figure 2. Spectra-cn plots of alternative HiFi assemblies, a) Hifiasm L=2, b) Hicanu, c) WTDBG2 and d) IPA**

The spectra-cn plots for the Hifiasm assemblies (only L=2 shown) fit with expectations. The first peak of heterozygous kmers is coloured roughly half red and half black, indicating half of the heterozygous kmers present in the reads are present in the assembly (**Figure 2**). This is as expected for a haplotype mosaic assembly, where one allele is represented at each heterozygous locus. The second peak of single copy genomic content is coloured red, indicating that the content is represented once in the assembly. Lastly the third peak of content shared between both sub genomes is coloured purple indicating that all this content is present twice in the assembly. Once again, this is concomitant with a veridical assembly of the genome. Kmer spectrum analysis shows a significant amount of missing content from the single copy portion of the kmer spectrum in both the WTDBG and Hicanu assemblies, indicating that these assemblies do not completely represent the unique content in the genome. No further analysis was conducted on the WTDBG and Hicanu assemblies.

The hifiasm assemblies are more contiguous than the IPA assembly but have more read alignment breakpoints. 10X genomics linked reads were used to assess Hifiasm and IPA assemblies. Linked read data was pre-processed using the Longranger Basic pipeline (https://github.com/10XGenomics/longranger) to trim reads and prepare fastq headers with barcode information. Pre-processed linked read data was passed to the Tigmint pipeline (Jackman et al., 2018) with default settings. Tigmint aligns reads to the assembly using longranger, infers molecule extents, and identifies misassembles using linked-read information. The number of breaks to contigs in different size fractions was used as a QC metric (**Table 2**).

**Table 2. Summary of 10X contig breaking**

| | IPA | Hifiasm L=1 | Hifiasm L=2 | Hifiasm L=2 Hihet = True |
|---|---|---|---|---|
| **Total Breaks** | 342 | 2099 | 1971 | 2016 |
| **Broken Contigs** | 132 | 474 | 463 | 477 |
| **Total Length Broken Contigs** | 104.8 Mb | 110.3 Mb | 70.9 Mb | 200.9 Mb |
| **Broken contigs > 1Mb** | 20 | 3 | 2 | 6 |
| **Broken contigs 100kb – 1Mb** | 9 | 30 | 27 | 29 |
| **Broken contigs 10-100kb** | 94 | 441 | 434 | 442 |
| **Broken contigs 1-10kb** | 9 | 0 | 0 | 0 |

Analysis of the Tigmint assembly breaking indicates that the IPA assembly contains more potential misassemblies in contigs over 1Mb based on alignment of 10X data (**Table 2**). Based on this powerful orthogonal data set and the contiguity and kmer analysis, it was decided to accept the Tigmint broken, hifiasm L2 assembly as the final contig assembly.

Breaks were introduced at positions where barcode coverage (i.e. the number of inferred DNA molecules covering each position in the assembly) dropped below 20X. These breaks were mostly introduced at the ends of long contigs.

The final contig assembly was provided to Dovetail Genomics for scaffolding with Omni-C data and the HiRise scaffolding algorithm (Putnam et al. 2016). The Hirise software made a total of 5 breaks and 33 joins to the input contigs. Dovetail Genomics made a total of 7 manual joins to the assembly. The linked read data again provides an orthogonal data set with which to validate the scaffolding process. The broken assembly was independently scaffolding using the 10X data and the ARCS algorithm (Yeo et al. 2018) and the results were compared with the results obtained from Hirise. Of the 33 joins made by Hirise, 20 were supported by 10X data. The ARCS software makes a total of 23 joins based on 10X data, i.e., an additional three which were not made by Hirise. Of these three additional 10X based joins, two supported manual edits made by Dovetail genomics while the final one would have linked two chromosome scale scaffolds. Since this last join is biologically implausible it was not incorporated into the final assembly.

The final contig set was screened for contaminants using Kraken2 v2.0.7 (Wood, Lu, and Langmead 2019) and an index built from the Refseq genomes database. Kraken2 was. Run using default settings. In addition, proteins from *Helianthus annuus* (sunflower) were aligned to the assembly with diamond (version 0.9.18) and parameters [--id 70 --subject-cover 80]. Diamond hits were filtered to retain only those with greater than 80% identity and alignment length within +/- 5% of the Sunflower protein length. Contigs with at least one filtered Sunflower gene hit, or which have a Kraken2 annotation of Viridiplantae were considered plant contigs. A total of 33 small contigs remained unclassified but no evidence was found of any contaminant contigs. The assembly was screened for organelle contigs via BLAST and with manual review of the alignments, a total of 697 contigs were removed, 622 (9136324 bp) designated chloroplast and 75 (1625390) designated mitochondria.

#### 3. Genome Annotation

Gene models were annotated using the Robust and Extendable eukaryotic Annotation Toolkit (REAT, https://github.com/EI-CoreBioinformatics/reat) and Minos (https://github.com/EI-CoreBioinformatics/minos) using a workflow incorporating repeat identification, RNA-Seq mapping / assembly, alignment of protein sequences from related species and evidence guided gene prediction with AUGUSTUS (see below). The REAT transcriptome workflow was run with RNA-Seq reads and IsoSeq transcripts from 3 tissues (disc, leaf and ray). Illumina RNA-Seq reads were mapped to the genome with HISAT2 v2.1.0 (Kim et al. 2019) and high-confidence splice junctions identified by Portcullis (Kim et al. 2019; Mapleson et al. 2018). The aligned reads were assembled for each tissue with StringTie2 v1.3.3 (Kovaka et al. 2019) and Scallop v0.10.2 (Shao and Kingsford 2017), IsoSeq transcripts were aligned with minimap2 (Li H., 2018). From the combined set of RNA-Seq assemblies and IsoSeq alignments a filtered set of non-redundant gene-models were derived using Mikado (https://github.com/EI-CoreBioinformatics/mikado; (Venturini et al. 2018)). The REAT homology workflow was used to generate gene models based on alignment of protein sequences from 11 Asteraceae species. These together with the transcriptome derived models were used to train the AUGUSTUS (Stanke and Morgenstern 2005), gene predictor, with transcript and protein alignments plus repeat annotation provided as hints in evidence guided gene prediction using the REAT prediction workflow. Three alternative AUGUSTUS gene builds were generated using different evidence inputs or weightings. These together with the gene models derived from the transcriptome and homology workflow were consolidated into a single set of gene models using Minos (https://github.com/EI-CoreBioinformatics/minos). The Minos pipeline scores alternative models based on the level of supporting evidence (protein homology, transcriptome data) and gene structure characteristics (e.g., CDS, UTR features) to select a representative gene model and alternative splice variants.

*Reference guided transcriptome reconstruction*

Gene models were derived from the RNA-Seq and IsoSeq transcripts using the REAT transcriptome workflow (https://github.com/EI-CoreBioinformatics/reat). HISAT2 (Kim et al. 2019) was selected as the short read aligner with IsoSeq transcripts aligned with minimap2 (Kim et al. 2019; Li 2018)), maximum intron length was set as 50,000 bp and minimum intron length to 20bp. IsoSeq alignments were required to meet 90% coverage and 95% identity. High-confidence splice junctions were identified by Portcullis. RNA-Seq reads were assembled for each tissue with StringTie2 (Kovaka et al. 2019) and Scallop (Shao and Kingsford 2017). Gene models were derived from the RNA-Seq assemblies and IsoSeq alignments with Mikado (https://github.com/EI-CoreBioinformatics/mikado; (Venturini et al. 2018)). Mikado was run with all Scallop, StringTie and IsoSeq alignments and a second run with only IsoSeq alignments.

**Table 3.** Reference guided transcriptome assembly statistics.

| Stat | CoDisc (StringTie) | CoDisc (scallop) | CoLeaf (StringTie) | CoDisc (scallop) | CoRay (StringTie) | CoRay (scallop) |
|---|---|---|---|---|---|---|
| Number of genes | 96657 | 79669 | 88418 | 79669 | 84618 | 68308 |
| Number of Transcripts | 162235 | 200517 | 141734 | 200517 | 138904 | 171285 |
| Transcripts per gene | 1.68 | 2.52 | 1.6 | 2.52 | 1.64 | 2.51 |
| Number of monoexonic genes | 21899 | 10074 | 23357 | 10074 | 20140 | 8672 |
| Monoexonic transcripts | 23268 | 14285 | 24490 | 14285 | 21229 | 12217 |
| Transcript mean size cDNA (bp) | 1669.59 | 1852.64 | 1743.52 | 1852.64 | 1705.18 | 1836.14 |
| Transcript median size cDNA (bp) | 1375 | 1593 | 1420 | 1593 | 1395 | 1586 |
| Min cDNA | 200 | 200 | 200 | 200 | 200 | 200 |
| Max cDNA | 29526 | 32610 | 25392 | 32610 | 36617 | 38844 |
| Total exons | 871337 | 1087218 | 763945 | 1087218 | 764548 | 958715 |
| Exons per transcript | 5.37 | 5.42 | 5.39 | 5.42 | 5.5 | 5.6 |
| Exon mean size (bp) | 310.86 | 341.69 | 323.47 | 341.69 | 309.8 | 328.05 |
| Intron mean size (bp) | 429.22 | 399.35 | 403.29 | 399.35 | 410.56 | 383.84 |

**Table 4.** Mikado consolidated gene sets, gene model statistics

| Stat | mikado (only IsoSeq) | mikado (IsoSeq + StringTie + scallop) |
|---|---|---|
| Number of genes | 40650 | 85954 |
| Number of Transcripts | 49415 | 144680 |
| Transcripts per gene | 1.22 | 1.68 |
| Number of monoexonic genes | 4664 | 10169 |

| | | |
|---|---|---|
| Monoexonic transcripts | 4683 | 10256 |
| Transcript mean size cDNA (bp) | 2004.98 | 1753.3 |
| Transcript median size cDNA (bp) | 1759 | 1539 |
| Min cDNA | 305 | 200 |
| Max cDNA | 12107 | 21329 |
| Total exons | 356703 | 821530 |
| Exons per transcript | 7.22 | 5.68 |
| Exon mean size (bp) | 277.76 | 308.77 |
| CDS mean size (bp) | 219 | 214.49 |
| Transcript mean size CDS (bp) | 1434.73 | 982.76 |
| Transcript median size CDS (bp) | 1236 | 801 |
| Min CDS | 0 | 0 |
| Max CDS | 11646 | 16149 |
| Intron mean size (bp) | 275.22 | 363.17 |
| 5'UTR mean size (bp) | 249.73 | 313.01 |
| 3'UTR mean size (bp) | 285.07 | 356.74 |

*Cross-species protein alignment*

Protein sequences from 5 Asteraceae species *Cynara cardunculus* (cardoon), *Mikania micrantha,* (bitter vine), *Lactuca sativa* (lettuce), *Helianthus annuus* (sunflower) and *Artemisia annua* (sweet wormwood) were aligned to the *C. officinalis* assembly using the reat homology workflow (https://github.com/EI-CoreBioinformatics/reat) with options --annotation_filters aa_len --alignment_species Eudicoty --filter_max_intron 20000 --filter_min_exon 10 --alignment_filters aa_len internal_stop intron_len exon_len splicing --alignment_min_coverage 90 --junction_f1_filter 40. The reat homology workflow aligns proteins with spaln v2.4.6 (Gotoh 2021) and filters and generates metrics to remove misaligned proteins. The aligned proteins are clustered into loci and a consolidated set of gene models are derived via Mikado. A second reat homology run was also performed with predicted protein sequences derived from *de novo* transcriptome assemblies and IsoSeq transcripts of *C. arvensis*. Illumina RNA-Seq reads were assembled with trinity v2.8.5, combined with IsoSeq transcripts and filtered and clustered with the tr2aacds pipeline (tr2aacds4.pl, -MINAA=90) of the EvidentialGene package. The filtered set of transcripts (tr2aacds.pl okayset) were further filtered to retain only transcripts with greater than 0.3 TPM as determined via Salmon quantification against the illumina RNA-Seq reads for each species. The CDS sequences of the retained transcripts were translated, and the protein sequences used as input to the reat homology workflow, aligning the proteins to the *C.officinalis* assembly and generating a consolidated set of protein derived gene models.

*Evidence guided gene prediction*

The evidence guided annotation of protein coding genes based on repeats, RNA-Seq mappings, transcript assembly and alignment of protein sequences was created using the reat prediction workflow. The pipeline has four main steps: (1) the reat transcriptome and homology Mikado models are categorised based on alignments to uniprot proteins to identify models with likely full-length CDS and which meet basic structural checks i.e. having complete but not excessively long UTR's and not exceeding a minimum CDS/cDNA ratio. A subset of gene models were then selected from the classified models and used to train the AUGUSTUS gene predictor (Stanke et al., 2005); (2) Augustus is run in both *ab initio* mode and with extrinsic evidence generated in the reat transcriptome and homology runs (repeats, protein alignments, RNA-Seq alignments, splice junctions, categorised Mikado models). Three evidence guided AUGUSTUS predictions were created using alternative bonus scores and priority based on evidence type. (3). AUGUSTUS models, reat transcriptome / homology models, protein and transcriptome alignments were provided to EVidenceModeler (EVM) (Haas et al. 2008) to generate consensus gene structures. (4) EVM models are processed through Mikado to add UTR features and splice variants.

*Gene model consolidation*

The final set of gene models was selected using Minos (https://github.com/EI-CoreBioinformatics/minos)**.** Minos is a pipeline that generates and utilises metrics derived from protein, transcript, and expression data sets to create a consolidated set of gene models. In this annotation, the three alternative evidence guided Augustus gene builds described earlier, and the gene models derived from the reat transcriptome and homology runs were filtered and consolidated into a single set of gene models.

Gene models were classified as biotypes protein_coding_gene, predicted_gene and transposable_element_gene, and assigned as high or low confidence based on the criteria below:

a) **High confidence protein_coding_gene:** Any protein coding gene where any of its associated gene models have a BUSCO v4.0.6 (Seppey, Manni, and Zdobnov 2019)protein status of Complete/Duplicated OR have blastp (v2.9.0+) coverage (average across query and target coverage) >= 80% against the list protein datasets mentioned in Section 4. Or alternatively have average blastp coverage (across query and target coverage) >= 60% against the list protein datasets AND have transcript alignment F1 score (average across nucleotide, exon and junction F1 scores based on RNA-Seq transcript assemblies) >= 40%.

b) **Low confidence protein_coding_gene**: Any protein coding gene where all of its associated transcript models do not meet the criteria to be considered as high confidence protein coding transcripts.

c) **High confidence transposable_element_gene**: Any protein coding gene where any of its associated gene models have coverage >= 40% against the combined interspersed repeats (see methods).

d) **Low confidence transposable_element_gene**: Any protein coding gene where all of its associated transcript models do not meet the criteria to be considered as high confidence and assigned as a transposable_element_gene (see c).

e) **Low confidence predicted_gene**: Any protein coding gene where all of its associated transcript models do not meet the criteria to be considered as high confidence protein coding transcripts. And in addition where any of the associated gene models have average blastp coverage (across query and target coverage) < 30% against the list protein datasets mentioned in Section 4 AND having a protein-coding potential score < 0.25 calculated using CPC2 0.1 (Kong et al. 2007).

f) **Discarded models**: Any models having no BUSCO protein hit AND no protein alignment score (average across nucleotide, exon and junction F1 scores based on protein alignments) AND no transcript alignment F1 score (average across nucleotide, exon and junction F1 scores based on RNA-Seq transcript assemblies) AND no blastp coverage (average across query and target coverage) AND Kallisto v0.44 (Bray et al. 2016) expression score <0.3 from across RNA-Seq reads OR having short CDS <30bps.

**Table 5.** Minos classified gene models

| **Gene Model Classification** | | | |
|---|---|---|---|
| **Biotype** | **Confidence** | **Gene** | **Transcript** |
| protein_coding_gene | High | 72324 | 110138 |
| transposable_element_gene | High | 14620 | 14900 |
| protein_coding_gene | Low | 9858 | 10221 |
| transposable_element_gene | Low | 5301 | 5318 |
| predicted_gene | Low | 2352 | 2383 |
| Total | | 104455 | 142960 |

*Functional annotation*

All proteins were annotated using EI-FunAnnot pipeline v1.2 (https://github.com/EI-CoreBioinformatics/eifunannot) utilising AHRD v.3.3.3

https://hdl.handle.net/20.500.11811/6420; https://github.com/groupschoof/AHRD/blob/master/README.textile). Sequences were blasted against the reference proteins (Arabidopsis thaliana TAIR10, TAIR10_pep_20101214_updated.fasta.gz - https://www.araport.org) and the UniProt viridiplantae sequences (data download 20Dec2021), both Swiss-Prot and TrEMBL datasets (The UniProt Consortium, 2014). Proteins were BLASTed (v2.6.0; blastp) with an e-value of 1e-5. We have also provided interproscan (v5.22.61) (Jones et al. 2014) results to AHRD. We adapted the standard AHRD example configuration file path test/resources/ahrd_example_input_go_prediction.yml, distributed with the AHRD tool, changing the following apart from the location of input and output files:

1. we included the GOA mapping from uniprot (ftp://ftp.ebi.ac.uk/pub/databases/GO/goa/UNIPROT/goa_uniprot_all.gaf.gz) as parameter 'gene_ontology_result',

2. we also included the interpro database (ftp://ftp.ebi.ac.uk/pub/databases/interpro/61.0/interpro.xml.gz) and provided as parameter 'interpro_database',

3. we changed the parameter 'prefer_reference_with_go_annos' to 'false' and did not use the parameter 'gene_ontology_result'

4. The regular expression used to analyse the reference protein fasta header was amended to the custom header format:

fasta_header_regex: "^>(?<accession>[aA][tT][0-9mMcC][gG]\\d+(\\.\\d+)?)\\s+\\|[^\\|]+\\|\\s+(?<description>[^\\|]+)(\\s*\\|.*)?$"

Kovaka, Sam, Aleksey V. Zimin, Geo M. Pertea, Roham Razaghi, Steven L. Salzberg, and

Mihaela Pertea. 2019. "Transcriptome Assembly from Long-Read RNA-Seq Alignments with StringTie2." *Genome Biology* 20 (1): 278.

Li, Heng. 2018. "Minimap2: Pairwise Alignment for Nucleotide Sequences." *Bioinformatics* 34 (18): 3094–3100.

Mapleson, Daniel, Gonzalo Garcia Accinelli, George Kettleborough, Jonathan Wright, and Bernardo J. Clavijo. 2017. "KAT: A K-Mer Analysis Toolkit to Quality Control NGS Datasets and Genome Assemblies." *Bioinformatics (Oxford, England)* 33 (4): 574–76.

Mapleson, Daniel, Luca Venturini, Gemy Kaithakottil, and David Swarbreck. 2018. "Efficient and Accurate Detection of Splice Junctions from RNA-Seq with Portcullis." *GigaScience* 7 (12). https://doi.org/10.1093/gigascience/giy131.

Nurk, Sergey, Brian P. Walenz, Arang Rhie, Mitchell R. Vollger, Glennis A. Logsdon, Robert Grothe, Karen H. Miga, Evan E. Eichler, Adam M. Phillippy, and Sergey Koren. 2020. "HiCanu: Accurate Assembly of Segmental Duplications, Satellites, and Allelic Variants from High-Fidelity Long Reads." *Genome Research* 30 (9): 1291–1305.

Putnam, Nicholas H., Brendan L. O'Connell, Jonathan C. Stites, Brandon J. Rice, Marco Blanchette, Robert Calef, Christopher J. Troll, et al. 2016. "Chromosome-Scale Shotgun Assembly Using an in Vitro Method for Long-Range Linkage." *Genome Research* 26 (3): 342–50.

Ruan, Jue, and Heng Li. 2020. "Fast and Accurate Long-Read Assembly with wtdbg2." *Nature Methods* 17 (2): 155–58.

Seppey, Mathieu, Mosè Manni, and Evgeny M. Zdobnov. 2019. "BUSCO: Assessing Genome Assembly and Annotation Completeness." *Methods in Molecular Biology (Clifton, N.J.)* 1962:227–45.

Shao, Mingfu, and Carl Kingsford. 2017. "Accurate Assembly of Transcripts through Phase-Preserving Graph Decomposition." *Nature Biotechnology* 35 (12): 1167–69.

Simão, Felipe A., Robert M. Waterhouse, Panagiotis Ioannidis, Evgenia V. Kriventseva, and Evgeny M. Zdobnov. 2015. "BUSCO: Assessing Genome Assembly and Annotation Completeness with Single-Copy Orthologs." *Bioinformatics (Oxford, England)* 31 (19): 3210–12.

Stanke, Mario, and Burkhard Morgenstern. 2005. "AUGUSTUS: A Web Server for Gene Prediction in Eukaryotes That Allows User-Defined Constraints." *Nucleic Acids Research* 33 (Web Server issue): W465–67.

Venturini, Luca, Shabhonam Caim, Gemy George Kaithakottil, Daniel Lee Mapleson, and David Swarbreck. 2018. "Leveraging Multiple Transcriptome Assembly Methods for Improved Gene Structure Annotation." *GigaScience* 7 (8). https://doi.org/10.1093/gigascience/giy093.

Wood, Derrick E., Jennifer Lu, and Ben Langmead. 2019. "Improved Metagenomic Analysis with Kraken 2." *Genome Biology* 20 (1): 257.

Yeo, Sarah, Lauren Coombe, René L. Warren, Justin Chu, and Inanç Birol. 2018. "ARCS: Scaffolding Genome Drafts with Linked Reads." *Bioinformatics (Oxford, England)* 34 (5): 725–31.
